# PROFET Predicts Continuous Gene Expression Dynamics from scRNA-seq Data to Elucidate Heterogeneity of Cancer Treatment Responses

**DOI:** 10.1101/2025.06.27.662030

**Authors:** Yu-Chen Cheng, Hyemin Gu, Thomas O. McDonald, Wenbo Wu, Shubham Tripathi, Cristina Guarducci, Douglas Russo, Daniel L. Abravanel, Madeline Bailey, Yue Wang, Yun Zhang, Yannis Pantazis, Herbert Levine, Rinath Jeselsohn, Markos A. Katsoulakis, Franziska Michor

## Abstract

Single-cell RNA sequencing profiles cellular heterogeneity but captures only static snapshots, limiting inference of gene expression dynamics. We developed PROFET (Particle-based Reconstruction Of generative Force-matched Expression Trajectories), a framework that reconstructs continuous, nonlinear single-cell trajectories from sparsely sampled scRNA-seq time series. PROFET combines a particle-based gradient-flow algorithm with simulation-free force matching to accurately infer cellular dynamics. Across mouse and human *in vitro* datasets and an *in vivo* axolotl regeneration dataset, PROFET achieved 2.6–12.5× lower prediction error than ten state-of-the-art trajectory inference methods. Applying PROFET to newly generated scRNA-seq data from a palbociclib-treated MCF7 cell line and three published breast cancer patient datasets, we reconstructed treatment-response trajectories and identified a resistant cell subpopulation exhibiting large phenotypic shifts and enrichment of the surface markers *UNC5B*, *TLR3*, *PCDH19*, *PROCR*, *SLITRK6*, and *SEMA6B*. PROFET provides a biologically grounded framework for reconstructing cell-state dynamics from static single-cell data across development, regeneration, and therapeutic response.

## Introduction

Modern live single-cell imaging technologies, such as those using fluorescently labeled reporter proteins, offer high temporal resolution for tracking gene expression dynamics at the single-cell level^1^. Despite this advantage, live-cell experimental approaches are limited by the small number of markers that can be tracked simultaneously, challenges in maintaining system stability, and the complexity of analyzing imaging data over prolonged periods^2^. In contrast, scRNA-seq technologies provide invaluable insights into cellular heterogeneity by capturing genome-wide expression patterns^3,4^, but face challenges as expression patterns can vary over time due to cell state changes, intrinsic gene expression noise and experimental variability. Moreover, scRNA-seq data are inherently cross-sectional, as individual cells must be lysed for sequencing at each time point. These constraints limit the ability to reconstruct true single-cell trajectories from noisy time-course scRNA-seq datasets^5^.

As a result, neither live-cell imaging nor scRNA-seq alone can provide both the high temporal resolution and broad gene expression profiling required to capture dynamic changes in cellular states. To bridge this gap, recent data-driven approaches have leveraged mathematical frameworks from optimal transport (OT)^6–13^ and spline interpolation^14^ to infer cellular trajectories by linking distributions of cell states across discrete time points. The majority of existing methods are grounded in OT theory, spanning a range of formulations: static OT maps (WOT^6^ and CellOT^13^), Schrödinger bridges (DeepRUOT^12^), Sinkhorn-based schemes (MIOFlow^8^, PI-SDE^11^, VGFM^10^), and Benamou–Brenier dynamic OT (TrajectoryNet^7^, TIGON^9^). Others construct trajectories via spline interpolation with OT-derived couplings (MMFM^14^). Many of these formulations employ a quadratic transport cost, leading to the Wasserstein-2 (W2) distance. In the dynamic OT, finding the most cost-efficient path can be viewed as a control-theoretic approach that interpolates between two probability measures by minimizing kinetic energy over the entire path, often augmented with additional constraints to enforce flow consistency or incorporate biological priors ^7,9^. While these approaches have enabled important advances in trajectory inference, W2-based formulations can remain sensitive to complex biological data geometries, including multimodal distributions, branching trajectories, heavy-tailed distributions, and manifold structure ^7,8,15^.

To address these limitations, we developed PROFET (Particle-based Reconstruction Of generative Force-matched Expression Trajectories), a computational framework that formulates trajectory inference as a Wasserstein gradient flow of the Lipschitz-regularized Kullback–Leibler (KL) divergence—mathematically equivalent to a Wasserstein-1 (W1) proximal regularized KL divergence—which enables robust trajectory inference in three respects. First, this divergence imposes no structural or moment conditions on the target distribution, requiring only a finite first-moment condition on the source—a mild assumption that holds in essentially all practical settings^15^. Second, the Lipschitz constraint on the dual potential ensures that the induced velocity field has uniformly bounded norm, yielding finite-speed dynamics that guarantee numerical stability under forward Euler discretization. Third, the dissipative structure of the Wasserstein gradient flow admits a greedy sequential implementation—each time step solves a local variational problem independently—rather than requiring global optimization over the entire trajectory, enabling adaptive time horizons and resolution sufficient to resolve transport through complex geometries.

In this study, we systematically benchmark PROFET against ten representative methods— most rooted in optimal transport but differing in training methodology: discretize-then-optimize (DTO) (MIOFlow^8^, PRESCIENT^16^, PI-SDE^11^, DeepROUT^12^), adjoint-based (TrajectoryNet^7^, TIGON^9^), simulation-free (MMFM^14^, VGFM^10^), and static transport maps(WOT^6^, CellOT^13^). Using *in vitro* and *in vivo* mouse, axolotl, and human datasets, we demonstrate that PROFET consistently outperforms competing methods across varying levels of biological complexity while remaining data-agnostic and scalable. Across all test datasets, PROFET achieved 2.6–12.5-fold lower W2 error than competing methods, based on the geometric mean of W2 error ratios, while maintaining comparable runtime and memory requirements.

For downstream analyses, we then applied PROFET to a scRNA-seq dataset from the MCF7 hormone receptor positive (HR+) breast cancer cell line^17^ as well as three datasets from HR+ metastatic breast cancer patients^18,19^ to reconstruct single-cell trajectories spanning pre- and post-palbociclib time points. Palbociclib, a CDK4/6 inhibitor, has extended progression-free survival in HR+ breast cancer; however, acquired resistance to CDK4/6 inhibition has been widely reported in both clinical and preclinical studies^20,21^. In this study of the MCF7 cell line, 16 weeks of palbociclib treatment led to experimentally confirmed drug resistance, accompanied by marked downregulation of cell cycle–related genes. To compare this response with that obtained from patient-derived data, we identified a subpopulation of cells in the patient samples that exhibited similarly pronounced phenotypic shifts, characterized by the same downregulation signature of cell cycle genes. Differential gene expression (DEG) analysis revealed that six surface markers— *UNC5B*, *TLR3*, *PCDH19*, *PROCR*, *SLITRK6*, and *SEMA6B* —were enriched in the pre-treatment states of these high-shift cells. These markers may serve as early predictors of phenotypic transitions in response to palbociclib treatment and potential targets for antibody-drug conjugates (ADCs)^22^. More broadly, PROFET provides a generalizable approach for reconstructing gene expression trajectories from time-series scRNA-seq data across phenotypically heterogeneous cell populations.

## Results

### Overview of PROFET

To reconstruct complete trajectories from discrete scRNA-seq measurements, PROFET employs a two-step framework to infer the underlying temporal dynamics using force-based generative models whose output a deep neural network is then trained on (Figure 1A). The first step of PROFET is based on the Generative Particle Algorithm (GPA)^23,24^, which interpolates gene expression patterns by propagating initial cell states (i.e. the source) along the gradient of a potential landscape (Figure 1B and Supplemental Notes 1–2). Unlike OT-based formulations that rely on W2-based interpolation, GPA defines trajectories as a Wasserstein gradient flow of a Lipschitz-regularized KL divergence with respect to the target distribution. Lipschitz-regularized KL divergence is well-defined on a broad class of distributions, imposing no structural or moment conditions on the target — in particular, requiring no finite second moment assumption — and yields a well-defined potential function for the gradient flow^15^. The Lipschitz continuity of the potential induces finite-speed dynamics, with the Lipschitz bound serving as a speed limit, ensuring numerical stability of the forward Euler discretization via the Courant–Friedrichs–Lewy (CFL) condition (Methods, Supplemental Note 2). Furthermore, the dissipative structure of the Wasserstein gradient flow naturally admits a greedy sequential implementation, where each time step solves an independent local variational problem. This enables adaptive time horizons (see Methods, “Stopping time for GPA”), providing sufficient resolution to capture transport through complex geometries — in contrast to control-theoretic formulations of dynamic OT, where the time horizon and integration resolution are fixed a priori and typically implemented as amortized flow maps. Collectively, these properties endow PROFET with both theoretical stability guarantees and empirical robustness against trajectory collapse or divergence during training.

**Figure 1.**
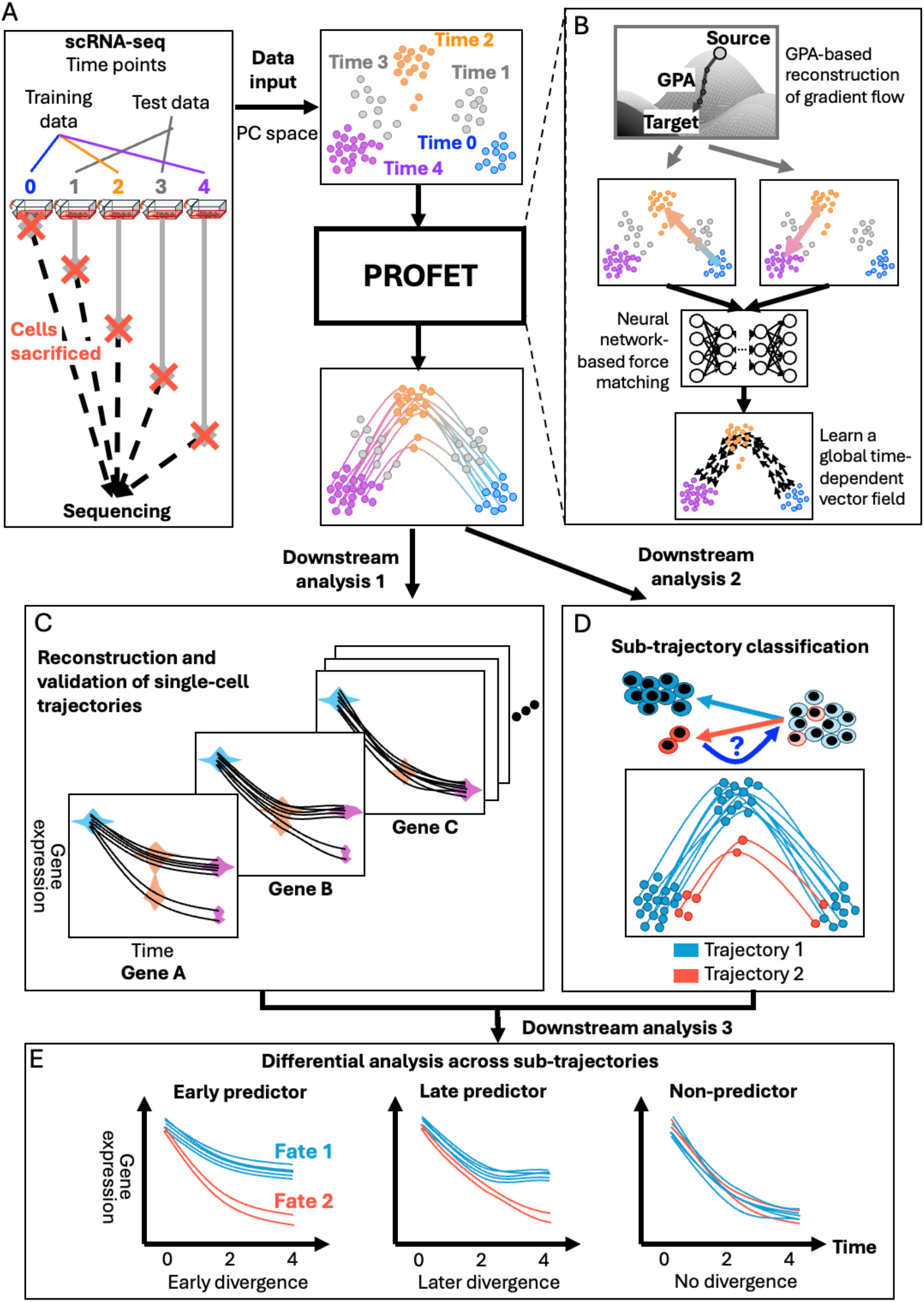
Reconstruction of continuous single-cell gene expression dynamics from discrete scRNA-seq data. (A) Study objective and approach. Left: Existing scRNA-seq technologies capture only static snapshots of cell states, limiting the ability to observe continuous trajectories. Right: To address this limitation, we developed PROFET to reconstruct continuous gene expression dynamics from sampled scRNA-seq time points (colored), while masking intermediate time points (gray) for validation. (B) Two-step architecture of PROFET. Step 1: Apply Generative Particle Algorithm (GPA) to infer gradient flows interpolating between observed distributions at adjacent time points. Step 2: Use a neural network to perform force-matching on these flows and learn a globally smooth, time-dependent vector field that models cell dynamics. (C–E) Three key downstream analyses enabled by our method. (C) Prediction of continuous single-cell gene expression dynamics. (D) Classification of predicted trajectories into terminal fates, enabling tracing fate-assigned cells to their origins. (E) Identification of diverse temporal gene expression patterns associated with distinct fates, illustrated by three representative genes: left panel shows early divergence, middle shows late divergence, and right shows no divergence.

In the second step, the outputs from GPA are used to train a neural network that learns a continuous, time-dependent vector field defined on a Eulerian grid using a force-matching scheme^25^, bridging the trajectory-based (Lagrangian) and field-based (Eulerian) representations of cell dynamics^26^ (Methods and Supplemental Note 3). In terms of training methodology, this step amortizes the piecewise particle trajectories from Step 1 into a globally smooth velocity field in a simulation-free manner. As a result, the complexity of the training signal scales naturally with the data geometry determined in Step 1. This procedure enforces temporal consistency across subintervals and yields a continuous vector field that can be integrated from arbitrary initial conditions. The resulting object is therefore a time-dependent flow map, analogous to those obtained in dynamic OT methods, but derived through a fundamentally different and more robust route. This design is biologically motivated: in many systems, particularly under perturbations (e.g., drug treatment or mutation), the underlying regulatory landscape is non-stationary and evolves over time.

Together, the principled gradient flow formulation and the two-step training pipeline enable robust modeling of complex biological dynamics—including multimodal splitting and merging processes, the emergence and disappearance of cell types, and strong external perturbations such as drug treatments—where trajectories become highly nonlinear and time-dependent, and accurate mode prediction is biologically critical. The amortized velocity field further enables downstream applications such as fate prediction, gene dynamics reconstruction, and phenotypic shift analysis from arbitrary initial conditions.

For PROFET development, we used synthetic data to guide hyperparameter selection, including the convergence criteria for particle dynamics, the Lipschitz constraint, and the dimensionality of the latent space. To this end, we generated synthetic single-cell expression trajectories using a stochastic gene regulatory network model^27^ of the epithelial–mesenchymal transition (EMT), producing temporally sparse observations that mimic time-series scRNA-seq measurements (Methods; Figure S1A). PROFET was trained on a subset of time points and evaluated by comparing the reconstructed to the withheld intermediate states, allowing us to identify stable parameter ranges that produced robust and non-divergent trajectories (Figure S1B and Supplemental Note 4). The synthetic data were used exclusively for hyperparameter optimization. For each biological dataset analyzed subsequently, PROFET was trained directly on the time-series scRNA-seq data using a subset of time points to learn temporal dynamics and reconstruct withheld intermediate states (Figure S1C).

Building on this foundation, we performed three key downstream analyses to interpret PROFET outputs in biologically meaningful ways (Methods). First, to reconstruct single-gene dynamics, we applied an inverse transformation to map the inferred single-cell trajectories—originally reconstructed in a low-dimensional principal component (PC) space—back to the full gene expression space defined by all input genes. This step enables recovery of continuous gene expression dynamics for every gene in every individual cell (Figure 1C). Second, we implemented two strategies to classify predicted trajectories. The first assigns trajectories to predefined cell subpopulations based on clustering cells at a specific reference time point, allowing interpretation in terms of fate or origin (Figure 1D illustrates this classification applied to terminal fates). The second strategy quantifies the magnitude of cell state transitions across time by computing trajectory-wise Euclidean distances, which are then thresholded into distinct transition groups. Finally, by integrating the above classifications, we performed time-resolved differential gene expression analysis across sub-trajectories. This approach enabled us to identify not only which marker genes distinguish different trajectories, but also when these genes begin to diverge. These analyses uncover temporal transcriptional programs that underlie divergent treatment responses (Figure 1E).

### Model Validation and Benchmarking Across Four Biological Datasets

For validation and benchmarking, we compared PROFET against ten representative trajectory inference methods broadly rooted in optimal transport realized through different mathematical models and categorized into four distinct training methodologies — static transport maps, adjoint-based (neural ODE), discretize-then-optimize (DTO), and simulation-free (flow-matching) — each with different individual features. WOT^6^ and CellOT^13^ compute static transport maps between pairs of timepoints where WOT employs unbalancedness. TrajectoryNet^7^ and TIGON^9^ are adjoint-based methods that learn continuous dynamics via neural ODEs with backpropagation through trajectories, incorporating unbalancedness. Among DTO methods, MIOFlow^8^ learns nonlinear data manifolds, PRESCIENT^16^ employs a probabilistic energy-based formulation with unbalancedness, DeepROUT^12^ combines unbalancedness with stochasticity linking to the Schrödinger bridge, and PI-SDE^11^ uses physics-informed SDE formulations. MMFM^14^ and VGFM^10^ adopt simulation-free flow-matching, with VGFM additionally incorporating unbalancedness. A comprehensive comparison of these methods, including their mathematical frameworks, training strategies, and methodological features, is provided in Tables S1–S2, while the methodological novelties of PROFET are discussed in detail in Supplemental Note 5. We also included a null model, defined as random coupling between cell states across time points, which serves as a baseline without learned temporal structure.

We systematically benchmarked PROFET against these methods across four biological datasets spanning diverse biological systems and a broad range of nonlinear biological dynamics, including *in vitro* datasets profiling mouse embryonic stem cell (mESC) differentiation^28,29^ and TGF-β–induced EMT in the human MCF10A cell line^30^, as well as *in vivo* Axolotl limb regeneration data^31^ and the LARRY mouse hematopoiesis dataset^32^. To ensure consistent evaluation, the same training–testing strategy was applied across datasets by withholding intermediate time points during training and evaluating each method based on its ability to reconstruct the held-out cellular distributions. The mESC dataset contains five time points, with 0 h, 24 h, and 48 h used for training and 12 h and 36 h withheld for testing. This dataset exhibits a curved, detour-like trajectory in gene-expression space rather than a direct linear progression. The EMT dataset contains three time points, with day 0 and day 4 used for training and day 2 withheld for testing. It exhibits redistribution of cell states, transitioning from a bimodal structure at day 0 to a unimodal structure at day 4. The axolotl dataset contains five time points, with days 0, 7, and 22 used for training and days 3 and 14 withheld for testing. This dataset displays oscillatory dynamics, with the number of modes increasing from day 0 to day 7 and subsequently decreasing from day 7 to day 22. Finally, the LARRY dataset contains three time points, with days 2 and 6 used for training and day 4 withheld for testing. It captures mouse hematopoiesis, in which hematopoietic progenitor cells progressively diverge into multiple mature blood-cell lineages over time (Figure 2A).

**Figure 2.**
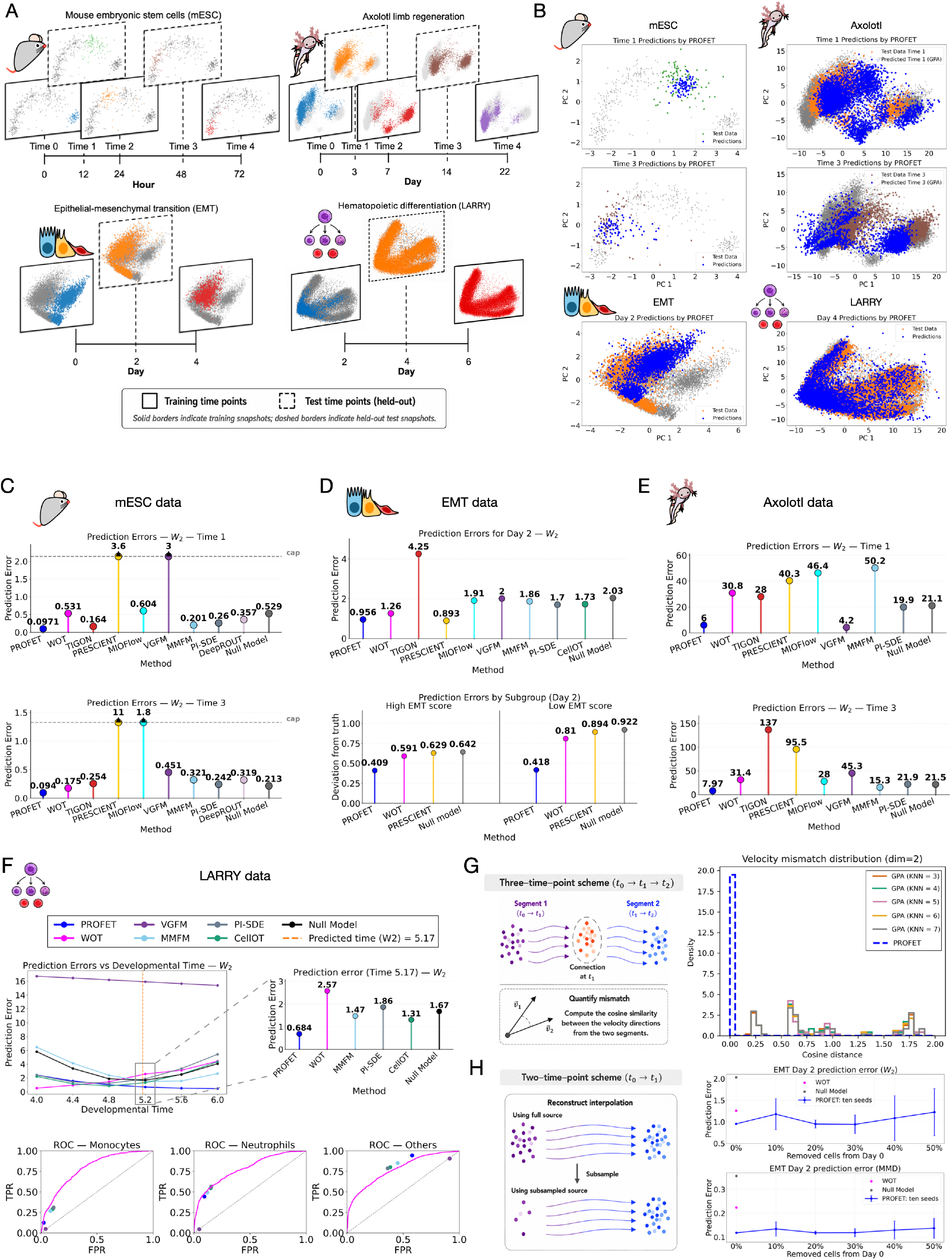
Validation and Benchmarking of PROFET. (A) Overview of the four time-series scRNA-seq datasets, indicating the time points used for model training and those reserved for testing. (B) Principal component (PC) plots of the mESC, EMT, axolotl, and LARRY scRNA-seq datasets comparing PROFET predictions with the experimentally observed ground-truth cell-state distributions at the held-out test time points. (C) mESC dataset: Quantitative prediction errors measured as Wasserstein-2 (W2) distances between the predicted cell-state distributions and the experimentally observed ground-truth distributions at time 1 (upper row) and time 3 (lower row). (D) EMT dataset: The upper row shows quantitative prediction errors between the predicted and experimentally observed cell-state distributions at the held-out intermediate time point (day 2), measured using the W2 metric. The lower row quantifies the absolute deviations in the predicted proportions of the two EMT subpopulations relative to the ground truth at day 2, corresponding to overestimation of the high-EMT subgroup and underestimation of the low-EMT subgroup. (E) Axolotl dataset: Quantitative prediction errors measured as Wasserstein-2 (W2) distances between the predicted cell-state distributions and the experimentally observed ground-truth distributions at time 1 (upper row) and time 3 (lower row). (F) LARRY dataset: The upper row shows prediction errors across developmental times from days 4–6, measured using the W2 metric. The vertical dashed line indicates the predicted developmental time, estimated from the ratio of W2 distances between the day 2–4 and day 4–6 intervals; the inset highlights prediction errors for competing methods at this estimated time point. The lower row shows prediction performance for the three cell fates using receiver operating characteristic (ROC) curves, where TPR (true positive rate) measures the fraction of correctly identified cells belonging to a given fate, and FPR (false positive rate) measures the fraction of incorrectly assigned cells from other fates. (G) Ablation analysis for three-time-point interpolation: The left panel illustrates the computation of vector field mismatch at the connected intermediate time point using cosine distance. The right panel shows the distribution of cosine distances for PROFET and GPA-only reconstructions. For GPA-only, the analysis depends on the choice of K-nearest neighbors (KNN); details are provided in Supplementary Note 7. (H) Ablation analysis for two-time-point interpolation: The left panel illustrates subsampling cells from the source time point to evaluate robustness under reduced training data. The right panel shows prediction performance under source-cell subsampling from 10% to 50%, compared with the baseline WOT and null models trained using the full dataset. To account for stochastic variability in the subsampling procedure, each subsampling level was repeated across ten independent random seeds, and the reported variation reflects this uncertainty.

Across the benchmark datasets, PROFET accurately predicted the held-out test time points (Figure 2B) and consistently achieved strong performance, with 2.6–12.5× lower W2 error than competing methods (geometric mean of error ratios across all datasets and timepoints; Table S3). Although some methods performed competitively on individual datasets or metrics, their performance varied substantially across biological systems (Figure 2C-E). In contrast, PROFET maintained stable performance across all benchmark datasets. In the mESC dataset, PROFET achieved the lowest prediction error at both held-out time points (12 h and 36 h) across all evaluated metrics, including W2 distance, Sinkhorn divergence, and MMD. The second-best method, TIGON, exhibited approximately 1.7-fold higher error, whereas all remaining methods showed more than twofold higher errors relative to PROFET (Figure 2C; Figure S2).

Moving from the mESC dataset to the EMT dataset, the complexity of the underlying dynamics increases, as the cellular distribution transitions from a bimodal to a unimodal structure over time. The number of modes is biologically meaningful: by computing an EMT score for each cell (Methods), we identified two distinct subpopulations (p = 0.0001; Wilcoxon rank-sum test; Figure S3). In this setting, although PRESCIENT achieves comparable W2 error to PROFET (1.06 vs 0.89; Figure 2D, upper panel; Figure S4), it fails to capture the underlying multimodal structure. Specifically, it predicts a single dominant mode, overestimating the abundance of the mesenchymal subpopulation by more than 60% relative to the ground truth while underestimating the epithelial subpopulation by approximately 90%. Similar behavior is observed for other methods, which also show strong bias toward the dominant mode (Figure 2D, lower panel). In contrast, PROFET more accurately reconstructs both populations, with estimated abundances closer to the ground truth (40% overestimation for the mesenchymal subpopulation and 30% underestimation for the epithelial subpopulation; Figure 2D, lower panel). Notably, at the held-out intermediate time point, PROFET retained approximately 70% of the epithelial subpopulation, whereas all competing methods retained only ∼10%, representing an approximately sevenfold improvement in recovering the transient minority population.

In the axolotl dataset—the most complex system analyzed here, featuring oscillatory dynamics with both divergence and convergence of cell states (Figure S5)—PROFET outperforms all methods except VGFM. Specifically, PROFET achieves W2 errors approximately 3- to 8-fold lower than competing models at both withheld time points (Time 1, day 7; and Time 3, day 14) across all distributional metrics (Figure 2E and Figure S5). Although VGFM shows comparable performance at Time 1 (0.9-fold relative to PROFET), its error grows to approximately 5-fold that of PROFET at Time 3. This degradation likely arises from VGFM’s tendency to evolve too slowly, preventing convergence to the target distribution within the required temporal window (Figure S5).

Furthermore, we performed two evaluations using the LARRY dataset. First, for interpolation benchmarking, we leveraged the three available time points (days 2, 4, and 6) in the hematopoietic system. Following the same setup as for the EMT dataset, we trained the models using days 2 and 6 and evaluated predictions on the held-out intermediate time point (day 4). Previous studies have shown that cells at days 4 and 6 are already close to mature states, with day 4 approaching a quasi-stationary distribution. To account for this observation, we further estimated an effective developmental time for the held-out distribution by comparing relative distances between distributions (day 2 → day 4 and day 6 → day 4) across multiple metrics, yielding consistent estimates (day 5.17 using W2 and day 5.66 using MMD). Across this broad range of estimated intermediate times (days 5–6), PROFET consistently achieves the lowest prediction error among all methods (Figure 2F, upper panel and Figure S6), demonstrating robustness to uncertainty in temporal alignment. Second, leveraging the lineage barcode information unique to the LARRY dataset, we evaluated the cell-to-cell matching performance of PROFET and the other methods. Following the original study, we assessed each method’s ability to recover progenitor–fate relationships for major hematopoietic lineages (e.g., neutrophils, monocytes and others) using the true positive rate (TPR) and the area under the ROC curve (AUC) metrics (Methods). Although lineage matching is not the primary objective of PROFET, we found that it achieves a performance comparable to established approaches such as WOT, as well as recent methods including CellOT, VGFM, and MMFM (Figure 2F, lower panel). These results indicate that, despite not being explicitly designed for lineage reconstruction, PROFET preserves biologically meaningful relationships between progenitor and descendant cells.

This analysis establishes that alternative methods exhibit clear limitations when applied to temporally sparse and highly nonlinear biological datasets. In particular, MMFM, WOT, TIGON, and PRESCIENT underestimate the number of cellular subpopulations present at transient states, such as day 2 in the EMT dataset. Capturing these cellular subpopulations is biologically essential, as they reflect heterogeneous cell identities and state transitions. In addition, VGFM and PRESCIENT show deficiencies in aligning learned dynamics with the underlying temporal scale. VGFM tends to evolve too slowly, often failing to reach even the first intermediate time point and not converging to the target distribution, whereas PRESCIENT exhibits the opposite behavior, overshooting the target distribution (e.g., in the mESC dataset, Figure S2). These mismatches indicate limitations of these methods in accurately modeling the temporal progression of cellular dynamics. Furthermore, PI-SDE, DeepROUT, and MIOFlow perform poorly on sparse and highly nonlinear datasets, such as mESC and axolotl (Figures 2C and 2E; Figures S2 and S5), frequently producing collapsed or distorted trajectory reconstructions. CellOT, which is designed for pairwise transport between two time points under a W2 optimal transport framework, achieves reasonable performance in two-time-point settings, similar to WOT. However, when evaluated on interpolation tasks involving three-time-point datasets (Figure 2D for EMT and Figure 2F for LARRY), CellOT exhibits lower accuracy in predicting the masked intermediate time point compared to PROFET. Finally, TrajectoryNet, a continuous normalizing flow (CNF)-based model, fails to converge across the benchmark datasets, using both the parameter settings recommended in the original study and additional hyperparameter configurations, and generates implausible trajectories that deviate from the withheld intermediate distributions (Supplemental Note 6).

### Ablation Analysis, Robustness, and Computational Scalability

To further assess the robustness and component-wise contributions of PROFET, we performed systematic ablation and sensitivity analyses. For the ablation analysis, we evaluated the contribution of the neural force-matching step (Step 2) to the GPA component (Step 1) under two data schemes: (i) datasets with more than two observed training time points (e.g., the mESC dataset) and (ii) datasets with only two observed training time points (e.g., the EMT dataset). This distinction is important because the force-matching step is expected to provide different benefits depending on the amount of temporal information available during training.

In multi-time-point settings, the primary role of the neural force-matching step, Step 2, is to ensure temporal consistency across adjacent trajectory segments. Using only the GPA component, Step 1, trajectories are reconstructed independently between neighboring time points, which can result in inconsistencies in the inferred velocity fields at shared intermediate states. To illustrate this effect, we evaluated the mESC dataset, where trajectories are independently reconstructed by GPA between time points 0–2 and 2–4. The intermediate time point (time 2) therefore serves as the connection point between the two GPA-derived trajectory segments. To quantify the degree of inconsistency between these independently reconstructed segments, we compared velocity vectors inferred from the two adjacent segments at this shared time point (Figure 2G, left panel). Because GPA does not provide explicit cell-to-cell correspondences between independently reconstructed segments, we estimated local correspondences using a K-nearest-neighbor (KNN) approach and computed the cosine distance between matched velocity vectors (Supplemental Note 7). We found that Step 2 reduces these velocity mismatches by integrating local GPA-derived flows into a unified continuous-time vector field, lowering cosine distances and improved alignment of the inferred dynamics (Figure 2G, right panel). These results indicate that Step 2 resolves local inconsistencies and produces smoother, more temporally coherent trajectories than using Step 1 alone.

In two-time-point settings, Step 1 alone is sufficient to generate interpolation trajectories, allowing us to isolate the contribution of Step 2 to data efficiency, robustness and generalization. To assess data efficiency, we performed systematic cell-subsampling experiments in which only 10–50% of the source cells were retained while preserving the overall distribution of cell states (Figure 2H, left panel). Under this setting, Step 1 learns trajectories only for the selected subset of cells. Step 2 then learns a global continuous-time vector field from these trajectories, enabling the model to generalize beyond the sampled cells and reconstruct dynamics throughout the broader state space. Across all subsampling levels, the full PROFET framework maintained prediction accuracy comparable to that obtained using the complete dataset, demonstrating reduced sensitivity to sample size and improved robustness under limited-data conditions (Figure 2H, right panel). Each subsampling level was repeated across ten independent random subsamples to account for stochastic variability. Together, these results demonstrate that the force-matching step (Step 2) not only improves temporal consistency but also enhances the stability and generalization of the learned dynamics.

In addition to the ablation analysis, we performed sensitivity analyses across the key hyperparameters of PROFET. Specifically, we varied the GPA stopping criterion, the dimensionality of the latent space, and the Lipschitz constraint parameter. Across both the EMT and mESC datasets, PROFET maintained stable performance over a broad range of parameter settings, indicating that the framework is robust to moderate variations in model configuration (Figure S7). In particular, performance remained stable across Lipschitz constraint values ranging from 1 to 5 in the EMT dataset, whereas only a mild degradation was observed at larger values in the mESC dataset (Figure S7). Overall, these results demonstrate that PROFET is not highly sensitive to hyperparameter selection and retains strong predictive performance across diverse settings. Detailed results for the sensitivity analyses are provided in Figure S7 and Supplemental Note 7.

To assess computational efficiency and scalability, we further evaluated runtime and memory usage across datasets of varying sample sizes and dimensions (Supplemental Note 8). PROFET’s runtime scales moderately with both dimension (487 s to 1,085 s from d=2 to d=16) and sample size (487 s to 2,743 s from n=278 to n=34,317), while peak memory remains constant across dimensions (∼0.36 GB) and grows to 3.95 GB only on the largest dataset — well below methods that retain full computational graphs (DeepRUOT: 7.41 GB, MIOFlow: 6.70 GB, VGFM: 6.83 GB). On the largest benchmark, the LARRY dataset, PROFET was 13× faster than TIGON and 34× faster than TrajectoryNet. Despite taking finer, geometry-adaptive integration steps, PROFET’s runtime remains comparable to discretize-then-optimize (DTO) methods (PRESCIENT, PI-SDE, DeepRUOT) because each forward Euler step is a greedy local computation with no backward pass through the trajectory. Methods that reduce to a single integration step per time interval (MIOFlow, VGFM) or bypass simulation entirely (MMFM) are faster but sacrifice temporal smoothness or modeling flexibility (Table S4).

### Predicting Gene Expression and Differentiation Pathways from Single-Cell Trajectories

PROFET enables reconstruction of continuous single-cell gene expression trajectories for all genes in a multi-time-point single-cell RNAseq dataset. We first set out to analyze the mESC differentiation dataset. To this end, we applied an inverse PCA transformation to project the reconstructed trajectories from the low-dimensional PCA space back to the original high-dimensional gene expression space, defined by all genes in the input data (Methods). By aligning the trajectories with subgroup classifications, PROFET then enables analysis of gene expression differences across distinct subtrajectories (Figure 3A).

**Figure 3.**
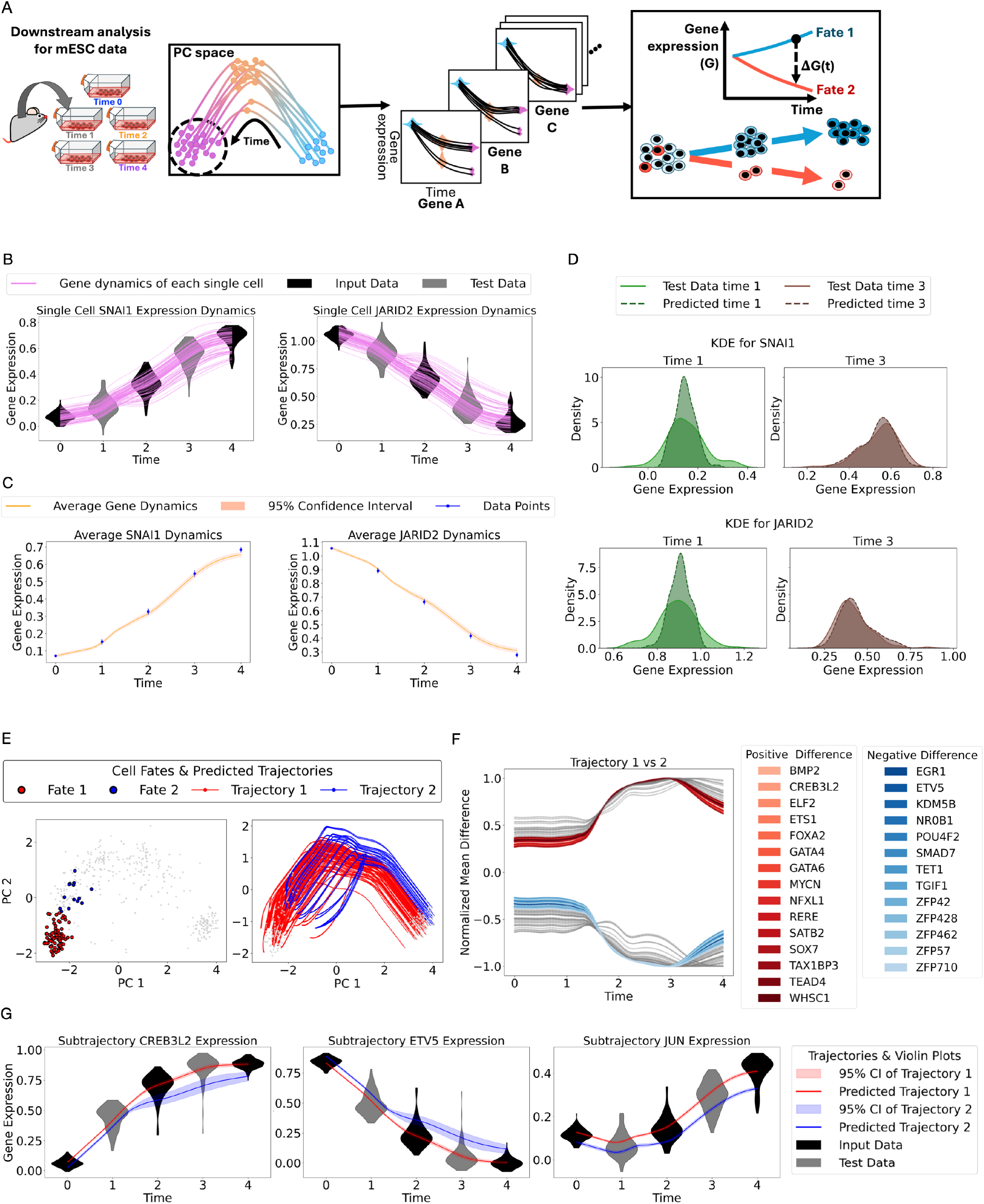
Downstream analysis of reconstructed single-cell trajectories for mESC data. (A) Overview of the downstream analysis pipeline for mESC differentiation. The process begins with PROFET-reconstructed single-cell trajectories in PCA space. These trajectories are then inverse-mapped back to the original gene expression space to recover dynamic gene expression profiles for each gene in each cell. Subsequently, we investigate gene expression differences across distinct sub-trajectories to identify the timing and magnitude of divergence associated with different terminal fates. ΔG(t) denotes the temporal difference in mean gene expression between sub-trajectories. (B) Single-cell, single-gene dynamics plotted as gene expression levels (y-axis) over time (x-axis), with each green curve representing an individual cell trajectory. The predicted trajectories are compared against real data distributions using violin plots (gray for test data, black for training data). The violin plots summarize the distributions of gene expression across all cells at the corresponding time points. (C) Predicted average gene expression trajectories (orange curves with shaded regions indicating 95% confidence intervals across reconstructed cell trajectories) compared to actual gene expression levels from real data (blue dots with error bars). (D) Comparison of predicted and actual gene expression distributions at day 1 (green) and day 3 (brown). Predicted distributions are shown as dashed lines, with real data distributions shown as solid lines. The distributions summarize gene expression across all cells at the corresponding time points. (E) Classification of differentiation fates and ancestral trajectory tracing. Left panel: Two distinct cell fate subgroups at day 4, identified through clustering analysis. Right panel: Ancestral trajectories of the two subgroups traced back to earlier time points. (F) Temporal quantification of gene expression divergence between fate groups. Normalized mean differences are plotted over time, with genes exceeding a divergence threshold of 0.5 highlighted. Each curve represents one gene. Positive values indicate higher expression in trajectory group 1, and negative values indicate higher expression in trajectory group 2. (G) Examples of gene expression divergence in the two subpopulations. Three representative genes are shown, with average gene expression trajectories plotted separately for each subgroup (shaded regions indicate 95% confidence intervals across reconstructed cell trajectories).

By performing this downstream analysis, we recovered the gene expression dynamics for all genes across individual cells in the mESC differentiation dataset. We highlight two representative genes, *JARID2* and *SNAI1*, both of which are known regulators of embryonic development^33,34^ (Figure 3B). Using three time points (time points 0, 2, and 4) for model input, PROFET successfully reconstructs full gene expression trajectories for individual cells, validated by accurate predictions at withheld time points (time points 1 and 3, treated as test data). To quantify prediction accuracy, we compared the mean gene expression along the reconstructed trajectories to the experimental data, finding that, for both test time points 1 and 3, the experimental means (with error bars) consistently fall within the 95% confidence intervals of our predicted means (Figure 3C). We further compared the full distributions of predicted and measured gene expression, observing no statistically significant differences for both genes (one-sided permutation test, *p* > 0.05 and Methods, Figure 3D). Complete results for all other genes are provided in Figures S8–S10 and Table S5.

We next sought to investigate whether single-cell trajectories diverge into distinct cell fates during mESC differentiation. We performed cluster analysis on the final time point of the scRNA-seq dataset (Methods), leading to the identification of two clusters representing different fate subpopulations. Differential gene expression analysis suggested that one trajectory group (fate 1) was enriched for endoderm- or extraembryonic endoderm-associated genes^35^, including *Gata4*, *Gata6*, *Sox7*, *Foxa2*, and *Bmp2*, whereas the other trajectory group (fate 2) retained higher expression of pluripotency- or early stem state-associated genes^36^, such as *Zfp42*/*Rex1*, *Nr0b1*, *Kdm5b*, *Tet1*, and *Zfp57* (Figure 3E left). Leveraging the reconstructed single-cell trajectories, we assigned the final state of each trajectory to one of the two fate clusters by computing its distance to the cluster centroids and selecting the closer one (Methods). This approach enabled us to group trajectories into two fate-specific subpopulations (Figure 3E right).

To identify genes that diverge between the two sub-trajectories, we computed the temporal difference of mean gene expression, normalized by the maximum difference between the groups from day 0 to day 4 (Methods). Genes with a normalized difference greater than 0.5 were classified as divergent (Figure 3F). The continuity of reconstructed trajectories also enabled us to pinpoint when these divergences emerged with high temporal resolution. We identified genes whose expression began to diverge between time points 1 and 2, shedding light on the timing of fate decisions (Figure 3F). Two representative genes, *CREB3L2* and *ETV5*, showed changes exceeding 0.5, in contrast to *JUN*, which exhibited a smaller change of 0.1. Their mean expression profiles across the two trajectory groups (Figure 3G) highlight distinct temporal patterns of gene expression divergence. Complete results for all other genes are provided in Figure S11.

### Predicting Cell-State Convergence in EMT Dynamics

We then performed downstream analyses on the reconstructed trajectories from the EMT dataset. This dataset exhibits cell-state convergence—transitioning from two distinct subpopulations at day 0 to a single population by day 4 (Figure 4A). However, with only discrete time points available in scRNA-seq data, the complete trajectory and timing of this convergence remain unclear. We thus used PROFET to reconstruct the full trajectories of cell states by interpolating between the initial (day 0) and final (day 4) distributions, while withholding the intermediate time point (day 2) for validation (Figure 4B and Figure S12A-D).

**Figure 4.**
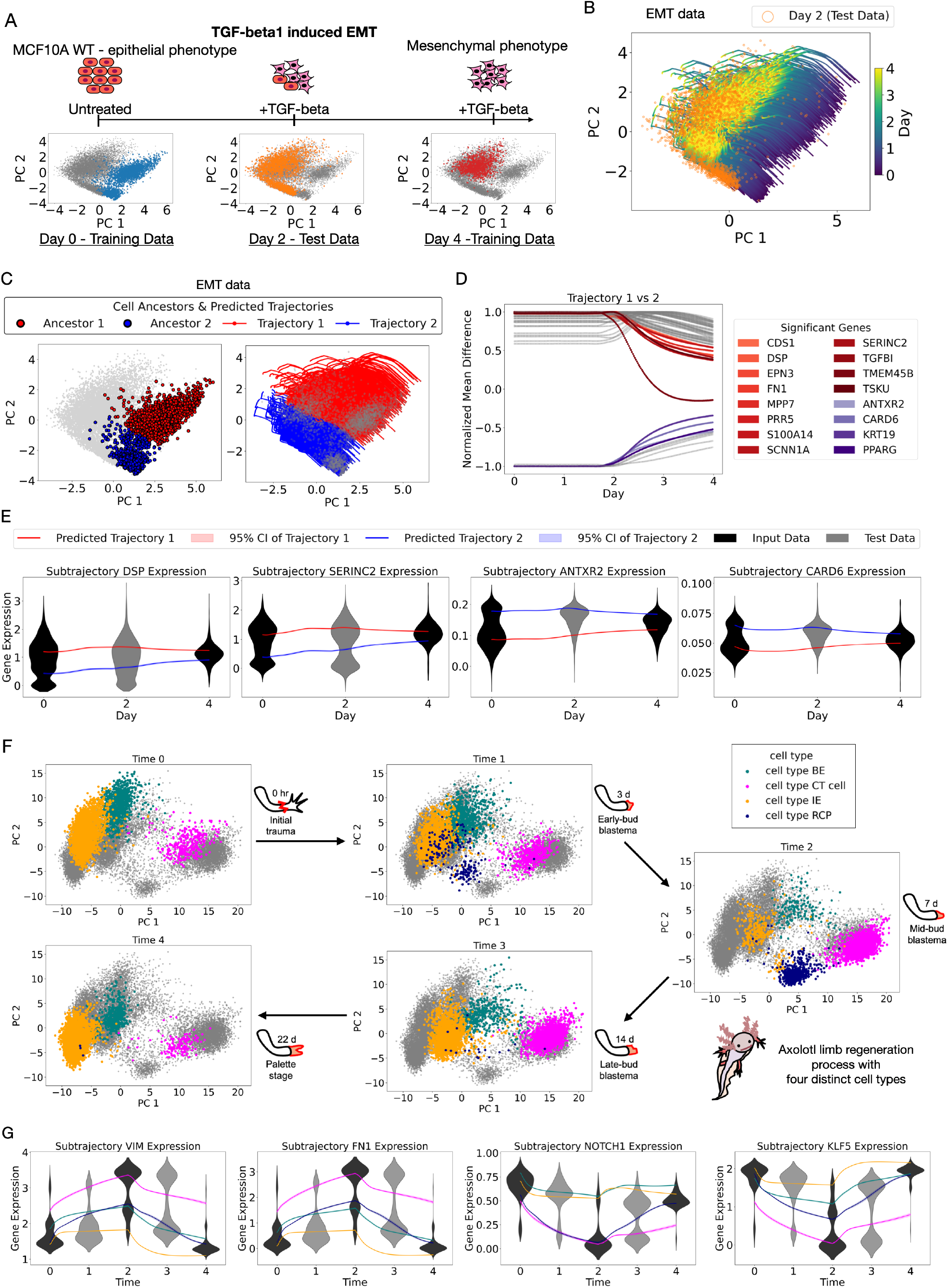
Downstream analysis of reconstructed single-cell trajectories for EMT and Axolotl data. (A) Illustration of EMT scRNA-seq data. (B) Application of PROFET to reconstruct continuous trajectories using only data from time points 0 and 4, with time point 2 held out for validation. The colormap represents the predicted time along the trajectory, ranging from time 0 to time 4, and the test data is highlighted by orange circles. (C) Classification of ancestral cell subgroups and forward trajectory predictions. Left: Two distinct cell subpopulations at time 0, identified via clustering analysis. Right: Forward-predicted trajectories of these two subpopulations. (D) Normalized mean differences in gene expression between the two trajectory groups are plotted over time. Each curve represents one gene. Positive values indicate higher expression in group 1, while negative values indicate higher expression in group 2. Genes exhibiting a drop greater than 0.5 in normalized mean difference at specific time points are highlighted, indicating pronounced convergence in expression levels between the two trajectories. (E) Representative examples of gene expression convergence. Four genes are shown, with average gene expression forward trajectories plotted separately for each ancestral subpopulation. Shaded regions indicate 95% confidence intervals across reconstructed cell trajectories. (F) Illustration of the Axolotl scRNA-seq dataset, highlighting the cell state distributions of four cell types across five time points (days 0, 3, 7, 14, and 22). (G) Representative examples of gene expression divergence. Four genes are shown, with average gene expression trajectories plotted separately for each cell type identified at the mid-bud blastema stage (day 7). Shaded regions indicate 95% confidence intervals across reconstructed cell trajectories. The color of each curve corresponds to the cell-type legend shown in panel (F).

We again applied an inverse PCA transformation to reconstruct continuous single-cell, single-gene trajectories for all genes in the EMT dataset. Analysis of two example genes, *DSP*^37^ and *SERINC2*^38^, showed that their trajectories converged from two distinct modes at day 0 into a single mode by day 4 (Figure S12E). PROFET also captured the intermediate bimodal structure at day 2, which had been withheld as test data (Figure S12F). One-sided permutation tests revealed no statistically significant differences between the predicted and observed distributions on day 2 (*p* > 0.05), supporting the accuracy of PROFET (Methods). Complete results for all other genes are provided in Figures S13–S14 and Table S6.

Unlike the mESC dataset, where trajectory divergence was observed, the EMT dataset reveals evidence of convergence in cellular states. To further explore this observation, we performed clustering analysis on the day 0 data and identified two ancestral subpopulations (Figure 4C left). By classifying each reconstructed trajectory based on its proximity to these ancestral clusters, we traced their respective differentiation paths toward a common final state (Figure 4C right).

To identify genes associated with the convergence process, we calculated the normalized difference in mean gene expression between the two trajectory groups, as described in the previous section. Genes whose expression differences declined by more than 0.5 —representing a reduction of over 50% from their maximal divergence—between day 0 and day 4 were considered strongly associated with convergence (Figure 4D). A key advantage of PROFET, which reconstructs continuous trajectories using force-matching (Methods), is its ability to precisely localize the timing of convergence. We observed that a major reduction in inter-trajectory expression differences occurred after day 2, as illustrated by four representative genes. Each of these genes exhibited a decline of over 50% in mean expression differences between the two sub-trajectories from day 2 to day 4, highlighting their involvement in the convergence process in the later phase of EMT (Figure 4E).

We next set out to investigate temporal phenotypic switching across diverse cell types in the Axolotl dataset. Unlike the *in vitro* EMT dataset, the Axolotl limb regeneration data include four pre-annotated cell types—basal epidermis (BE), intermediate epidermis (IE), connective tissue (CT), and Prdx2⁺ blastema cells (RCP)—originally defined in the source study^31^ (Figure 4F). Divergence among these four cell types is most prominent at the mid-bud blastema stage (time 2, day 7; Figure 4F). However, the full gene-expression dynamics of these cell types, including the early EMT-like phase following injury (from day 0 to day 7) and the later MET-like phase (from day 7 to day 22), were not resolved in the original analysis^31^.

Using PROFET, we reconstructed continuous trajectories across these stages in PC space and applied the same downstream analyses described previously to map the trajectories back to the original gene-expression space, obtaining dynamic expression profiles for every gene. By examining EMT- and stemness-related genes identified from the Molecular Signatures Database (MSigDB)^39^, we predicted complete EMT- and stemness-associated gene dynamics spanning the EMT-to-MET transition (Figure 4G and Figures S15-S16). Four representative genes are shown in Figure 4G: *VIM* and *FN1*, markers of mesenchymal cells, and *NOTCH1* and *KLF5*, associated with stemness. PROFET disentangled their gene-expression trajectories across all four cell types, revealing clear divergence during the EMT phase (from time 0 to 2) and convergence during the MET phase (from time 2 to 4).

### Tracing Phenotypic Shifts Associated with Palbociclib Therapy in HR+ Breast Cancer

We then sought to deploy PROFET to investigate heterogeneity in treatment response to palbociclib, a CDK4/6 inhibitor used in HR+ breast cancer that is known to induce variable treatment responses^19,40^. This analysis was performed across four datasets: one *in vitro* cell line model and three patient-derived samples. For the *in vitro* model, we analyzed scRNA-seq data from an MCF7 breast cancer cell line in which palbociclib resistance was established by escalating drug doses over a 16-week period^17^ (Methods). The dataset includes both parental and resistant cells (Figure 5A).

**Figure 5.**
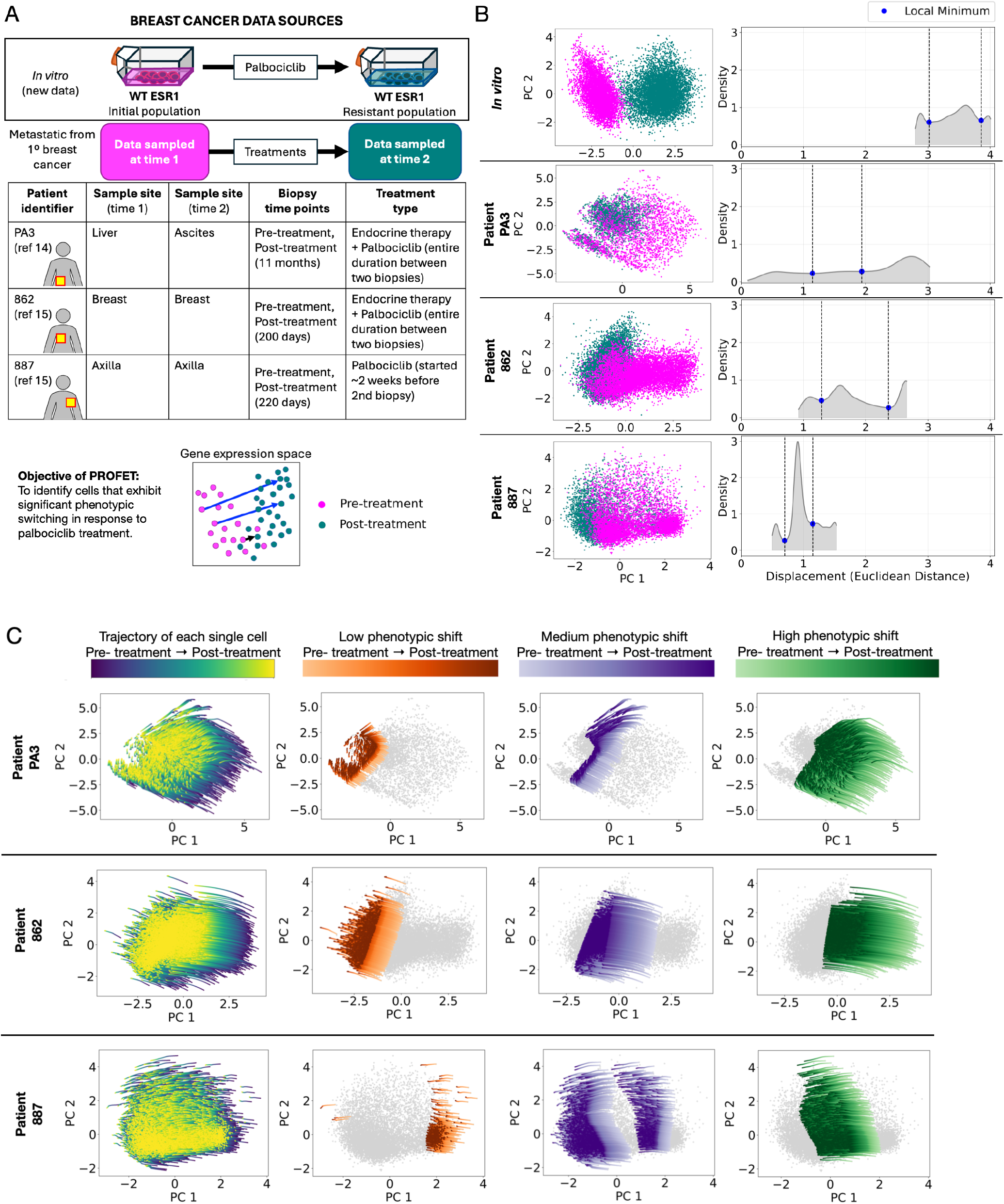
Reconstructed single-cell trajectories for palbociclib resistance. (A) Overview of the datasets analyzed (upper table): one *in vitro* MCF7 breast cancer scRNA-seq dataset (GEO: GSE326012) and three patient-derived datasets—one from Luo *et al.* and two from Klughammer *et al.* The lower panel illustrates the objective of applying PROFET to these datasets: to identify single cells that undergo significant phenotypic switching in response to palbociclib treatment. (B) Each row represents one dataset. Left: PCA projection of scRNA-seq data, with pre-treatment cells in magenta and post-treatment cells in teal. Right: Kernel density estimates (KDE) of phenotypic shifts, defined as the Euclidean distances between initial and final cell states for each single-cell trajectory inferred by PROFET. C) Trajectories of resistance in HR+ breast cancer. Each row corresponds to one dataset. The first panel presents the full set of continuous single-cell trajectories reconstructed by PROFET. The subsequent three panels display subsets of trajectories grouped by their magnitudes of phenotypic shift, quantified as the Euclidean distances between initial and final cell states, as defined in (B).

In parallel, we analyzed longitudinal scRNA-seq data from three HR+ metastatic breast cancer patients. The first patient (PA3) was part of the study by Luo et al^18^. Her initial biopsy was collected prior to palbociclib treatment, and two additional biopsies were both obtained 11 months after initiating palbociclib plus endocrine therapy. These latter samples were collected from ascitic fluid and confirmed disease progression; in our analysis, they were combined as a single post-treatment sample. The second (862) and third (887) patients were part of the study by Klughammer et al^18^. For patient 862, biopsies were collected before and after 200 days of treatment with endocrine therapy plus palbociclib; the second biopsy was performed for research purposes. Patient 887 initially responded to therapy following the first biopsy, but clinical progression was observed prior to the second biopsy, which was collected at day 220. The patient started receiving palbociclib approximately two weeks before the second biopsy (Figure 5A; Methods).

PROFET was developed and validated on datasets with lower dimensionality: a synthetic dataset with 26 genes, the mESC dataset with 100 genes, and the EMT dataset with 72 genes. In contrast, the four palbociclib-treated samples were assayed with genome-wide scRNA-seq, necessitating dimensionality reduction prior to analysis. However, due to high noise levels in these datasets, the first two principal components (PCs) captured only 5–10% of the total variance across samples, making direct PCA-based reduction insufficient.

To address this issue, we adopted a two-step dimensionality reduction strategy. First, we preselected 121 genes previously implicated in palbociclib response, including genes involved in cell cycle regulation, MYC and estrogen receptor (ER) signaling, growth factor pathways, the Hippo pathway, and inflammatory processes^41,42^ (Table S7). In the second step, we applied PCA to the gene expression matrix restricted to this curated gene set. In the resulting PCA space, we observed that the *in vitro* dataset exhibited the most distinct separation between pre- and post-treatment samples while patient-derived datasets showed less clear separation, indicating heterogeneous responses to palbociclib (Figure 5B). To quantify these differences, we leveraged the continuous single-cell trajectories reconstructed by PROFET. For each cell, we computed the extent of phenotypic shift as the Euclidean distance between its inferred pre-treatment and post-treatment states in the gene expression space (Methods). As expected, the *in vitro* dataset exhibited the largest average shift with low variability, while the patient samples displayed broader, more heterogeneous distributions (Figure 5B): the coefficients of variation (CVs) were 0.10 for the *in vitro* data compared to 0.43, 0.30, and 0.26 for patients PA3, 862, and 887, respectively (Methods and Table S8), confirming greater phenotypic variability in the clinical setting.

Based on the relatively homogeneous treatment response observed in the MCF7 *in vitro* dataset—specifically in genes relevant to palbociclib treatment—we used this dataset as a reference to dissect heterogeneity in resistant cell states in patient samples. We focused on identifying subpopulations of patient-derived cells that underwent substantial phenotypic shifts analogous to those observed *in vitro*. To this end, we examined the distributions of phenotypic shift magnitudes and identified three major modes using local minima (Methods and Figure 5B), allowing us to stratify cells in each patient sample into three subgroups: those marked by high, medium, and low phenotypic shifts, respectively. This classification, visualized in PCA space (Figure 5C), supports our hypothesis of heterogeneous treatment responses and enables quantification of which cells undergo large state transitions — an analysis not accessible from unpaired scRNA-seq data alone. This single-cell level classification provides a framework for dissecting phenotypic heterogeneity in response to palbociclib and offers a powerful strategy for identifying candidate cell populations likely to undergo substantial phenotypic shifts in response to treatment.

### Trajectory-Based Dissection of Treatment Response Heterogeneity and Early Biomarker Discovery

Using these trajectory-based subgroups, we next aimed to identify genes with the largest expression changes within the high-shift subgroup, and to further pinpoint surface markers uniquely enriched in pre-treatment cells within this group (Figure 6A). We performed DEG analysis for all four datasets by comparing average gene expression levels between post-treatment samples to those from pre-treatment samples (Methods). This analysis revealed that most cell cycle-related genes were consistently downregulated following treatment across all datasets (Figure 6B). In particular, several members of the centromere protein family—*CENPE*, *CENPF*, *CENPM*, and *CENPV*—were significantly downregulated in the MCF7 cell line following palbociclib treatment (-2.41 < log2 fold change < –1.36; p < 5e–10, two-sided Welch’s t-test), and showed similar downregulation trends in the patient samples (Figure 6B). Complete DEG results are provided in Table S9.

**Figure 6.**
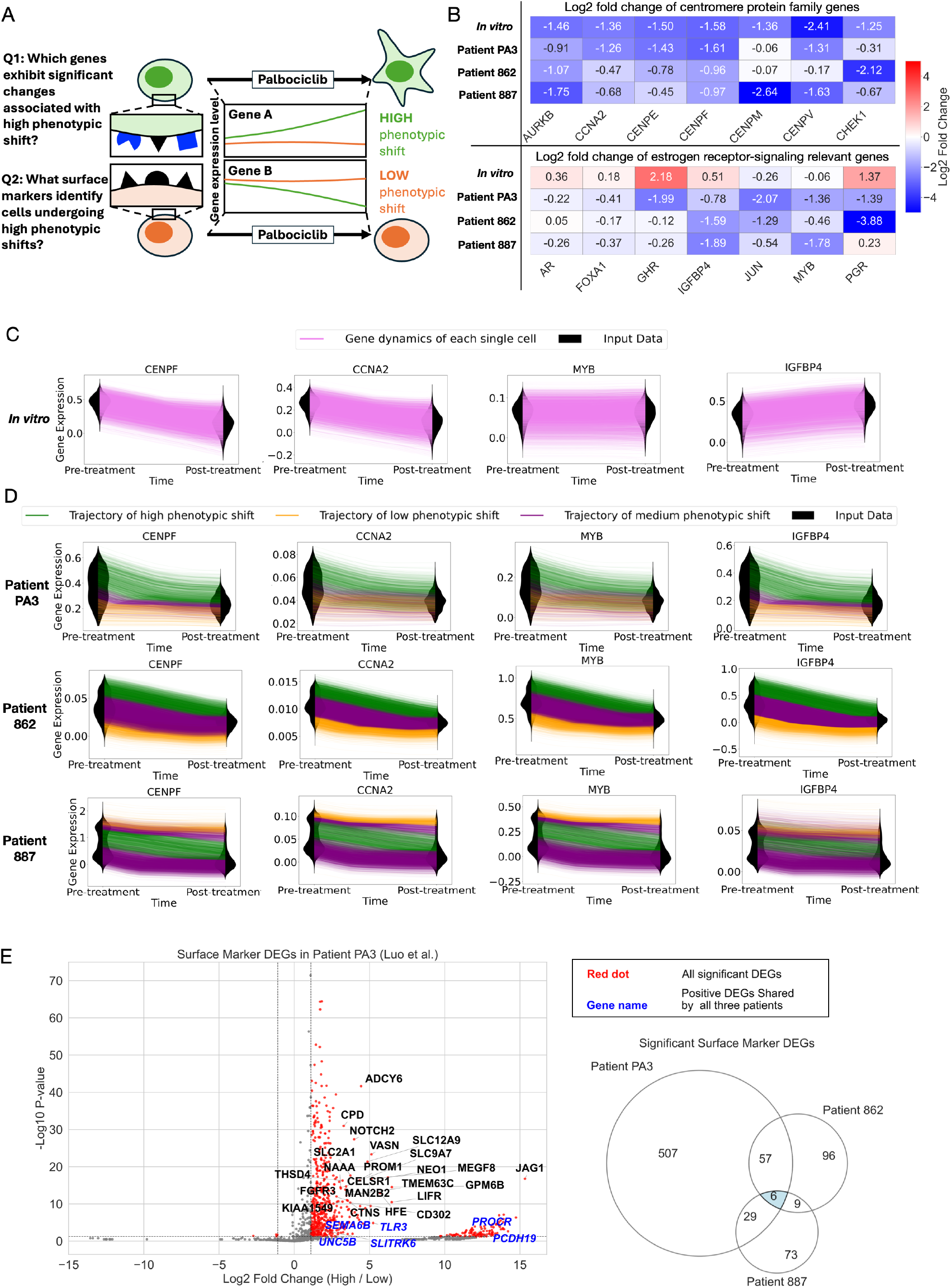
Downstream analysis of reconstructed single-cell trajectories in palbociclib resistance. (A) Schematic overview of the key biological questions addressed through downstream analyses of reconstructed single-cell trajectories that link pre-treatment and post-treatment states. (B) Differential gene expression analysis comparing post-treatment versus pre-treatment time points for each dataset. Each row corresponds to one dataset. Color indicates the direction and magnitude of log₂ fold change (red: upregulated, blue: downregulated). (C) Representative examples of gene expression dynamics for individual cells in the *in vitro* MCF7 breast cancer dataset (GEO: GSE326012). Each magenta curve represents one reconstructed single-cell trajectory. Violin plots show the distributions of observed gene expression across all cells at each time point used as input for trajectory inference. (D) Representative gene dynamics for individual cells across three patient datasets. Violin plots show real expression data; colored lines represent predicted trajectories from three subgroups of cells defined by low, medium, and high phenotypic shift levels, as described in Figure 5. (E) Differential expression analysis of surface marker genes in pre-treatment cells from patient PA3 (Luo et al.), comparing high versus low phenotypic shift subgroups. Statistical significance was assessed using a two-sided Welch’s t-test. Red dots indicate significantly differentially expressed genes (*p* < 0.05, |log₂ fold change| > 1.1). Blue-labeled genes represent significantly upregulated genes (positive log₂ fold change) in the high phenotypic shift subgroup of PA3 and also in the corresponding subpopulations of the other two patients (862 and 887) from Klughammer et al. The accompanying Venn diagram summarizes the overlap of differentially expressed surface-marker genes across all three datasets.

In contrast, we observed divergent expression patterns of estrogen receptor (ER) signaling–relevant genes between the cell line and patient samples PA3 and 862, both of whom received combination therapy with palbociclib and endocrine treatment (Figure 5A). In the cell line data, *AR*, *FOXA1*, *GHR*, *IGFBP4*, and *PGR* were upregulated, whereas these same genes were downregulated in the patient samples. *JUN* and *MYB* were downregulated in both the cell line and patient datasets, with more pronounced downregulation observed in the patient samples (Figure 6B for the full fold change heatmap and tables). These results indicate that the profound downregulation of ER signaling–relevant genes is specific to the patient samples, while centromere genes were consistently downregulated in both the cell line and patient data. The well-established effects of these treatments—palbociclib primarily inhibiting cell cycle–related genes^19,43^, and endocrine therapy downregulating ER signaling–related genes^44,45^— validate our observed trajectories, aligned with the treatment regimens applied to each sample (Figure 5A).

Since this DEG analysis reflects only population-average changes, it does not resolve the specific subgroups within each tumor that exhibit these phenotypic changes induced by the combination therapy. To further dissect this treatment response heterogeneity, we used PROFET to reconstruct gene expression dynamics at the single-cell, single-gene level. Here, we present results for two representative cell cycle–related genes (*CENPF* and *CCNA2*) and two ER signaling–related genes (*MYB* and *IGFBP4*) (Figure 6C-D). Complete results for additional genes are provided in Figures S17–S20. In the *in vitro* dataset, *CENPF* and *CCNA2* showed uniform downregulation, *MYB* expression remained consistently steady, and *IGFBP4* was uniformly upregulated (Figure 6C). In contrast, the patient datasets displayed substantial variability, with the high phenotypic shift subgroup exhibiting more pronounced downregulation of all four genes compared to the low-shift subgroup (Figure 6D).

We then sought to identify early markers of the subpopulation of cells undergoing high phenotypic shifts in response to palbociclib resistance. To identify such markers, we performed DEG analysis of surface marker genes between the high and low phenotypic shift groups at the pre-treatment time point^37^. This analysis revealed six genes— *UNC5B*, *TLR3*, *PCDH19*, *PROCR*, *SLITRK6*, and *SEMA6B* —that were consistently enriched in the high-shift subgroup across all three patient datasets (Figure 6E).

Among these genes, *UNC5B* has been reported to be significantly upregulated in breast cancer and is correlated with poor overall survival in breast cancer patients^46^. Activation of *TLR3* has been shown to increase cancer stem cell (CSC) markers, which are associated with breast cancer progression^47^. *PROCR* is known to activate several signaling pathways in breast cancer cells, including ERK, PI3K–Akt–mTOR, and RhoA–ROCK–p38 pathways^48^. While *PCDH19* has not previously been implicated in metastatic breast cancer, it belongs to the protocadherin family; another member of this family, *PCDHA1*, has been associated with poor prognosis and immune cell infiltration in breast cancer^49^. Although *SEMA6B* and *SLITRK6* have not yet been linked to breast cancer, *SLITRK6* has been identified as a novel biomarker in hepatocellular carcinoma^50^, while *SEMA6B* has been associated with poor prognosis and an immunosuppressive tumor microenvironment in colorectal cancer^51^—suggesting, based on our findings, that both genes may serve as candidate markers of disease progression in HR+ metastatic breast cancer.

Taken together, these studies demonstrate that overexpression of these genes is strongly associated with more aggressive breast cancer and poorer prognosis. Building on this evidence, our findings—based on reconstructed phenotypic trajectories before and after treatment—further suggest that these genes may serve novel roles as surface markers capable of predicting which pre-treatment cells are likely to undergo profound phenotypic changes in response to palbociclib and endocrine therapy. In both the *in vitro* dataset and patient samples PA3^18^ and 887^19^, where treatment resistance developed, these markers show promise for the early identification of treatment-resistant cells, paving the way for more targeted and proactive therapeutic strategies.

## Discussion

Here, we introduce PROFET, a computational framework for reconstructing continuous, nonlinear single-cell trajectories from limited and discrete scRNA-seq time points. PROFET addresses a fundamental limitation of traditional OT approaches, which interpolate between observed cellular distributions through cost-minimizing W2 geodesics. While such W2-based methods provide efficient displacement between snapshots, many biological systems exhibit multimodal distributions, branching trajectories, and rare outlier cell states that remain challenging for W2-based trajectory inference. First, the W2 distance requires finite second moments; recent work showed that W2-regularized objectives in general failed to learn distributions lacking this property^15^, whereas W1-regularized objectives succeeded. Second, dynamic OT formulations penalize kinetic energy, biasing inferred trajectories toward displacement-minimizing paths that can shortcut across branching or merging structures commonly observed in cell differentiation trajectories, rather than follow them^7^. Even OT on learned nonlinear latent manifolds can fail to recover a smooth geometry from limited samples, and may produce unreliable interpolations when distinct cell populations from different time points overlap in space^52^. Third, the control-theoretic formulation of dynamic OT is naturally implemented in amortized flow maps, where the time horizon and integration resolution are fixed a priori and may be insufficient for capturing complex geometries. Together, these limitations make it difficult for W2-based methods to learn realistic flows when the underlying biological dynamics involve complex geometry.

To address these limitations, we formulate trajectory inference as a Wasserstein gradient flow of the Lipschitz-regularized KL divergence—mathematically equivalent to a W1 proximal regularized KL divergence. The framework discretizes the Wasserstein gradient flow via forward Euler: at each step, the variational dual formulation of the Lipschitz-regularized KL functional is optimized, particle positions are updated by taking the gradient of the dual potential greedily. The number of steps adapts to the complexity of the data geometry—more steps for branching or merging dynamics, fewer for simpler transitions. This discretization depth determines the resolution of the particle trajectories from which a neural velocity field is subsequently learned in a simulation-free manner, so the complexity of the training signal scales naturally with the data geometry. A subsequent force-matching step then amortizes these piecewise particle trajectories into a globally smooth, time-dependent velocity field that captures the entire cell state transition process, ensuring temporal consistency across subintervals. This two-step framework enables modeling of highly nonlinear and time-inhomogeneous biological dynamics, including multimodal splitting and merging processes, transient cellular states, and strong perturbation responses such as drug treatment.

To further position PROFET within the rapidly growing landscape of trajectory inference methods, we constructed a structured comparison of representative approaches across mathematical framework and training methodology. This comparison highlights that PROFET is uniquely grounded in a variational formulation of Wasserstein gradient flow. While PRESCIENT also assumes gradient-flow dynamics, it does not directly optimize an energy functional, instead learning a time-independent potential through Sinkhorn-based distribution matching. In contrast, most other dynamics-based approaches rely on optimal transport either as a primary training objective or, in the case of MMFM, for sample coupling during spline-based trajectory construction. Methodologically, PROFET combines greedy (backpropagation-free) optimization with simulation-free force matching, avoiding both the backpropagation through unrolled dynamics required by adjoint-based methods and DTO-based methods. These distinctions contribute to PROFET’s scalable and flexible modeling of complex biological dynamics.

Consistent with these theoretical distinctions, we comprehensively benchmarked PROFET against those widely used and recently developed trajectory inference methods for scRNA-seq data and observed strong performance across diverse datasets. Our benchmarking and validation are conducted primarily on real datasets, spanning a range of biological systems (in vitro and in vivo; mouse, axolotl and human), including EMT, stem cell differentiation, regeneration, and hematopoiesis. These datasets exhibit diverse and complex dynamical patterns, including nonlinear trajectories (mESC), merging dynamics (EMT), combined splitting and merging behaviors (axolotl), and lineage divergence (LARRY), which together reduce bias toward any specific modeling assumption and allow us to assess the general applicability of PROFET. To quantify performance, we employ multiple complementary evaluation metrics, including W2 distance, Sinkhorn divergence, and MMD, to assess distributional accuracy. Across the benchmark datasets, PROFET consistently achieved strong performance, with 2.6–12.5× lower W2 error than competing methods. In addition, we incorporate biologically motivated analyses—such as the ability to capture multimodal structure (e.g., emergence and disappearance of cell populations)—to provide a more comprehensive evaluation of model performance.

Given PROFET’s strength in analyzing systems with phenotypic heterogeneity, we applied it to study variability in response to palbociclib, a CDK4/6 inhibitor used in HR+ breast cancer. In an *in vitro* breast cancer cell line treated with escalating doses of palbociclib, we observed a relatively uniform response characterized by downregulation of cell cycle genes and eventual drug resistance. In contrast, patient samples treated with combined palbociclib and endocrine therapy exhibited greater heterogeneity, with some cells showing strong downregulation of both cell cycle and ER signaling genes, while others remained largely unresponsive. Using PROFET to reconstruct single-cell trajectories, we identified a subset of pre-treatment cells that later underwent pronounced phenotypic shifts. Differential expression analysis revealed surface markers—including *UNC5B*, *TLR3*, *PCDH19*, *PROCR*, *SEMA6B*, and *SLITRK6*—that were enriched in this high-shift group, suggesting their potential as early predictors of treatment-associated transitions. Both the *in vitro* model and two patient samples (PA3 and 887) showed signs of treatment resistance or disease progression, implicating these markers in the emergence of resistance. These findings point to the utility of these surface proteins not only for early response prediction in HR+ breast cancer but also as promising targets for ADCs, particularly *SLITRK6*, which is currently under clinical evaluation in ADC-based therapies^43^.

Despite outperforming existing trajectory analysis tools and demonstrating strong utility in biological applications, our approach has several limitations. First, although we have performed a comparison with a broad set of widely used methods and evaluated prediction accuracy using multiple complementary distributional metrics (e.g., W2, Sinkhorn divergence, and MMD), no single quantitative metric can fully capture biological relevance. To address this limitation, we complemented these unbiased distribution-based evaluations with biologically grounded downstream analyses, such as cell-type/subpopulation prediction, to assess whether accurate distributional reconstruction also translates into meaningful biological interpretation. Second, PROFET is specifically designed for time-resolved scRNA-seq data and requires multiple observed time points. It is not intended for single-snapshot datasets, which are typically analyzed using pseudotime inference or stationary optimal transport methods. These approaches rely on additional assumptions (e.g., predefined root states) and aim to infer latent progression rather than reconstruct dynamics in physical time, thus addressing a distinct problem. Third, PROFET is designed for interpolation within observed time intervals and does not perform extrapolation beyond the provided time range. Predicting future states would require additional assumptions about the underlying dynamics and remains an important direction for future work.

In summary, PROFET offers a biologically grounded framework for reconstructing continuous, time-inhomogeneous, and nonlinear single-cell trajectories from discrete time-series scRNA-seq data. This characteristic makes it particularly valuable for analyzing systems with phenotypic heterogeneity, especially in the context of cell state changes and treatment response. By enabling high-resolution inference of gene expression dynamics and fate transitions, PROFET offers a powerful and robust approach for uncovering hidden heterogeneity, tracing phenotypic plasticity, and informing time-sensitive interventions in both developmental and disease contexts.

## Methods

### EXPERIMENTAL MODEL AND SUBJECT DETAILS

#### Synthetic EMT Data

We generated a synthetic dataset mimicking an experimental setup wherein a population of epithelial cells is treated with an inducer of EMT such as TGF-*β* for a fixed duration of time. For this purpose, we started with a previously characterized 26-node gene regulatory network that is known to establish gene expression patterns characteristic of epithelial and mesenchymal cells and to govern the transition between these phenotypic states^53^. The gene regulatory network is described by a connection matrix *J* whose elements represent the regulatory interactions between genes:

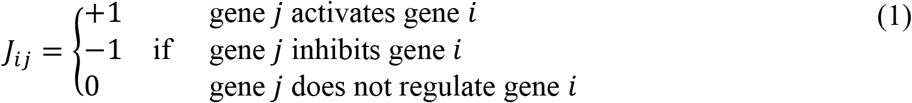

Given the connection matrix *J*, we can write a system of stochastic differential equations to describe the dynamics of the EMT gene expression vector *x* ≡ {*x_i_*}, 1 ≤ *i* ≤ 26:

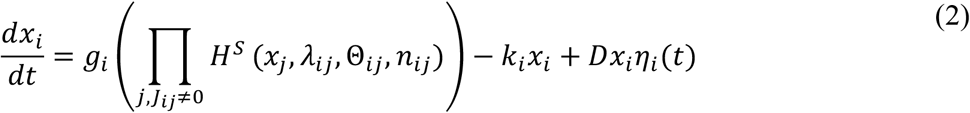

Here, *g_i_* and *k_i_* are the production and degradation rates for gene *i*, respectively. The term *Dx_i_η_i_*(*t*) corresponds to multiplicative noise where *D* = 1 is the noise magnitude and *η_i_*(*t*) is a delta correlated noise term with zero mean and unit variance. *H^S^* is the shifted Hill function:

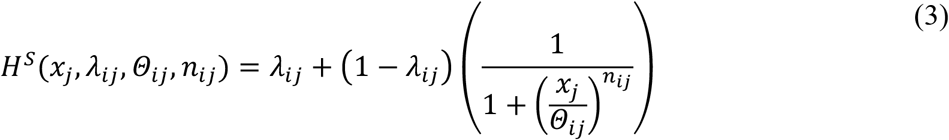

The parameters *λ_i_*_j_, *Θ_i_*_j_, and *n_i_*_j_ together characterize the nature of the regulation of gene *i* by gene *j*:

1. *λ_i_*_j_ is the maximum fold change in the production rate of gene *i* that gene *j* can cause. Thus, *λ_i_*_j_ > 1 if *J_i_*_j_ = +1 and 0 < *λ_i_*_j_ < 1 if *J_i_*_j_ = −1.
2. *Θ_i_*_j_ is the threshold parameter: the regulatory edge form *j* to *i* is active if *x*_j_ > *Θ_i_*_j_. The regulatory interaction is very weak otherwise.
3. *n_i_*_j_ is the Hill coefficient: *H^S^* varies more steeply with *x*_j_ for higher values of *n_i_*_j_.

Given a parameter set ({*g_i_*}, {*k_i_*}, {*λ_i_*_j_}, {*Θ_i_*_j_}, {*n_i_*_j_}) and the initial condition *x*_0_, Eq. (2) can be solved using the Euler-Maruyama method^54^.

#### Preprocessing of mESC data

The dataset analyzed consists of single-cell RNA-sequencing data from mouse embryonic stem cells collected at five time points—0, 12, 24, 48, and 72 hours—comprising 92, 91, 93, 87, and 93 cells, respectively^28,29^. To preprocess the data, we applied a log-transformation to the raw read counts, replacing each value x with log(1+x). Following the same gene selection strategy as in the original study^29^, we calculated the variance of each gene across all cells and selected the top 100 genes with the highest variance for downstream temporal analysis.

#### Preprocessing of EMT data induced by TGF-*β*

The single-cell RNA-sequencing dataset analyzed in this study was obtained from a previous publication on MCF10A cells undergoing TGF-*β*-induced epithelial-to-mesenchymal transition (EMT)^30^. We utilized the processed gene expression matrix provided by the authors; raw sequencing reads are available through the NCBI Sequence Read Archive (BioProject accession PRJNA698642)^55^. For dimensionality reduction, we selected a subset of 72 genes that overlap with a previously established list of 76 genes known to be strongly associated with the EMT gene expression signature^56,57^. The dataset includes cells collected at three time points: day 0 (n = 2,734), day 2 (n = 2,381), and day 4 (n = 1,147).

#### Preprocessing of Axolotl Regeneration Data

The single-cell RNA-sequencing dataset analyzed in this study was obtained from a previously published study on Axolotl limb regeneration^31^. We used the processed gene-expression matrix provided by the authors. For cell-type annotations, we adopted the exact annotation files from the original publication and confirmed their consistency with the classifications reported in the study.

#### Preprocessing of breast cancer cell line data

Samples were aligned with 10x Genomics Cell Ranger (v3.1.0) pipeline^58^ with the cellranger-count command with parameter ‘expect-cells=8000’ to the reference genome refdata-cellranger-GRCh38-3.0.0 provided by 10x Genomics. Filtered feature barcode matrices were imported into R using the Read10X function from Seurat (v5.2.1)^59^ and empty droplets and dead cells were filtered out by exploratory data analysis of QC metrics and manually choosing thresholds to keep cells with percent mitochondrial reads <= 25%, minimum number of genes with nonzero reads per cell >= 100, and minimum total number reads per cell >= 10000. Normalized expression values were calculated with Seurat’s NormalizeData function with default ‘LogNormalize’ options. The dataset comprises cells collected at two time points: pre-treatment (day 0, n = 6,249) and post-treatment (week 16, n = 7,911).

#### Preprocessing of breast cancer patient data

For the Klughammer dataset, we used the original cell annotations to select malignant epithelial cells for downstream analysis^19^. Specifically, for patient 862, the dataset includes cells collected at two time points: pre-treatment (day 0, n = 8,953) and post-treatment (day 200, n = 8,307). For patient 887, cells were also collected at two time points: pre-treatment (day 0, n = 5,536) and post-treatment (day 220, n = 4,638).

For the Luo dataset, we focused on patient PA3 and followed the published inferCNV pipeline^60^ to identify malignant cells. Prior to running inferCNV, we performed unsupervised clustering using Seurat^61^, which identified 15 distinct cell clusters (Figure S21A). Based on the expression of known immune cell marker genes (Table S10), a subset of clusters —specifically clusters 2, 3, 4, 5, 7, 9, 11, 13, and 14 (Figure S21A)— was identified as immune cell populations and designated as reference normal cells for the inferCNV analysis. We then applied inferCNV to infer copy number variations (CNVs) across the remaining cells. The analysis revealed consistently abnormal CNV burden across all clusters except one (Figure S21B).

Further inspection of this exception group (cluster 1) revealed elevated expression of multiple collagen family genes (Figure S21C), suggesting these cells were fibroblasts. As a result, this fibroblast cluster was excluded, and the remaining clusters were retained as malignant epithelial cells. To validate this classification, we examined the expression of EPCAM, a well-established marker of malignant epithelial cells^62^, and confirmed its overexpression in the inferred malignant population (Figure S21D). This confirmed both the malignant nature and epithelial identity of the selected cells. The dataset includes cells collected at two time points: pre-treatment (day 0, *n* = 2,157) and post-treatment (month 11, *n* = 2,535).

### METHOD DETAILS

#### Gradient flow simulation with Generative Particles Algorithm (GPA)

We simulate data-driven trajectories using the Generative Particles Algorithm (GPA)^24^, a deterministic method that iteratively transports particles from a source distribution toward a target. In our setting, each particle represents a cell’s gene expression profile in reduced-dimensional space. GPA uses Lipschitz-regularized gradient flows to interpolate between arbitrary distributions—including empirical, discrete, or irregular ones—in a fully agnostic and mathematically justified manner. This provable flexibility^15^ is essential in single-cell analysis, where the underlying data structure is unknown.

At each time step, GPA estimates a potential function based on the current particle configuration and computes an update direction via its gradient. This approach leads to ODE-based dynamics:

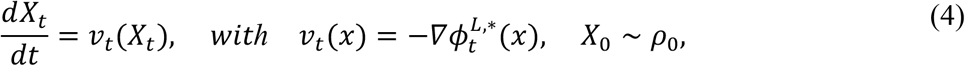

where *ϕ_t_^L^*^,∗^ is the time-dependent potential learned from the data. The underlying Wasserstein gradient flow formulation and the variational structure of the Lipschitz-regularized KL divergence are provided in detail in Supplemental Note 1.

To solve this ODE numerically, GPA uses a forward Euler scheme over *n_T_* steps of size *Δt*, with a greedy optimization at each step:

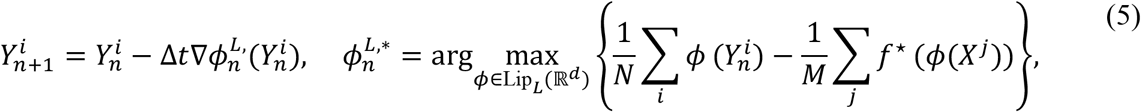

starting from samples 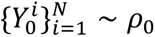. The inner optimization approximates the KL divergence between the current particle distribution and the target distribution using the variational representation of *f*-divergence, where *f*(*x*) = *xlogx* and *f*\* denotes its Legendre transform.

At each iteration, GPA updates the particle positions and records the time-labeled velocities

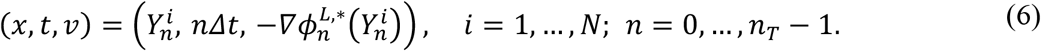

The evolution naturally slows as the particle distribution approaches the target, which is quantified by the decay of the expected kinetic energy 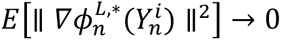 (Theorem 2, Lipschitz-regularized dissipation; Gu et al.^24^).

#### The role of Lipschitz regularization in empirical gradient flow modeling

Lipschitz regularization on the KL divergence serves *three critical purposes* in modeling empirical gradient flows:

1. **Theoretical well-posedness:** Empirical distributions—being finite samples from unknown distributions on *R^d^*—are generally not absolutely continuous with respect to each other. Consequently, the KL divergence between them may become ill-defined or divergent. The Lipschitz-regularized KL divergence resolves this issue by defining a well-posed variational objective even when absolute continuity fails. This approach ensures that the first variation and corresponding flow velocity remain well-defined. See Supplemental Note 2 for a detailed discussion of this issue.
2. **Velocity control:** Lipschitz continuity ensures that the velocity field induced by the flow has a finite speed bound, specifically bounded by the Lipschitz constant *L*. In the limit as *L* → ∞, the Lipschitz-regularized divergence recovers the unregularized KL divergence and its associated gradient flow, confirming that the regularization is a consistent approximation that introduces no fundamental distortion to the KL-based dynamics. In the unregularized case, the velocity grows without bound as the support gets unbounded (Supplemental Note 9). For empirical distributions with compact support, a finite *L* suffices to approximate the KL-induced dynamics well.
3. **Numerical stability and stiffness control:** Bounding the velocity field by a finite Lipschitz constant *L* ensures that the transport speed is controlled, enabling stable time integration under the Courant–Friedrichs–Lewy (CFL) condition for Euler-type methods (2). However, larger *L* values allow the velocity to vary more rapidly across space, which can lead to stiff dynamics that pose numerical instability, as demonstrated in (Gu et al. 2024, Figures 2 and 3). This stiffness–stability trade-off motivates the use of moderate values of *L* that are large enough to capture meaningful dynamical behavior (e.g., convergence rate) but small enough to maintain stable, efficient simulations at practical step sizes. In effect, *L* acts as a dial that balances dynamical accuracy and numerical feasibility in GPA.

In our method, we set *L* = 1 to ensure numerical stability across diverse datasets, providing a universally robust baseline; a sensitivity analysis on *L* is provided in Supplemental Note 7. The step size *Δt* is then adjusted per dataset to accommodate variations in scale and complexity.

#### Handling high-dimensionality via latent-space modeling

Gene expression datasets are intrinsically high-dimensional, with each cell represented by expression levels across thousands of genes. To make learning tractable while preserving meaningful structure, we apply Principal Component Analysis (PCA) to project the data into a lower-dimensional latent space, where GPA is used to model the dynamics. For downstream analysis, we reconstruct the trajectories in the original space using the inverse PCA transform.

Across our 5 datasets — with ambient dimensions *D* = 26,100,72,117,116,115,115 corresponding to the number of genes in synthetic data, mESC data, EMT data, breast cancer cell line, patient PA3, patient 862, and patient 887, respectively — we reduce the dimension of each of them to *d* = 2,8,32, … , *D* and observe that GPA consistently performs well up to moderate latent dimensions (e.g., *d* ≤ 32), capturing temporal structure without overfitting or numerical instability. A sensitivity analysis on prediction errors across PCA dimensionalities is provided in Supplemental Note 7.

To justify latent modeling of high-dimensional biological applications, we provide a theoretical guarantee based on the Lipschitz-regularized divergence in Supplemental Note 10.

#### Stopping time for GPA

As the time horizon for gradient flows is infinite, numerical time-discretized algorithms such as forward Euler require a stopping criterion to determine when particle transport has sufficiently converged. The stopping time *n_T_* determines the effective time horizon, which is then rescaled to the physical time scale between consecutive snapshots, thereby adjusting the resolution of the transport dynamics— more steps for complex dynamics such as branching or merging, fewer for simpler transitions. We use a threshold value of *D_f_^LipL^* < 0.05 (with f=KL and L=1). This threshold was calibrated on the synthetic dataset by computing the Sinkhorn divergence^63^ between the generated and true distributions at held-out intermediate timepoints, and was found to provide optimal alignment for the synthetic dataset example. This threshold provides a practical and computationally efficient criterion for early stopping that balances convergence and overfitting to the endpoint distributions. A sensitivity analysis on stopping time is provided in Supplemental Note 7.

#### Extending GPA to capture nonlinear temporal dynamics

While GPA accurately simulates local transport between pairs of gene expression snapshots, many biological processes—such as drug response or lineage bifurcation—exhibit temporally nonlinear or branched dynamics that cannot be represented by a single global trajectory. To address this issue, we partition the total time horizon [0, *T*] into subintervals and apply GPA independently within each, using the corresponding pair of snapshots.

This strategy yields a sequence of locally consistent gradient flows that adapt to complex temporal structures while preserving interpretability through KL-consistent dynamics. These local simulations serve as the foundation for constructing a unified, continuous-time model in the next stage of our framework.

#### Force-matching for learning a unified velocity field

To amortize the locally simulated gradient flows from multiple GPA subintervals into a global dynamic model, we adopt a force-matching approach^64^. Classically used to learn coarse-grained force fields from trajectory data in molecular simulations^25^, force-matching is here extended to the data-driven regime: instead of relying on predefined physical models, we use GPA-inferred particle trajectories—obtained separately on each subinterval—to supervise the learning of a continuous-time Eulerian velocity field. We require a globally defined velocity field over the entire time horizon to support downstream analyses—such as tracing fate bifurcations (Figure 3E), identifying genes with temporally localized divergence or convergence (Figures 3F–G, 4H–I), and quantifying phenotypic shifts in patient-derived cells (Figure 5C)—all of which rely on continuous integration from arbitrary time points.

We define a neural velocity field *v_θ_*(*x*, *s*) over space and time and train it by minimizing the deviation between the learned velocity and the GPA-estimated transport velocity at randomly sampled time points across all particles:

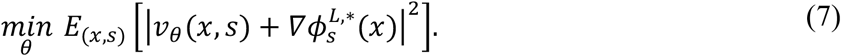

Here, *ϕ*_s_*^L^*^,∗^ is the potential estimated from GPA on the subinterval that covers time *s*. To ensure temporal consistency across subintervals, the physical time variable *s* ∈ [0, *T*] is defined by proportionally aligning each subinterval’s local GPA simulation time *t* ∈ [0, *n_T_*^(*i*)^*Δt*^(*i*)^] to a shared physical time scale. All subintervals are reparameterized relative to the longest GPA simulation horizon *max_i_n_T_*^(*i*)^*Δt*^(*i*)^, allowing the learned velocity field *v_θ_*(*x*, *s*) to be defined on a unified time domain.

The resulting training process unifies the velocity information from all subintervals and fits a smooth vector field that generalizes beyond the discrete particle paths. This formulation distills the localized Lagrangian dynamics simulated by GPA into a global Eulerian flow model, ensuring smooth dynamics at junctions of adjacent time intervals where independently run GPA subintervals may produce mismatched velocities (Figure 2G) and enabling robust interpolation even under limited training data (Figure 2H).

Unlike conventional flow-matching methods^65^, our formulation uses empirical velocity targets derived from simulated particle dynamics. This approach enables the learned field to capture nonlinear biological transitions such as convergence, divergence, or bifurcations in cell states.

#### Overview of the two-step framework

Our method infers continuous-time gene expression dynamics from a small number of time-stamped single-cell snapshots by combining particle-based simulation and neural flow modeling in two successive stages.

In the first stage, we apply GPA independently to each consecutive pair of gene expression distributions {*P_ti_*, *P_ti_*_+1_ }. GPA simulates discrete-time particle trajectories that approximate the Wasserstein gradient flow minimizing the Lipschitz-regularized KL divergence. At each iteration, GPA solves the greedy optimization problem to estimate the dual potential *ϕ_n_^L^*^,∗^ and updates particle positions via forward Euler steps as in Eq. (5). This process yields a time series of particle states and their corresponding velocity vectors.

In the second stage, we train a time-dependent velocity field *v_θ_*(*x*, *s*) parameterized by a neural network to match the simulated velocities from GPA. This is done by minimizing the squared loss between predicted and simulated velocities over all time-labeled particle states. The result is a continuous vector field that generalizes beyond particle locations and interpolates temporal dynamics across the full time horizon.

#### Model architecture and training algorithm

To estimate the potential *ϕ_n_^L^*^,∗^, we use a fully connected feedforward network (MLP) with 3 hidden layers of 128 units each and ReLU activations, followed by a projection layer to enforce the 1-Lipschitz constraint via spectral normalization. Within each forward Euler step of the gradient flow, this potential is optimized for 5 inner gradient-descent (Adam) steps after which the computational graph is discarded before proceeding to the next step, avoiding backpropagation through the trajectory. The velocity field *v_θ_*(*x*, *s*) is modeled as another MLP with 4 hidden layers of 64 units and Tanh activations, taking both position *x* and time *s* as inputs. This velocity field is trained via simulation-free force matching on the particle trajectories obtained from Step 1. Spectral normalization is applied to all layers to ensure Lipschitz continuity, which improves numerical stability and supports temporally smooth interpolation across time points. These architectures were selected based on preliminary experiments indicating reliable scalability to high-dimensional settings—up to ∼100 dimensions in the GPA step and ∼32 dimensions in the force-matching step. We further explored various hyperparameters including learning rates, iteration steps, network size, and regularization thresholds (Figure S22A–D). For the velocity field network, the chosen width and depth provided sufficient expressivity for modeling nonlinear dynamics while avoiding underfitting issues observed in larger architectures, which were prone to optimization instability and diminished learning efficacy.

The overall procedure is summarized in Supplemental Note 11 and is illustrated in Figure 1.

#### Downstream Analysis Using PROFET

To extract biologically meaningful insights from the PROFET-inferred trajectories, we performed three main downstream analyses. First, we reconstructed the dynamic expression profiles of all genes provided in the input data for each individual cell. This step enables gene-level resolution of temporal behavior along each cell’s trajectory. Second, although PROFET outputs trajectories at the single-cell level, we further enhanced interpretability by grouping these trajectories into subpopulations corresponding to distinct cellular behaviors or phenotypic states. Finally, by integrating trajectory classification with single-gene dynamics, we extended conventional differential gene expression analysis—typically performed at discrete time points—to a trajectory-aware framework. This approach allows us to identify key gene markers associated with divergent fate transitions, thereby uncovering regulators that may drive phenotypic bifurcations over time.

#### Reconstruction and evaluation of single-gene dynamics

To ensure stability and computational efficiency, PROFET was applied to reconstruct cell state trajectories in a reduced-dimensional space. Benchmarking against WOT and a null model demonstrated reliable performance with dimensions up to 30 (Figures S23 and S24). Accordingly, we first performed PCA on the gene expression matrix compiled across all input time points to reduce dimensionality. The resulting principal components were then used to represent the data in a lower-dimensional space for trajectory reconstruction. To recover gene-level information, the reconstructed trajectories were mapped back to the original gene expression space using the inverse PCA transformation^66^, thereby yielding predicted gene expression profiles for each individual cell.

To evaluate the accuracy of these reconstructions, we compared the predicted gene expression levels at held-out test time points with the actual expression data. We assessed whether the overall distribution of predicted cell states was statistically indistinguishable from that of the observed test data. We conducted permutation tests using three distributional distance metrics: Total Variation (TV) distance, Kullback-Leibler (KL) divergence, and Sinkhorn divergence^67^. For each metric, we first computed the observed distance between the predicted and observed distributions. Next, we pooled the two datasets and randomly permuted the group labels, recomputing the distance between two groups of equal size for each permutation. This process was repeated 1,000 times to generate a null distribution of distances under the hypothesis that both datasets originate from the same underlying distribution. A one-sided p-value was calculated as the fraction of permutations in which the permuted distance exceeded the observed distance. This approach enabled us to quantitatively assess the alignment between predicted and actual cell state distributions over time. Results for the mESC and EMT datasets, including all p-values, are provided in Table S5 and Table S6, respectively. Finally, we summarized the number of genes for which predicted and observed distributions were statistically indistinguishable (*p* < 0.05), based on one, two, or all three metrics (Figure S25).

#### Classification of sub-trajectories

In this study, we implemented two complementary strategies to classify predicted single-cell trajectories, providing distinct biological interpretations of cell fate dynamics. The first strategy involves classification based on cell subpopulations defined at a specific time point. These subpopulations can be identified using unsupervised clustering alone or in combination with supervised annotations such as known cell types or marker-based labels.

In our analysis, we adopted an unsupervised approach using *k*-means clustering^68^ to define subpopulations from the observed data. Specifically, we first reduced the dimensionality of gene expression matrices using PCA and then applied the *k*-means algorithm to the cell states at a chosen reference time point (e.g., the final day for mESC or the initial day for EMT). We used the *kmeans.fit* function from the scikit-learn Python package^69^ to train the *k*-means model on the PCA-transformed gene expression profiles, which partitions the data into *k* clusters by minimizing within-cluster variance and learns the associated cluster centroids in the reduced feature space. The optimal number of clusters (*k*) was determined using the silhouette score (Figure S26).

Once the *k*-means model was trained on the observed cells, we then used it to assign predicted cell states from PROFET to the learned subpopulations. This was done using the *kmeans.predict* function, which computes the nearest centroid in Euclidean space for each predicted cell state at the reference time point and assigns it the corresponding cluster label. This strategy enables trajectory classification by linking each predicted cell to a biologically meaningful subpopulation derived from the data. For instance, in the mESC dataset, this allowed us to identify sub-trajectories that diverge into distinct terminal fates; in the EMT dataset, it enabled classification of sub-trajectories based on their inferred origins.

The second strategy classifies trajectories based on the extent of cell state transitions between two selected time points. For each trajectory, we computed the Euclidean distance between its predicted cell states at the early and late stages, using this distance as a quantitative proxy for the magnitude of phenotypic change or differentiation. Applying this procedure across all trajectories yielded a distribution of transition distances, which reflects the heterogeneity in dynamic behavior among cells. To stratify trajectories, we categorized them into low, intermediate, or high transition groups based on thresholds identified from local minima in the histogram of transition distances. This data-driven thresholding approach enables separation of distinct transition regimes. Unlike the first strategy, which aligns trajectories with known subpopulations at a fixed reference time point, this approach captures cell state transitions and highlights trajectories undergoing varying degrees of phenotypic change over time.

#### Differential analysis across sub-trajectories

By integrating the reconstructed gene expression dynamics along each predicted trajectory with the sub-trajectory classifications, we conducted two complementary types of differential analysis to identify gene expression differences associated with divergent cell fates. The first approach focused on quantifying temporal divergence in mean gene expression between subpopulations. For each gene, we computed the mean expression within each sub-trajectory group at every time point, then calculated the difference in means between the two groups. These differences were subsequently normalized by the maximum observed value across all time points, producing a relative divergence profile. This method allowed us to pinpoint when specific genes began to diverge between subpopulations, thereby highlighting key temporal features of gene regulation along distinct cell fates.

The second approach involved standard differential gene expression (DGE) analysis^70^ at selected time points to identify gene markers that distinguish sub-trajectory groups. For each gene, we calculated the log_2_ fold change in expression between and assessed statistical significance using a two-sided Welch’s t-test, which accounts for unequal variances between groups. These comparisons were performed at individual time points using sets of cells assigned to different sub-trajectories. By interpreting the results within the framework of sub-trajectory definitions—such as the degree of phenotypic transition or final cell identity—we were able to identify gene expression features associated with distinct developmental paths or treatment responses.

#### Setup of Benchmarked Methods

##### WOT

We employed the Waddington-OT (WOT) method^6^ to analyze cell-state transition probabilities over time in the scRNA-seq data, using normalized expression matrices and day annotations as inputs. Cell growth rates were estimated using a logistic function based on the cells’ proliferation and apoptosis signatures derived from MSigDB gene sets^39^. These growth rates were incorporated into an unbalanced optimal transport formulation to model transitions between consecutive days, using parameters validated in the original study^6^. Following the procedure described in the WOT tutorial, we then performed interpolation between time points. Specifically, an interpolating distribution was computed by first estimating a transport map between the two empirical distributions and then connecting each point in the first distribution to its mapped points in the second distribution according to the transport plan. Finally, a family of interpolations was constructed that continuously sweep from the first distribution to the second, generating a smooth trajectory across time.

##### TrajectoryNet

We employed the TrajectoryNet method^7^ to model continuous-time cellular dynamics using neural ODEs. The model was trained and evaluated on the same PCA basis as used in our framework to ensure a consistent coordinate representation across methods. The architecture of the neural network parameterizing the vector field was set to match ours, using three hidden layers with dimensions specified as --dims “128-128-128”. All other hyperparameters were fixed to the default settings from the original implementation, and the model was trained without incorporating any growth rate estimation or biological priors. To ensure consistent temporal resolution, we linearly interpolated 100 evenly spaced time points along our physical time axis for trajectory generation and evaluation.

##### TIGON

We employed the MIOFlow framework^8^, adapting the implementation provided in the original repository using the Dyngen example. For the PCA-based configuration, we used the same two-dimensional basis as in our experiments. When using PHATE or graph autoencoder (GAE) embeddings, the data were first projected into a 50-dimensional PCA space and then further reduced to two dimensions for visualization and analysis. Training parameters were set following the Embryoid Body Data configuration in the official notebook *01_mioflow.ipynb*, specifically: *n_local_epochs = 40* (phase 1), *n_epochs = 40* (phase 2), *n_post_local_epochs = 0* (phase 3), *reverse_schema = True*, and *reverse_n = 2*. The network architecture followed the default configuration of three layers with hidden dimensions of 16–32–16. Finally, trajectories were interpolated along the physical time axis using 100 evenly spaced time points.

##### PRESCIENT

We employed the PRESCIENT framework^16^ to model cell-state transitions in the same PCA coordinate system used across all methods for consistency. The model was trained with growth weights fixed to zero, ensuring that no external biological priors were introduced. The neural network parameterizing the potential function followed the default architecture of three layers with hidden dimensions of 32–32–32. To ensure consistent temporal resolution, we linearly interpolated 100 evenly spaced time points along our physical time axis for trajectory generation and evaluation.

##### MIOFlows

We employed the TIGON framework^9^ to model dynamic unbalanced optimal transport on the same PCA coordinate system used across all methods for consistency. The default neural network architecture, consisting of four layers with 16–16–16–16 hidden units, was used to parameterize the velocity field. To ensure consistent temporal resolution, we linearly interpolated 100 evenly spaced time points along our physical time axis for trajectory generation and evaluation.

##### CellOT

We applied the CellOT framework using the same two-dimensional PCA coordinate system used across all methods. The transport map was parameterized by an input convex neural network (ICNN) with four hidden layers of dimension 64–64–64–64 and a latent dimension of 50. The model was trained for 100,000 iterations with 10 inner discriminator iterations per step using the Adam optimizer (lr=1e-4, β₁=0.5, β₂=0.9) and a batch size of 256. Trajectories were generated by applying the learned transport map between consecutive observed time points.

##### DeepRUOT

We employed the DeepRUOT framework on the same PCA coordinate system used across all methods. The joint network (FNet) consists of velocity, growth, score, and diffusion sub-networks, each with three hidden layers of dimension 128–128–128 and LeakyReLU activation. Training proceeded in three phases: a velocity/growth pretraining phase using OT loss, a score model training phase (3,001 iterations) using the Schrödinger Bridge conditional flow matching objective with σ=0.05, and a joint fine-tuning phase. Trajectories were generated using stochastic differential equation (SDE) integration with torchsde at 100 evenly spaced time points along the physical time axis.

##### MMFM

We applied the Multi-Marginal Flow Matching (MMFM) framework on the same PCA coordinate system used across all methods. The vector field was parameterized by a network with three output layers of dimension 64 for the data, time, and condition embeddings, using SELU activation and Xavier initialization. The model was trained for 300 epochs with the Adam optimizer (lr=1e-2) and a cosine annealing learning rate schedule, using cubic interpolation and a flow variance of σ=1.0. Trajectories were generated at 100 evenly spaced time points using Euler integration.

##### VGFM

We applied the VGFM framework on the same PCA coordinate system used across all methods. The velocity field was parameterized by a network with three hidden layers of dimension 128–128–128 and LeakyReLU activation. Training consisted of two phases: a pretraining phase minimizing the VGFM objective for 2,000 epochs (Adam, lr=1×10⁻³, UOT regularization reg=0.5, reg_m=5, batch size=60), followed by a refinement phase incorporating an OT loss for 30 epochs (Adam, lr=1×10⁻⁴). Trajectories were generated at 10 evenly spaced time points along the physical time axis.

##### PI-SDE

We applied the PI-SDE framework on the same PCA coordinate system used across all methods. The drift was derived from a time-dependent potential network with two hidden layers of dimension 400–400 and softplus activation, while diffusion was set to a constant σ=0.1. The model was trained for 3,000 epochs using the Adam optimizer (lr=5×10⁻³) with a step learning rate scheduler (step size=100, γ=0.9), Sinkhorn OT loss (blur=0.1, scaling=0.7), and gradient clipping at 0.1. Trajectories were generated at 100 evenly spaced time points along the physical time axis.

### QUANTIFICATION AND STATISTICAL ANALYSIS

#### Computing distributional distances between predicted and test-time distributions

To quantify how closely the predicted cell states aligned with the experimentally observed distributions at withheld time points, we computed distributional distances using three metrics: W2 distance, Sinkhorn divergence, and maximum mean discrepancy (MMD). We approximated the W2 distance using the entropic optimal transport cost computed via the *SamplesLoss* function from the *GeomLoss* library^71^. This regularized OT cost is defined as:

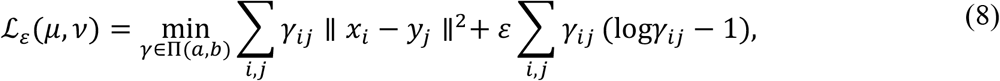

where *γ* is a transport plan between discrete empirical distributions *μ* = ∑*_i_ a_i_ δ_xi_* and *ν* = ∑_j_ *b*_j_ *δ_y_*_j_ with marginals *a* and *b*, and *ε* > 0 controls the strength of entropy regularization. This formulation provides a smooth approximation of the W2 distance that is computationally stable and scalable^72^. To obtain a proper divergence, we computed the Sinkhorn divergence by correcting for the bias introduced by the entropy term:

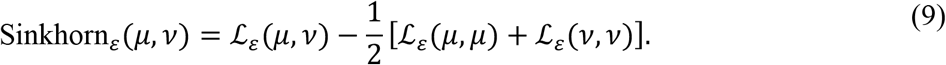

We tested a range of entropy regularization parameters *ε* ∈ {0.01,0.1,1,10,100} to assess the sensitivity of this metric. Additionally, we computed MMD between predicted and observed distributions using a radial basis function (RBF) kernel:

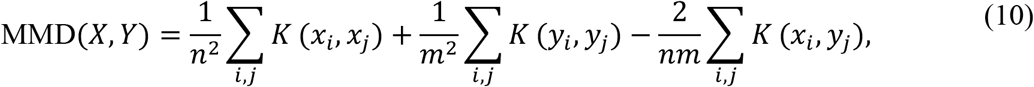

where *K*(*x*, *y*) = exp(−*γ* ∥ *x* − *y* ∥^2^) is the RBF kernel with bandwidth *γ* = 1.072. All computations were performed in Python using the *GeomLoss*^73^ and *POT (Python Optimal Transport)*^74^ libraries.

#### Permutation testing of predicted versus observed distributions

To assess the agreement between predicted gene expression distributions and experimentally measured distributions at held-out time points, we conducted permutation-based hypothesis testing for each gene. Specifically, we evaluated whether the predicted and observed distributions could be considered samples from the same underlying distribution (null hypothesis), using three complementary distance metrics: total variation distance (TV), Kullback–Leibler divergence (KL), and Sinkhorn divergence. For each gene and each time point, we computed the observed value of the respective distance metric between the predicted and test distributions. We then permuted the pooled samples 1,000 times to construct a null distribution and estimated the empirical *p*-value as the proportion of permuted statistics greater than or equal to the observed one.

To evaluate the robustness of our predictions, we quantified the number of genes for which the null hypothesis could not be rejected (*p* > 0.05) under at least one, at least two, or all three distance metrics. In the mESC dataset (100 genes), at the held-out time point 1, 78 genes passed the test with at least one metric, 50 genes with at least two, and 37 genes with all three. At time point 3, these numbers increased to 100, 99, and 91, respectively. In the EMT dataset (72 genes), when predicting time point 2, 62 genes passed with at least one metric, 8 with at least two, and only 1 gene with all three. These results—summarized in Figure S25, with all *p*-values reported in Tables S5 and S6—highlight the consistency of PROFET’s predictions and underscore the value of using multiple statistical metrics to validate distributional similarity across time points.

#### Displacement-based quantification of phenotypic shift

To quantify the extent of phenotypic shifts inferred by PROFET, we computed the Euclidean displacement for each cell between its predicted pre-treatment and post-treatment states. Summary statistics—including the mean, standard deviation, interquartile range (IQR), coefficient of variation (CV), and entropy of the displacement distribution—were recorded for each dataset (Table S8). The full implementation can be found in the *plot_X1_hat_displacement_distribution* function, which loads the trained model and PCA parameters, computes displacements, saves statistical outputs as CSV files, and generates visualizations.

#### KDE-based stratification of cellular subpopulations

To identify subpopulations with distinct phenotypic shifts, we used kernel density estimation (KDE) to smooth the histogram of displacement values^75^. The density curves were computed using the Scott bandwidth method, and local minima were detected via peak-finding on the inverse KDE curve. The two smallest local minima (based on the *x*-axis values) were selected as cutoffs to stratify cells into three groups: low, medium, and high phenotypic shift. This analysis was performed independently for each patient dataset using the corresponding histogram of cell displacements. The KDE and local minima detection steps are implemented in a Python pipeline that reads per-sample histograms, applies KDE smoothing via *scipy.stats.gaussian_kde*, and uses *scipy.signal.find_peaks* to locate valleys in the density curve^76^.

#### Assessing EMT Score

We computed the EMT score as a weighted sum of the expression levels of 76 EMT-related genes^56^. The weight assigned to each gene was determined by its correlation with the expression of CDH1 (E-cadherin). The resulting scores were normalized to have a mean of zero, such that negative values indicate a cell state closer to the epithelial (E) phenotype and positive values correspond to the mesenchymal (M) phenotype. To ensure consistent interpretation, we further rescaled the scores by multiplying them by –1, thereby aligning their direction with the progression from the epithelial to mesenchymal state.

## RESOURCE AVAILABILITY

### Lead Contact

Further information and requests for resources should be directed to and will be fulfilled by the Lead Contact, Franziska Michor.

### Materials Availability

This study did not generate new unique reagents or materials.

### Data and Code Availability

● This study analyzes existing, publicly available single-cell RNA-sequencing datasets. Accession numbers for all datasets are provided in the Key Resources Table.
● All original code used to implement PROFET, perform downstream analyses, benchmark competing methods, and generate the figures reported in this study has been deposited in a public repository and is publicly available as of the date of publication. The archived version of the code is available through Zenodo, and the corresponding GitHub repository (https://github.com/HyeminGu/PROFET) and Zenodo DOI are listed in the Key Resources Table.
● Any additional information required to reanalyze the data reported in this study is available from the Lead Contact upon request.

### Author Contributions

Y.-C.C. and H.G. contributed equally to this work. Y.-C.C., H.G., T.O.M., S.T., Y.W., Y.P., H.L., M.A.K., and F.M. conceived and designed the study. Y.-C.C., H.G., T.O.M., S.T., Y.P., M.A.K., and F.M. developed the methodology. Y.-C.C., H.G., and M.B. developed the computational software. Y.-C.C., H.G., C.G., Y.W., Y.Z., H.L., R.J., M.A.K., and F.M. validated the results. Y.-C.C., H.G., W.W., S.T., M.B., and Y.W. performed the computational and statistical analyses. C.G., D.R., D.L.A., and R.J. generated experimental data and provided biological resources. Y.-C.C., H.G., S.T., C.G., D.R., D.L.A., Y.W., H.L., R.J., and F.M. curated and interpreted the data. Y.-C.C., H.G., W.W., S.T., M.B., M.A.K., and F.M. prepared the figures and visualizations. Y.-C.C., H.G., R.J., M.A.K. and F.M. wrote the original manuscript draft. All authors reviewed, edited, and approved the manuscript. T.O.M., H.L., R.J., M.A.K., and F.M. supervised the research. Y.-C.C., H.G., R.J., M.A.K., and F.M. coordinated the project. R.J., M.A.K. and F.M. acquired funding for the study.

## Acknowledgements

Y.-C.C. and F.M. thank the Center for Cancer Evolution at Dana-Farber Cancer Institute, Jacob Geisberg, Simona Cristea, and members of the Michor lab for their support and helpful discussions. Y.-C.C. further acknowledges insightful conversations with Helen Byrne (University of Oxford), Tom Chou (UCLA), and Hong Qian (University of Washington). M.K. and H.G. are partially funded by AFOSR grant FA9550-21-1-0354 and NSF grant DMS-2307115. W.W. is grateful for the support from the National Institute of General Medical Sciences (grant T32GM144273). The content of this study is solely the responsibility of the authors and does not necessarily represent the official views of the National Institute of General Medical Sciences or the National Institutes of Health. D.L.A. acknowledges support from the DF/HCC Breast Specialized Program in Research Excellence (SPORE) grant 1P50CA168504, Cancer Couch Foundation, Hope Scarves, the Jamieson Family Fund for Early Career Breast Cancer Researchers, and the Saverin Family Fund. H.L. acknowledges funding from the Center for Theoretical Biological Physics through NSF PHY-2019745 and NSF DMS-245957. C.G. was supported by fellowships from the Associazione Italiana Ricerca sul Cancro (AIRC) and the American-Italian Cancer Foundation (AICF) during her postdoctoral training in R.J.’s lab. Y.Z. is supported by the National Natural Science Foundation of China (82372750), the CAMS Innovation Fund for Medical Sciences (2023-I2M-3-003 and 2022-I2M-1-008), and the National Key R&D Program of China (2023YFC2507000).

## Supplemental information

Figure S1, PROFET model development and validation process, related to the section “Overview of PROFET.”

Figure S2, Validation and benchmarking of PROFET using mESC data, related to Figure 2AB. Figure S3, Classification of two EMT cell types at days 0 and 2, related to Figure 2D.

Figure S4, Validation and benchmarking of PROFET using EMT data, related to Figure 2D. Figure S5, Validation and benchmarking of PROFET using Axolotl data, related to Figure 2E.

Figure S6, Validation and benchmarking of PROFET using the LARRY hematopoiesis scRNAseq dataset, related to Figure 2F.

Figure S7, Sensitivity analysis of PROFET hyperparameters, related to the section “Ablation Analysis, Robustness, and Computational Scalability.”

Figure S8, Reconstruction of single-cell, single-gene dynamics in the mESC dataset, related to Figure 3B.

Figure S9, Predicted average gene expression trajectories in the mESC dataset, related to Figure 3C.

Figure S10, Comparison of predicted and observed gene expression distributions at held-out time points in the mESC dataset, related to Figure 3D.

Figure S11, Average gene expression trajectories for each identified subgroup in the mESC dataset, related to Figure 3G.

Figure S12, Downstream analysis of reconstructed single-cell trajectories for EMT data, related to Figure 4AB.

Figure S13, Reconstruction of single-cell, single-gene dynamics in the EMT dataset, related to Figure 4B.

Figure S14, Comparison of predicted and observed gene expression distributions at held-out time points in the EMT dataset, related to Figure 4B.

Figure S15, Average EMT relevant gene expression trajectories plotted separately for each subgroup, related to Figure 4G.

Figure S16, Average stemness relevant gene expression trajectories plotted separately for each subgroup, related to Figure 4G.

Figure S17, Single-cell, single-gene expression dynamics reconstructed by PROFET in the MCF7 cell line dataset, related to the section “Trajectory-Based Dissection of Treatment Response Heterogeneity and Early Biomarker Discovery.”

Figure S18, Single-cell, single-gene expression dynamics reconstructed by PROFET in the patient PA3 dataset, related to the section “Trajectory-Based Dissection of Treatment Response Heterogeneity and Early Biomarker Discovery.”

Figure S19, Single-cell, single-gene expression dynamics reconstructed by PROFET in the patient 862 dataset, related to the section “Trajectory-Based Dissection of Treatment Response Heterogeneity and Early Biomarker Discovery.”

Figure S20, Single-cell, single-gene expression dynamics reconstructed by PROFET in the patient 887 dataset, related to the section “Trajectory-Based Dissection of Treatment Response Heterogeneity and Early Biomarker Discovery.”

Figure S21, inferCNV analysis of patient PA3 from Luo et al, related to the section “Methods: Preprocessing of breast cancer patient data.”

Figure S22, Sensitivity of trajectory prediction performance to hyperparameter selection, related to the section “Methods: Model architecture and training algorithm.”

Figure S23, Effect of latent dimensionality on trajectory prediction performance in the mESC dataset, related to the section “Methods: Downstream Analysis Using PROFET.”

Figure S24, Effect of latent dimensionality on trajectory prediction performance in the EMT dataset, related to the section “Methods: Downstream Analysis Using PROFET.”

Figure S25, Overlap of genes with no significant differences identified by multiple distributional metrics, related to the section “Methods: Downstream Analysis Using PROFET.”

Figure S26, Silhouette scores from clustering analysis across different numbers of clusters, related to the section “Methods: Downstream Analysis Using PROFET.”

Figure S27, Simulation setup to model epithelial-mesenchymal transition in a population of epithelial cells, related to Supplemental Note 4.

Figure S28, The learned trajectories from TrajectoryNet on synthetic data (top), EMT data (center), and stem cell differentiation data (bottom) diverge substantially from the true trajectories, related to Supplemental Note 6.

Figure S29, Convergence speed of Lipschitz-regularized KL GPA with different Lipschitz constants $L=1, 10$ and Ornstein–Uhlenbeck (OU) process, related to Supplemental Note 9.

Table S1. Comparison of trajectory inference methods via mathematical framework and theoretical properties, related to the section “Model Validation and Benchmarking Across Four Biological Datasets”.

Table S2. Comparison of trajectory inference methods via training methodology, related to the section “Model Validation and Benchmarking Across Four Biological Datasets”.

Table S3. Wasserstein-2 prediction error across benchmark models and datasets (five intermediate-timepoint snapshots) and comparison with PROFET, related to the section “Model Validation and Benchmarking Across Four Biological Datasets”.

Table S4. Comparison of trajectory inference methods via runtime and memory, related to the section “Ablation Analysis, Robustness, and Computational Scalability”.

Table S5. Comparison of predicted and observed gene expression distributions at the held-out time points (time points 1 and 3) in the mESC dataset, related to Figure 3G.

Table S6. Comparison of predicted and observed gene expression distributions at the held-out time point (day 2) in the EMT dataset, related to Figure 4E.

Table S7. List of 121 genes previously implicated in the response to palbociclib, including genes involved in cell cycle regulation, MYC and estrogen receptor (ER) signaling, growth factor pathways, the Hippo pathway, and inflammatory processes, related to the section “Tracing Phenotypic Shifts Associated with Palbociclib Therapy in HR+ Breast Cancer”.

Table S8. Coefficients of variation (CVs) of the predicted cell-state trajectory displacements for the in vitro dataset and patients PA3, 862, and 887, related to Figure 5B.

Table S9. Differential gene expression (DEG) analysis comparing post-treatment and pre-treatment cells in the in vitro dataset and patients PA3, 862, and 887, related to Figure 6E.

Table S10. Immune cell marker genes used for inferCNV analysis of patient PA3 from Luo et al, related to the section “Methods: Preprocessing of breast cancer patient data”.

Supplemental Note 1: Wasserstein Gradient Flow and Lipschitz-Regularized KL Divergence, related to the sections “Overview of PROFET” and “Gradient flow simulation with Generative Particles Algorithm (GPA)”.

Supplemental Note 2: Robustness through Lipschitz regularization, related to the sections “Overview of PROFET” and “The role of Lipschitz regularization in empirical gradient flow modeling”.

Supplemental Note 3: Comparison to flow matching in score-based generative modeling, related to the section “Overview of PROFET”.

Supplemental Note 4: Simulation setup to model epithelial-mesenchymal transition, related to the section “Overview of PROFET”.

Supplemental Note 5: Full description of novelty, related to the section “Model Validation and Benchmarking Across Four Biological Datasets”.

Supplemental Note 6: Simulation results on TrajectoryNet, related to the section “Model Validation and Benchmarking Across Four Biological Datasets”.

Supplemental Note 7: Ablation studies and robustness analyses, related to the sections “Ablation Analysis, Robustness, and Computational Scalability”, “The role of Lipschitz regularization in empirical gradient flow modeling” and “Stopping time for GPA”.

Supplemental Note 8: Computational Efficiency and Scalability, related to the section “Ablation Analysis, Robustness, and Computational Scalability”.

Supplemental Note 9: Velocity growth in KL gradient flow and the role of Lipschitz regularization, related to the section “The role of Lipschitz regularization in empirical gradient flow modeling”.

Supplemental Note 10: Theoretical guarantee for latent-space modeling, related to the section

“Handling high-dimensionality via latent-space modeling”.

Supplemental Note 11: PROFET algorithm, related to the section “Model architecture and training algorithm”.

