## Supplemental Materials for "PROFET Predicts Continuous Gene Expression Dynamics from scRNA-seq Data to Elucidate Heterogeneity of Cancer Treatment Responses"

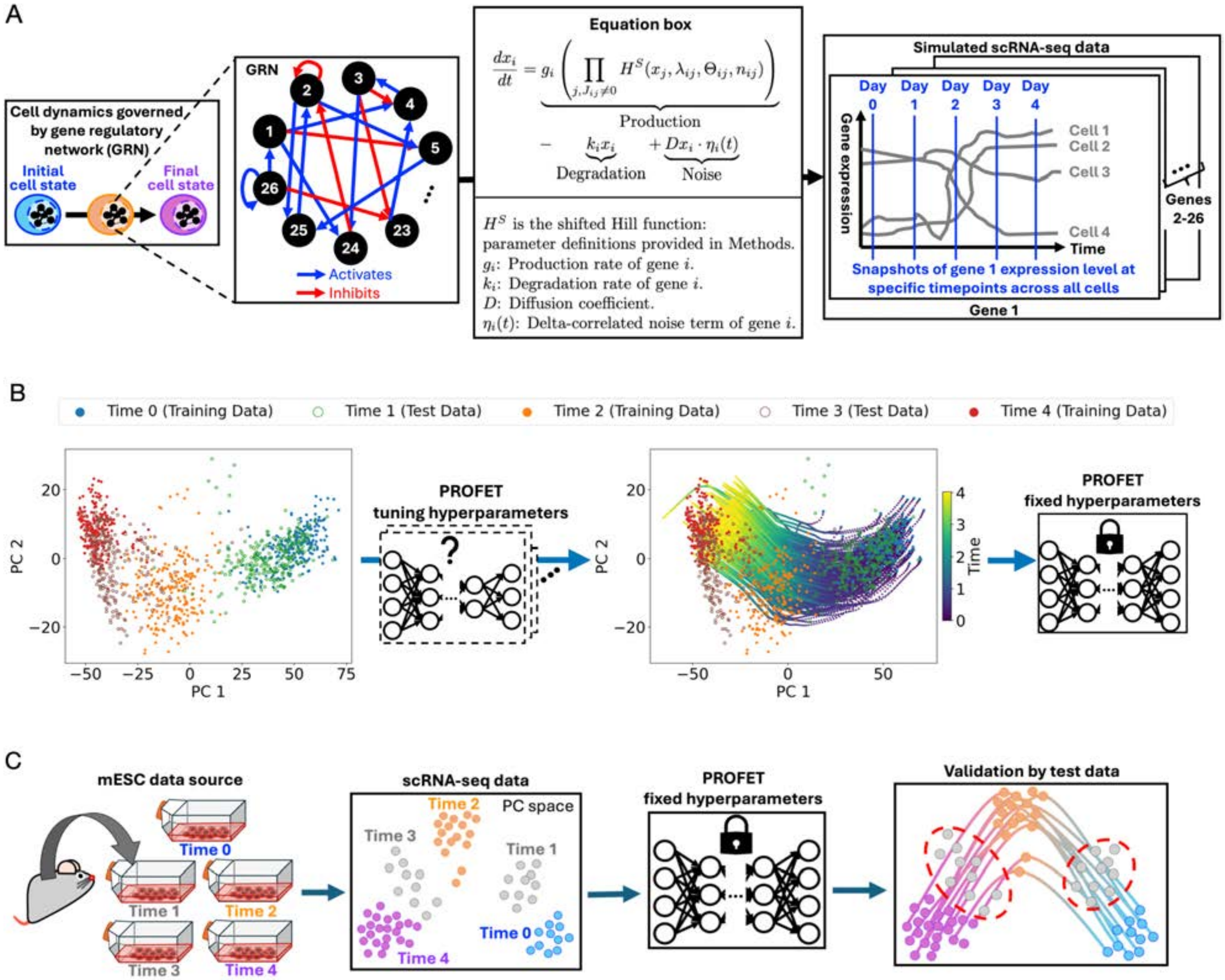

Figure S1, PROFET model development and validation process, related to the section “Overview of PROFET.” (A) Overview of the synthetic data generation process simulating TGF-β–induced EMT using a gene regulatory network model. (B) PROFET development using synthetic EMT data. The model is trained using input from time points 0, 2, and 4 to reconstruct full single-cell trajectories. Hyperparameters were selected to ensure stable, non-divergent trajectories that align with the distributions at the withheld test time points. (C) Schematic of the validation process using mESC differentiation data. The model, with fixed hyperparameters, is applied to reconstruct trajectories from input data at time points 0, 2, and 4, and evaluated using held-out data from time points 1 and 3.

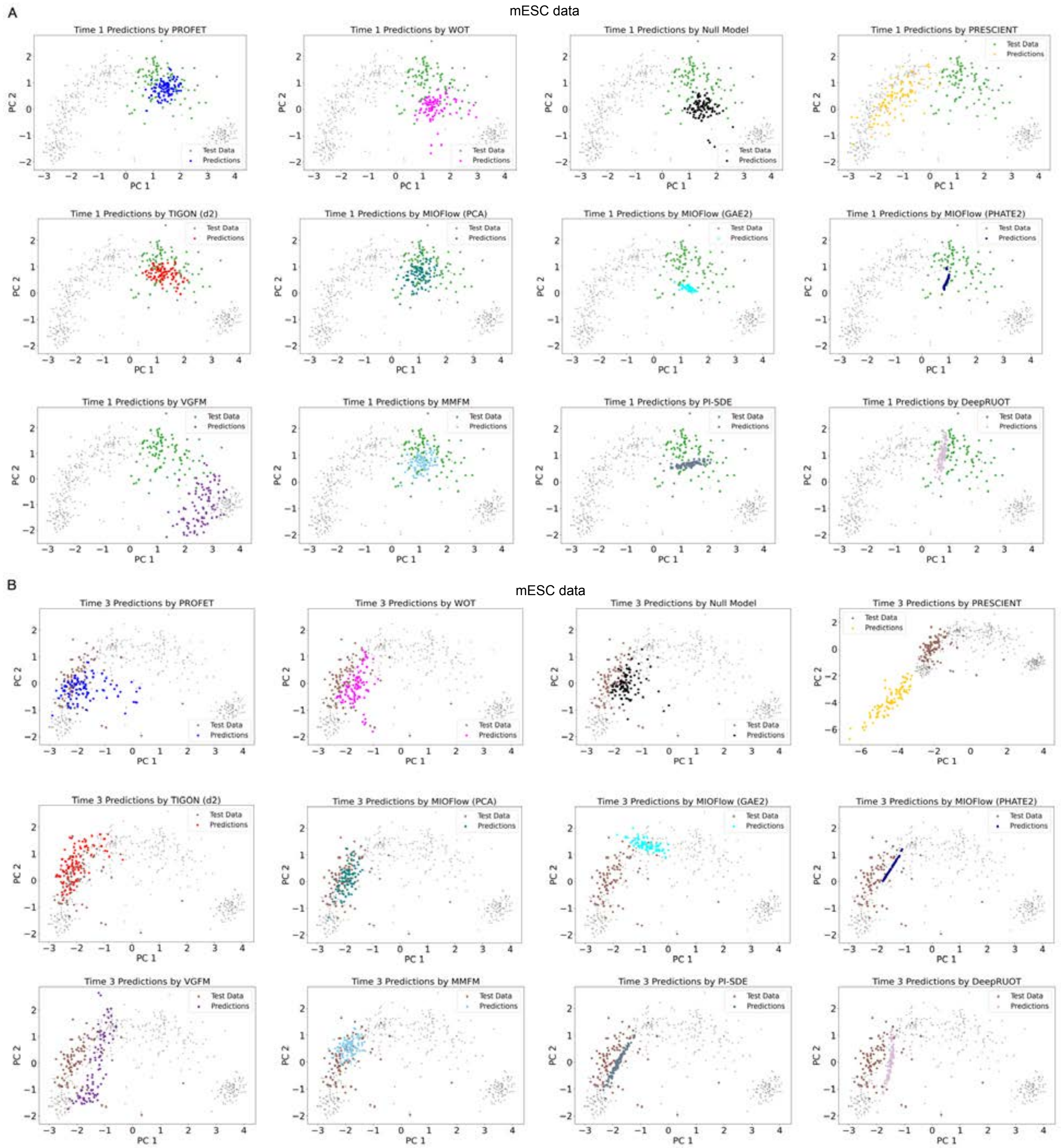

Figure S2, Validation and benchmarking of PROFET using mESC data, related to Figure 2AB. (A) Comparison of predicted test data at time 1 (12 hours; green) across ten methods. (B) Comparison of predicted test data at time 3 (48 hours; brown) across the same ten methods.

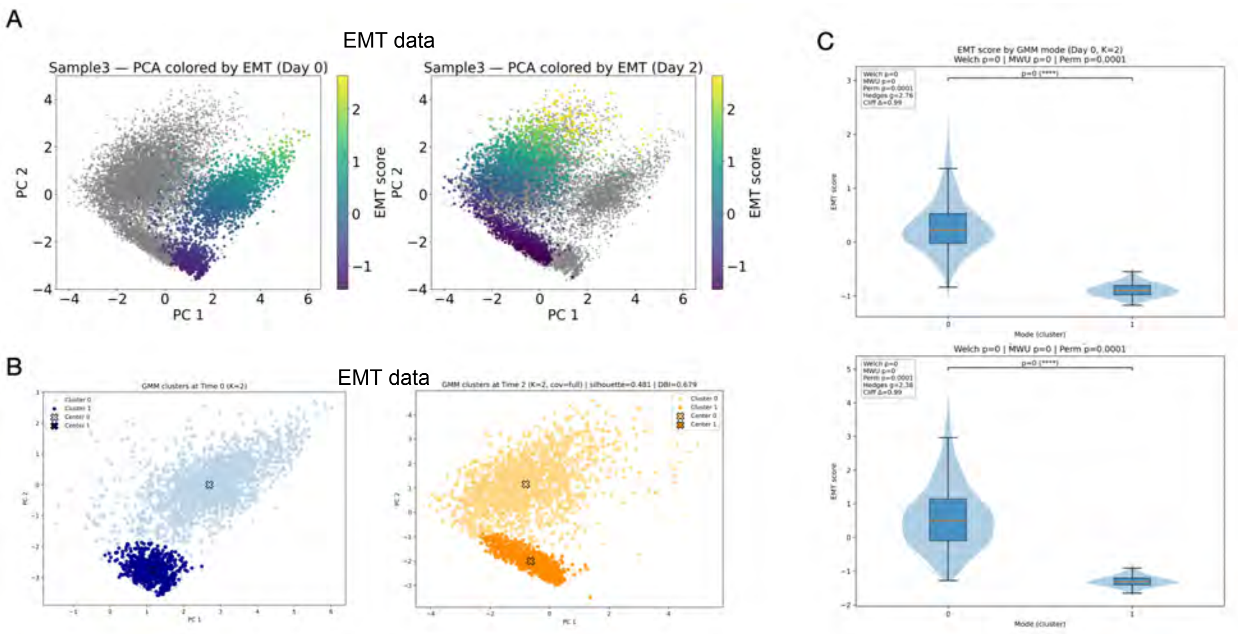

Figure S3, Classification of two EMT cell types at days 0 and 2, related to Figure 2D. (A) Colormap of single-cell EMT scores computed using the 76GS method. (B) Classification of two EMT subpopulations using Gaussian mixture models (GMM). (C) Distribution of EMT scores for the two subpopulations. Each point represents one cell. Mean EMT scores are indicated, and statistical significance was assessed using a two-sided Welch's t-test (replace with the actual test if different). P-values indicate the significance of the difference between the two subpopulations.

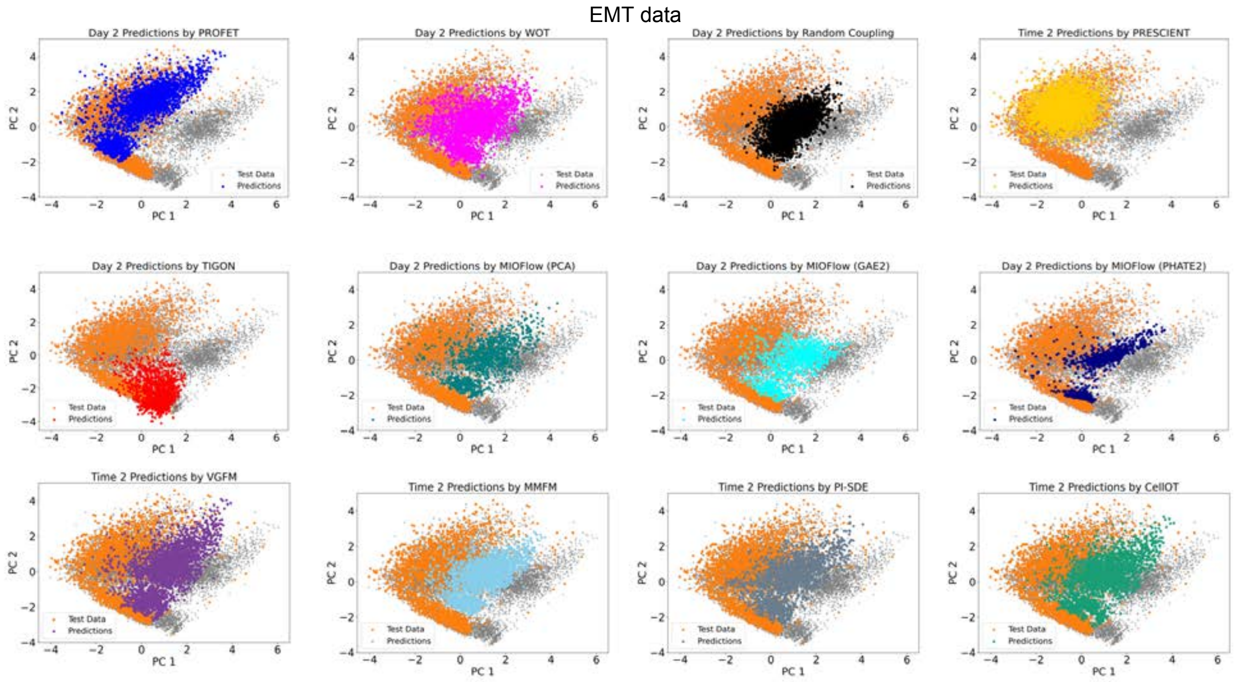

Figure S4, Validation and benchmarking of PROFET using EMT data, related to Figure 2D. Comparison of predicted test data at day 2(green) across ten methods.

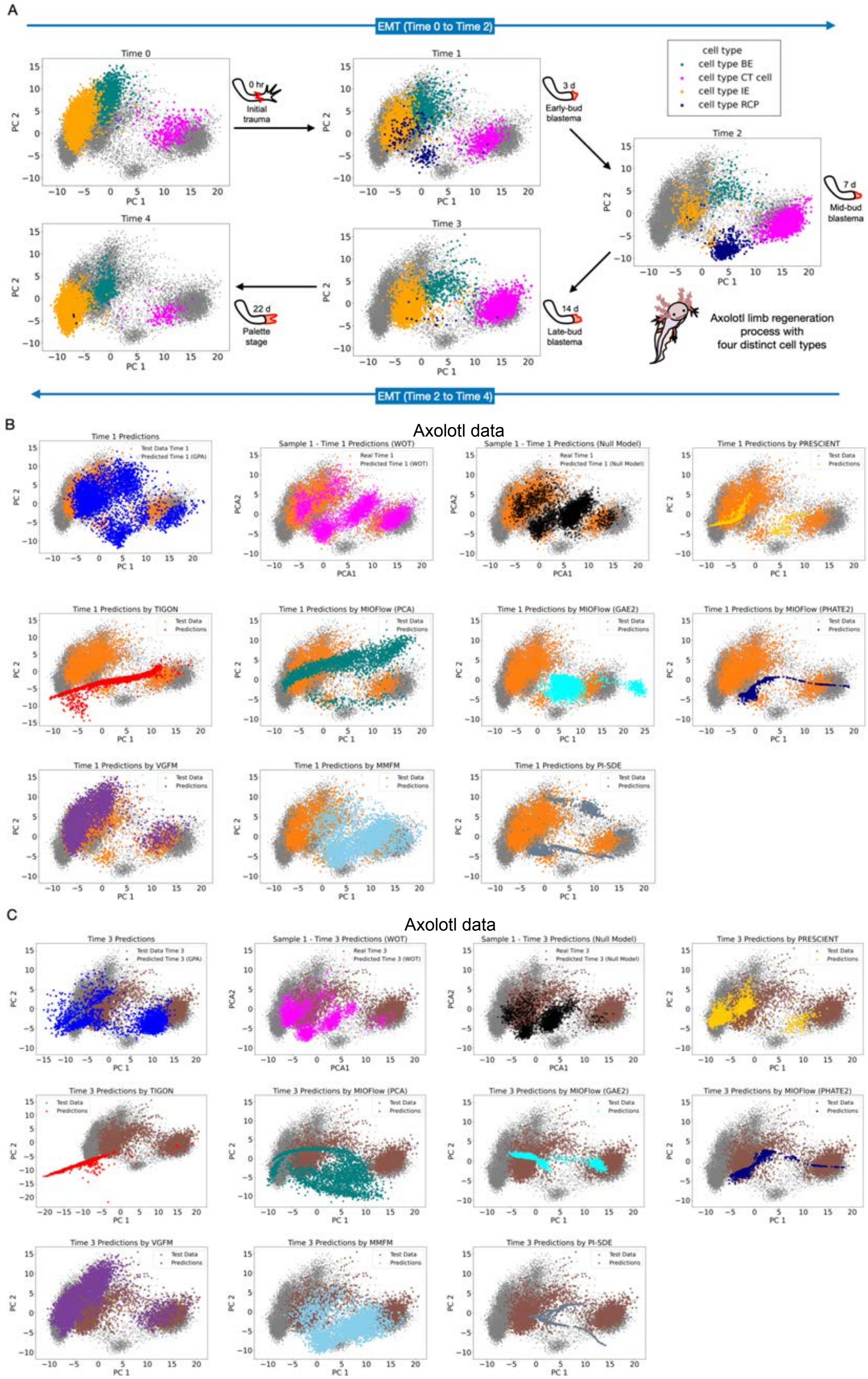

A
EMT (Time 0 to Time 2)
Time 0
0 hr
Initial trauma
Time 1
3 d
Early-bud blastema
cell type
cell type BE
cell type CT cell
cell type IE
cell type RCP
Time 2
7 d
Mid-bud blastema
Time 4
22 d
Palette stage
Time 3
14 d
Late-bud blastema
Axolotl limb regeneration process with four distinct cell types
EMT (Time 2 to Time 4)
PC 1
PC 2
B
Axolotl data
Time 1 Predictions
Test Data Time 1
Predicted Time 1 (GPA)
Sample 1 - Time 1 Predictions (WOT)
Real Time 1
Predicted Time 1 (WOT)
Sample 1 - Time 1 Predictions (Null Model)
Predicted Time 1 (Null Model)
Time 1 Predictions by PRESCIENT
Time 1 Predictions by TIGON
Time 1 Predictions by MIOFlow (PCA)
Time 1 Predictions by MIOFlow (GAE2)
Time 1 Predictions by MIOFlow (PHATE2)
Time 1 Predictions by VGFM
Time 1 Predictions by MMFM
Time 1 Predictions by PI-SDE
Test Data
Predictions
PCA1
PCA2
C
Axolotl data
Time 3 Predictions
Test Data Time 3
Predicted Time 3 (GPA)
Sample 1 - Time 3 Predictions (WOT)
Real Time 3
Predicted Time 3 (WOT)
Sample 1 - Time 3 Predictions (Null Model)
Predicted Time 3 (Null Model)
Time 3 Predictions by PRESCIENT
Time 3 Predictions by TIGON
Time 3 Predictions by MIOFlow (PCA)
Time 3 Predictions by MIOFlow (GAE2)
Time 3 Predictions by MIOFlow (PHATE2)
Time 3 Predictions by VGFM
Time 3 Predictions by MMFM
Time 3 Predictions by PI-SDE

Figure S5, Validation and benchmarking of PROFET using Axolotl data, related to Figure 2E. (A) Illustration of the Axolotl scRNA-seq dataset, highlighting the cell state distributions of four cell types across five time points (days 0, 3, 7, 14, and 22). (B) Comparison of predicted test data at time 1 (day 3; orange) across 9 methods. (C) Comparison of predicted test data at time 3 (day 14; brown) across the same 9 methods.

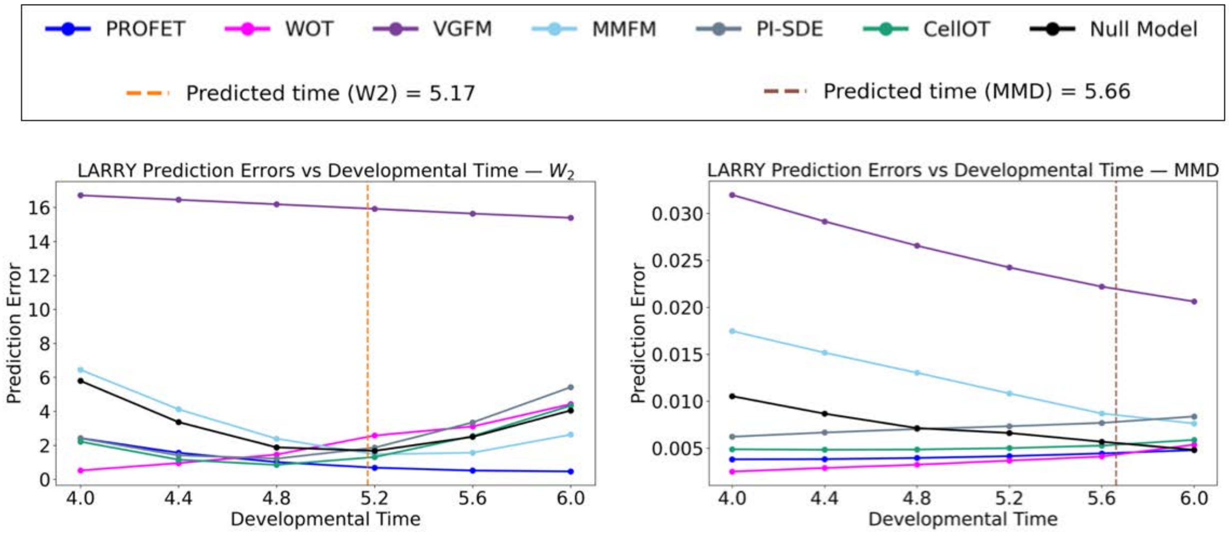

Figure S6, Validation and benchmarking of PROFET using the LARRY hematopoiesis scRNAseq dataset, related to Figure 2F. Prediction errors across interpolated developmental times between days 4 and 6 are shown using the W2 metric (left panel) and the MMD metric (right panel). The vertical dashed line indicates the predicted developmental time, estimated by matching the ratio of W2 distances (left) or MMD distances (right) between the day 2–4 and day 4–6 intervals. Insets show the prediction errors of PROFET and benchmark methods at the estimated developmental time.

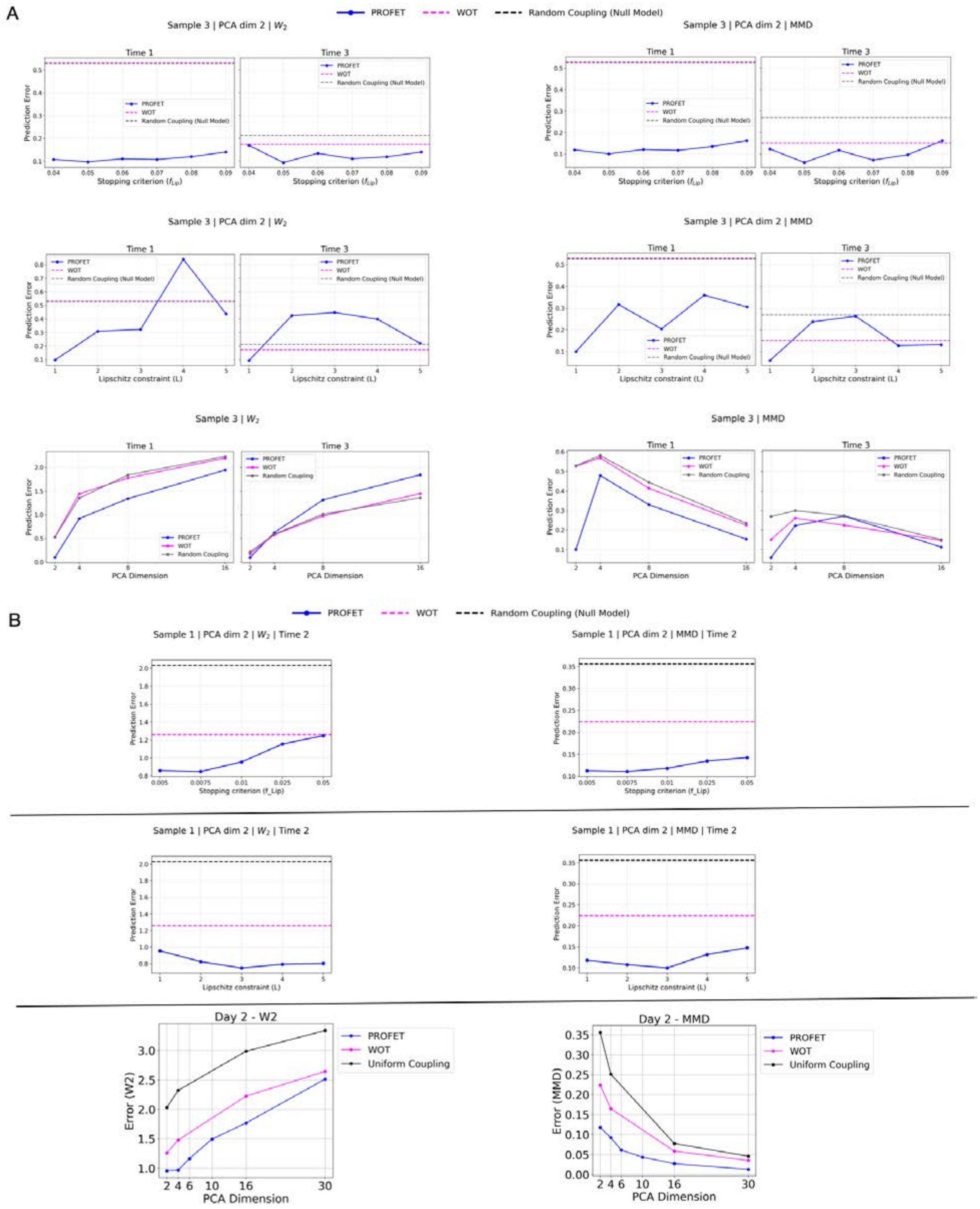

Figure S7, Sensitivity analysis of PROFET hyperparameters, related to the section "Ablation Analysis, Robustness, and Computational Scalability."**.** Prediction errors measured by W2 distance (left panels) and MMD (right panels) for the mESC dataset at time points 1 and 3 (A) and the EMT dataset at day 2 (B). Performance is evaluated under varying hyperparameter settings, including the GPA stopping criterion (defined by the *f*-Lipschitz bound), the Lipschitz constant (*L*), and the dimensionality of the PCA-reduced feature space. Lower values indicate better agreement between predicted and observed cell-state distributions at the held-out time points.

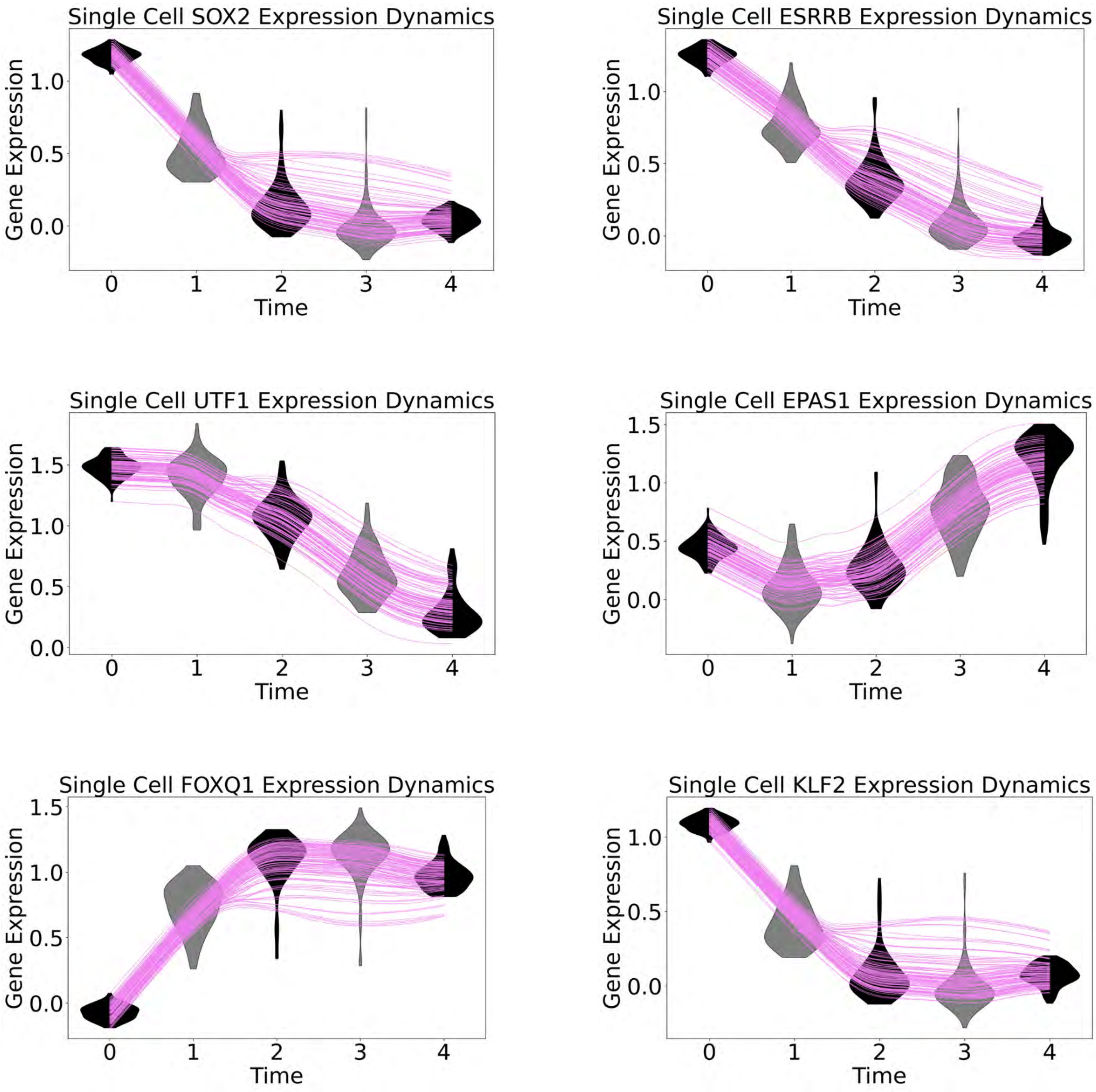

Single Cell SOX2 Expression Dynamics
Gene Expression
0.0
0.5
1.0
0
1
2
3
4
Time
Single Cell ESRRB Expression Dynamics
Gene Expression
0.0
0.5
1.0
0
1
2
3
4
Time
Single Cell UTF1 Expression Dynamics
Gene Expression
0.0
0.5
1.0
1.5
0
1
2
3
4
Time
Single Cell EPAS1 Expression Dynamics
Gene Expression
0.0
0.5
1.0
1.5
0
1
2
3
4
Time
Single Cell FOXQ1 Expression Dynamics
Gene Expression
0.0
0.5
1.0
1.5
0
1
2
3
4
Time
Single Cell KLF2 Expression Dynamics
Gene Expression
0.0
0.5
1.0
0
1
2
3
4
Time

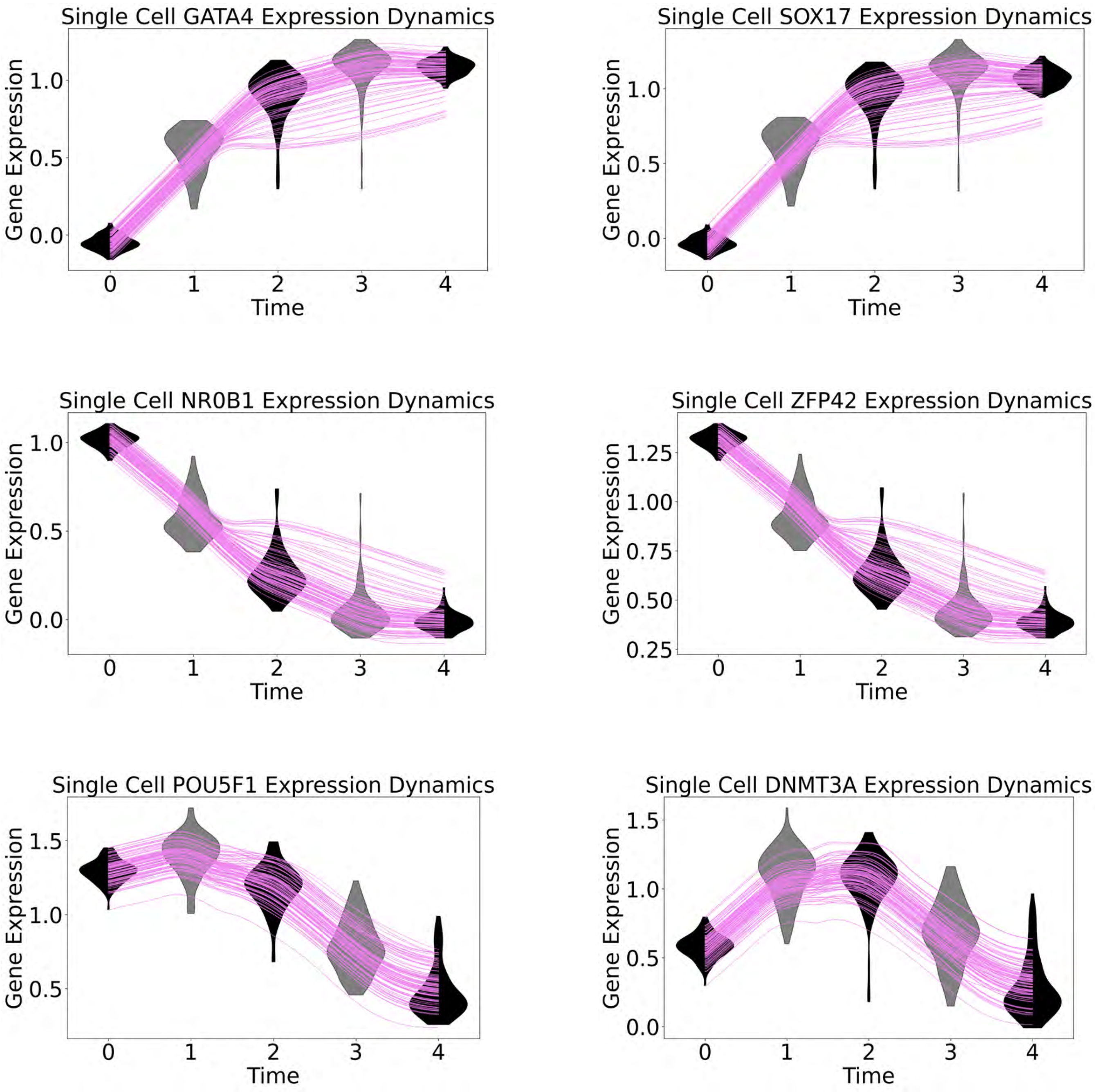

Single Cell GATA4 Expression Dynamics
Single Cell SOX17 Expression Dynamics
Single Cell NR0B1 Expression Dynamics
Single Cell ZFP42 Expression Dynamics
Single Cell POU5F1 Expression Dynamics
Single Cell DNMT3A Expression Dynamics
Gene Expression
Time

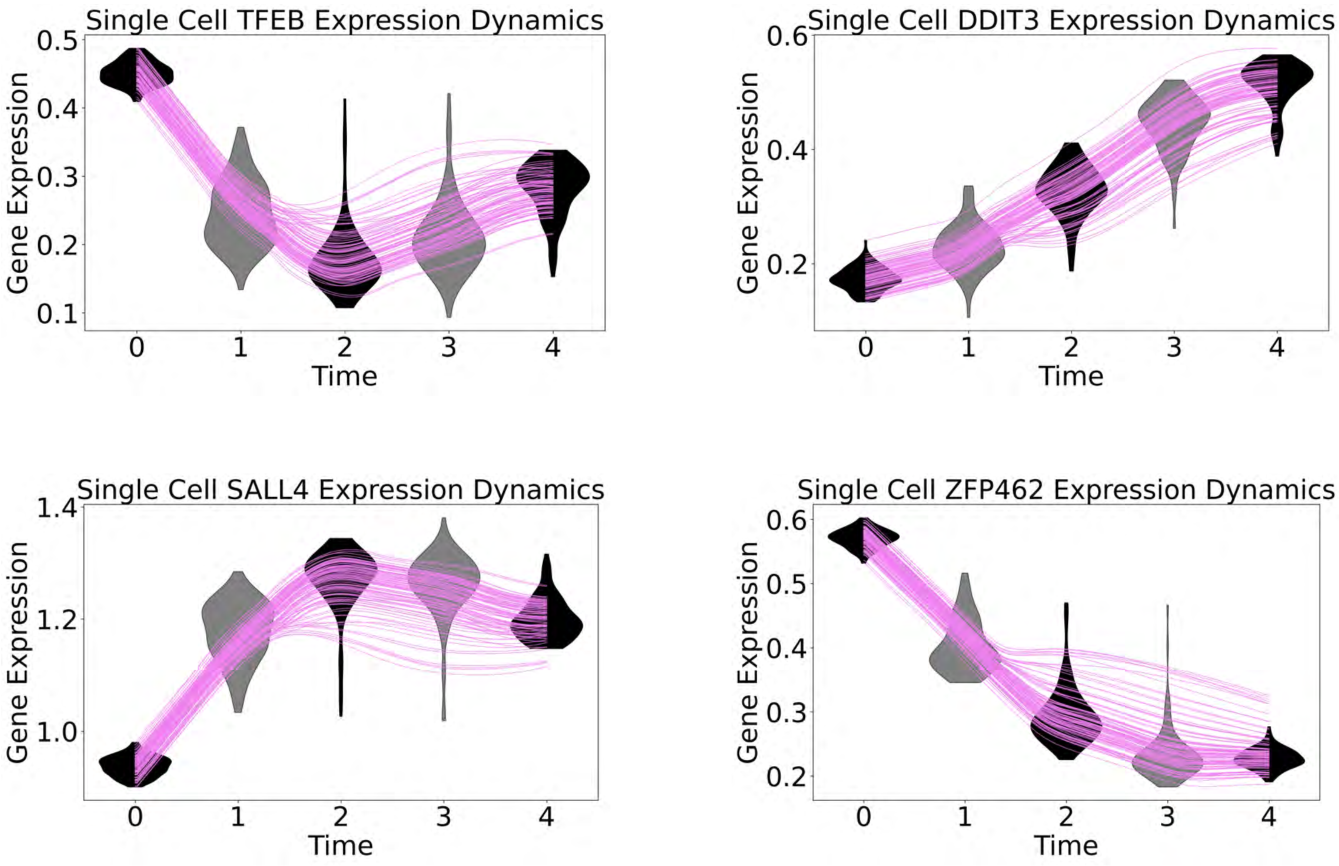

Figure S8, Reconstruction of single-cell, single-gene dynamics in the mESC dataset, related to Figure 3B. Single-cell, single-gene dynamics plotted as gene expression levels (y-axis) over time (x-axis), with each green curve representing an individual cell trajectory. The predicted trajectories are compared against real data distributions using violin plots (gray for test data, black for training data).

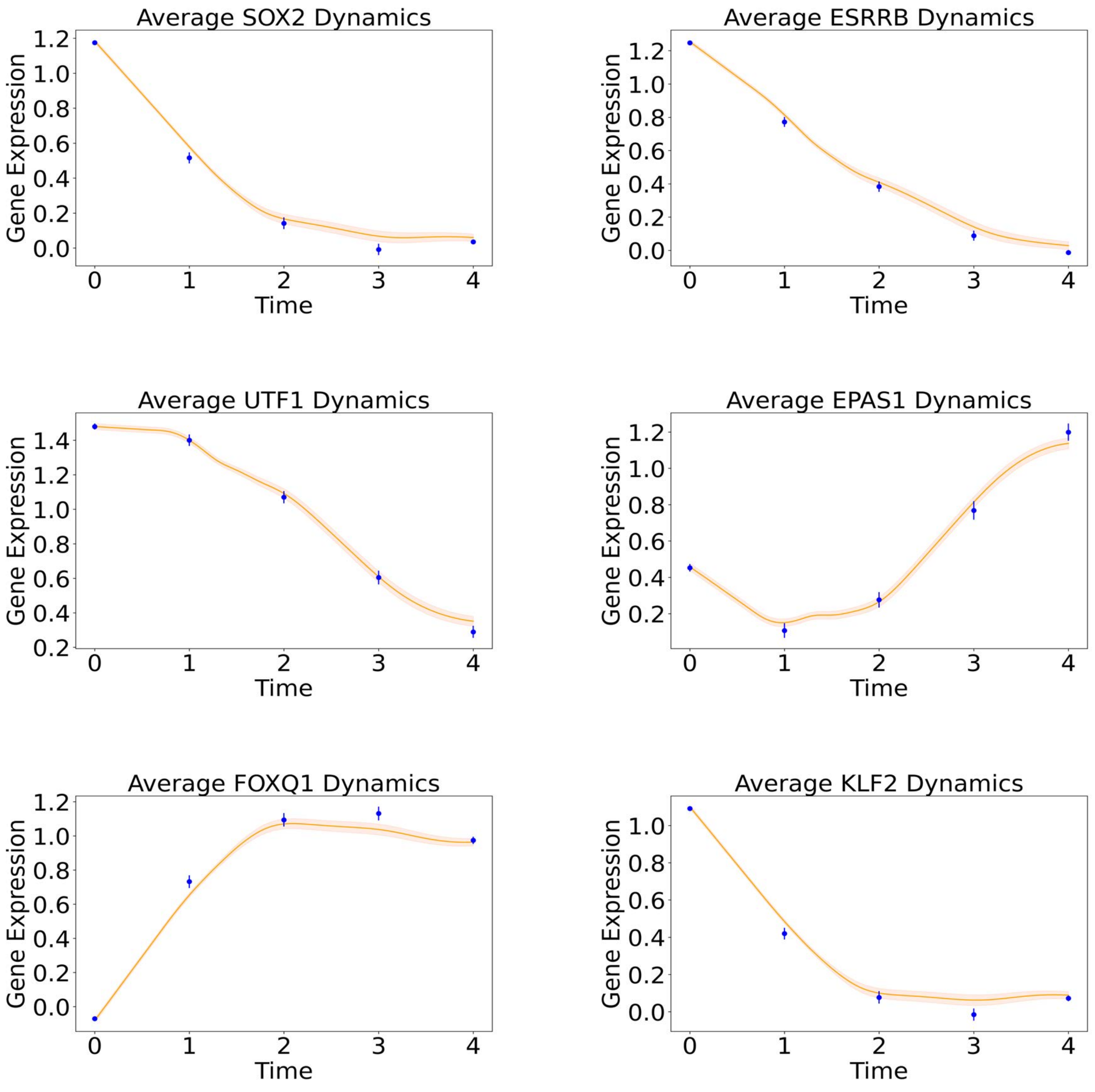
Average SOX2 Dynamics
Average ESRRB Dynamics
Average UTF1 Dynamics
Average EPAS1 Dynamics
Average FOXQ1 Dynamics
Average KLF2 Dynamics
Gene Expression
Time

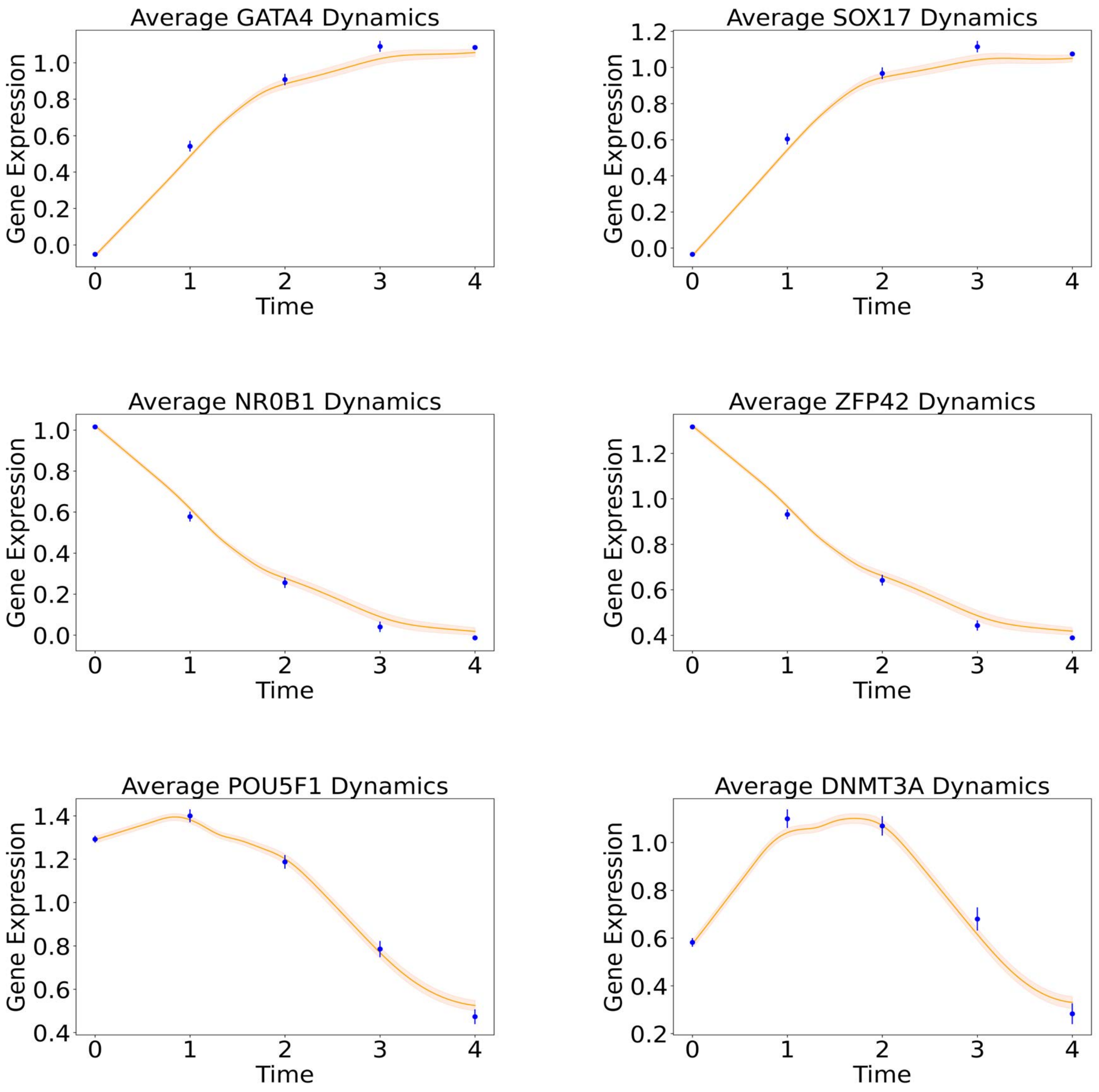

Average GATA4 Dynamics
Average SOX17 Dynamics
Average NR0B1 Dynamics
Average ZFP42 Dynamics
Average POU5F1 Dynamics
Average DNMT3A Dynamics
Gene Expression
Time

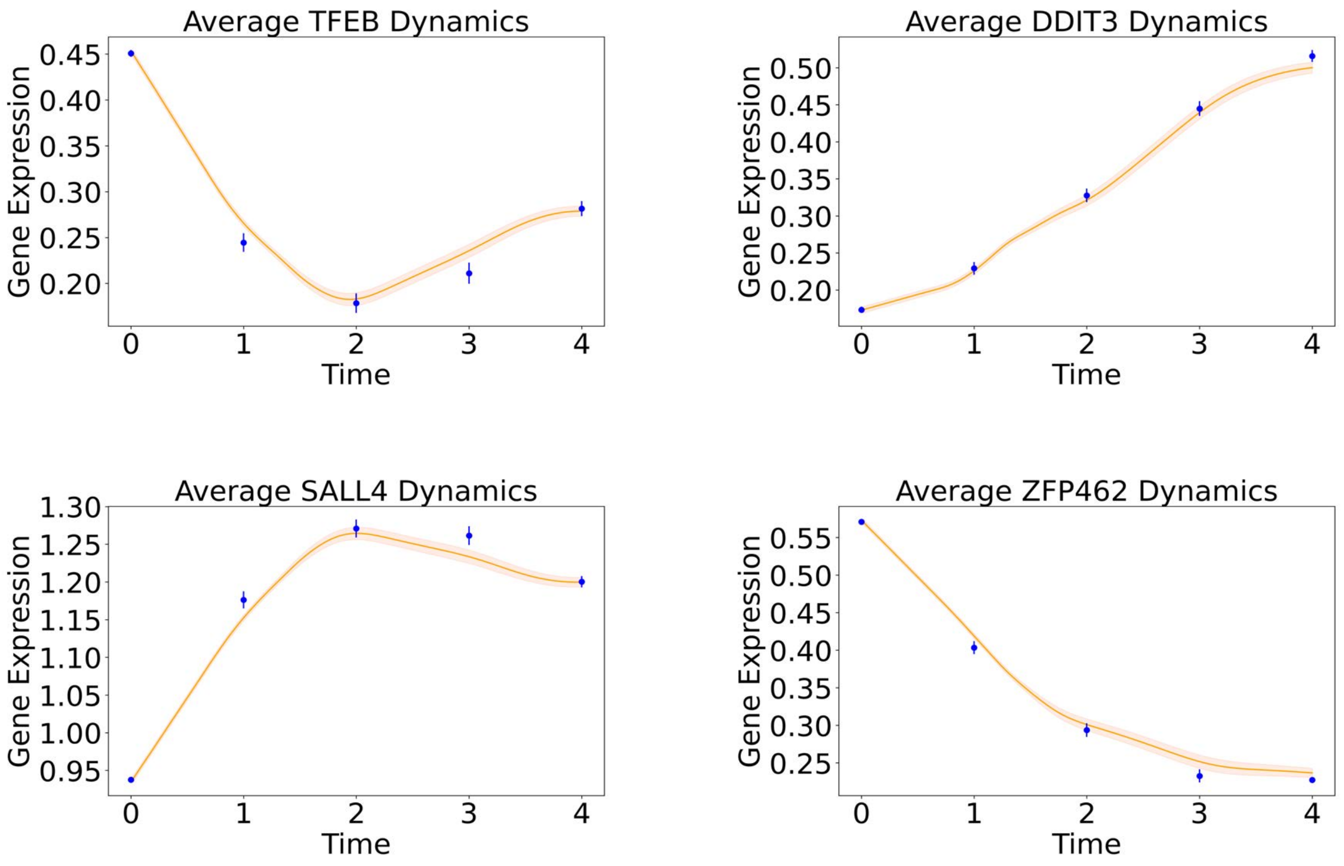

Figure S9, Predicted average gene expression trajectories in the mESC dataset, related to Figure 3C. Predicted average gene expression trajectories (orange curves with shaded regions indicating 95% confidence intervals) compared to actual gene expression levels from real data (blue dots with error bars).

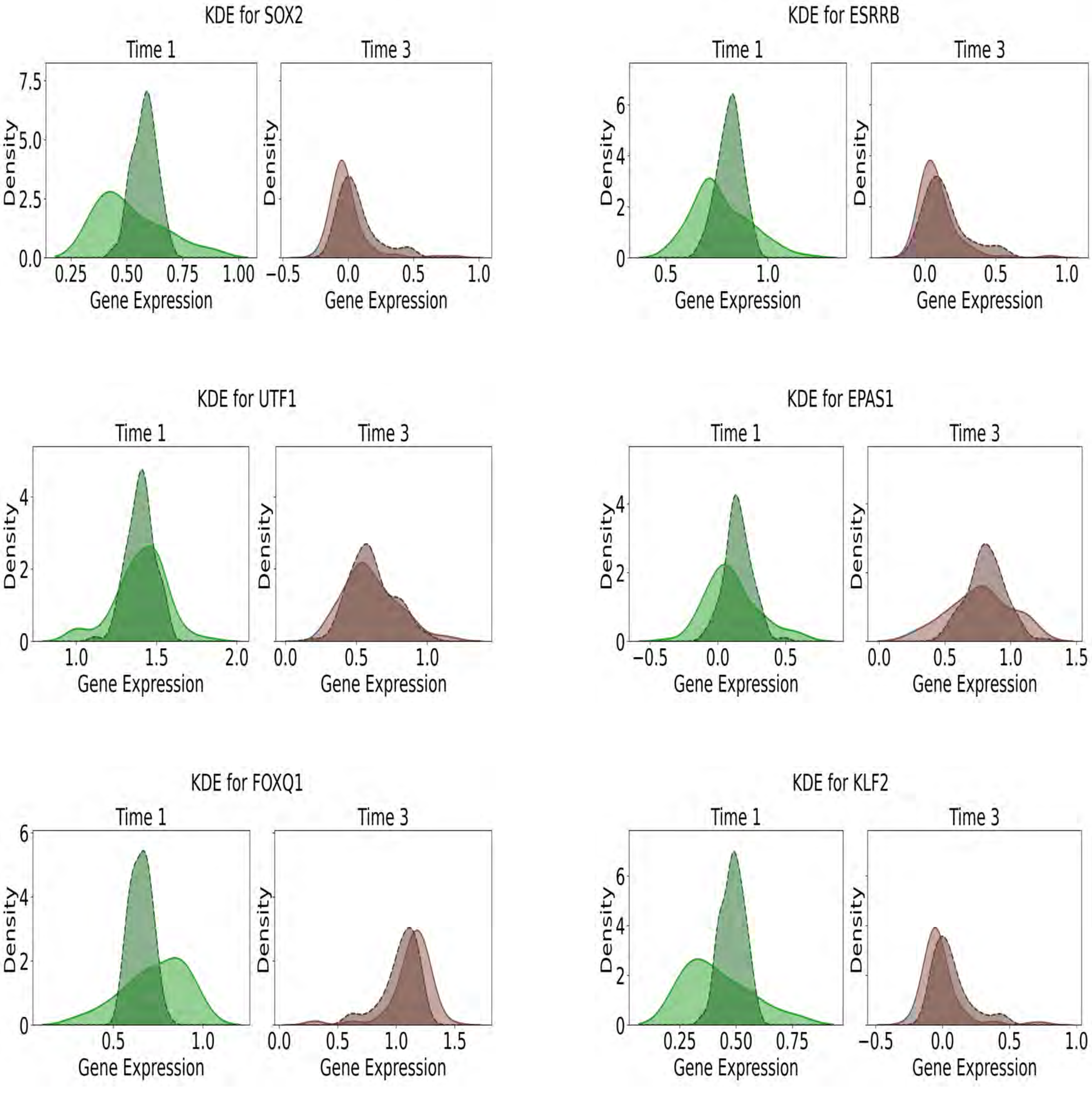

KDE for SOX2
Time 1
Time 3
Density
Gene Expression
KDE for ESRRB
Time 1
Time 3
Density
Gene Expression
KDE for UTF1
Time 1
Time 3
Density
Gene Expression
KDE for EPAS1
Time 1
Time 3
Density
Gene Expression
KDE for FOXQ1
Time 1
Time 3
Density
Gene Expression
KDE for KLF2
Time 1
Time 3
Density
Gene Expression

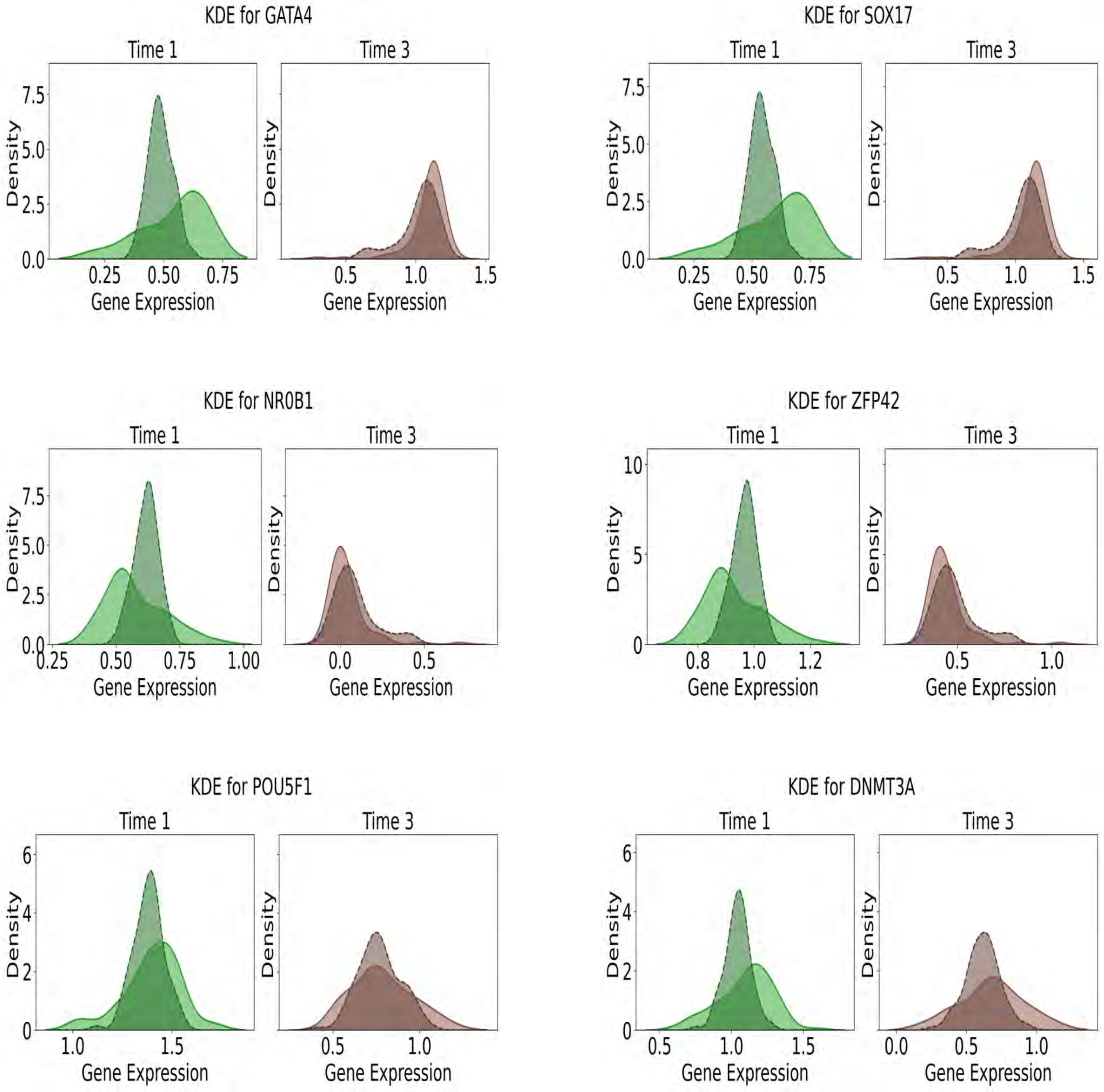
KDE for GATA4
Time 1
Time 3
Density
Gene Expression
0.0 2.5 5.0 7.5
0.25 0.50 0.75
0.5 1.0 1.5
KDE for SOX17
Time 1
Time 3
Density
Gene Expression
0.0 2.5 5.0 7.5
0.25 0.50 0.75
0.5 1.0 1.5
KDE for NR0B1
Time 1
Time 3
Density
Gene Expression
0.0 2.5 5.0 7.5
0.25 0.50 0.75 1.00
0.0 0.5
KDE for ZFP42
Time 1
Time 3
Density
Gene Expression
0 5 10
0.8 1.0 1.2
0.5 1.0
KDE for POU5F1
Time 1
Time 3
Density
Gene Expression
0 2 4 6
1.0 1.5
0.5 1.0
KDE for DNMT3A
Time 1
Time 3
Density
Gene Expression
0 2 4 6
0.5 1.0 1.5
0.0 0.5 1.0

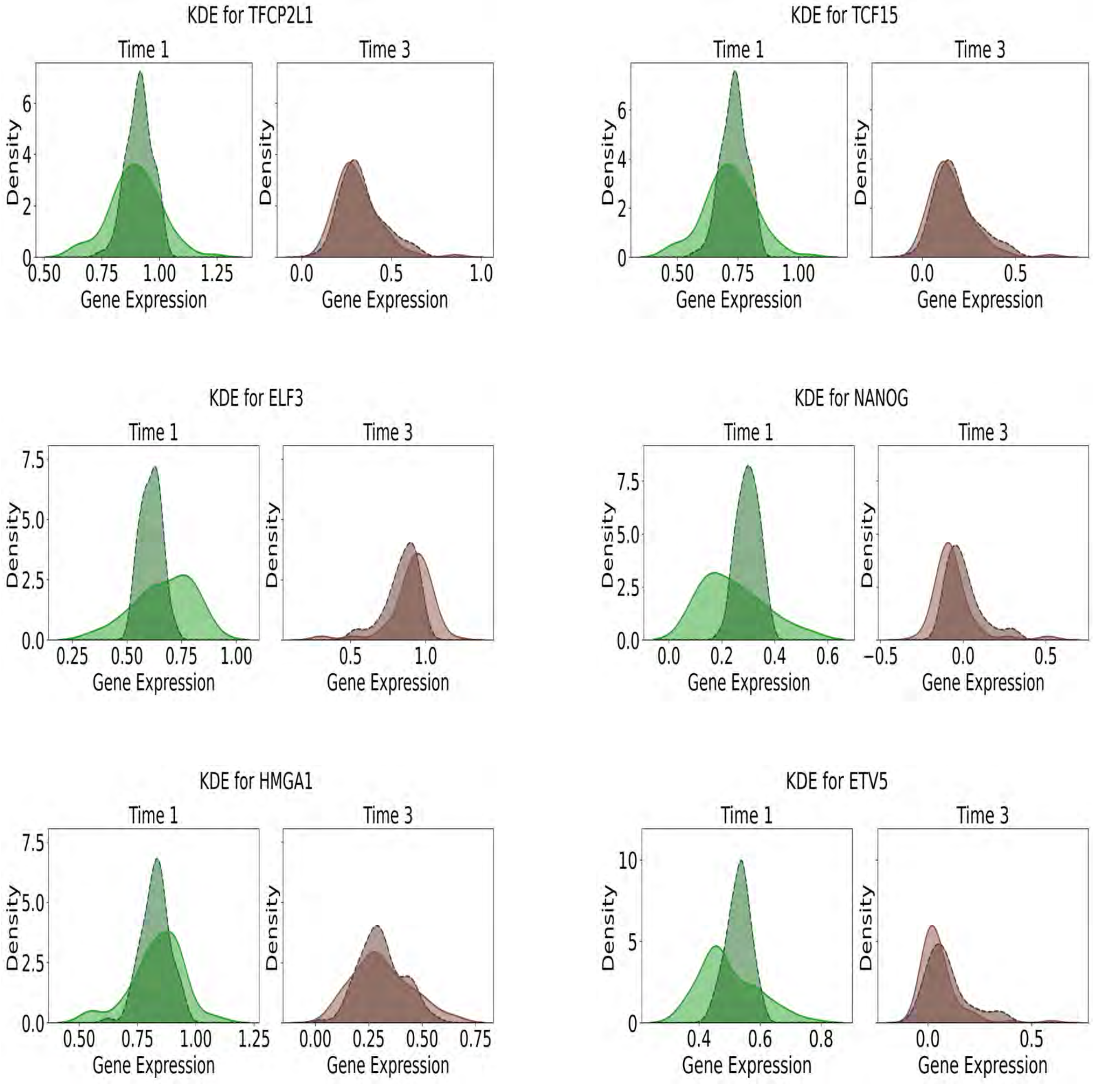

KDE for TFCP2L1
Time 1
Time 3
Density
Gene Expression
0.50 0.75 1.00 1.25
0.0 0.5 1.0
0 2 4 6
KDE for TCF15
Time 1
Time 3
Density
Gene Expression
0.50 0.75 1.00
0.0 0.5
0 2 4 6
KDE for ELF3
Time 1
Time 3
Density
Gene Expression
0.25 0.50 0.75 1.00
0.5 1.0
0.0 2.5 5.0 7.5
KDE for NANOG
Time 1
Time 3
Density
Gene Expression
0.0 0.2 0.4 0.6
−0.5 0.0 0.5
0.0 2.5 5.0 7.5
KDE for HMGA1
Time 1
Time 3
Density
Gene Expression
0.50 0.75 1.00 1.25
0.00 0.25 0.50 0.75
0.0 2.5 5.0 7.5
KDE for ETV5
Time 1
Time 3
Density
Gene Expression
0.4 0.6 0.8
0.0 0.5
0 5 10

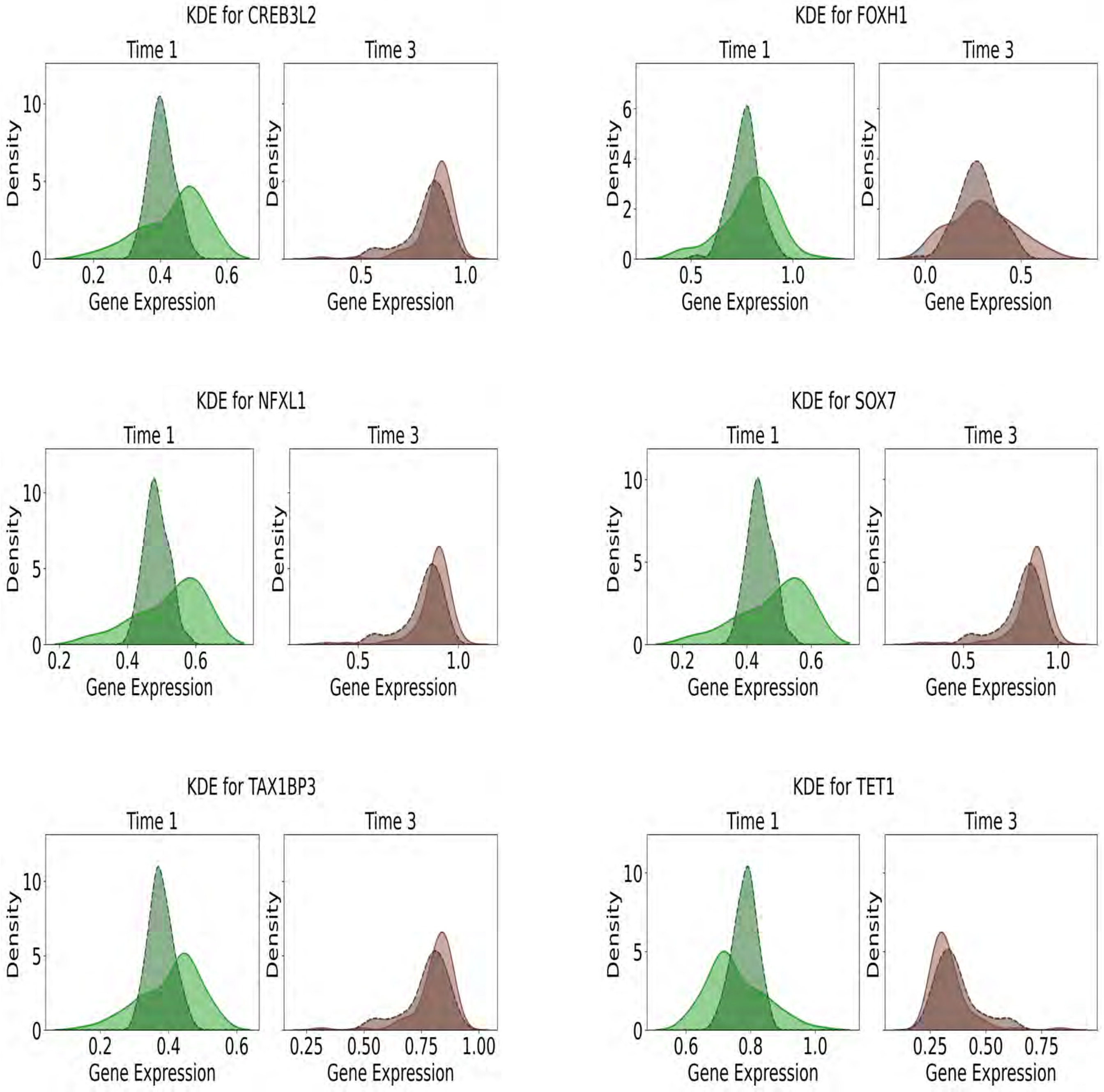

KDE for CREB3L2
Time 1
Time 3
Density
Gene Expression
KDE for FOXH1
Time 1
Time 3
Density
Gene Expression
KDE for NFXL1
Time 1
Time 3
Density
Gene Expression
KDE for SOX7
Time 1
Time 3
Density
Gene Expression
KDE for TAX1BP3
Time 1
Time 3
Density
Gene Expression
KDE for TET1
Time 1
Time 3
Density
Gene Expression

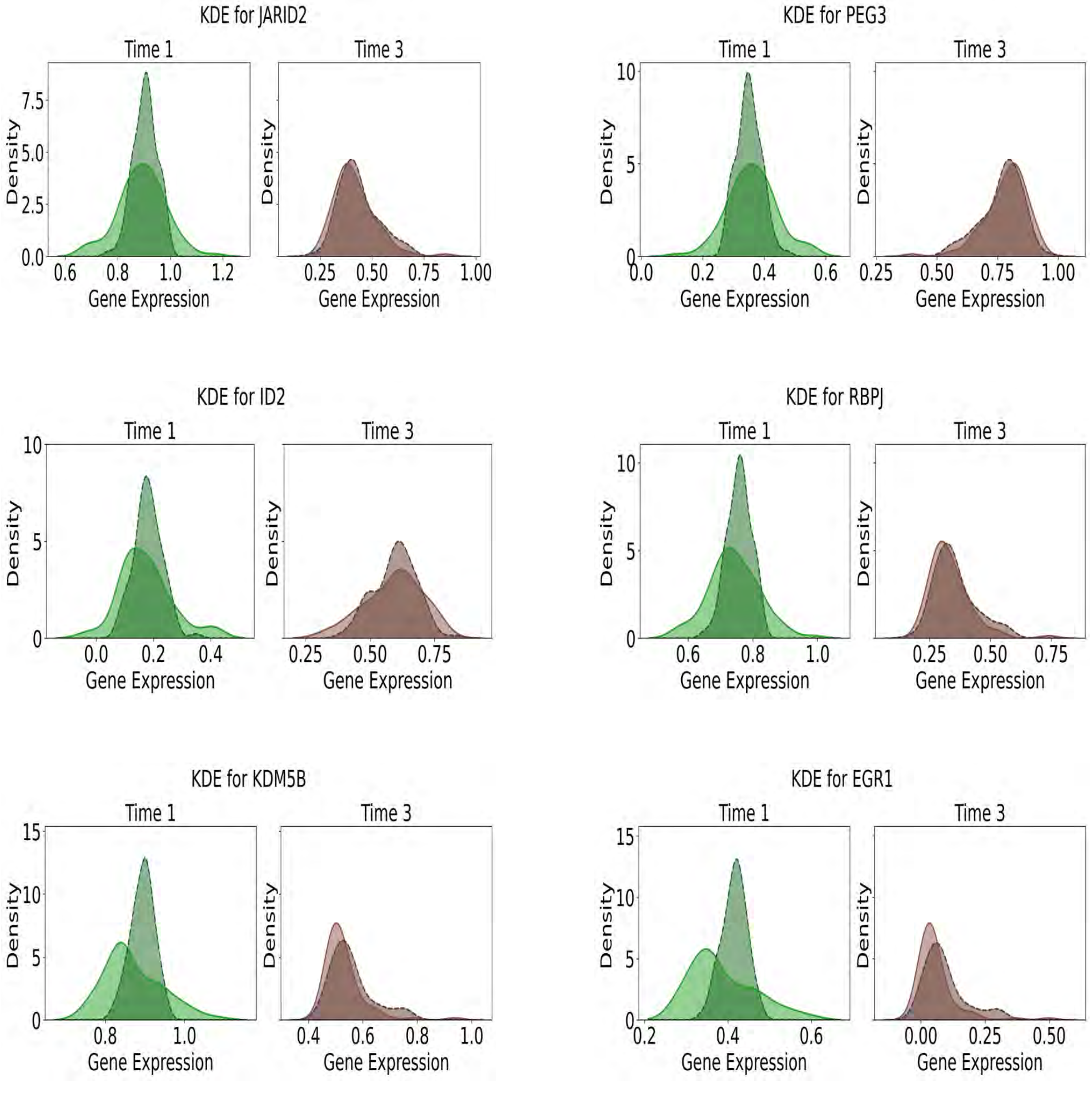

KDE for JARID2
Time 1
Time 3
Density
Gene Expression
KDE for PEG3
Time 1
Time 3
Density
Gene Expression
KDE for ID2
Time 1
Time 3
Density
Gene Expression
KDE for RBPJ
Time 1
Time 3
Density
Gene Expression
KDE for KDM5B
Time 1
Time 3
Density
Gene Expression
KDE for EGR1
Time 1
Time 3
Density
Gene Expression

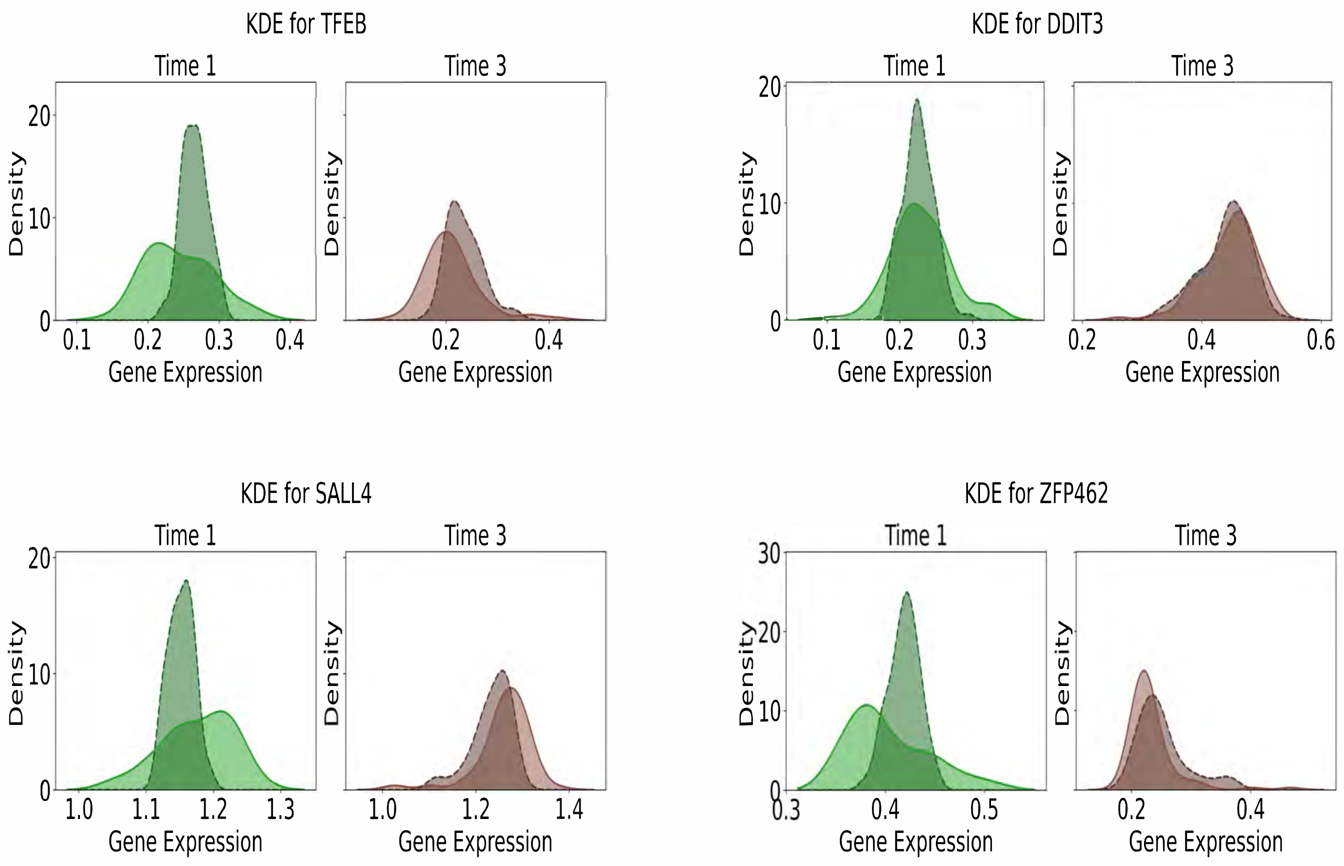

Figure S10, Comparison of predicted and observed gene expression distributions at held-out time points in the mESC dataset, related to Figure 3D. Comparison of predicted and actual gene expression distributions at time 1 (green) and time 3 (brown). Predicted distributions are shown as dashed lines, with real data distributions shown as solid lines

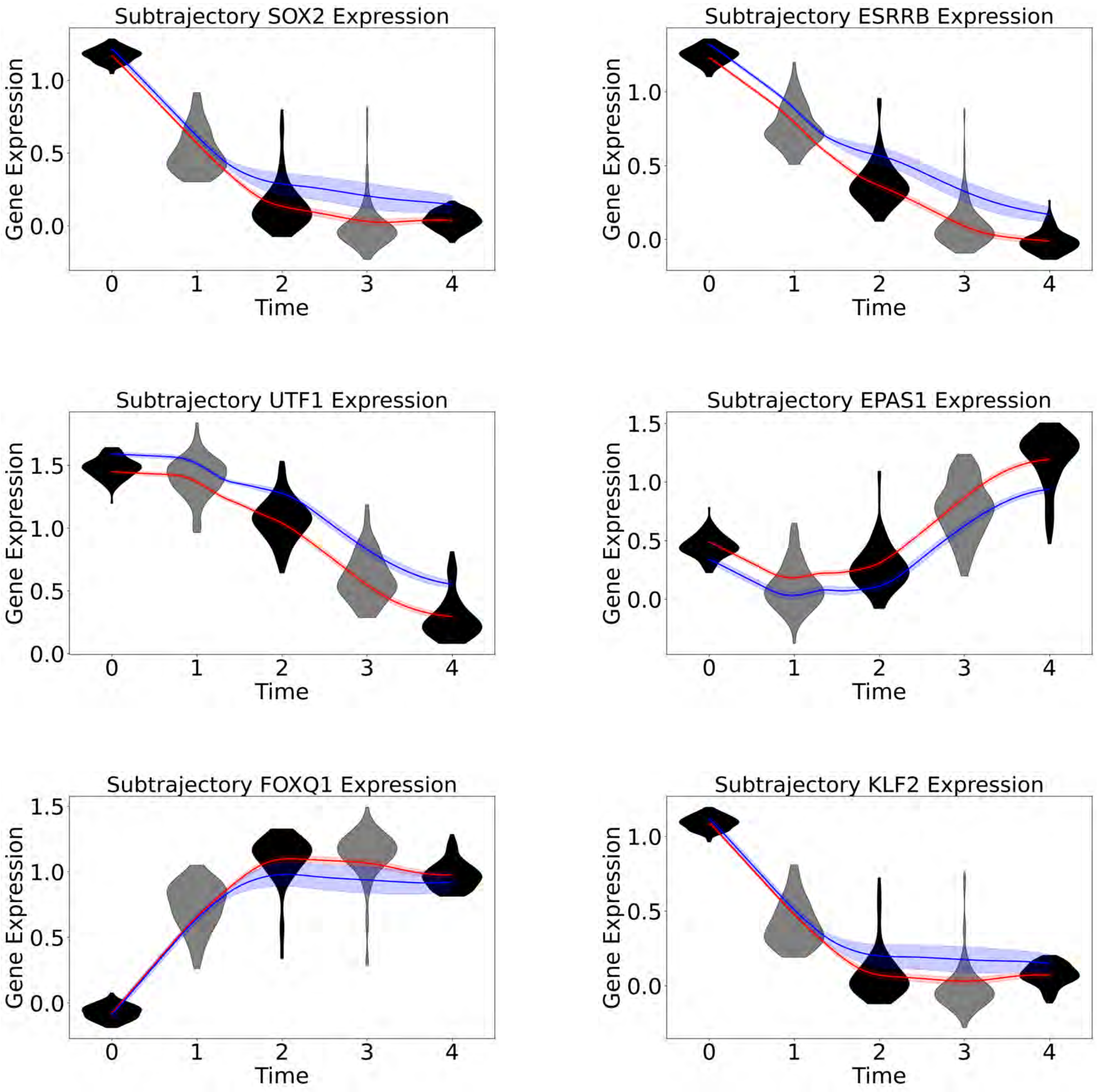

Subtrajectory SOX2 Expression
Subtrajectory ESRRB Expression
Subtrajectory UTF1 Expression
Subtrajectory EPAS1 Expression
Subtrajectory FOXQ1 Expression
Subtrajectory KLF2 Expression
Gene Expression
Time
0
1
2
3
4
0.0
0.5
1.0
1.5

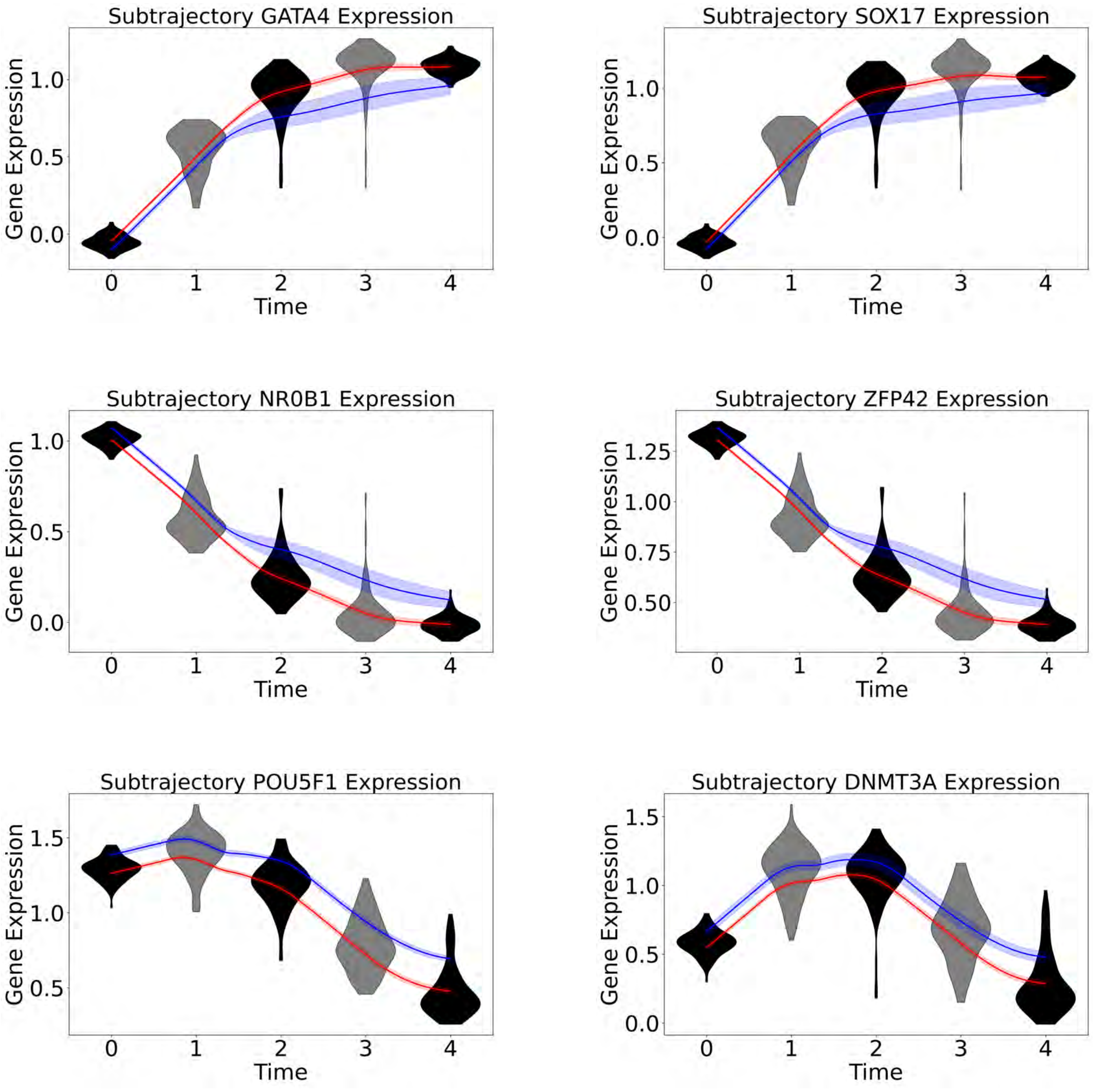
Subtrajectory GATA4 Expression
Subtrajectory SOX17 Expression
Subtrajectory NR0B1 Expression
Subtrajectory ZFP42 Expression
Subtrajectory POU5F1 Expression
Subtrajectory DNMT3A Expression
Gene Expression
Time

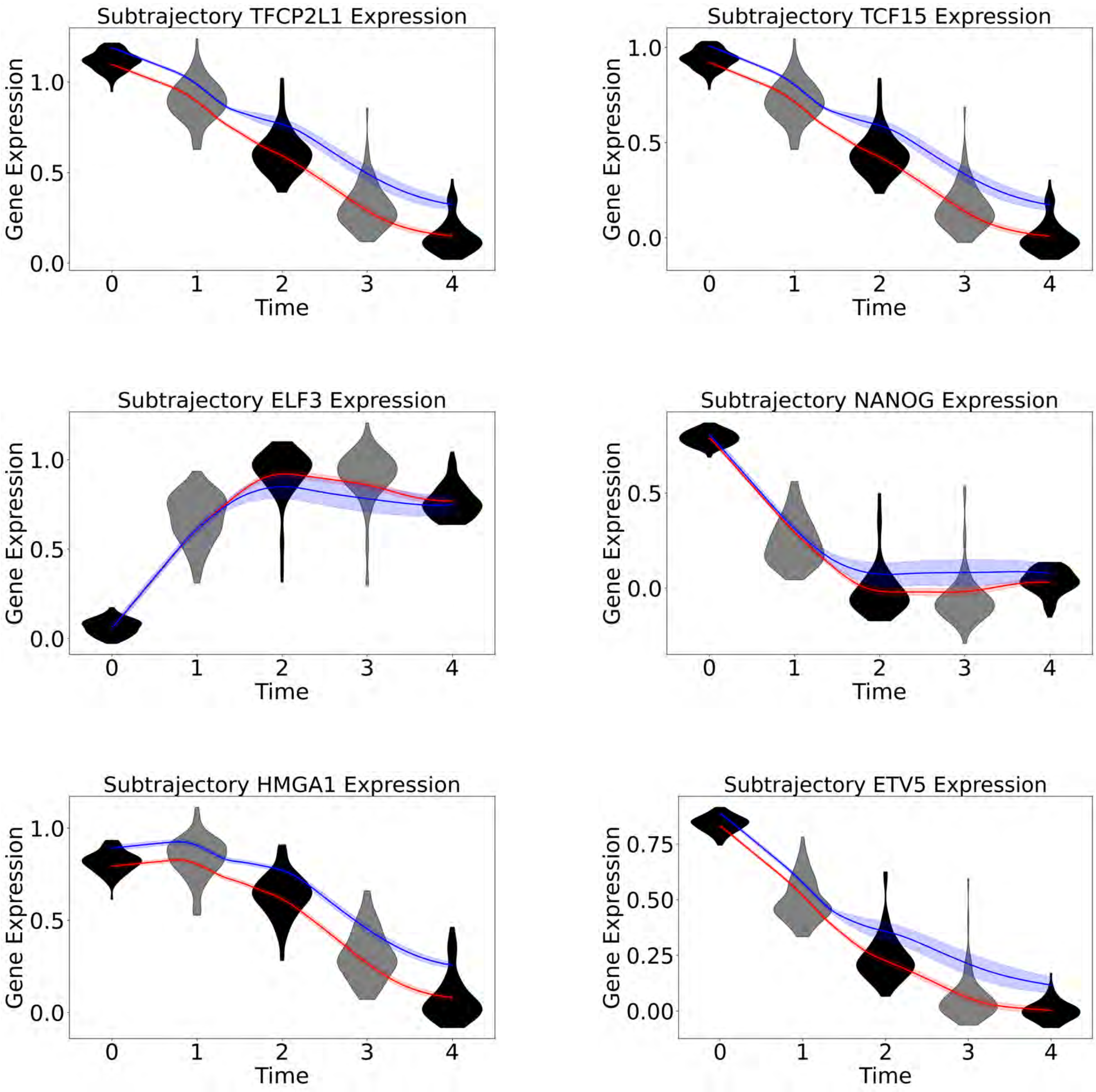
Subtrajectory TFCP2L1 Expression
Gene Expression
0.0
0.5
1.0
0
1
2
3
4
Time
Subtrajectory TCF15 Expression
Gene Expression
0.0
0.5
1.0
0
1
2
3
4
Time
Subtrajectory ELF3 Expression
Gene Expression
0.0
0.5
1.0
0
1
2
3
4
Time
Subtrajectory NANOG Expression
Gene Expression
0.0
0.5
0
1
2
3
4
Time
Subtrajectory HMGA1 Expression
Gene Expression
0.0
0.5
1.0
0
1
2
3
4
Time
Subtrajectory ETV5 Expression
Gene Expression
0.00
0.25
0.50
0.75
0
1
2
3
4
Time

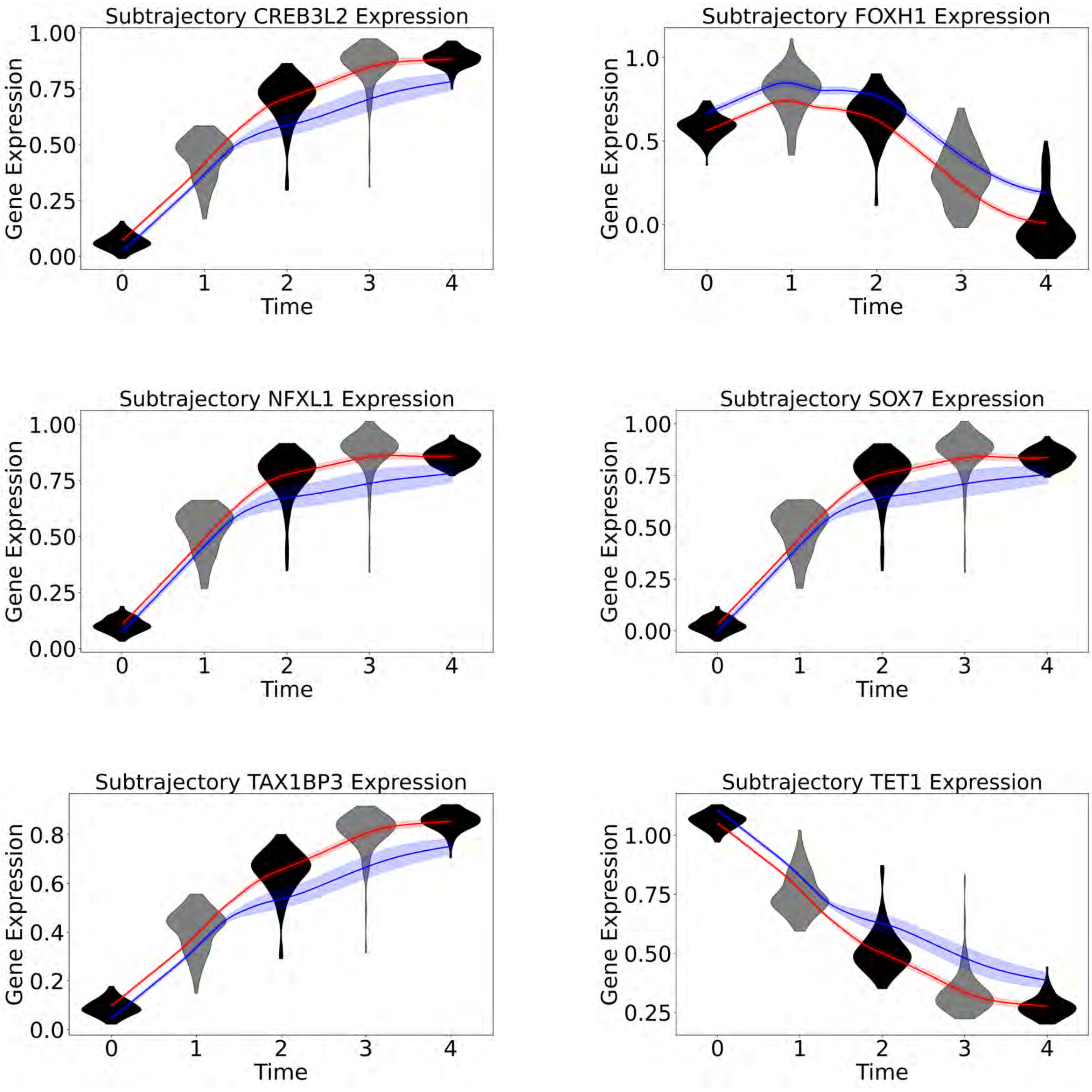
Subtrajectory CREB3L2 Expression
Gene Expression
Time
Subtrajectory FOXH1 Expression
Gene Expression
Time
Subtrajectory NFXL1 Expression
Gene Expression
Time
Subtrajectory SOX7 Expression
Gene Expression
Time
Subtrajectory TAX1BP3 Expression
Gene Expression
Time
Subtrajectory TET1 Expression
Gene Expression
Time

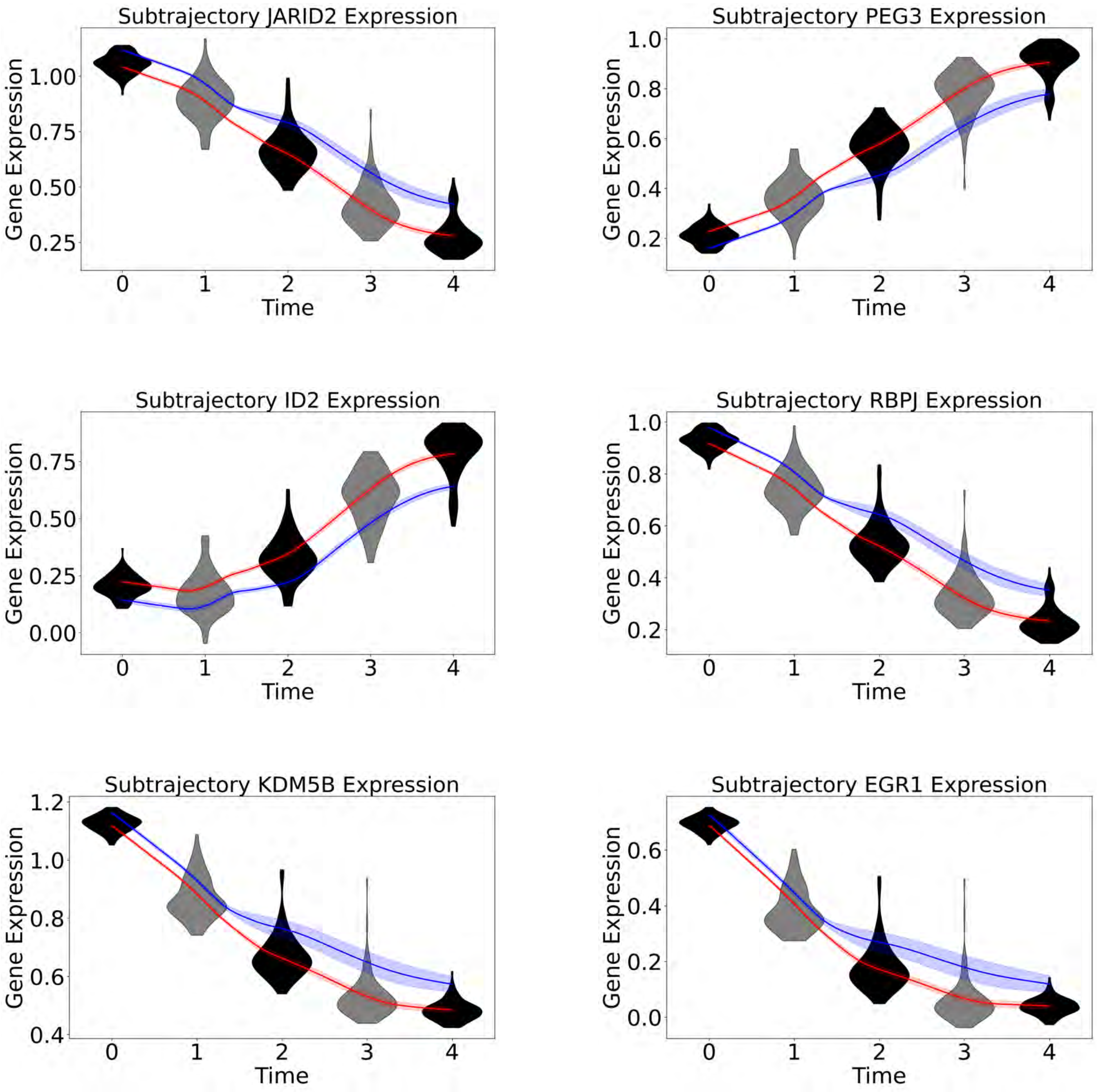

Subtrajectory JARID2 Expression
Gene Expression
0.25 0.50 0.75 1.00
0 1 2 3 4
Time
Subtrajectory PEG3 Expression
Gene Expression
0.2 0.4 0.6 0.8 1.0
0 1 2 3 4
Time
Subtrajectory ID2 Expression
Gene Expression
0.00 0.25 0.50 0.75
0 1 2 3 4
Time
Subtrajectory RBPJ Expression
Gene Expression
0.2 0.4 0.6 0.8 1.0
0 1 2 3 4
Time
Subtrajectory KDM5B Expression
Gene Expression
0.4 0.6 0.8 1.0 1.2
0 1 2 3 4
Time
Subtrajectory EGR1 Expression
Gene Expression
0.0 0.2 0.4 0.6
0 1 2 3 4
Time

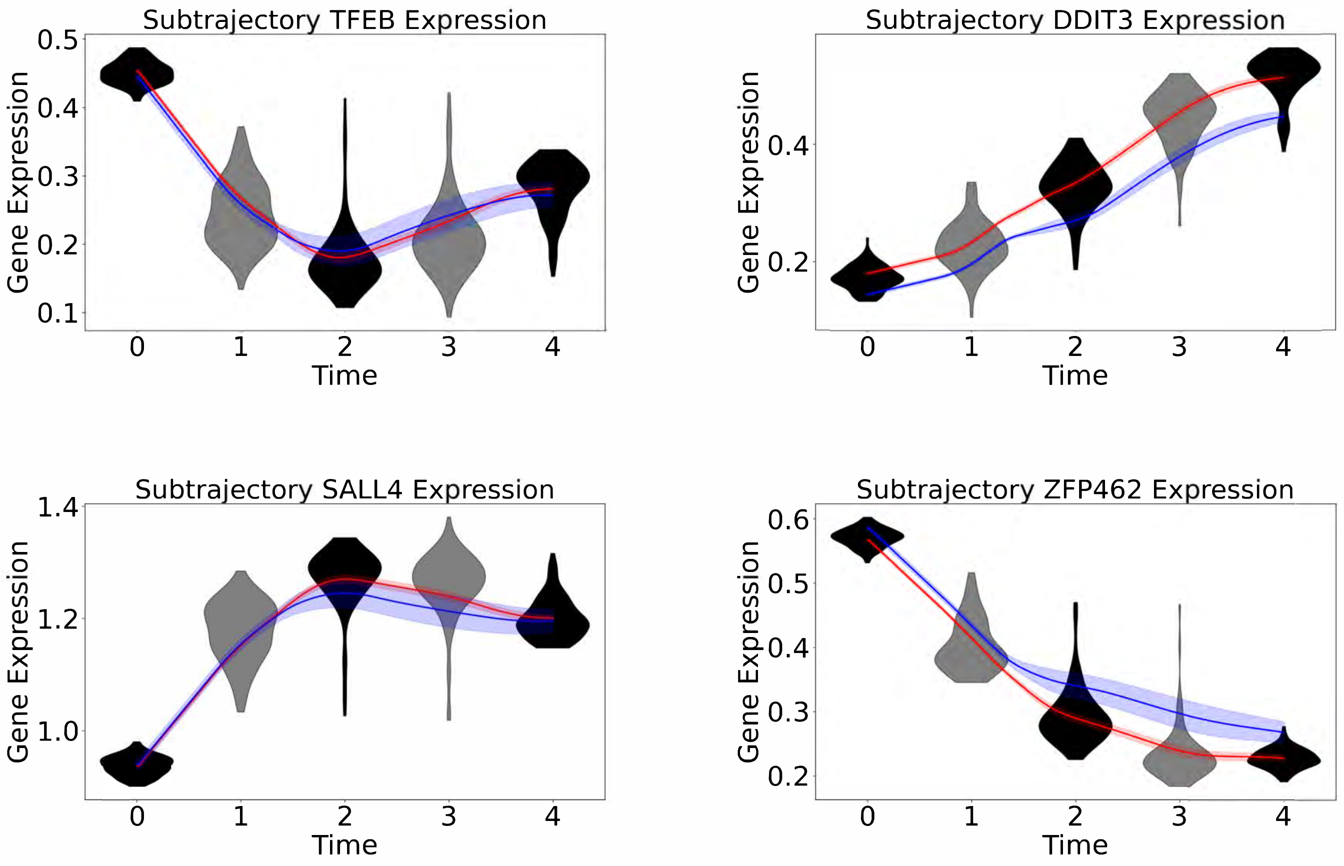

Figure S11, Average gene expression trajectories for each identified subgroup in the mESC dataset, related to Figure 3G. Average gene expression trajectories plotted separately for each subgroup (shaded regions indicate 95% confidence intervals).

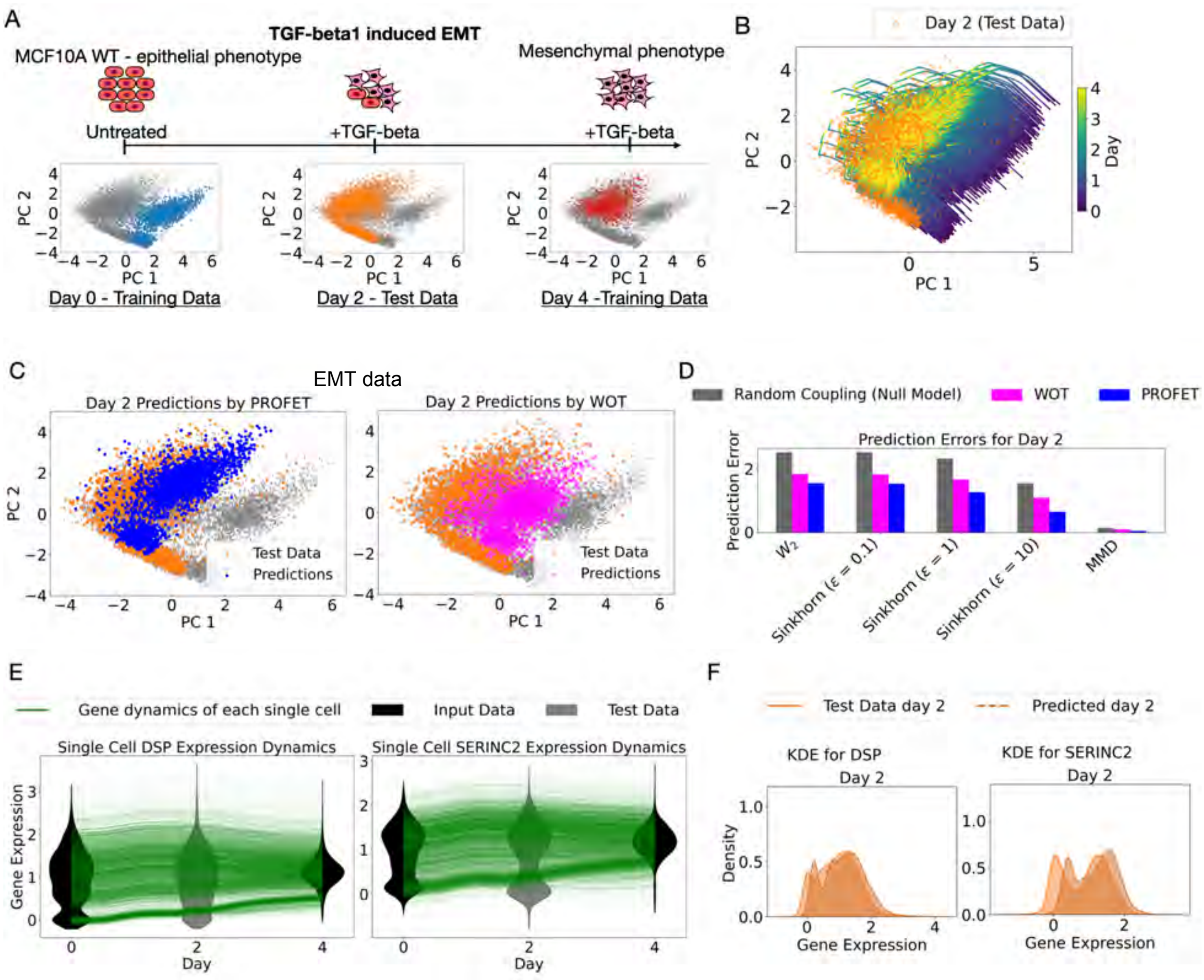

Figure S12, Downstream analysis of reconstructed single-cell trajectories for EMT data, related to Figure 4AB. (A) Illustration of EMT scRNA-seq data. (B) Application of PROFET to reconstruct continuous trajectories using only data from time points 0 and 4, with time point 2 held out for validation. The colormap represents the predicted time along the trajectory, ranging from time 0 to time 4, and the test data is highlighted by orange circles. (C) Comparison of test data predictions (time 2) between the PROFET (blue dots) and the WOT method (magenta crosses). (D) Quantitative comparison of prediction errors at time 2 across three methods—PROFET (blue), WOT (magenta), and random coupling (black)—measured using W2, Sinkhorn (entropy = 0.1, 1, 10), and MMD metrics. (E) Single-cell, single-gene dynamics, showing gene expression levels (y-axis) over time (x-axis). Each green curve represents an individual cell trajectory, with predicted trajectories compared against real data distributions using violin plots (gray for test data, black for training data). (F) Comparison of predicted and actual gene expression distributions at time 2. Predicted distributions are shown as dashed lines, while real data distributions are shown as solid lines.

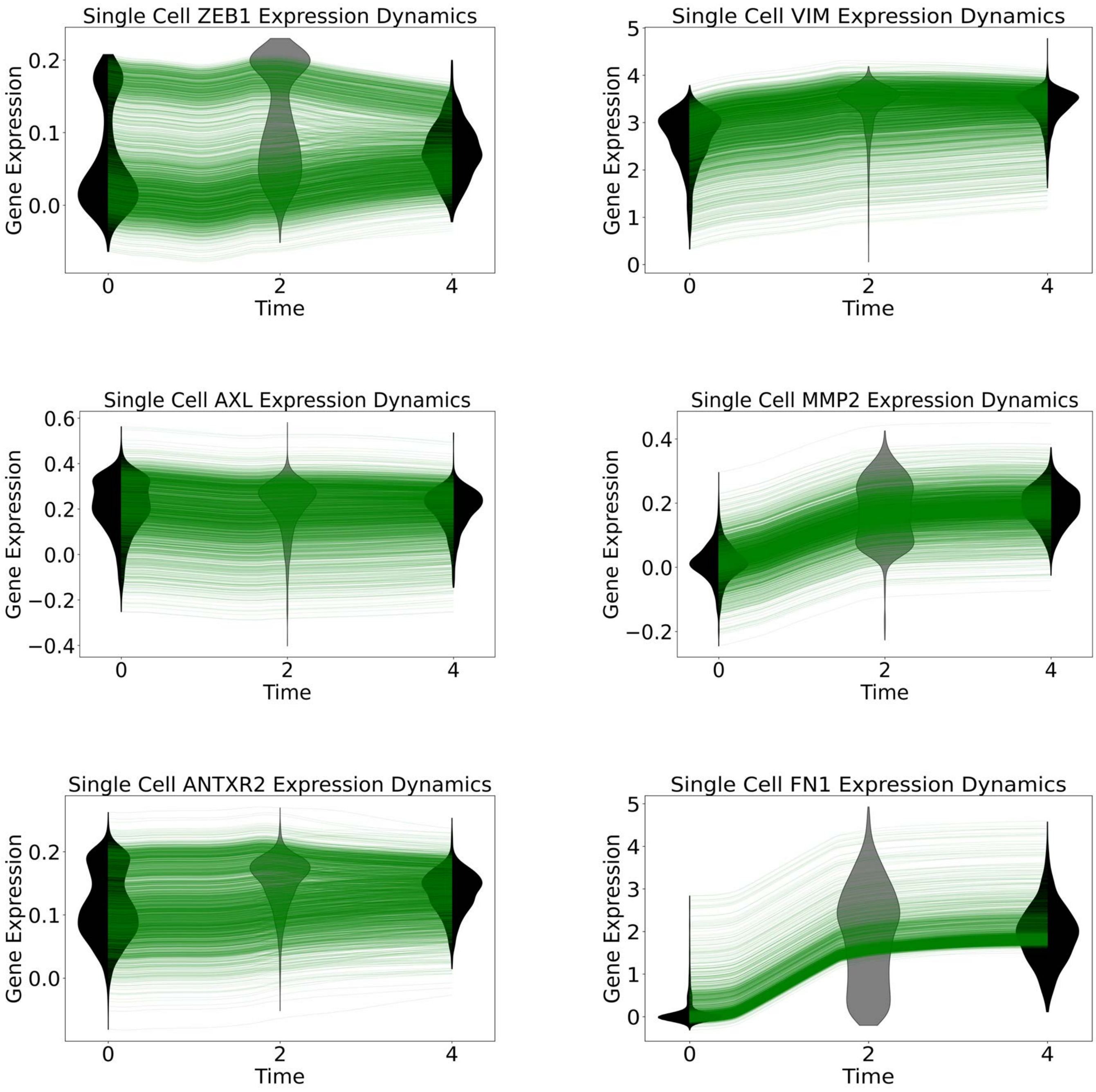

Single Cell ZEB1 Expression Dynamics
Gene Expression
0.2
0.1
0.0
0
2
4
Time
Single Cell VIM Expression Dynamics
Gene Expression
5
4
3
2
1
0
0
2
4
Time
Single Cell AXL Expression Dynamics
Gene Expression
0.6
0.4
0.2
0.0
−0.2
−0.4
0
2
4
Time
Single Cell MMP2 Expression Dynamics
Gene Expression
0.4
0.2
0.0
−0.2
0
2
4
Time
Single Cell ANTXR2 Expression Dynamics
Gene Expression
0.2
0.1
0.0
0
2
4
Time
Single Cell FN1 Expression Dynamics
Gene Expression
5
4
3
2
1
0
0
2
4
Time

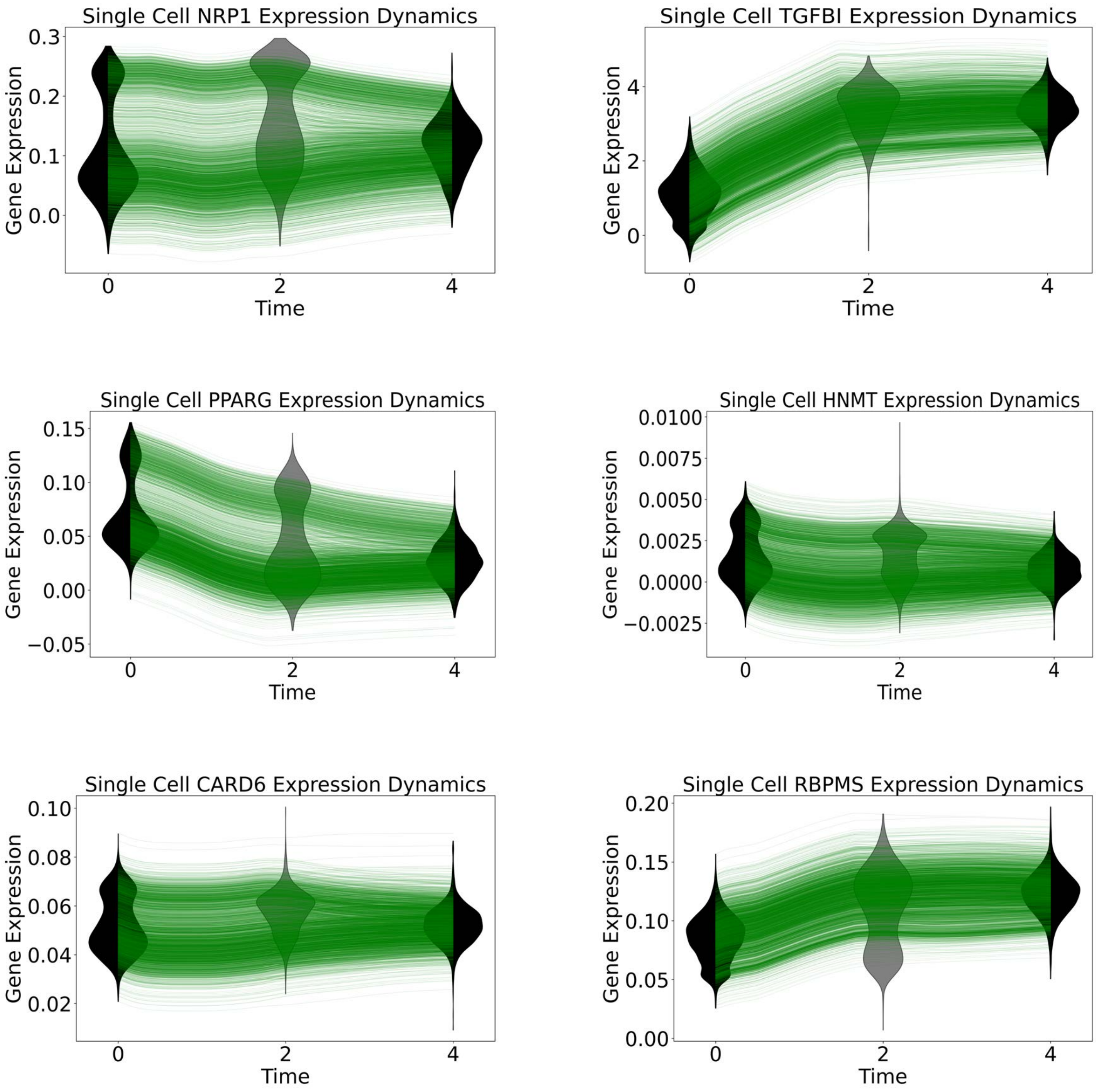

Single Cell NRP1 Expression Dynamics
Gene Expression
Time
Single Cell TGFBI Expression Dynamics
Gene Expression
Time
Single Cell PPARG Expression Dynamics
Gene Expression
Time
Single Cell HNMT Expression Dynamics
Gene Expression
Time
Single Cell CARD6 Expression Dynamics
Gene Expression
Time
Single Cell RBPMS Expression Dynamics
Gene Expression
Time

Figure S13, Reconstruction of single-cell, single-gene dynamics in the EMT dataset, related to Figure 4B. Single-cell, single-gene dynamics plotted as gene expression levels (y-axis) over time (x-axis), with each green curve representing an individual cell trajectory. The predicted trajectories are compared against real data distributions using violin plots (gray for test data, black for training data).

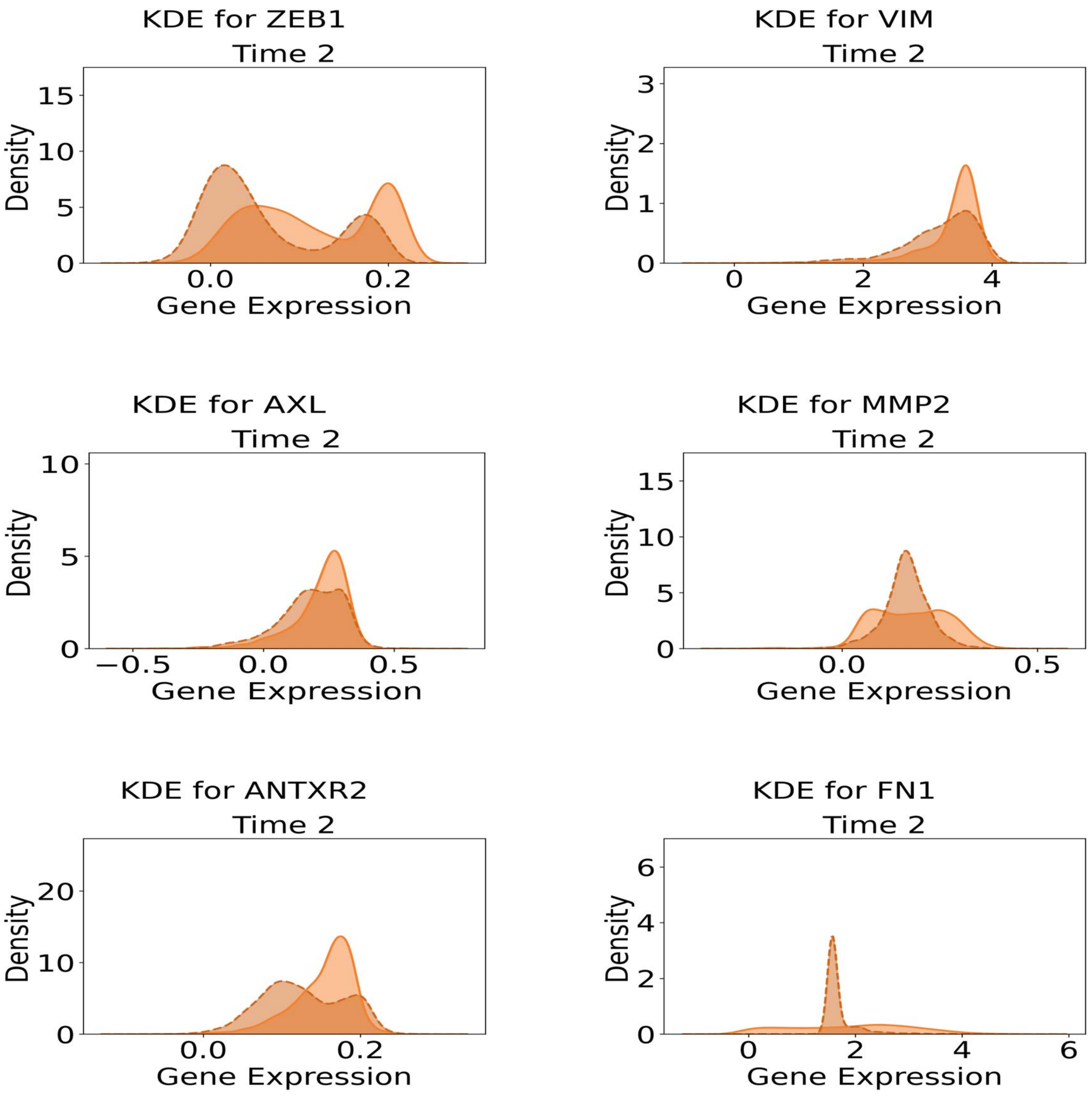
KDE for ZEB1
Time 2
Density
Gene Expression
KDE for VIM
Time 2
Density
Gene Expression
KDE for AXL
Time 2
Density
Gene Expression
KDE for MMP2
Time 2
Density
Gene Expression
KDE for ANTXR2
Time 2
Density
Gene Expression
KDE for FN1
Time 2
Density
Gene Expression

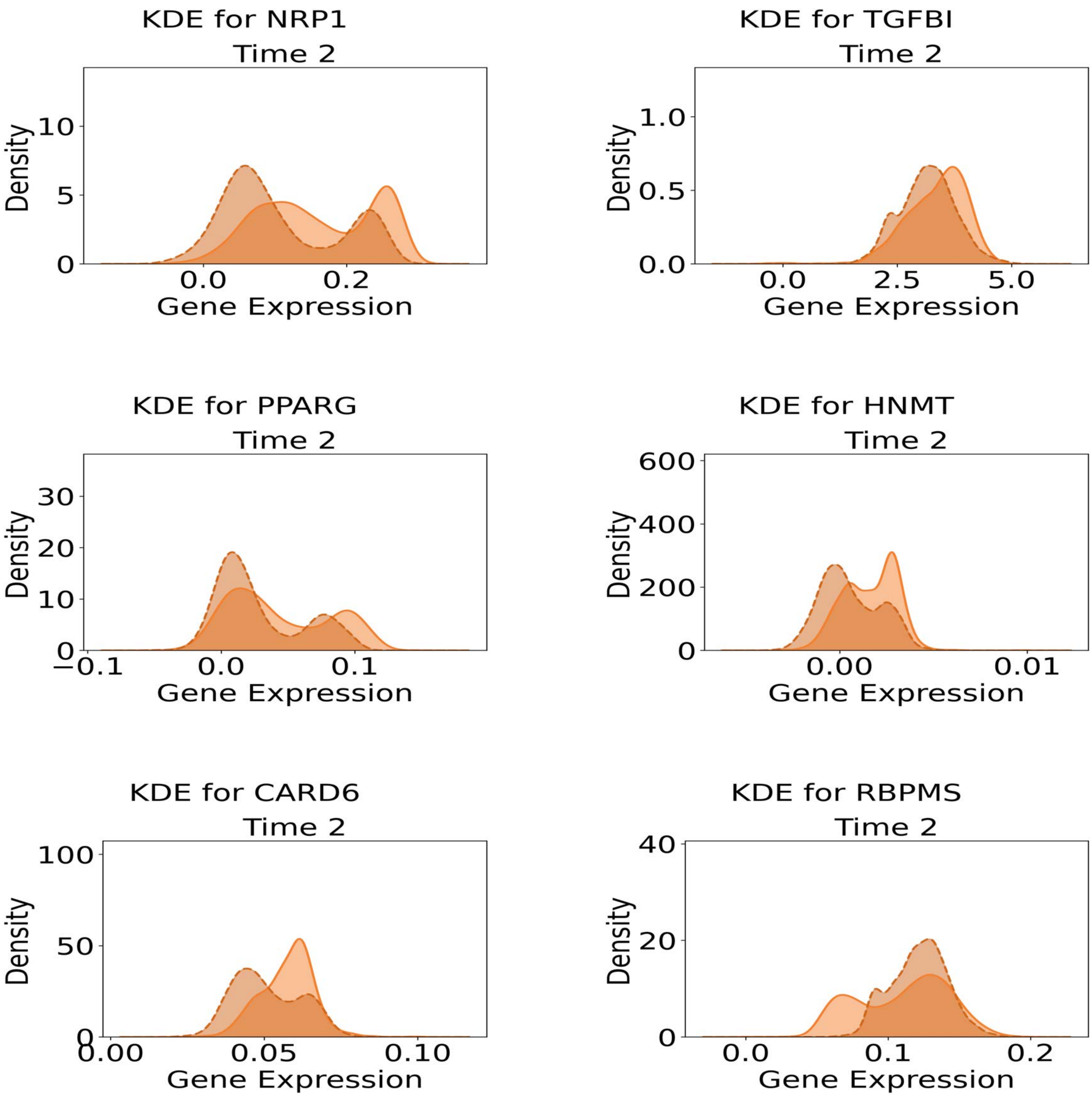
KDE for NRP1
Time 2
Density
Gene Expression
KDE for TGFBI
Time 2
Density
Gene Expression
KDE for PPARG
Time 2
Density
Gene Expression
KDE for HNMT
Time 2
Density
Gene Expression
KDE for CARD6
Time 2
Density
Gene Expression
KDE for RBPMS
Time 2
Density
Gene Expression

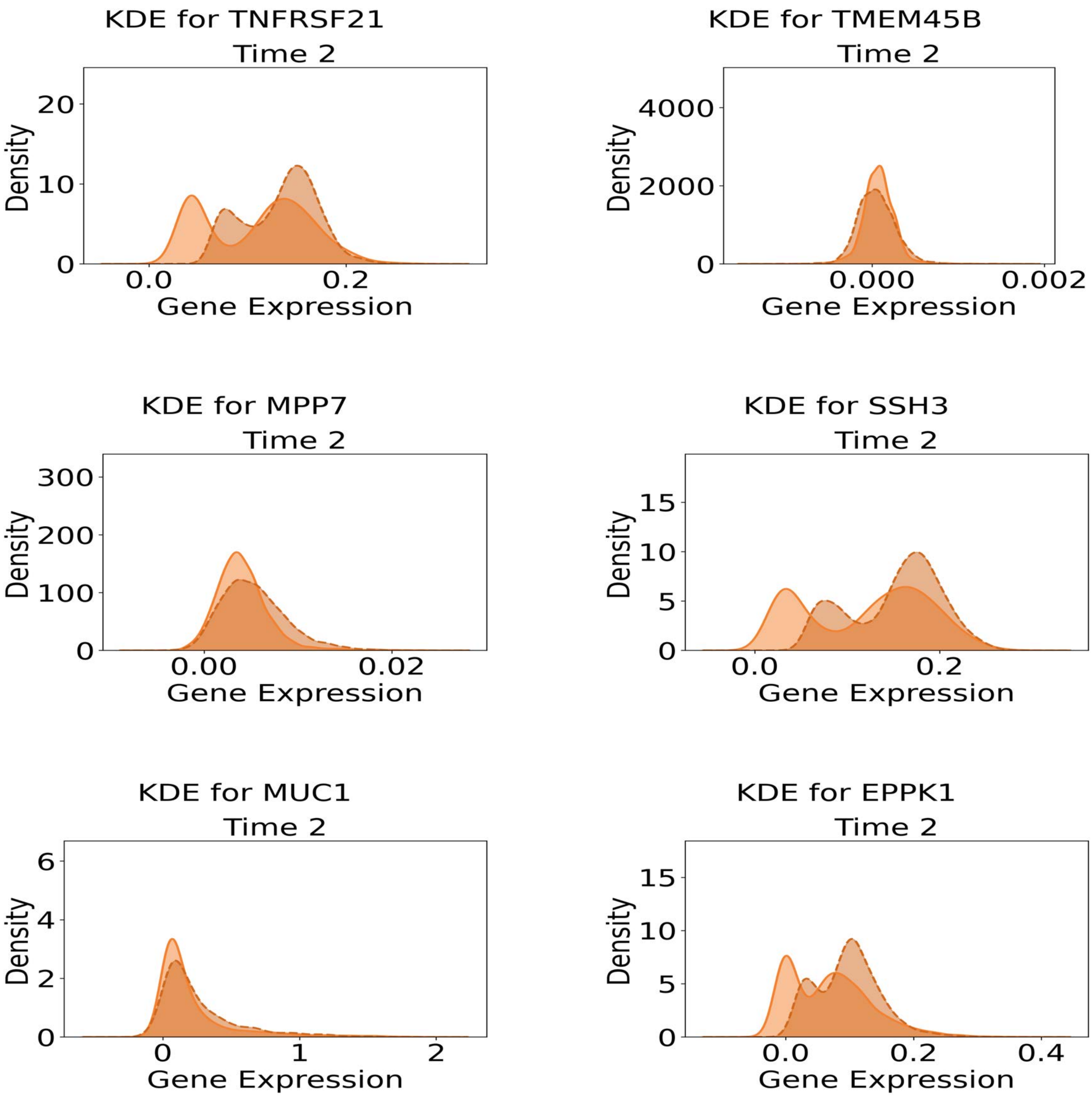

KDE for TNFRSF21
Time 2
Density
0
10
20
0.0
0.2
Gene Expression
KDE for TMEM45B
Time 2
Density
0
2000
4000
0.000
0.002
Gene Expression
KDE for MPP7
Time 2
Density
0
100
200
300
0.00
0.02
Gene Expression
KDE for SSH3
Time 2
Density
0
5
10
15
0.0
0.2
Gene Expression
KDE for MUC1
Time 2
Density
0
2
4
6
0
1
2
Gene Expression
KDE for EPPK1
Time 2
Density
0
5
10
15
0.0
0.2
0.4
Gene Expression

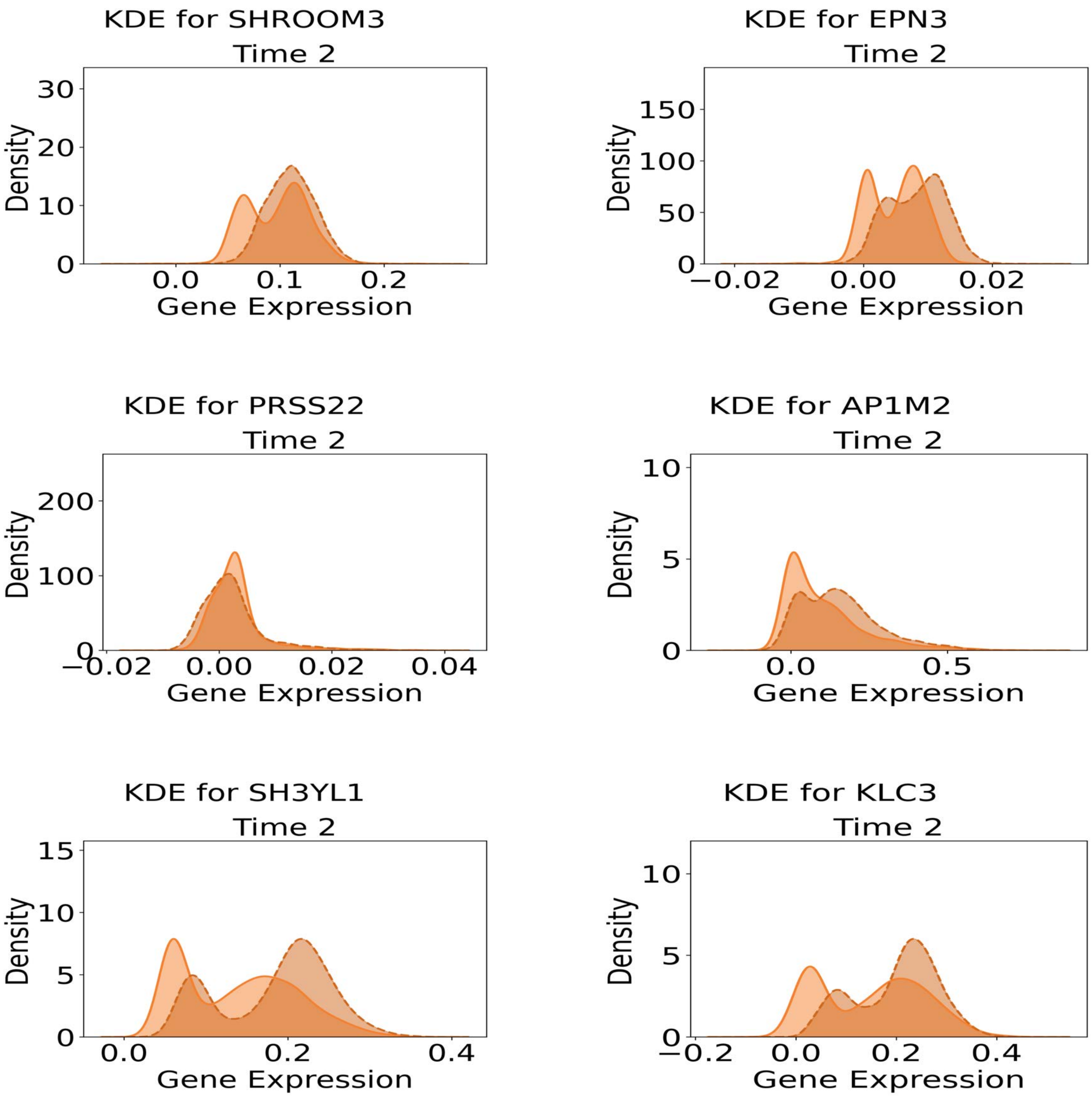
KDE for SHROOM3
Time 2
Density
Gene Expression
KDE for EPN3
Time 2
Density
Gene Expression
KDE for PRSS22
Time 2
Density
Gene Expression
KDE for AP1M2
Time 2
Density
Gene Expression
KDE for SH3YL1
Time 2
Density
Gene Expression
KDE for KLC3
Time 2
Density
Gene Expression

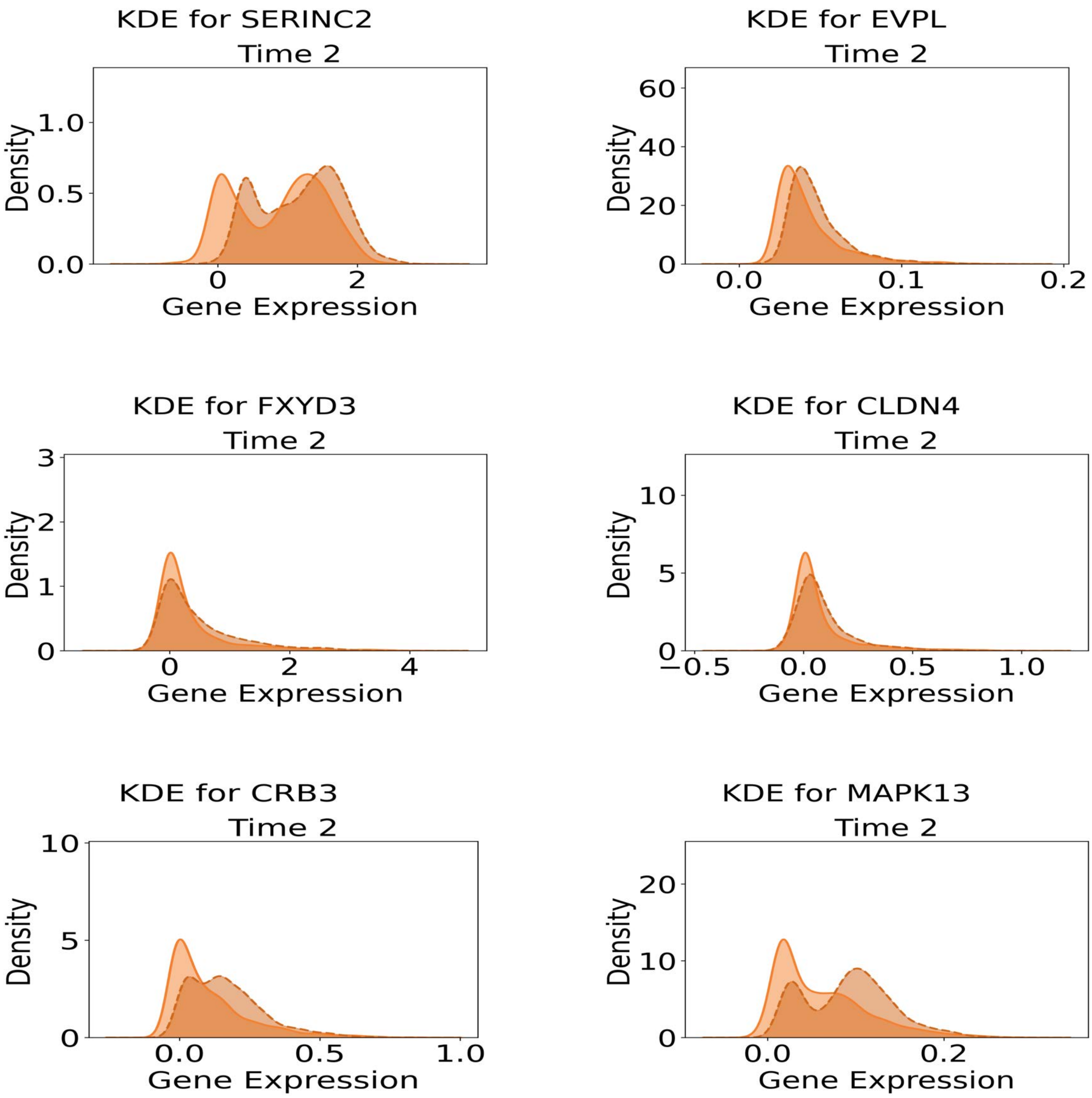

KDE for SERINC2
Time 2
Density
Gene Expression
0.0
0.5
1.0
0
2
KDE for EVPL
Time 2
Density
Gene Expression
0
20
40
60
0.0
0.1
0.2
KDE for FXYD3
Time 2
Density
Gene Expression
0
1
2
3
0
2
4
KDE for CLDN4
Time 2
Density
Gene Expression
0
5
10
−0.5
0.0
0.5
1.0
KDE for CRB3
Time 2
Density
Gene Expression
0
5
10
0.0
0.5
1.0
KDE for MAPK13
Time 2
Density
Gene Expression
0
10
20
0.0
0.2

Figure S14, Comparison of predicted and observed gene expression distributions at held-out time points in the EMT dataset, related to Figure 4B. Comparison of predicted and actual gene expression distributions at time 2. Predicted distributions are shown as dashed lines, while real data distributions are shown as solid lines.

AXL

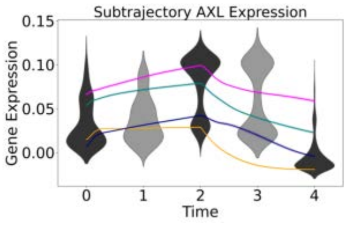

CXCR4

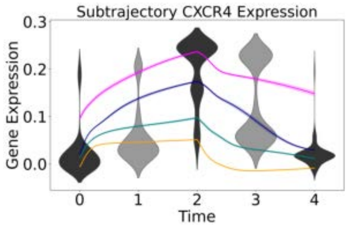

FN1

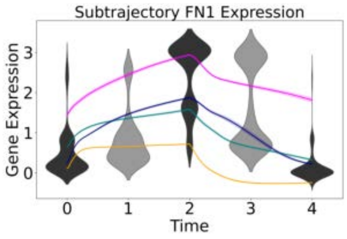

ITGA11

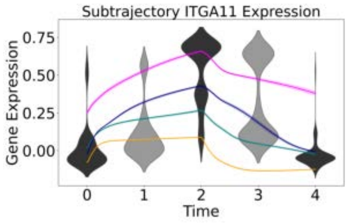

MMP2

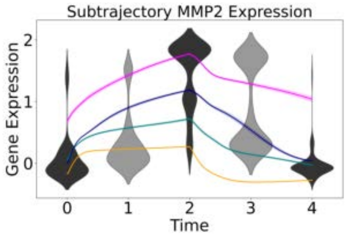

MMP9

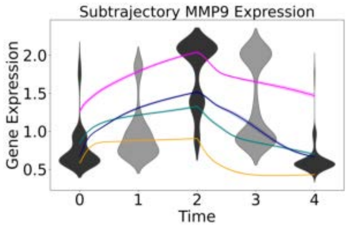

SNAI2

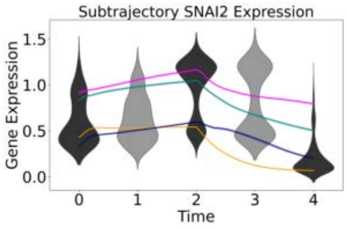

VIM

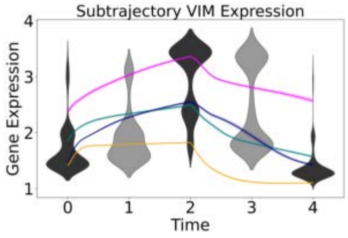

ZEB1

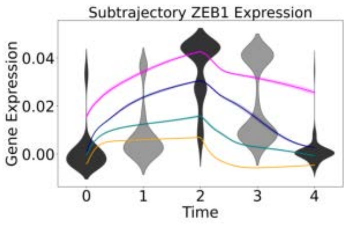

ZEB2

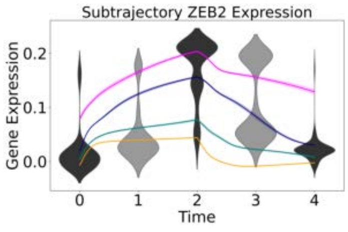

Figure S15, Average EMT relevant gene expression trajectories plotted separately for each subgroup, related to Figure 4G.

ABCG2

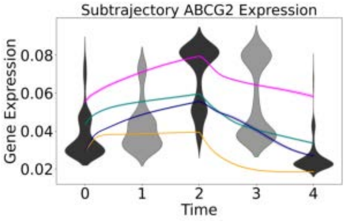

ALDH1A2

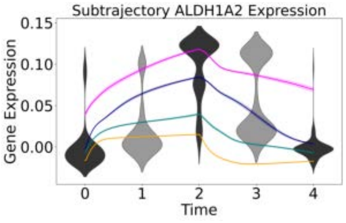

ALDH1A3

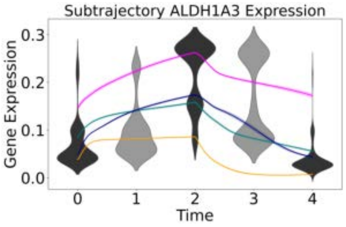

JAG1

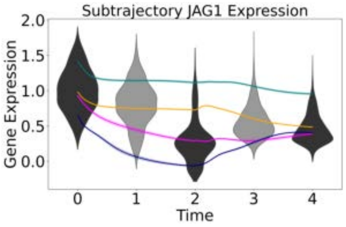

KLF5

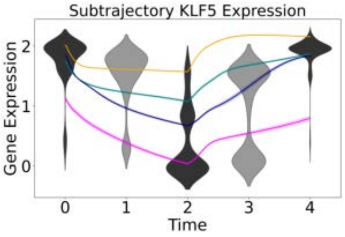

KLF7

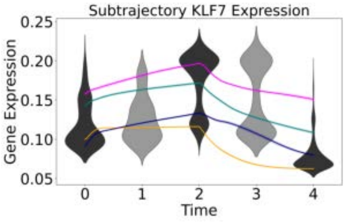

NOTCH1

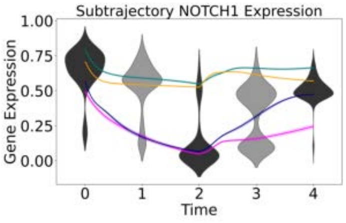

SOX2

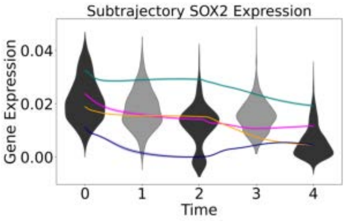

TCF7L2

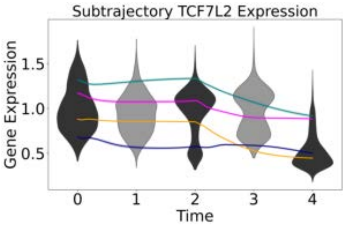

WNT5A

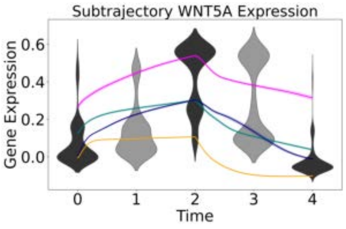

Figure S16, Average stemness relevant gene expression trajectories plotted separately for each subgroup, related to Figure 4G.

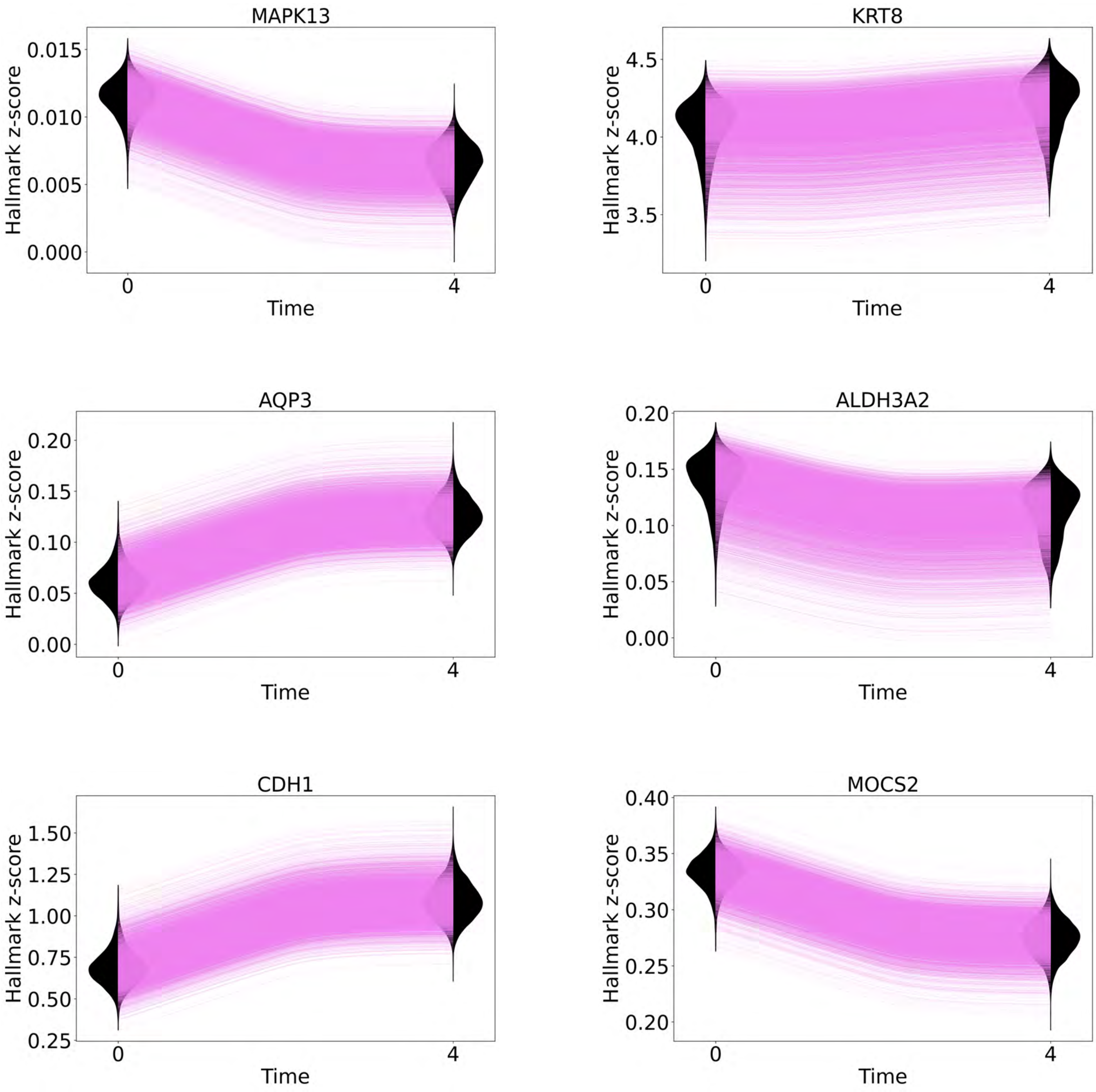

MAPK13
KRT8
AQP3
ALDH3A2
CDH1
MOCS2
Hallmark z-score
Time
0
4
0.015
0.010
0.005
0.000
4.5
4.0
3.5
0.20
0.15
0.10
0.05
0.00
1.50
1.25
1.00
0.75
0.50
0.25
0.40
0.35
0.30
0.25
0.20

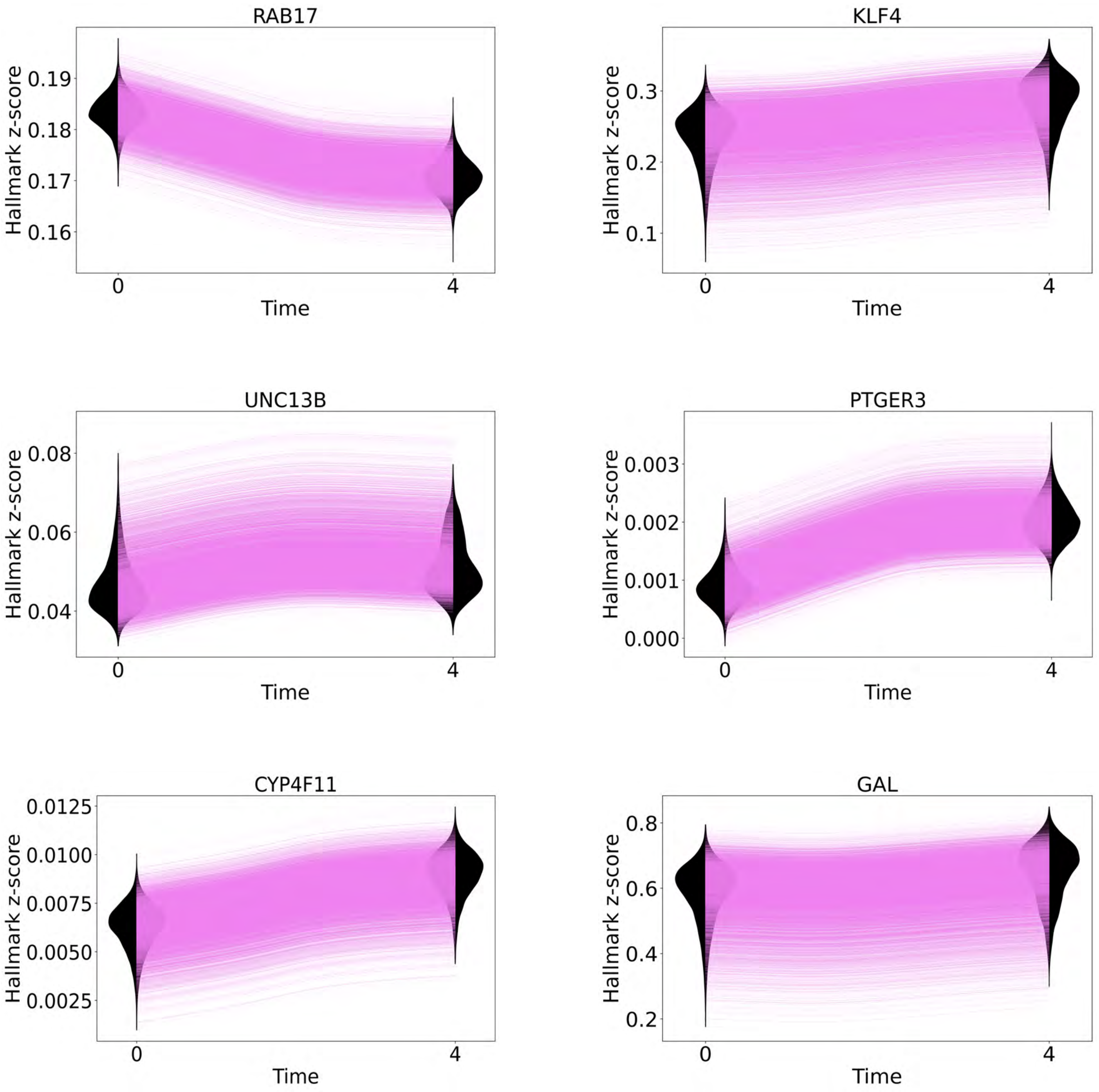

RAB17
KLF4
UNC13B
PTGER3
CYP4F11
GAL
Hallmark z-score
Time
0
4
0.19
0.18
0.17
0.16
0.3
0.2
0.1
0.08
0.06
0.04
0.003
0.002
0.001
0.000
0.0125
0.0100
0.0075
0.0050
0.0025
0.8
0.6
0.4
0.2

Figure S17, Single-cell, single-gene expression dynamics reconstructed by PROFET in the MCF7 cell line dataset, related to the section "Trajectory-Based Dissection of Treatment Response Heterogeneity and Early Biomarker Discovery." Single-cell, single-gene dynamics plotted as gene expression levels (y-axis) over time (x-axis), with each green curve representing an individual cell trajectory. The predicted trajectories are compared against real data distributions using violin plots (gray for test data, black for training data).

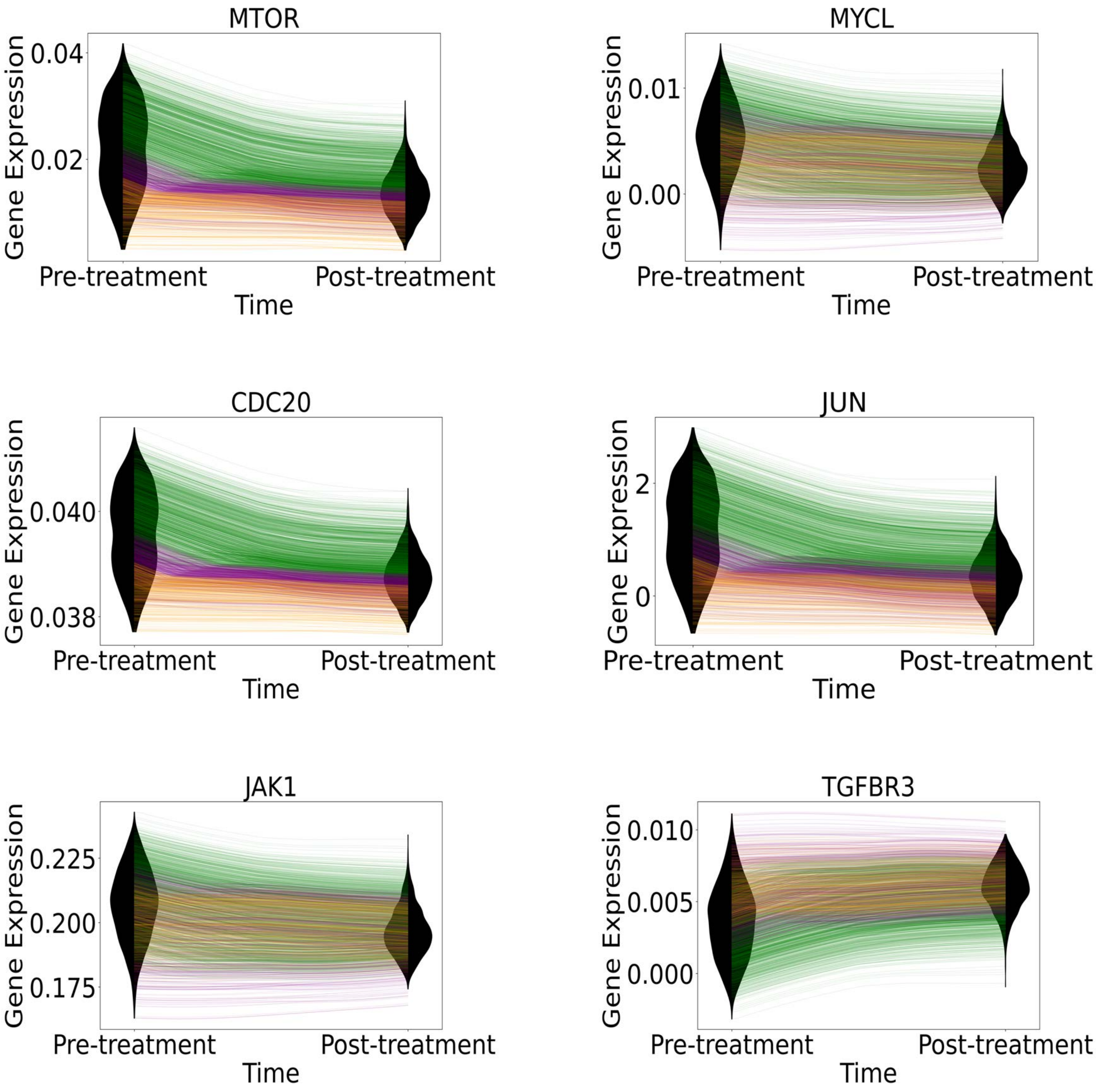
MTOR
Gene Expression
0.04
0.02
Pre-treatment
Post-treatment
Time
MYCL
Gene Expression
0.01
0.00
Pre-treatment
Post-treatment
Time
CDC20
Gene Expression
0.040
0.038
Pre-treatment
Post-treatment
Time
JUN
Gene Expression
2
0
Pre-treatment
Post-treatment
Time
JAK1
Gene Expression
0.225
0.200
0.175
Pre-treatment
Post-treatment
Time
TGFBR3
Gene Expression
0.010
0.005
0.000
Pre-treatment
Post-treatment
Time

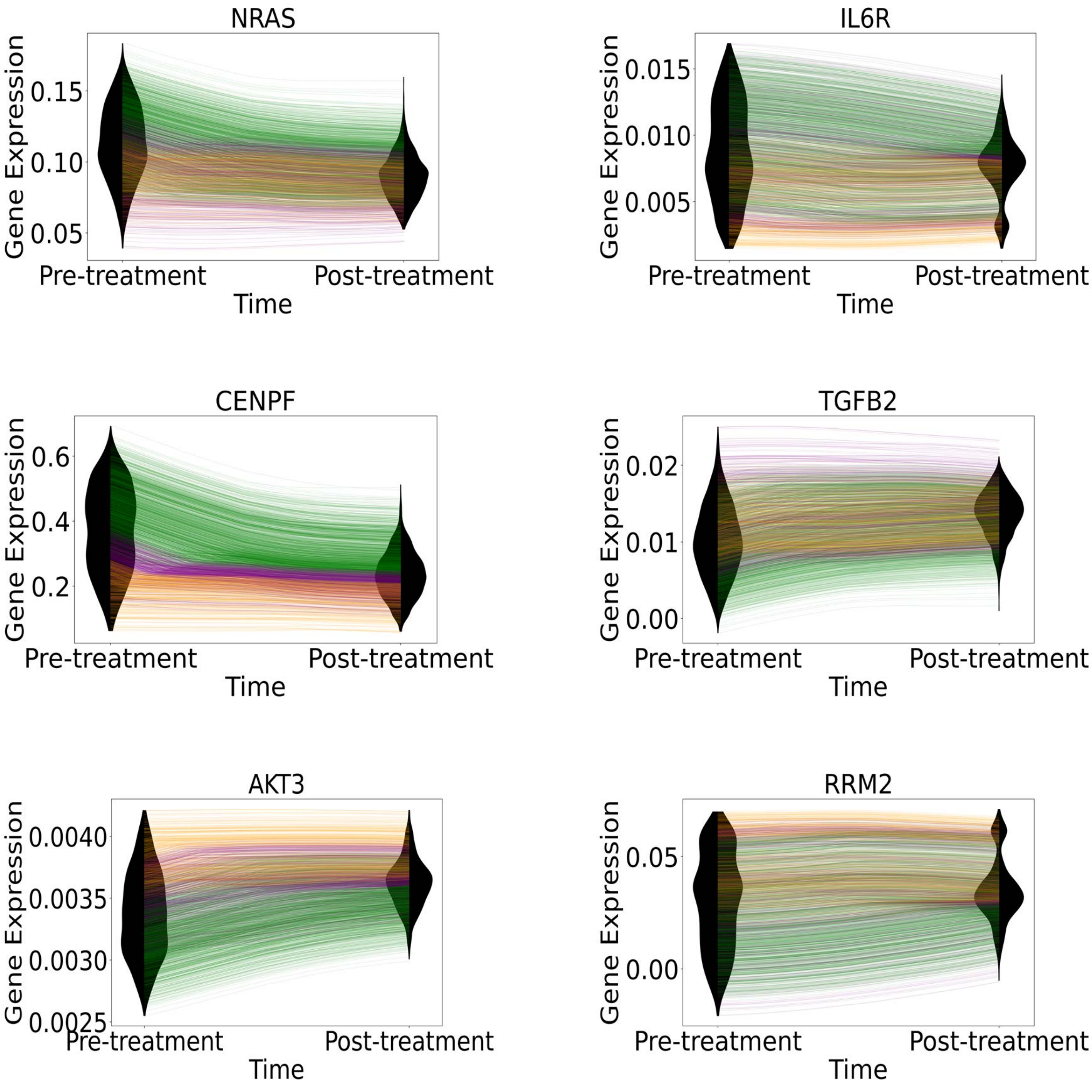

NRAS
Gene Expression
0.15
0.10
0.05
Pre-treatment
Post-treatment
Time
IL6R
Gene Expression
0.015
0.010
0.005
Pre-treatment
Post-treatment
Time
CENPF
Gene Expression
0.6
0.4
0.2
Pre-treatment
Post-treatment
Time
TGFB2
Gene Expression
0.02
0.01
0.00
Pre-treatment
Post-treatment
Time
AKT3
Gene Expression
0.0040
0.0035
0.0030
0.0025
Pre-treatment
Post-treatment
Time
RRM2
Gene Expression
0.05
0.00
Pre-treatment
Post-treatment
Time

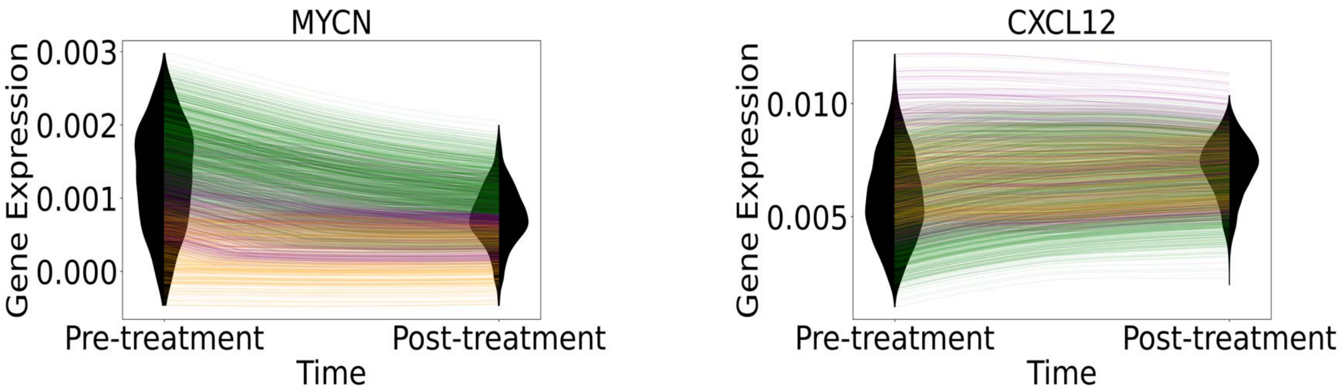

Figure S18, Single-cell, single-gene expression dynamics reconstructed by PROFET in the patient PA3 dataset, related to the section "Trajectory-Based Dissection of Treatment Response Heterogeneity and Early Biomarker Discovery." Violin plots show real expression data; colored lines represent predicted trajectories from three subgroups of cells defined by low, medium, and high phenotypic shift levels, as described in Figure 5.

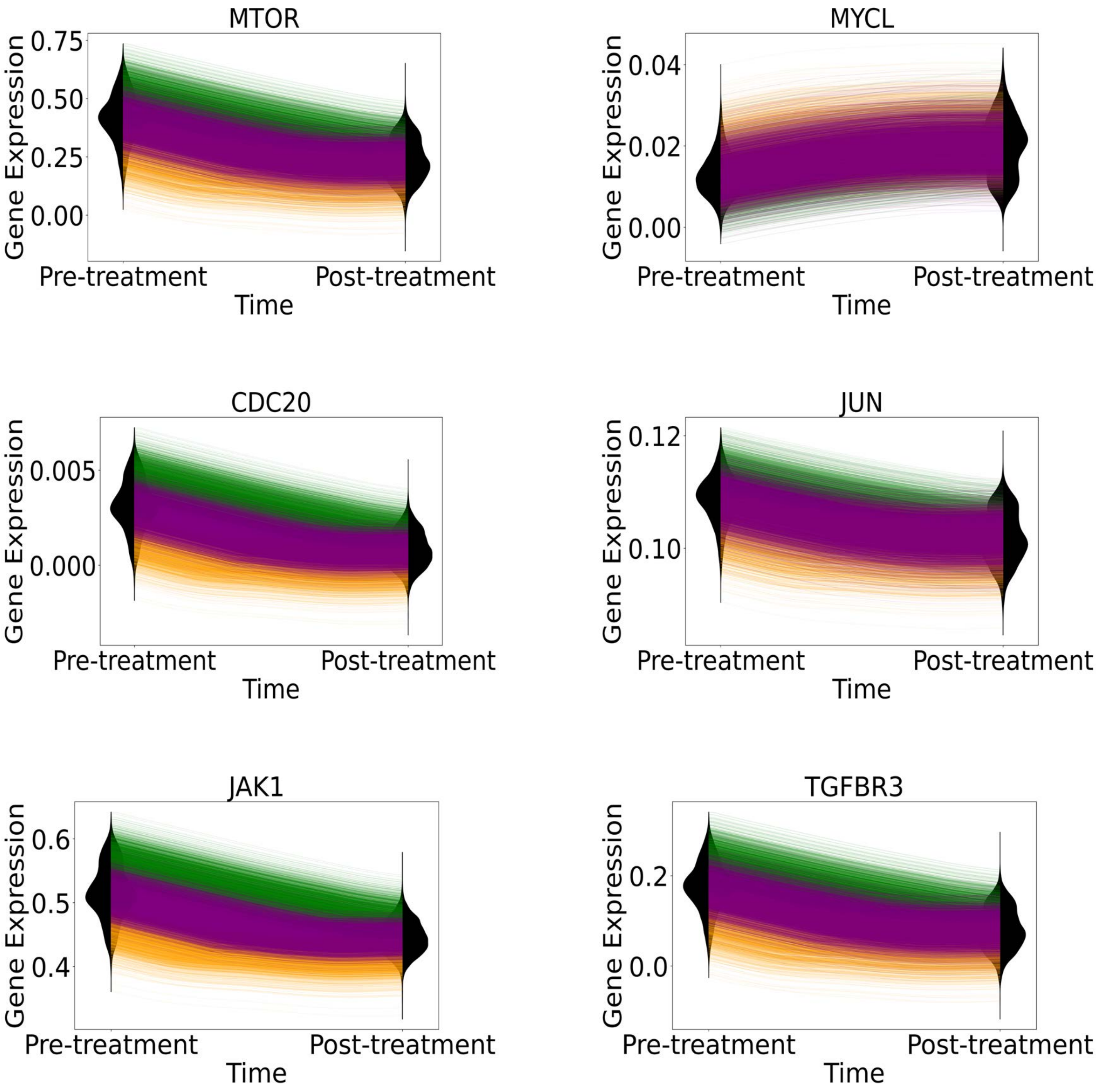

MTOR
Gene Expression
0.75
0.50
0.25
0.00
Pre-treatment
Post-treatment
Time
MYCL
Gene Expression
0.04
0.02
0.00
Pre-treatment
Post-treatment
Time
CDC20
Gene Expression
0.005
0.000
Pre-treatment
Post-treatment
Time
JUN
Gene Expression
0.12
0.10
Pre-treatment
Post-treatment
Time
JAK1
Gene Expression
0.6
0.5
0.4
Pre-treatment
Post-treatment
Time
TGFBR3
Gene Expression
0.2
0.0
Pre-treatment
Post-treatment
Time

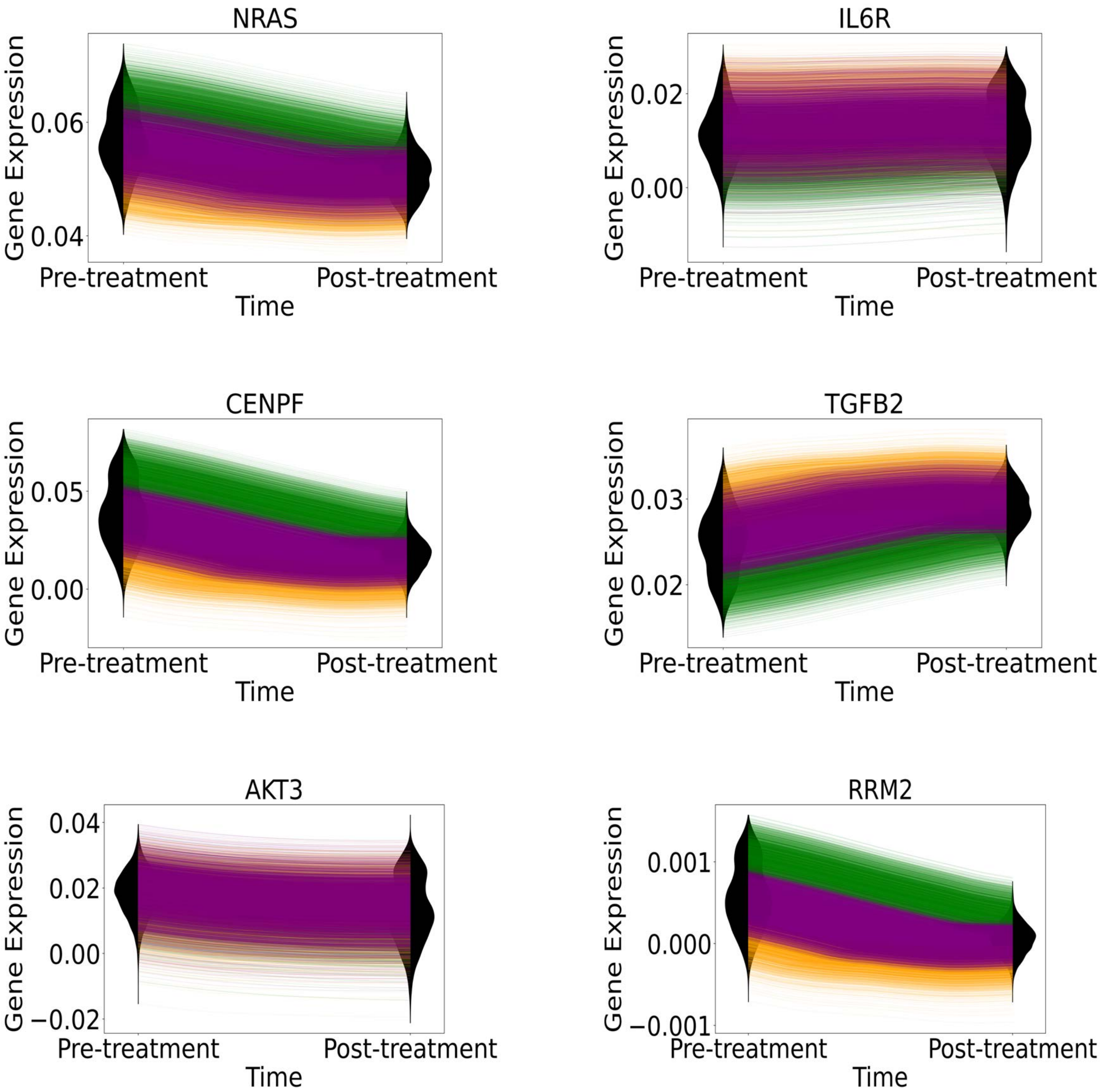

NRAS
IL6R
CENPF
TGFB2
AKT3
RRM2
Gene Expression
Time
Pre-treatment
Post-treatment
0.06
0.04
0.02
0.00
0.05
0.03
0.04
−0.02
0.001
0.000
−0.001

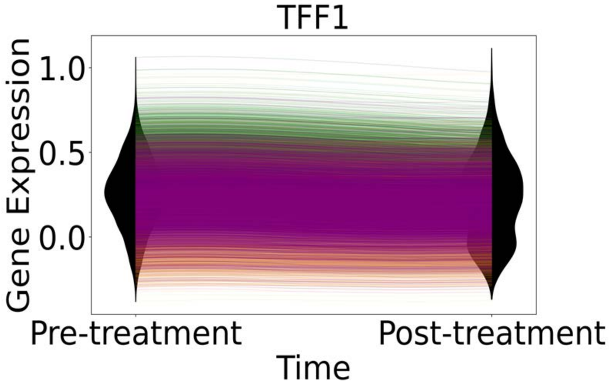

Figure S19, Single-cell, single-gene expression dynamics reconstructed by PROFET in the patient 862 dataset, related to the section “Trajectory-Based Dissection of Treatment Response Heterogeneity and Early Biomarker Discovery.” Violin plots show real expression data; colored lines represent predicted trajectories from three subgroups of cells defined by low, medium, and high phenotypic shift levels, as described in Figure 5.

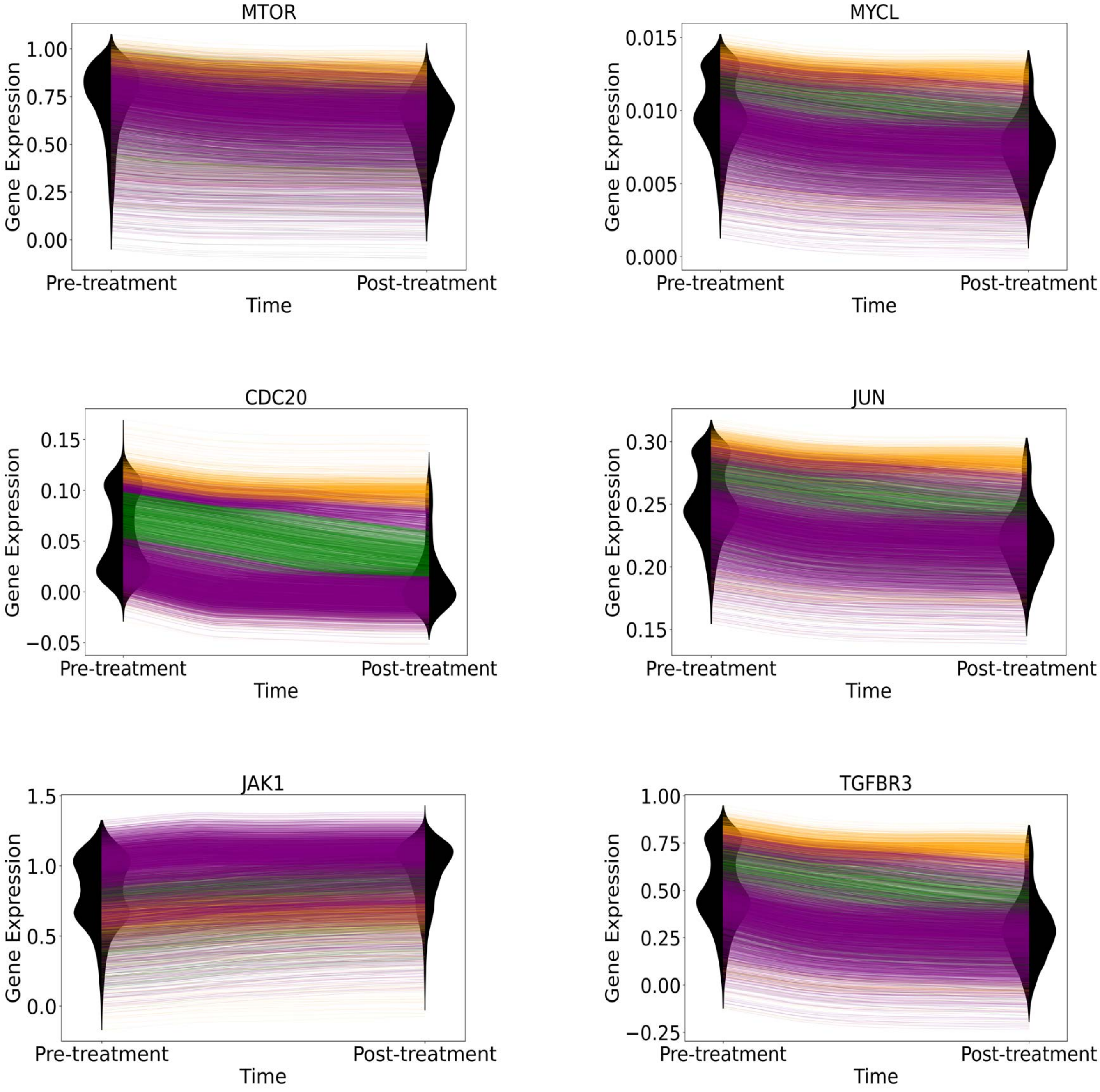
MTOR
MYCL
CDC20
JUN
JAK1
TGFBR3
Gene Expression
Time
Pre-treatment
Post-treatment
1.00
0.75
0.50
0.25
0.00
0.015
0.010
0.005
0.000
0.15
0.10
0.05
0.00
−0.05
0.30
0.25
0.20
0.15
1.5
1.0
0.5
0.0
1.00
0.75
0.50
0.25
0.00
−0.25

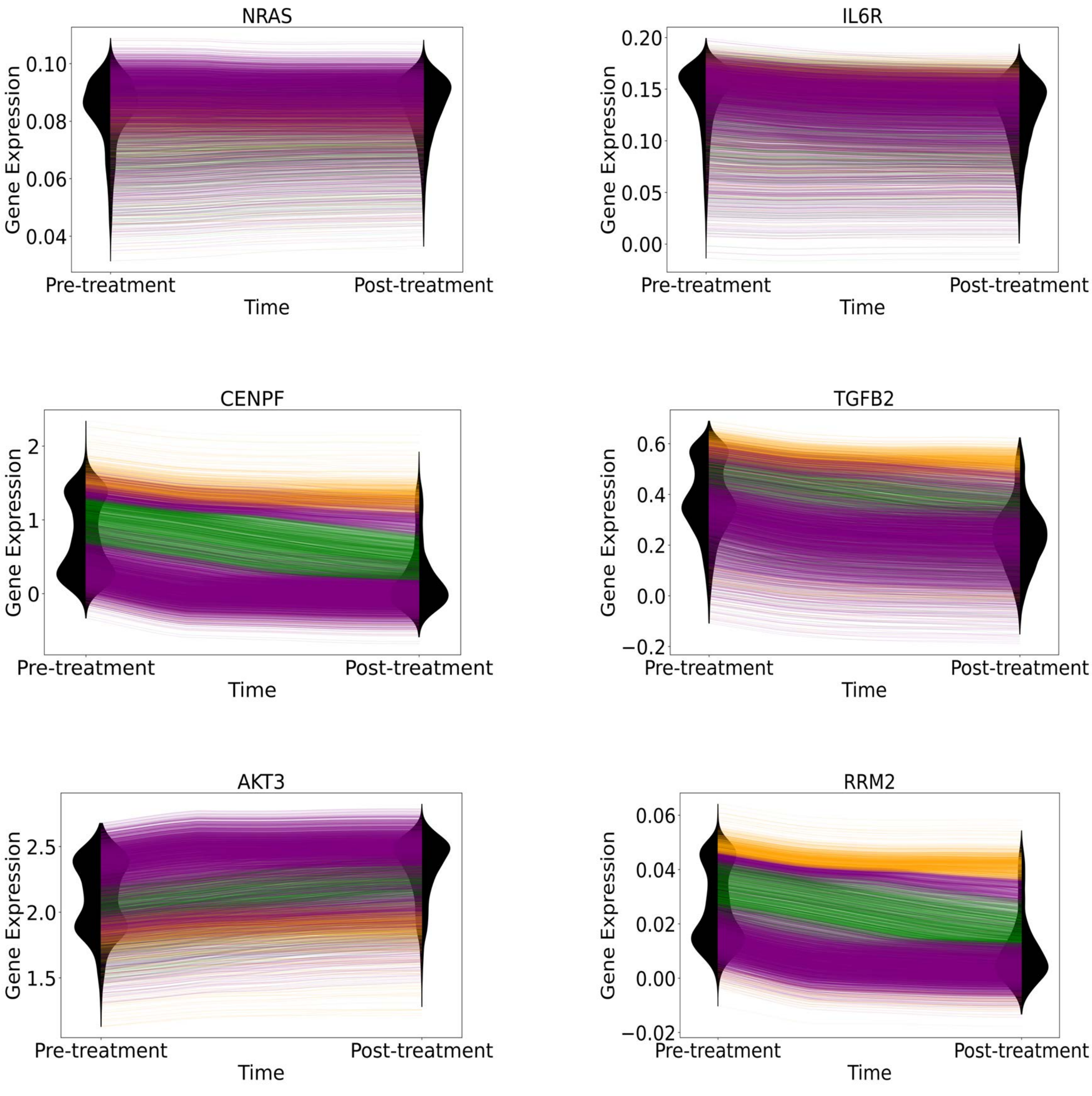
NRAS
Gene Expression
0.10
0.08
0.06
0.04
Pre-treatment
Post-treatment
Time
IL6R
Gene Expression
0.20
0.15
0.10
0.05
0.00
Pre-treatment
Post-treatment
Time
CENPF
Gene Expression
2
1
0
Pre-treatment
Post-treatment
Time
TGFB2
Gene Expression
0.6
0.4
0.2
0.0
−0.2
Pre-treatment
Post-treatment
Time
AKT3
Gene Expression
2.5
2.0
1.5
Pre-treatment
Post-treatment
Time
RRM2
Gene Expression
0.06
0.04
0.02
0.00
−0.02
Pre-treatment
Post-treatment
Time

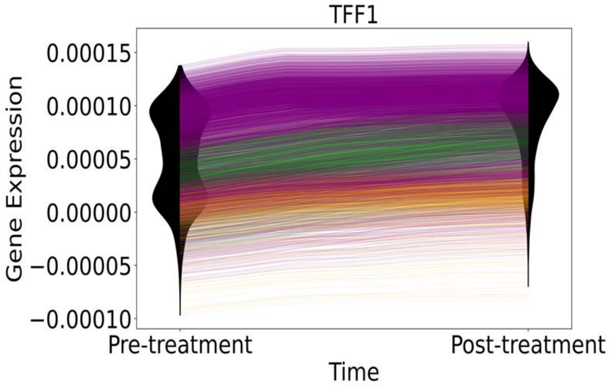

Figure S20, Single-cell, single-gene expression dynamics reconstructed by PROFET in the patient 887 dataset, related to the section "Trajectory-Based Dissection of Treatment Response Heterogeneity and Early Biomarker Discovery." Individual cells across three patient datasets. Violin plots show real expression data; colored lines represent predicted trajectories from three subgroups of cells defined by low, medium, and high phenotypic shift levels, as described in Figure 5.

A

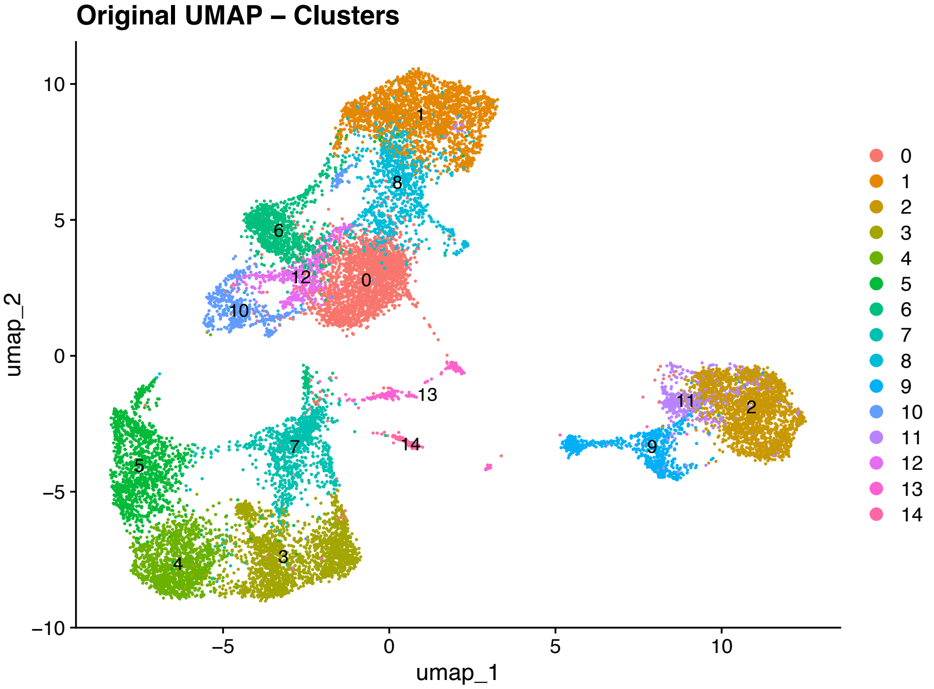
Original UMAP – Clusters
umap_2
umap_1
10
5
0
−5
−10
−5
0
5
10

B

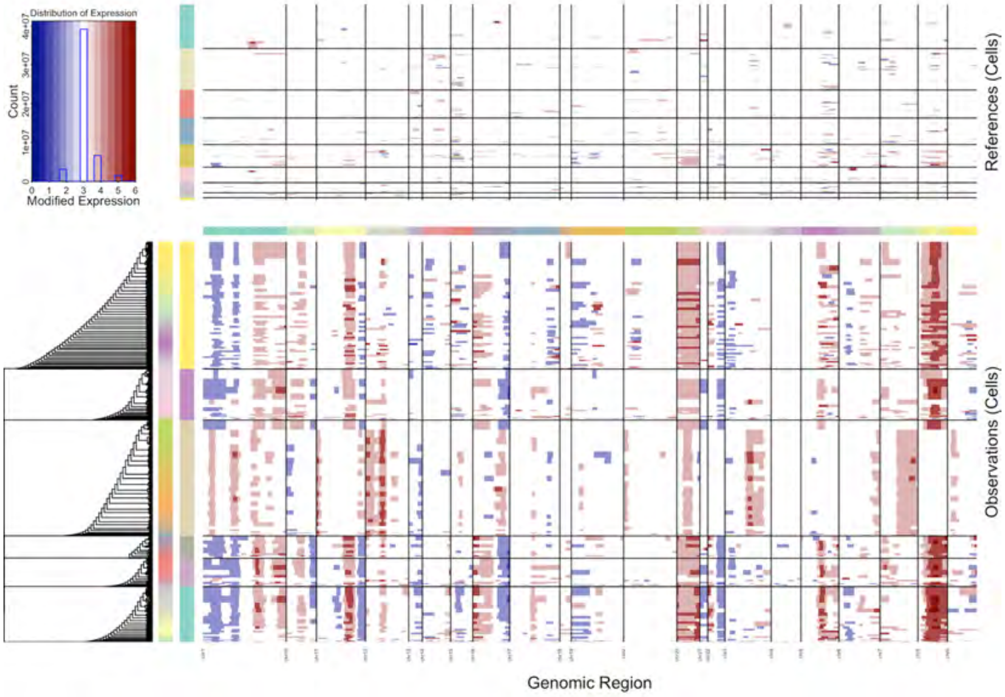
Distribution of Expression
Count
Modified Expression
References (Cells)
Observations (Cells)
Genomic Region

C

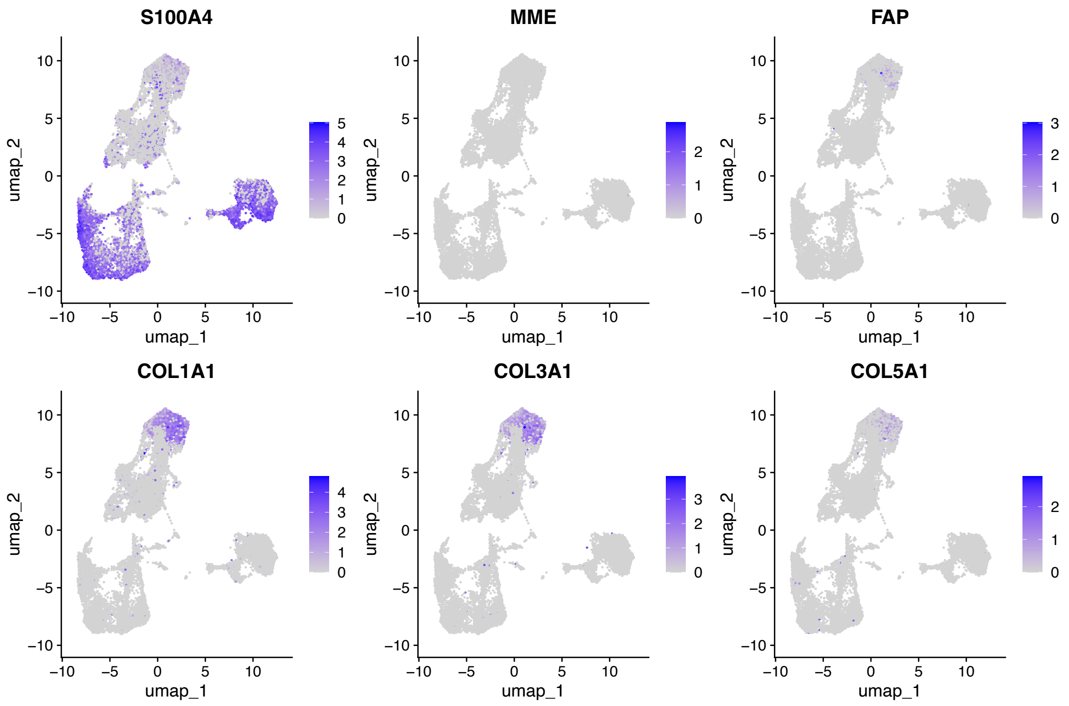
S100A4
MME
FAP
COL1A1
COL3A1
COL5A1
umap_1
umap_2

D

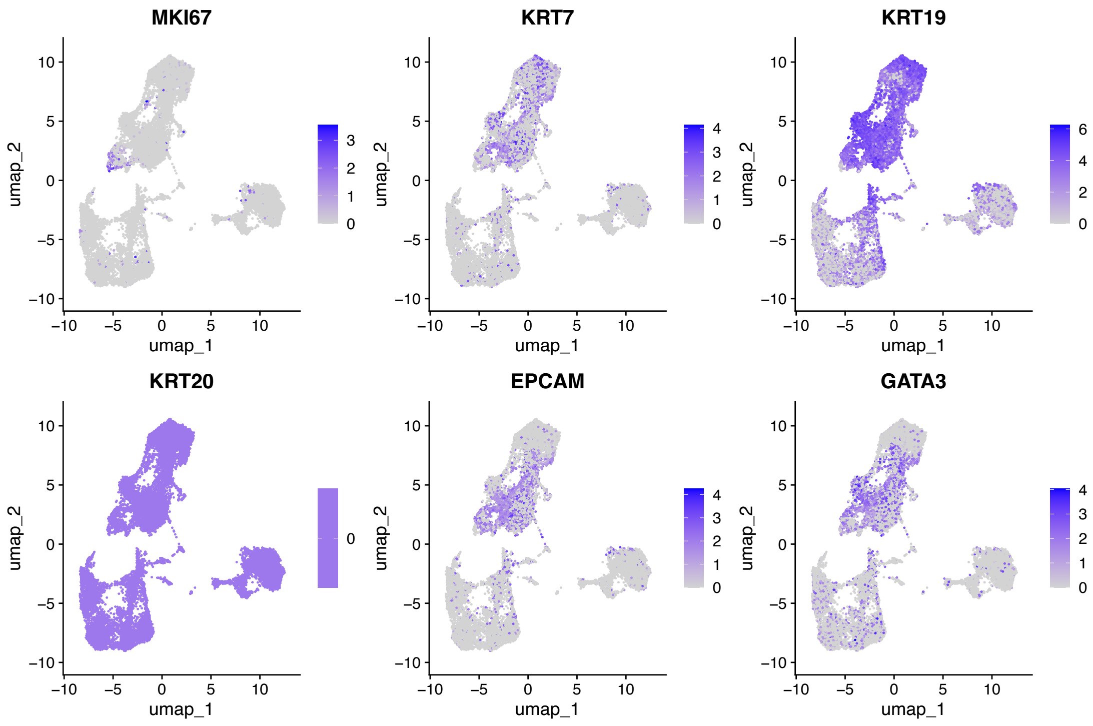
MKI67
KRT7
KRT19
KRT20
EPCAM
GATA3
umap_1
umap_2

Figure S21, inferCNV analysis of patient PA3 from Luo et al, related to the section "Methods: Preprocessing of breast cancer patient data."

(A) Clustering results of single-cell data.
(B) inferCNV analysis for identifying large-scale chromosomal copy number variations.
(C) Gene expression heatmap for fibroblast marker genes.
(D) Gene expression heatmap for epithelial marker genes.

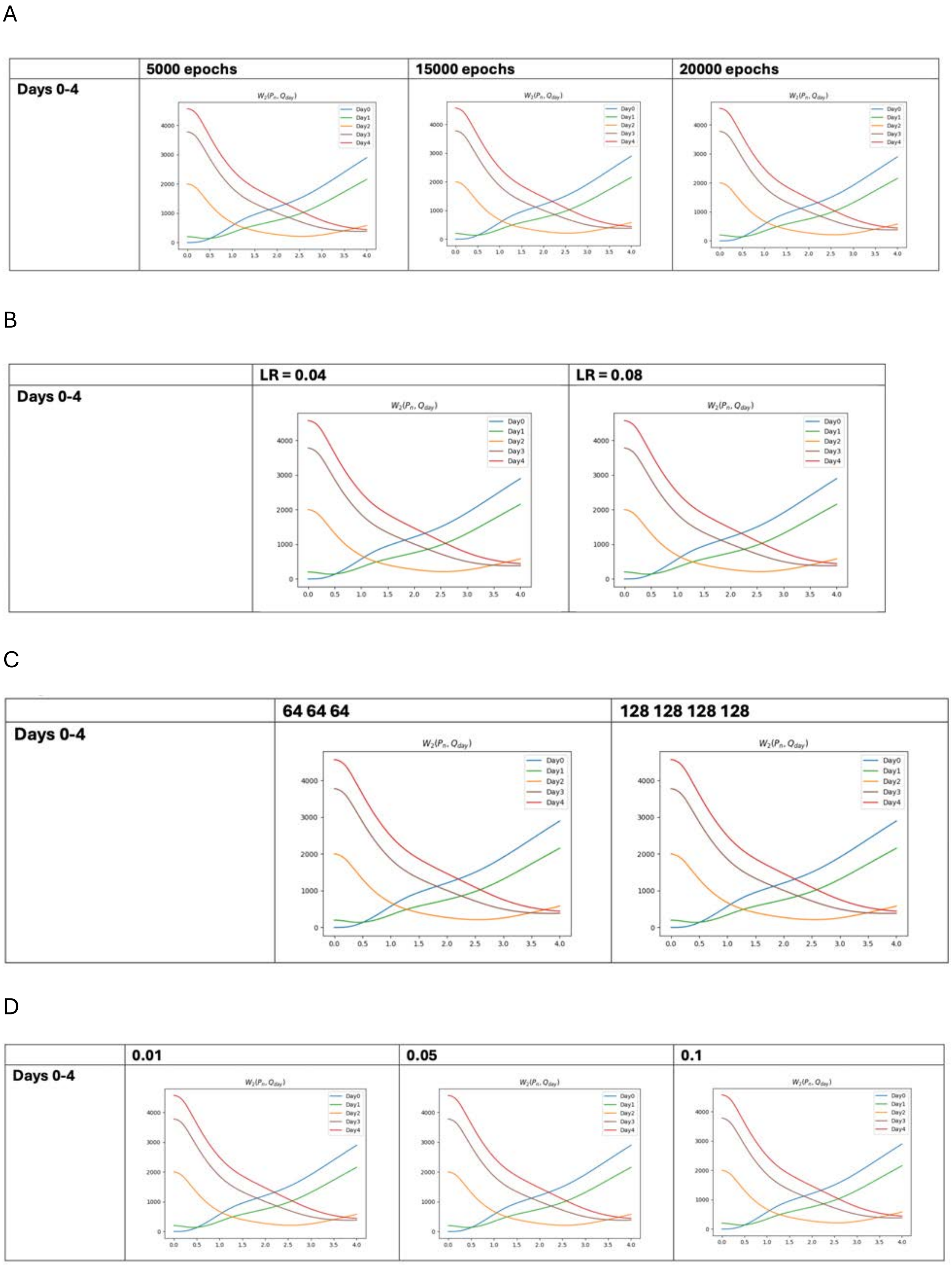

Figure S22, Sensitivity of trajectory prediction performance to hyperparameter selection, related to the section “Methods: Model architecture and training algorithm.” Effect of tuning hyperparameters on the Wasserstein distance between predicted trajectories and real data over time. Lower distances indicate closer alignment between predicted and observed trajectories.

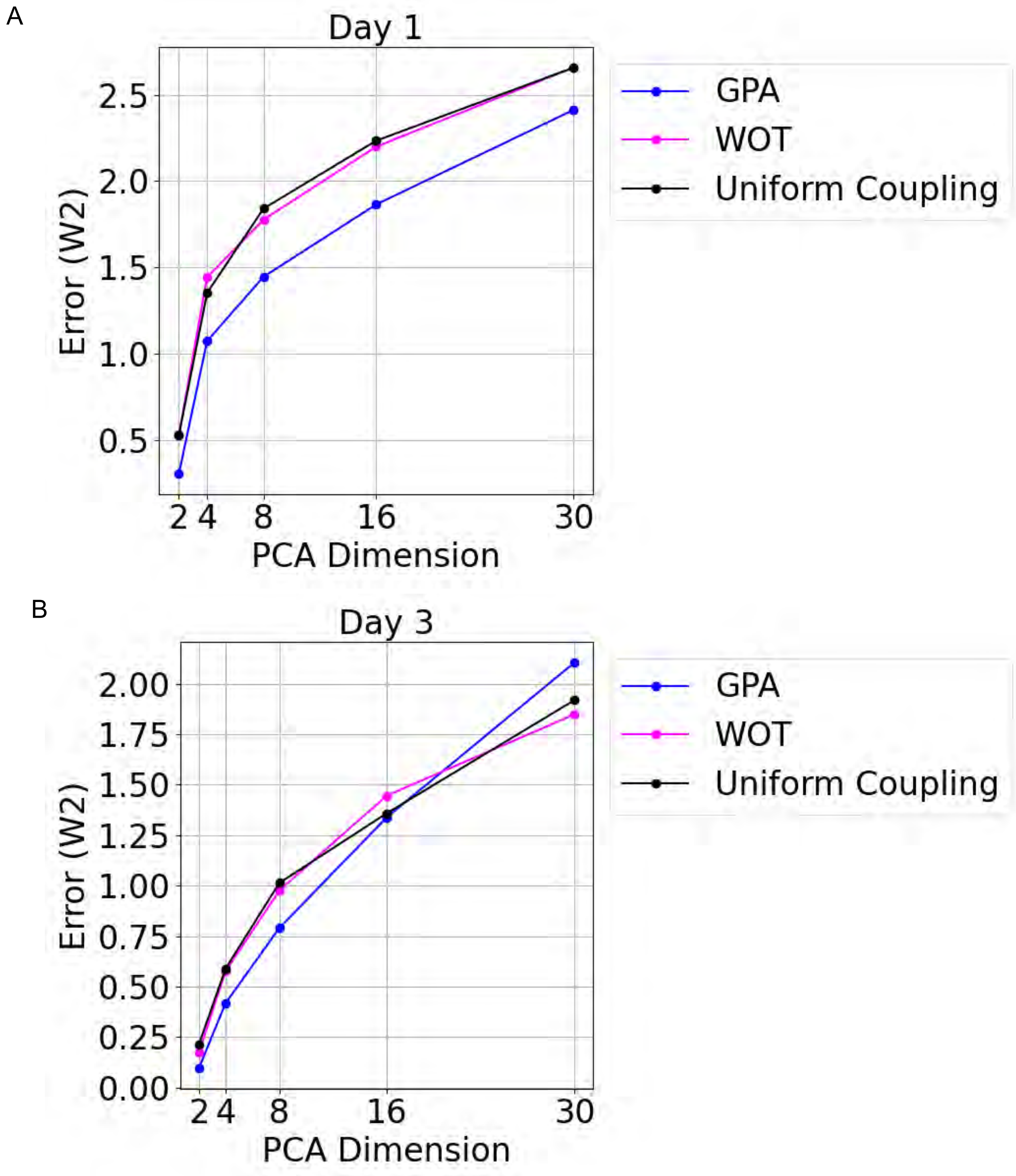
A
Day 1
Error (W2)
2.5
2.0
1.5
1.0
0.5
2 4 8 16 30
PCA Dimension
GPA
WOT
Uniform Coupling
B
Day 3
Error (W2)
2.00
1.75
1.50
1.25
1.00
0.75
0.50
0.25
0.00
2 4 8 16 30
PCA Dimension
GPA
WOT
Uniform Coupling

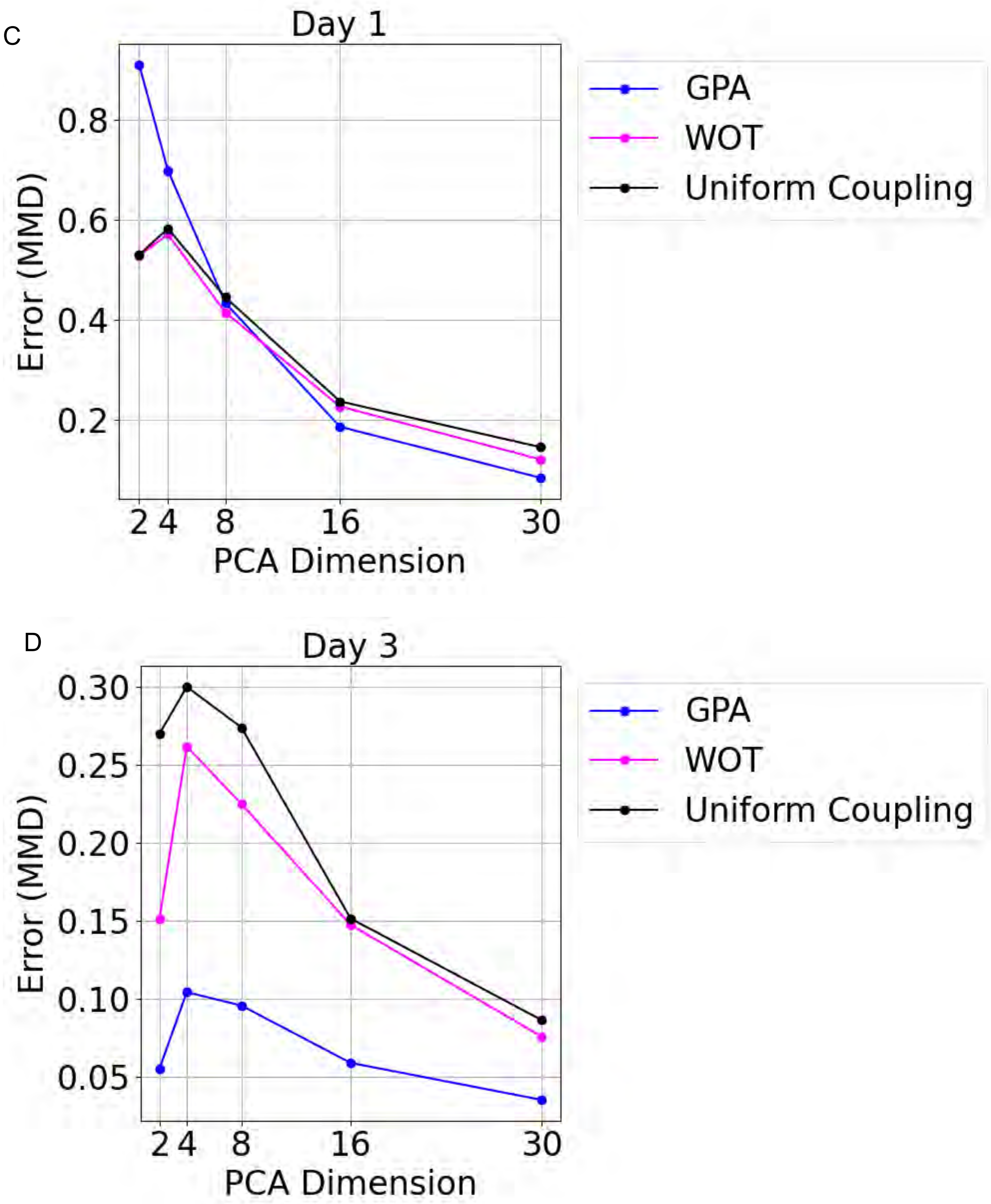
C
Day 1
Error (MMD)
0.8
0.6
0.4
0.2
2 4 8 16 30
PCA Dimension
GPA
WOT
Uniform Coupling
D
Day 3
Error (MMD)
0.30
0.25
0.20
0.15
0.10
0.05
2 4 8 16 30
PCA Dimension
GPA
WOT
Uniform Coupling

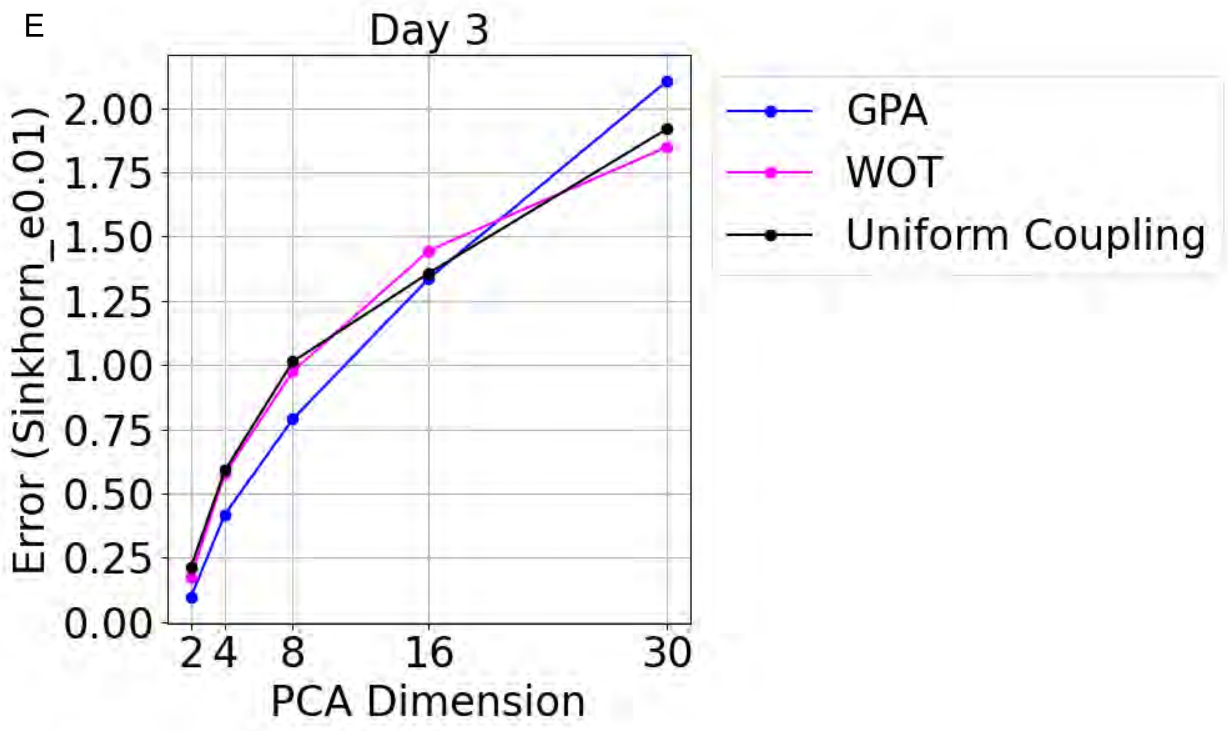
E
Day 3
Error (Sinkhorn_e0.01)
2.00
1.75
1.50
1.25
1.00
0.75
0.50
0.25
0.00
2
4
8
16
30
PCA Dimension
GPA
WOT
Uniform Coupling

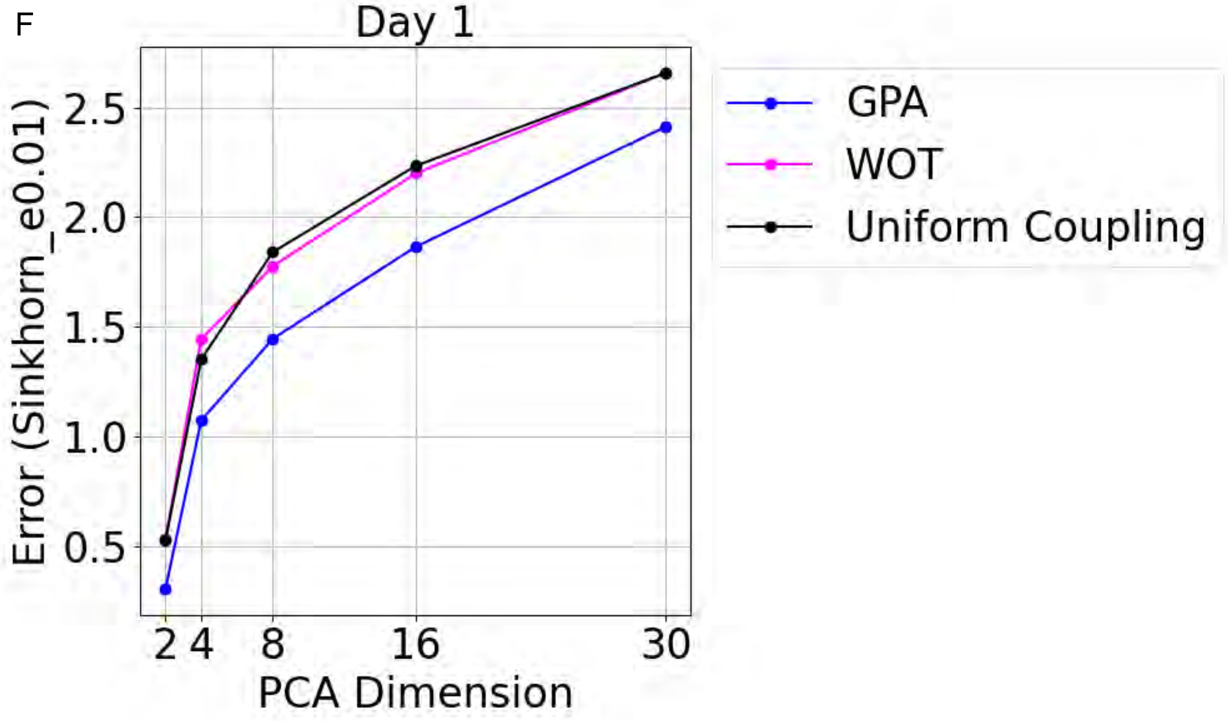
F
Day 1
Error (Sinkhorn_e0.01)
2.5
2.0
1.5
1.0
0.5
2
4
8
16
30
PCA Dimension
GPA
WOT
Uniform Coupling

G

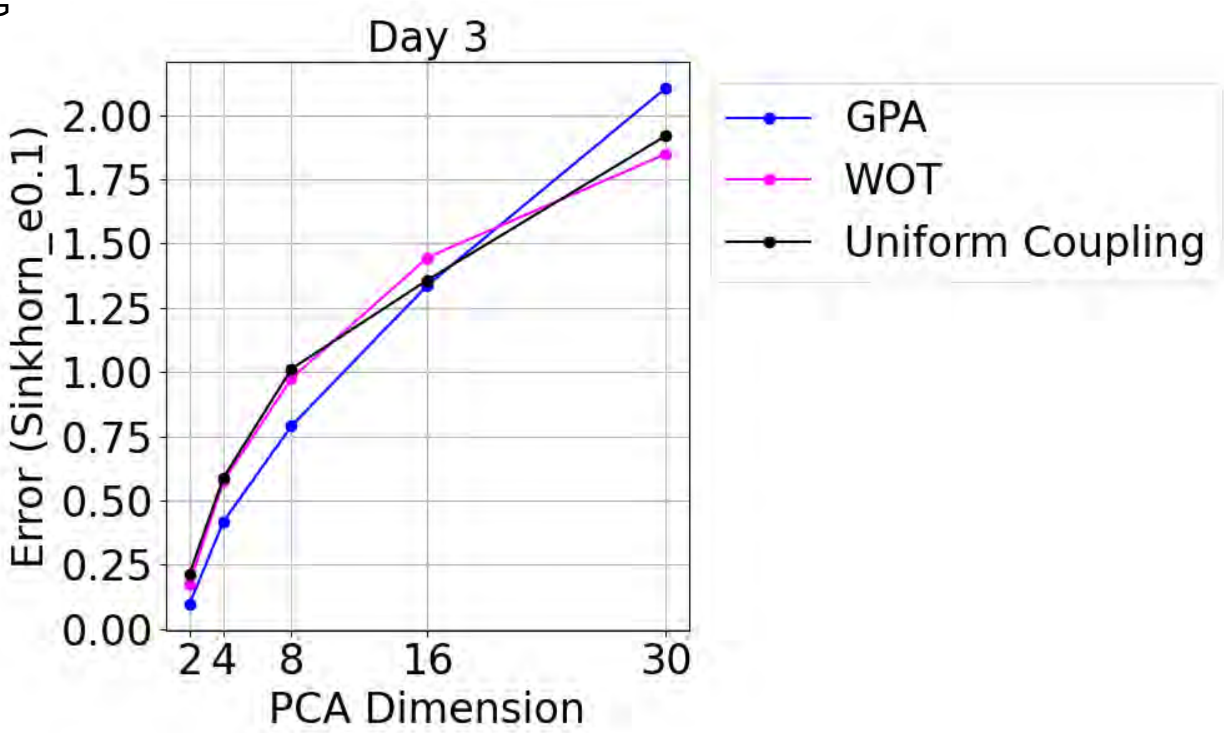

Day 3
Error (Sinkhorn_e0.1)
0.00
0.25
0.50
0.75
1.00
1.25
1.50
1.75
2.00
2 4 8 16 30
PCA Dimension
GPA
WOT
Uniform Coupling

H

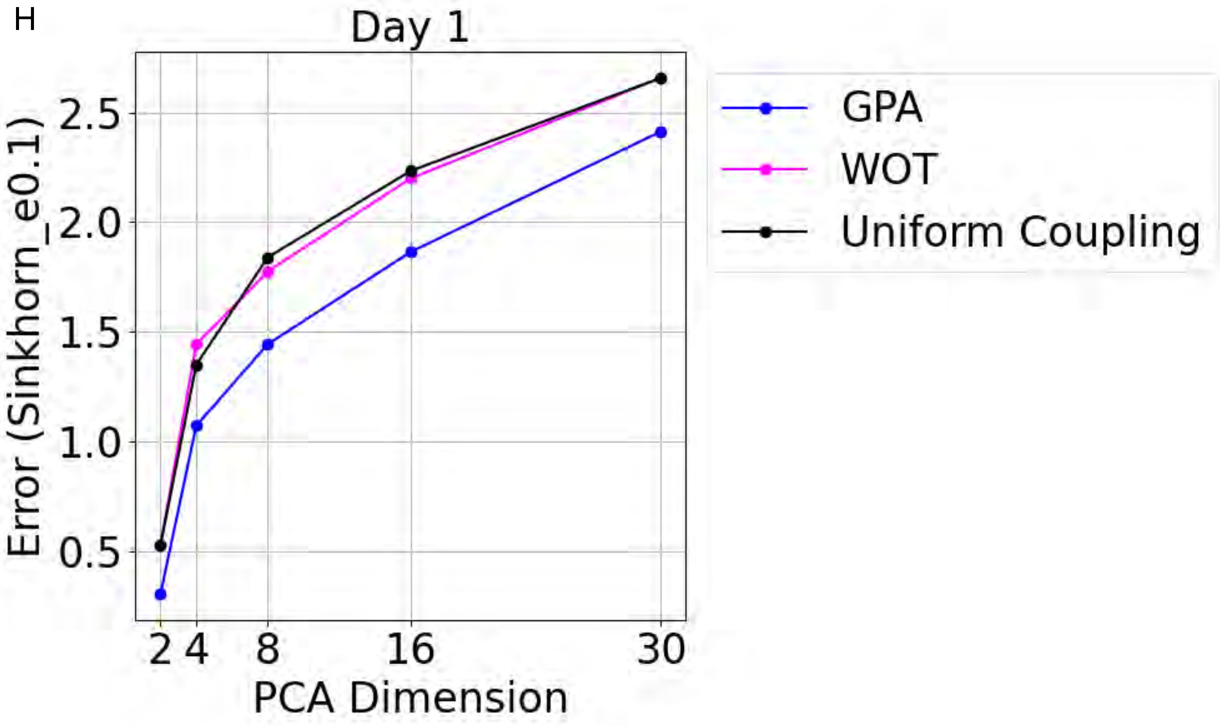

Day 1
Error (Sinkhorn_e0.1)
0.5
1.0
1.5
2.0
2.5
2 4 8 16 30
PCA Dimension
GPA
WOT
Uniform Coupling

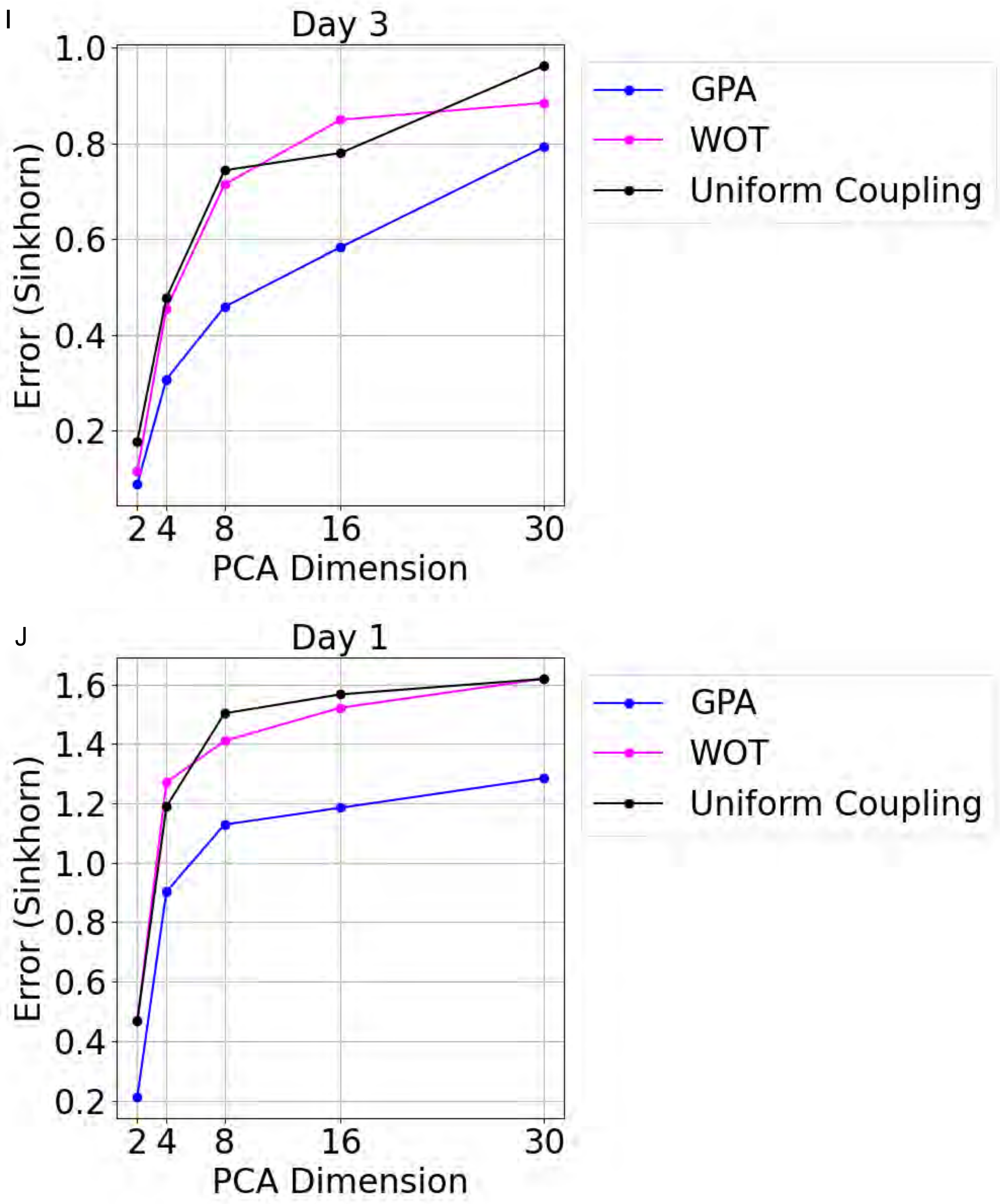
I
Day 3
1.0
0.8
0.6
0.4
0.2
Error (Sinkhorn)
2 4 8 16 30
PCA Dimension
GPA
WOT
Uniform Coupling
J
Day 1
1.6
1.4
1.2
1.0
0.8
0.6
0.4
0.2
Error (Sinkhorn)
2 4 8 16 30
PCA Dimension
GPA
WOT
Uniform Coupling

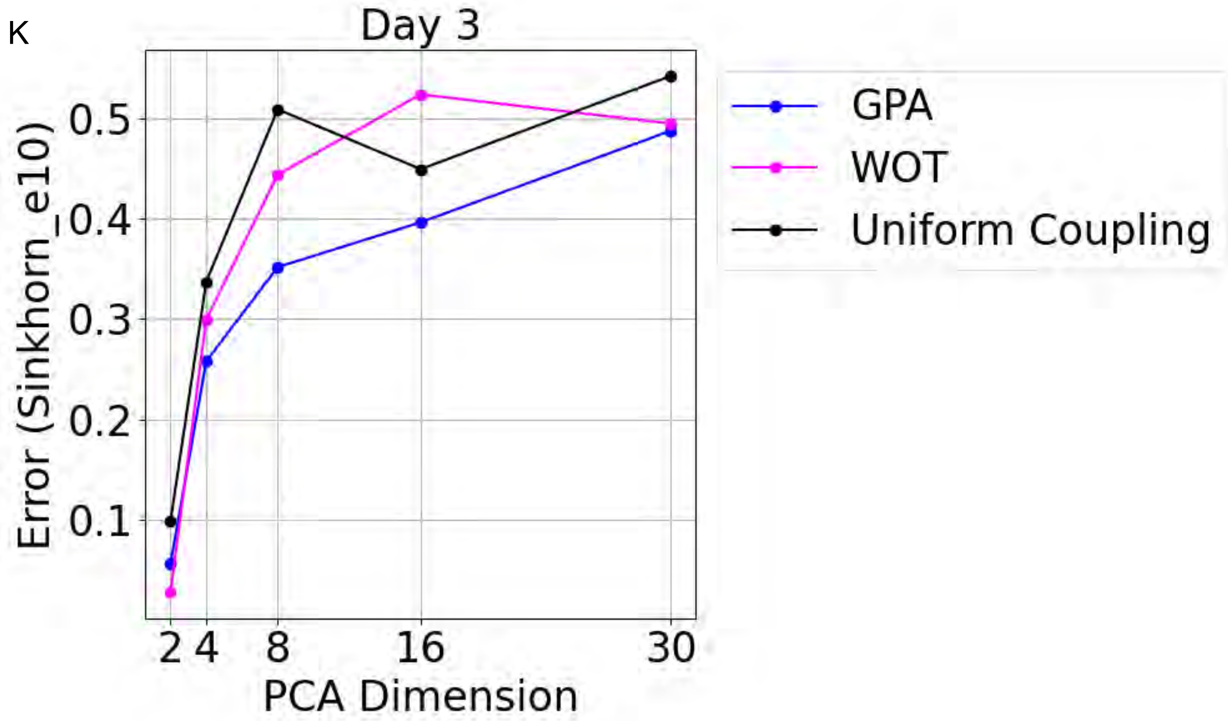
K
Day 3
Error (Sinkhorn_e10)
0.1
0.2
0.3
0.4
0.5
2 4 8 16 30
PCA Dimension
GPA
WOT
Uniform Coupling

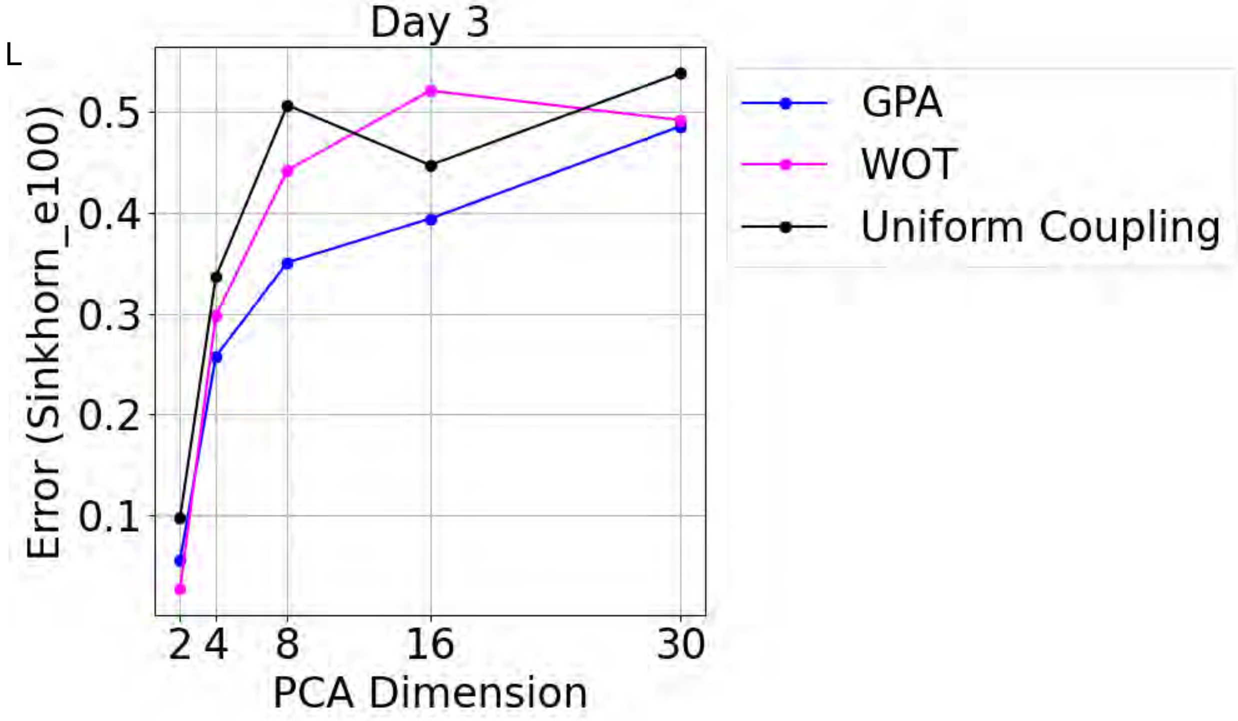
L
Day 3
Error (Sinkhorn_e100)
0.1
0.2
0.3
0.4
0.5
2 4 8 16 30
PCA Dimension
GPA
WOT
Uniform Coupling

M

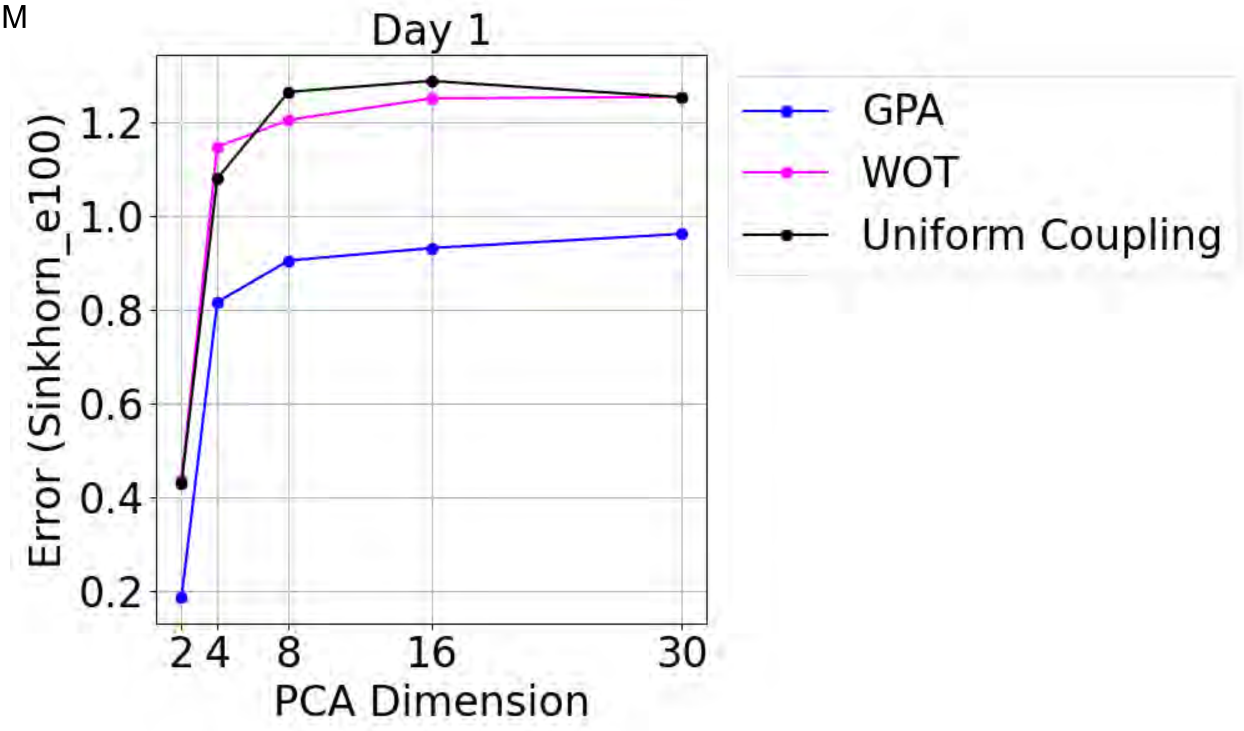

N

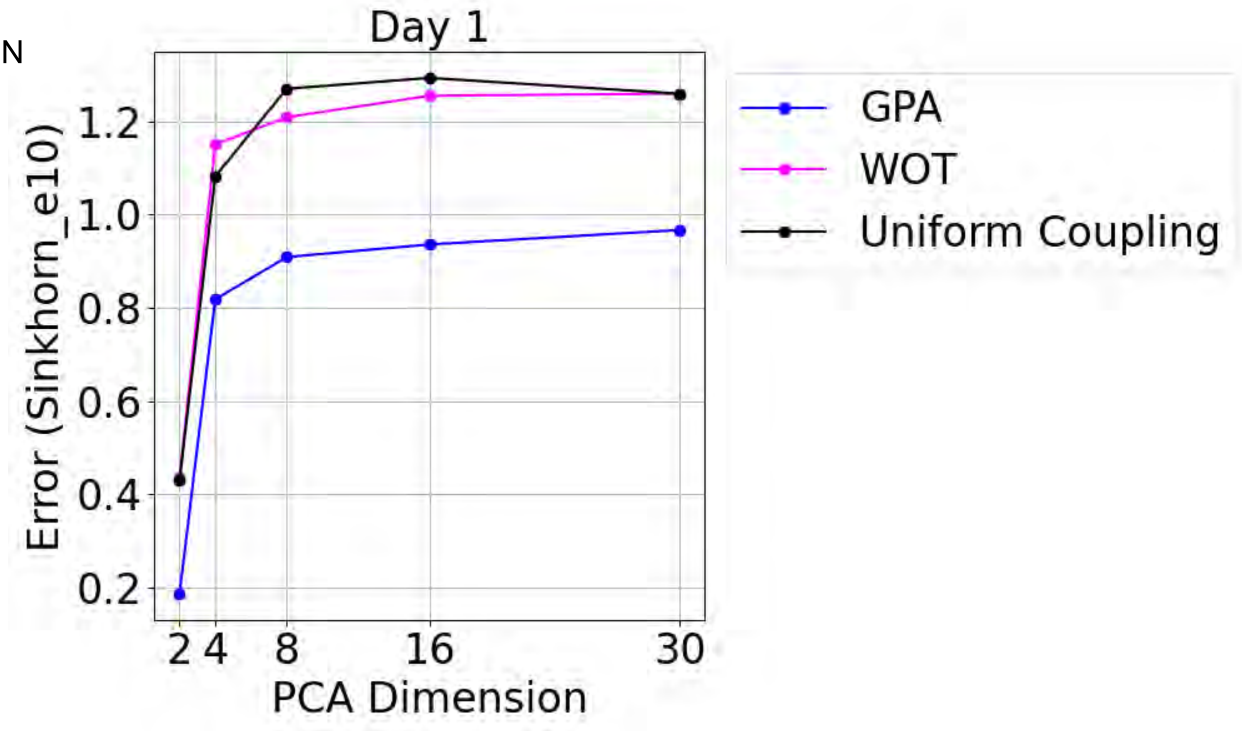

Figure S23, Effect of latent dimensionality on trajectory prediction performance in the mESC dataset, related to the section “Methods: Downstream Analysis Using PROFET.”

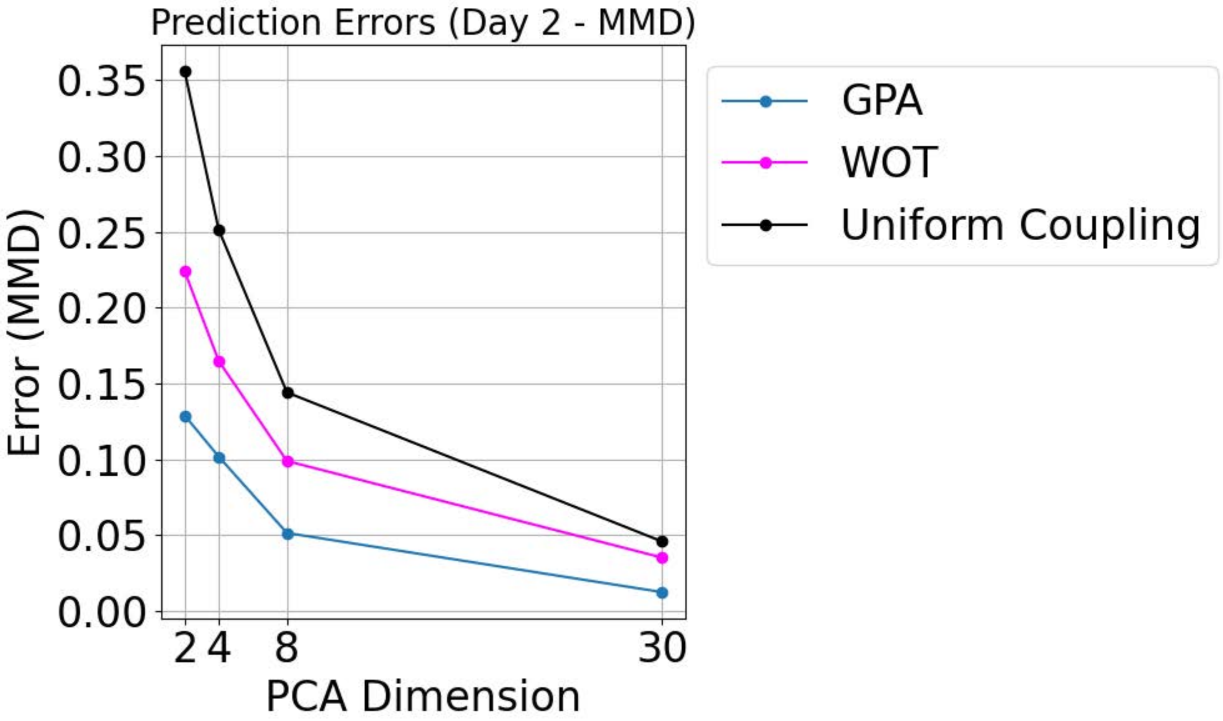
Prediction Errors (Day 2 - MMD)
Error (MMD)
0.00
0.05
0.10
0.15
0.20
0.25
0.30
0.35
PCA Dimension
2
4
8
30
GPA
WOT
Uniform Coupling

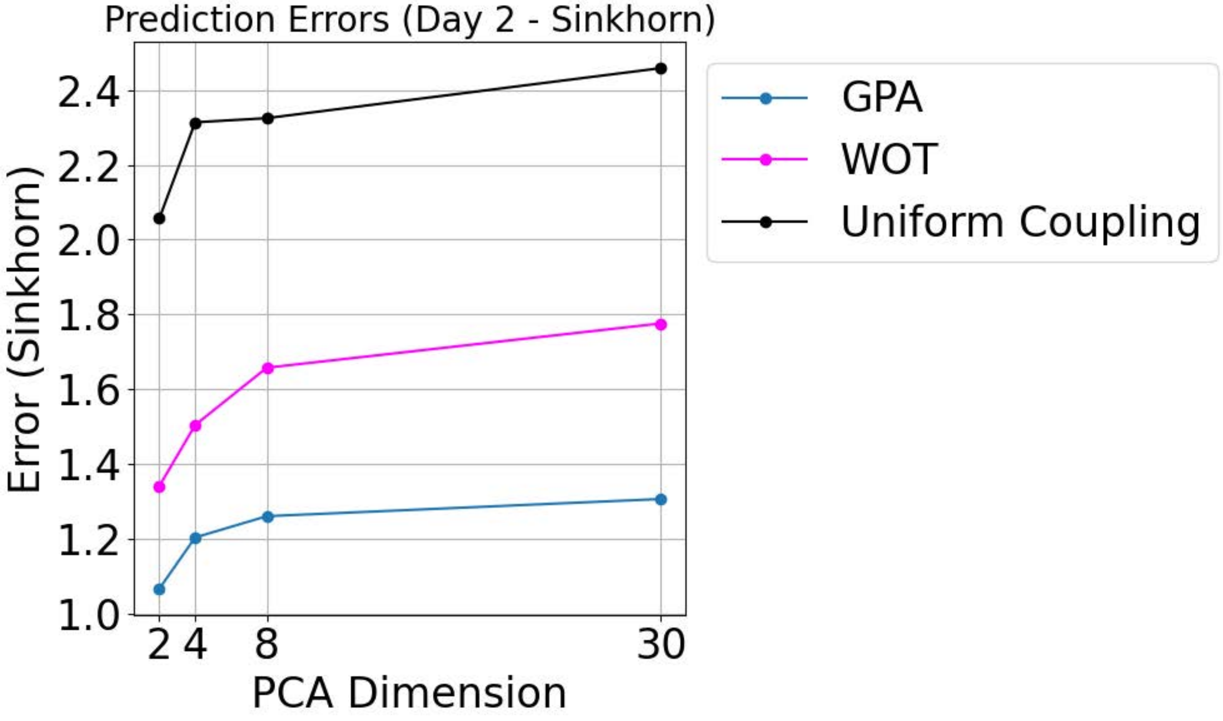
Prediction Errors (Day 2 - Sinkhorn)
Error (Sinkhorn)
1.0
1.2
1.4
1.6
1.8
2.0
2.2
2.4
PCA Dimension
2
4
8
30
GPA
WOT
Uniform Coupling

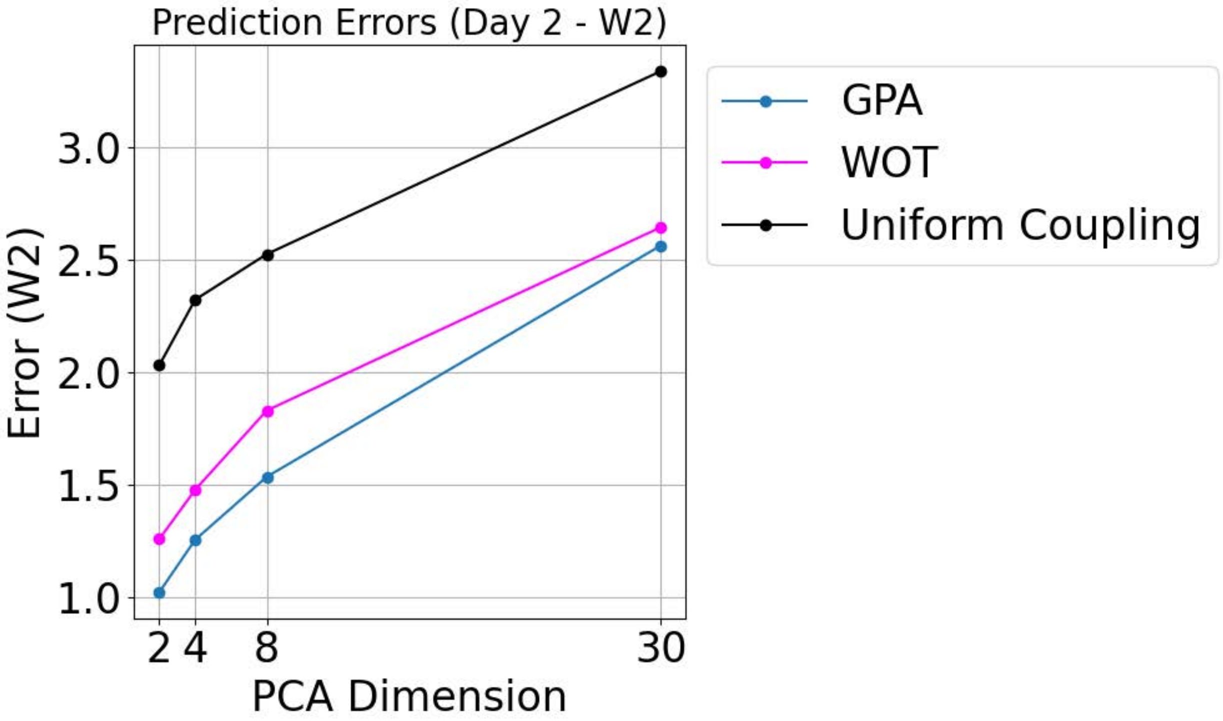

Figure S24, Effect of latent dimensionality on trajectory prediction performance in the EMT dataset, related to the section “Methods: Downstream Analysis Using PROFET.”

mESC permutation test for time 1 and time 2 distribution predictions

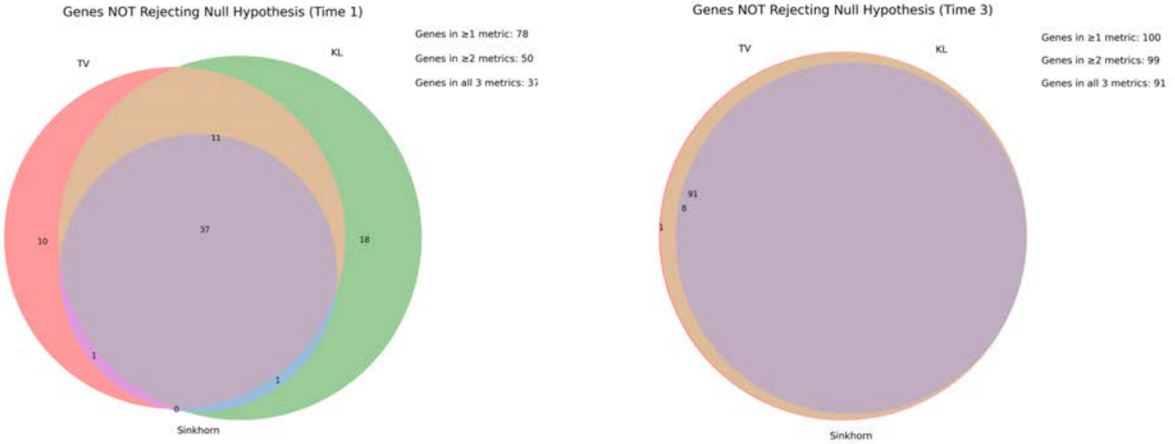

At least one metric (TV, KL, or Sinkhorn) showing No AFF1, BCL3, BHLHE40, BMP2, CARHSP1, CREB3, DDIT3, DNMT3A, DNMT3B, DNMT3L, EGR1, ELF3, EPAS1, ERF, ESRRB, ETV4, ETV5, FOXD3, FOXH1, GLI1, GLI2, HMGA1, HMGA2, HNRNPK, HOPX, ID2, ID3, JARID2, JUN, KDM5B, KLF10, KLF2, KLF3, KLF4, KLF6, L3MBTL3, MSC, MSX2, MTF2, MYBL2, NANOG, NR0B1, PARP1, PEG3, PHB, POU4F2, POU5F1, PURB, RARG, RBPJ, REST, RHOX6, RUNX1, SIX1, SMAD7, SNAI1, SOX2, TBX3, TCF15, TCF7L2, TET1, TET2, TFCP2L1, TFEB, TGIF1, TRP53, UTF1, ZBTB10, ZBTB44, ZC3H7A, ZFHX3, ZFP296, ZFP42, ZFP428, ZFP462, ZFP532, ZFP57, ZFP710

At least two metrics (TV, KL, or Sinkhorn) showing No
BCL3, BHLHE40, CARHSP1, CREB3, DDIT3, EPAS1, ERF, ESRRB, ETV4, ETV5, FOXH1, HMGA1, HNRNPK, HOPX, ID2, JARID2, JUN, KDM5B, KLF10, KLF3, KLF4, KLF6, L3MBTL3, MSC, MSX2, MTF2, MYBL2, PARP1, PEG3, PHB, POU5F1, PURB, RBPJ, REST, RHOX6, SNAI1, TCF15, TCF7L2, TET1, TET2, TFCP2L1, TRP53, UTF1, ZBTB44, ZC3H7A, ZFHX3, ZFP296, ZFP42, ZFP532, ZFP710

All three metrics (TV, KL, and Sinkhorn) showing No
BCL3, CARHSP1, CREB3, DDIT3, ERF, ETV4, HMGA1, HNRNPK, ID2, JARID2, JUN, KLF10, KLF3, KLF6, L3MBTL3, MSC, MSX2, MTF2, MYBL2, PARP1, PEG3, POU5F1, PURB, RBPJ, REST, RHOX6, SNAI1, TCF15, TCF7L2, TET2, TFCP2L1, TRP53, UTF1, ZBTB44, ZFHX3, ZFP296, ZFP532

EMT permutation test for time 1 and time 2 distribution predictions

At least one metric (TV, KL, or Sinkhorn) showing No ANKRD22, AP1M2, AP1M2.1, BSPRY, C1orf116, C1orf172, CDH1, CDH3, CDS1, CLDN4, CLDN7, CRB3, DSP, ELMO3, ENPP5, EPB41L5, EPCAM, EPHA1, EPN3, EPPK1, ERBB3, ESRP1, EVPL, F11R, FN1, FXYD3, GALNT3, GPR110, GPR56, GRHL1, INADL, ITGB6, KLC3, KRTCAP3, MAL2, MAPK13, MMP2, MPP7, MPZL2, MUC1, PRR5, PRSS22, PRSS8, RAB25, RBPMS, S100A14, SCNN1A, SERINC2, SH3YL1, SHROOM3, SPINT2, SSH3, ST14, STAP2, TACSTD2, TC2N, TJP3, TMC4, TMEM125, TMEM45B, TNFRSF21, TSKU

At least two metrics (TV, KL, or Sinkhorn) showing No
DSP, ENPP5, EPB41L5, KRTCAP3, MMP2, RAB25, SERINC2, TMEM45B

All three metrics (TV, KL, and Sinkhorn) showing No
RAB25

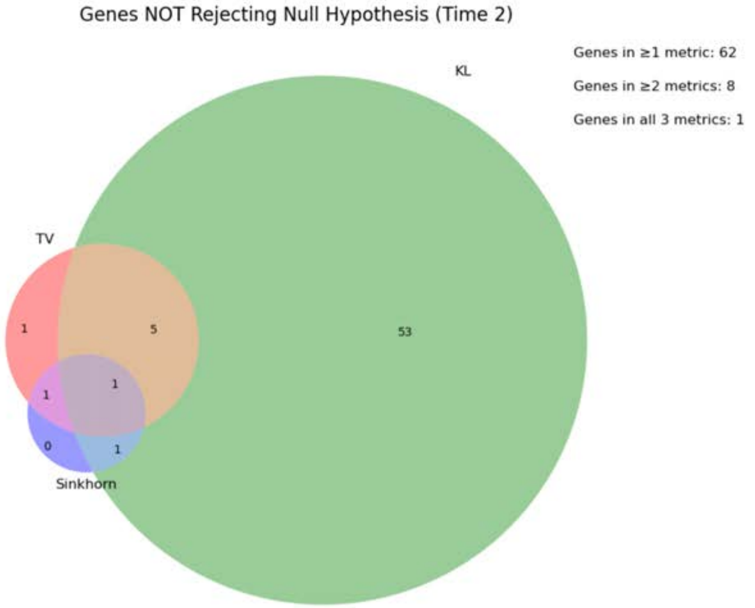

Figure S25, Overlap of genes with no significant differences identified by multiple distributional metrics, related to the section “Methods: Downstream Analysis Using PROFET.” Venn diagram and the list of genes which has not statistically difference through permutation tests under using metrics of KL, TV and Sinkhorm.Venn diagram and gene lists showing overlap among genes with no statistically significant differences based on permutation tests using KL divergence, total variation (TV), and Sinkhorn distance metrics.

A

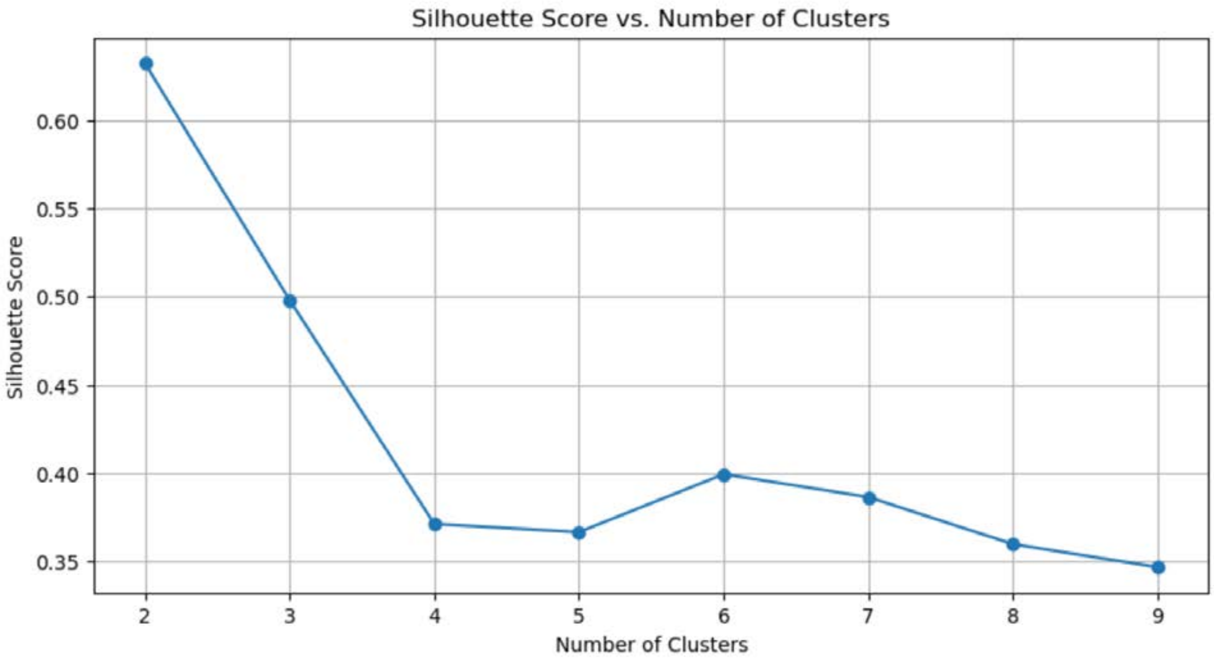

B

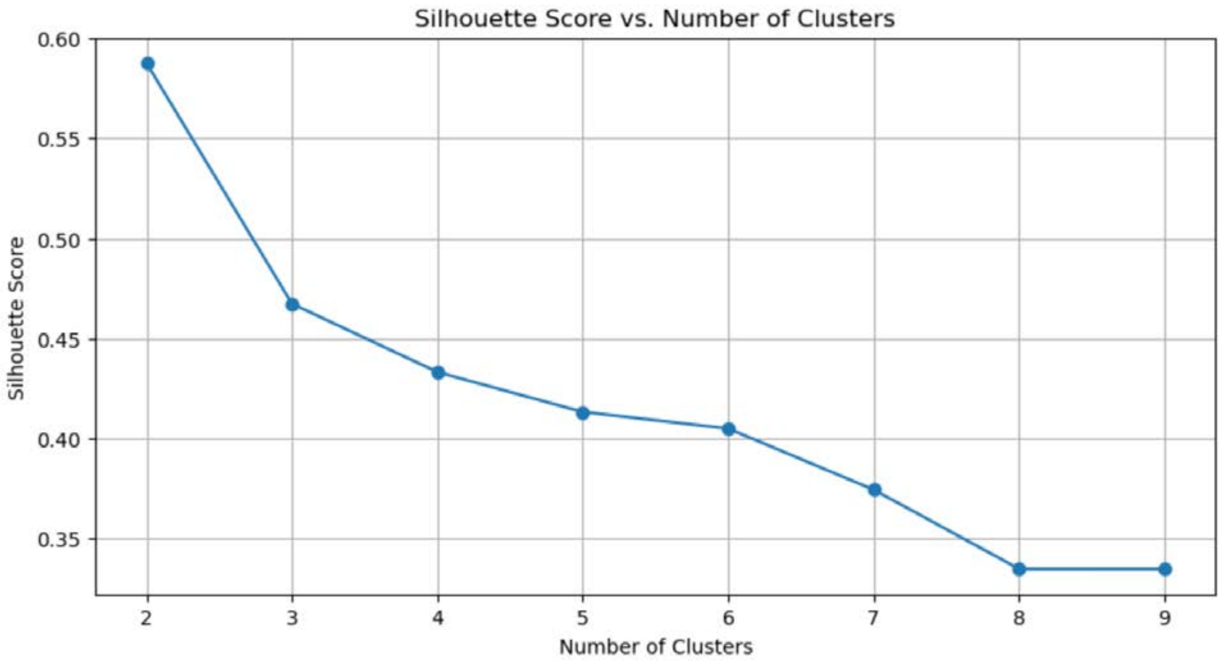

Figure S26, Silhouette scores from clustering analysis across different numbers of clusters, related to the section “Methods: Downstream Analysis Using PROFET.”

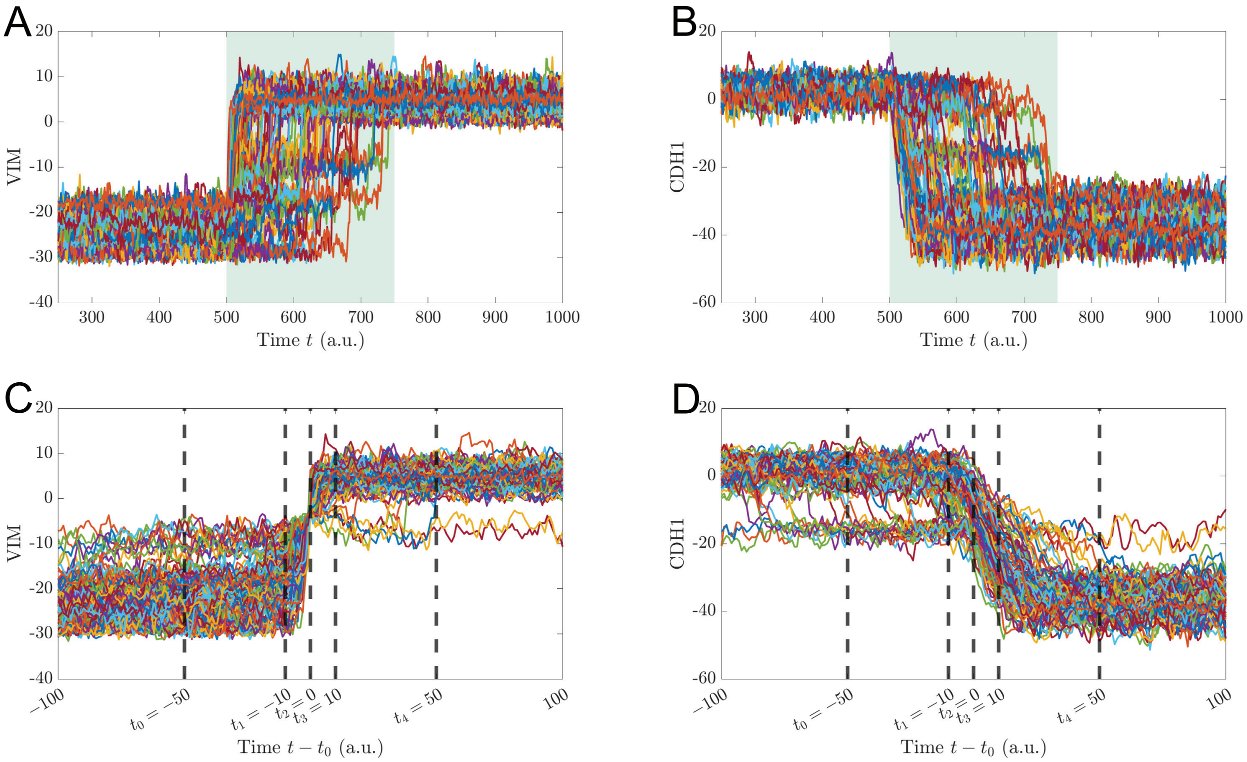

Figure S27: Simulation setup to model epithelial-mesenchymal transition in a population of epithelial cells, related to Supplemental Note 4. (A) Simulated expression levels (log2-transformed) of VIM, a mesenchymal marker, and (B) of CDH1, an epithelial marker, along the simulated trajectories. Each trajectory shown can be interpreted as representing the behavior of a single cell in the population. The shaded region indicates the time period for which the EMT-inducing perturbation (SNAIL up-regulation, miR-34 down-regulation) was active. (C) Same trajectories as in A and (D) as in B shown with shifted time. Each trajectory is shifted by $t_0$, the time point at which the VIM expression along the trajectory (log2-transformed) becomes positive for the first time during the perturbation period.

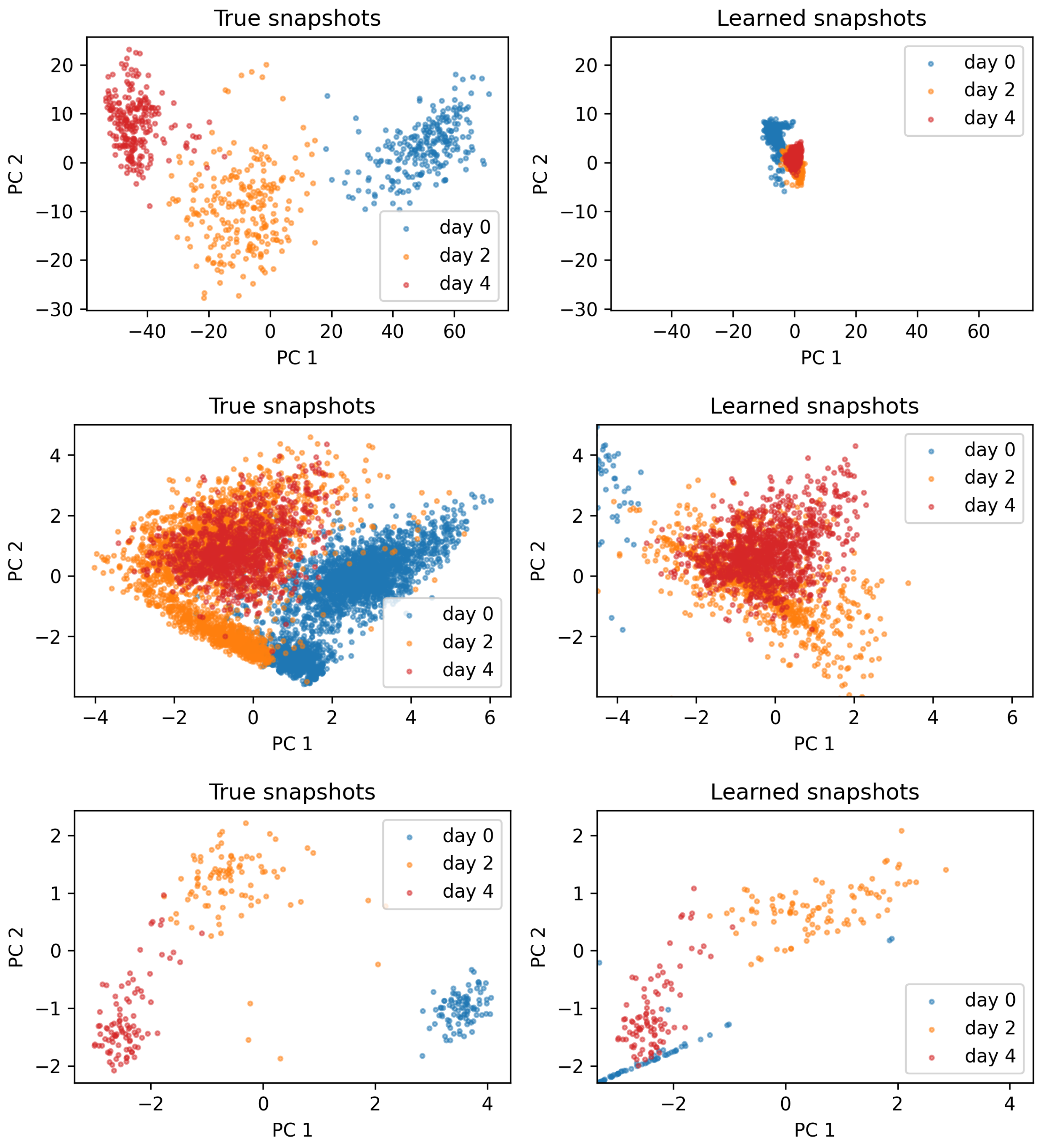

Figure S28: The learned trajectories from TrajectoryNet[9] on synthetic data (top), EMT data (center), and stem cell differentiation data (bottom) diverge substantially from the true trajectories, related to Supplemental Note 6. This trajectory bias worsens as the data support becomes broader, as continuous normalizing flow (CNF)-based methods transport particles through a Gaussian prior, inducing stiffness in the dynamics and pulling trajectories off the data manifold.

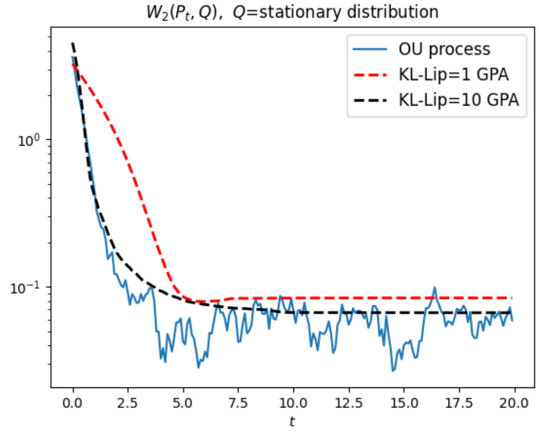

Figure S29: Convergence speed of Lipschitz-regularized KL GPA with different Lipschitz constants $L = 1, 10$ and Ornstein–Uhlenbeck (OU) process, related to Supplemental Note 9. The stationary distribution is Gaussian. With $L = 10$, GPA approximately recovers the convergence rate of the OU process. With $L = 1$, convergence is slower and the converged distribution reflects a proximal solution of the regularized functional, resulting in slightly larger error relative to the true stationary distribution.

**Table S1. Comparison of trajectory inference methods via mathematical framework and theoretical properties, related to the section "Model Validation and Benchmarking Across Four Biological Datasets".** Methods: CellOT, DeepRUOT, MIOFlow, MMFM, PI-SDE, PRESCIENT, TIGON, TrajectoryNet, VGFM, PROFET. Symbols: ✓ = present; ✗ = absent; ~ = partial/limited. N/A = not applicable.

| Feature | CellOT | DeepRUOT | MIOFlow | MMFM | PI-SDE | PRESCIENT | TIGON | TrajectoryNet | VGFM | PROFET |
|---|---|---|---|---|---|---|---|---|---|---|
| **Optimal Transport / Schrodinger Bridge** | ✓ Brenier theorem (Kantorovich dual) [core objective] | ✓ Pretraining: static OT; main: Benamou–Brenier + unbalanced Sinkhorn divergence; Schrodinger bridge connection through added stochasticity [core objective] | ✓ Sinkhorn [core objective] | ✓ Pairwise minibatch OT for coupling only (not in training objective) | ✓ Sinkhorn [core objective] | ✗ | ✓ Benamou–Brenier [core objective] | ✓ Benamou–Brenier [core objective] | ✓ Semi-relaxed OT (Sinkhorn) [core objective] | ✗ |
| **Spline-Based Path Construction** | ✗ | ✗ | ✗ | ✓ Cubic spline interpolation across multiple time-marginals; classifier-free guided flow matching | ✗ | ✗ | ✗ | ✗ | ✗ | ✗ |
| **Wasserstein Gradient Flow** | ✗ | ✗ | ✗ | ✗ | ✗ | ✓ Modeling assumption (does not solve variational problem) | ✗ | ✗ | ✗ | ✓ On a Lipschitz-regularized f-divergence landscape |
| **Unbalancedness** | ✗ | ✓ General convex growth penalty | ✗ | ✗ | ✗ | ✓ Pre-estimated growth weights | ✓ Through WFR | ✓ Pre-estimated growth rates | ✓ growthNet: g = d(ln w)/dt; UOT derives per-cell | ✗ |

| Feature | CellOT | DeepRUOT | MIOFlow | MMFM | PI-SDE | PRESCIENT | TIGON | TrajectoryNet | VGFM | PROFET |
|---|---|---|---|---|---|---|---|---|---|---|
| | | | | | | | | | growth targets from coupling matrix | |
| **Variational Principle** | ✗ | ✓ Action functional: kinetic energy + Fisher information + growth penalty | ✗ | ✗ | ✓ Partially (HJ regularization) | ✗ | ✓ WFR action functional | ✓ Kinetic energy as action functional | ✗ | ✓ Gradient descent on f-divergence functional |
| **Static / Dynamic** | Static | ODE, SDE | ODE (optional SDE) | ODE | SDE | SDE | Deterministic ODE (CNF + growth) | ODE | Deterministic ODE (velocity + growth) | Deterministic ODE (gradient flow + force matching) |

**Table S2. Comparison of trajectory inference methods via training methodology, related to the section "Model Validation and Benchmarking Across Four Biological Datasets".** Methods: CellOT, DeepRUOT, MIOFlow, MMFM, PI-SDE, PRESCIENT, TIGON, TrajectoryNet, VGFM, PROFET. Symbols: ✓ = present; ✗ = absent; ~ = partial/limited. N/A = not applicable.

| Feature | CellOT | DeepRUOT | MIOFlow | MMFM | PI-SDE | PRESCIENT | TIGON | TrajectoryNet | VGFM | PROFET |
|---|---|---|---|---|---|---|---|---|---|---|
| **Statics: Map-based** | ✓ Transport map ∇g(x) | N/A | N/A | N/A | N/A | N/A | N/A | N/A | N/A | N/A |
| **Statics: Coupling-based** | ✗ | N/A | N/A | N/A | N/A | N/A | N/A | N/A | N/A | N/A |
| **Dynamics: Adjoint-based** | N/A | ✗ | ✗ | ✗ | ✗ | ✗ | ✓ | ✓ | ✗ | ✗ |
| **Dynamics: DTO (discretize-then-optimize, global backprop)** | N/A | ✓ Main training stage | ✓ | ✗ | ✓ | ✓ | ✗ | ✗ | ✓ Refinement step (ODE simulation + distribution fitting) | ✗ |
| **Dynamics: Greedy forward (backprop-free)** | N/A | ✗ | ✗ | ✗ | ✗ | ✗ | ✗ | ✗ | ✗ | ✓ GPA step |
| **Dynamics: Simulation-free (flow matching)** | N/A | ✓ Pretraining stage | ✗ | ✓ | ✗ | ✗ | ✗ | ✗ | ✓ | ✓ Force-matching step |

**Table S3. Wasserstein-2 prediction error across benchmark models and datasets (five intermediate-timepoint snapshots) and comparison with PROFET, related to the section "Model Validation and Benchmarking Across Four Biological Datasets".** The MIN and MAX values were computed excluding the PROFET row so that they could be compared with PROFET.

| Models | mESC Day 1 | | mESC Day 3 | | EMT Day 2 | | Axolotl Day 1 | | Axolotl Day 3 | | Geometric mean |
|---|---|---|---|---|---|---|---|---|---|---|---|
| | W2 error | ratio | W2 error | ratio | W2 error | ratio | W2 error | ratio | W2 error | ratio | ratio |
| PROFET | 0.0971 | 1.0000 | 0.0940 | 1.0000 | 0.9556 | 1.0000 | 5.9992 | 1.0000 | 7.9746 | 1.0000 | 1.0000 |
| WOT | 0.5311 | 5.4689 | 0.1747 | 1.8589 | 1.2602 | 1.3188 | 30.8480 | 5.1420 | 31.3764 | 3.9345 | 3.0667 |
| TIGON | 0.1644 | 1.6933 | 0.2545 | 2.7078 | 4.2535 | 4.4511 | 28.0051 | 4.6681 | 136.7753 | 17.1514 | 4.3919 |
| PRESCIENT | 3.6045 | 37.1153 | 10.6093 | 112.8979 | 0.8933 | 0.9348 | 40.3402 | 6.7242 | 95.5057 | 11.9762 | 12.5831 |
| MIOFlow | 0.6040 | 6.2199 | 1.7971 | 19.1239 | 1.9054 | 1.9939 | 46.3555 | 7.7269 | 27.9711 | 3.5075 | 5.7758 |
| VGFM | 2.9638 | 30.5186 | 0.4514 | 4.8032 | 2.0040 | 2.0970 | 4.1996 | 0.7000 | 45.2601 | 5.6755 | 4.1435 |
| MMFM | 0.2010 | 2.0698 | 0.3211 | 3.4168 | 1.8650 | 1.9516 | 50.2037 | 8.3683 | 15.3080 | 1.9196 | 2.9455 |
| PI-SDE | 0.2604 | 2.6816 | 0.2419 | 2.5740 | 1.6987 | 1.7776 | 19.8972 | 3.3166 | 21.9103 | 2.7475 | 2.5686 |
| DeepRUOT | 0.3570 | 3.6757 | 0.3189 | 3.3933 | 1.7260 | 1.8062 | N/A | N/A | N/A | N/A | 2.8243 |
| Null model | 0.5287 | 5.4443 | 0.2130 | 2.2666 | 2.0325 | 2.1269 | 21.0868 | 3.5149 | 21.4725 | 2.6926 | 3.0132 |
| TrajectoryNet | N/A | N/A | N/A | N/A | N/A | N/A | N/A | N/A | N/A | N/A | N/A |
| MIN | 0.1644 | 1.6933 | 0.1747 | 1.8589 | 0.8933 | 0.9348 | 4.1996 | 0.7000 | 15.3080 | 1.9196 | **2.5686** |
| MAX | 3.6045 | 37.1153 | 10.6093 | 112.8979 | 4.2535 | 4.4511 | 50.2037 | 8.3683 | 136.7753 | 17.1514 | **12.5831** |

**Table S4. Comparison of trajectory inference methods via runtime and memory, related to the section "Ablation Analysis, Robustness, and Computational Scalability".** Runtime and memory analyses on three benchmarks (stem_cell_differentiation, emt_72, Larry_3000) across dimensions d=2–16. **Abbreviations:** OT = Optimal Transport; WFR = Wasserstein–Fisher–Rao; CNF = Continuous Normalizing Flow; SDE = Stochastic Differential Equation; ODE = Ordinary Differential Equation; HJ = Hamilton–Jacobi; GPA = Gradient Particle Approximation; DTO = Discretize-then-Optimize; UOT = Unbalanced Optimal Transport.

##### a. Summary

| Feature | CellOT | DeepRUOT | MIOFlow | MMFM | PI-SDE | PRESCIENT | TIGON | TrajectoryNet | VGFM | PROFET |
|---|---|---|---|---|---|---|---|---|---|---|
| **Runtime Range (across benchmarks)** | 2,024–2,349 s | 93–5,243 s | 70–2,172 s | 99–187 s | 417–2,622 s | 384–1,356 s | 1,406–36,388 s | 16,508–92,686 s | 16–343 s | 423–2,743 s |
| **Peak CPU RAM Range** | 0.43–6.91 GB | 0.87–7.41 GB | 1.13–6.72 GB | 0.62–0.84 GB | 0.32–0.59 GB | 0.24–0.60 GB | 0.52–2.15 GB | 0.29 GB (partial) | 0.63–6.83 GB | 0.36–3.95 GB |
| **Scalability Notes** | Static map; no ODE integration | DTO; OOM risk in eval (autograd graph accumulation at large scale) | DTO, Coarse ODE step size (RK4); scales with dataset | Fully simulation-free; fast and memory-stable | DTO (Euler-Maruyama); low memory | DTO; low memory | Adjoint-based; Slow, runtime scales steeply with data size | Adjoint-based; very slow, 10–100x slower than others | Simulation-free pretraining + coarse Euler; fast but memory scales with data | Greedy forward (backprop-free) + simulation-free Step 2; moderate speed |

##### b. Time and Computational Resources

| Method | Benchmark (dim) | Training sample size | Runtime | Peak CPU RAM | Notes |
|---|---|---|---|---|---|
| **MMFM** | stem_cell_diff (d=2) | 278 | 183 s | 0.620 GB | Fully simulation-free training |
| | stem_cell_diff (d=4) | 278 | 184 s | 0.623 GB | |
| | stem_cell_diff (d=8) | 278 | 185 s | 0.622 GB | |
| | stem_cell_diff (d=16) | 278 | 186 s | 0.619 GB | |
| | emt_72 (d=2) | 3,881 | 159 s | 0.637 GB | |
| | Larry_3000 (d=2) | 34,317 | 99 s | 0.839 GB | |
| **MIOFlow-PCA** | stem_cell_diff (d=2) | 278 | 71 s | 1.282 GB | Coarse ODE step size (RK4) |
| | stem_cell_diff (d=4) | 278 | 71 s | 1.240 GB | |
| | stem_cell_diff (d=8) | 278 | 71 s | 1.264 GB | |
| | stem_cell_diff (d=16) | 278 | 71 s | 1.270 GB | |
| | emt_72 (d=2) | 3,881 | 246 s | 1.467 GB | |
| | Larry_3000 (d=2) | 34,317 | 2,172 s | 6.704 GB | |

| Method | Benchmark (dim) | Training sample size | Runtime | Peak CPU RAM | Notes |
|---|---|---|---|---|---|
| **MIOFlow-PHATE** | stem_cell_diff (d=2) | 278 | 70 s | 1.271 GB | Coarse ODE step size (RK4) |
| | stem_cell_diff (d=4) | 278 | 70 s | 1.279 GB | |
| | stem_cell_diff (d=8) | 278 | 73 s | 1.153 GB | |
| | stem_cell_diff (d=16) | 278 | 73 s | 1.198 GB | |
| | emt_72 (d=2) | 3,881 | 245 s | 1.816 GB | |
| | Larry_3000 (d=2) | 34,317 | 2,129 s | 6.720 GB | |
| **MIOFlow-GAE** | stem_cell_diff (d=2) | 278 | 104 s | 1.263 GB | Coarse ODE step size (RK4) |
| | stem_cell_diff (d=4) | 278 | 105 s | 1.282 GB | |
| | stem_cell_diff (d=8) | 278 | 107 s | 1.125 GB | |
| | stem_cell_diff (d=16) | 278 | 117 s | 1.167 GB | |
| | emt_72 (d=2) | 3,881 | 235 s | 1.704 GB | |
| | Larry_3000 (d=2) | 34,317 | 2,141 s | 5.713 GB | |
| **PRESCIENT** | stem_cell_diff (d=2) | 278 | 384 s | 0.482 GB | Explicit Euler, no adjoint |
| | stem_cell_diff (d=4) | 278 | 401 s | 0.484 GB | |
| | stem_cell_diff (d=8) | 278 | 428 s | 0.484 GB | |
| | stem_cell_diff (d=16) | 278 | 466 s | 0.480 GB | |
| | emt_72 (d=2) | 3,881 | 517 s | 0.602 GB | |
| | Larry_3000 (d=2) | 34,317 | 1,356 s | 0.242 GB | |
| **VGFM** | stem_cell_diff (d=2) | 278 | 16 s | 0.628 GB | Simulation-free pretraining + coarse Euler main |
| | stem_cell_diff (d=4) | 278 | 16 s | 0.630 GB | |
| | stem_cell_diff (d=8) | 278 | 16 s | 0.628 GB | |
| | stem_cell_diff (d=16) | 278 | 16 s | 0.629 GB | |
| | emt_72 (d=2) | 3,881 | 24 s | 0.834 GB | |
| | Larry_3000 (d=2) | 34,317 | 343 s | 6.831 GB | |
| **DeepRUOT** | stem_cell_diff (d=2) | 278 | 100 s | 0.879 GB | Pretraining: 100 steps/interval; main: 10 steps, no adjoint |
| | stem_cell_diff (d=4) | 278 | 99 s | 0.873 GB | |
| | stem_cell_diff (d=8) | 278 | 93 s | 0.879 GB | |
| | stem_cell_diff (d=16) | 278 | 113 s | 0.887 GB | |
| | emt_72 (d=2) | 3,881 | 2,162 s | 2.176 GB | OOM risk in eval (autograd graph accumulation) |

| Method | Benchmark (dim) | Training sample size | Runtime | Peak CPU RAM | Notes |
|---|---|---|---|---|---|
| | Larry_3000 (d=2) | 34,317 | 5,243 s | 7.412 GB | OOM risk in eval |
| **PI-SDE** | stem_cell_diff (d=2) | 278 | 418 s | 0.328 GB | Euler SDE, no adjoint |
| | stem_cell_diff (d=4) | 278 | 422 s | 0.320 GB | |
| | stem_cell_diff (d=8) | 278 | 417 s | 0.326 GB | |
| | stem_cell_diff (d=16) | 278 | 424 s | 0.321 GB | |
| | emt_72 (d=2) | 3,881 | 985 s | 0.369 GB | |
| | Larry_3000 (d=2) | 34,317 | 2,622 s | 0.591 GB | |
| **TIGON** | stem_cell_diff (d=2) | 278 | 1,809 s | 0.525 GB | |
| | stem_cell_diff (d=4) | 278 | 1,406 s | 0.515 GB | |
| | stem_cell_diff (d=8) | 278 | 2,061 s | 0.573 GB | |
| | stem_cell_diff (d=16) | 278 | 2,950 s | 0.658 GB | |
| | emt_72 (d=2) | 3,881 | 6,725 s | 0.743 GB | |
| | Larry_3000 (d=2) | 34,317 | 36,388 s | 2.146 GB | |
| **TrajectoryNet** | stem_cell_diff (d=2) | 278 | 42,436 s | N/A | |
| | stem_cell_diff (d=4) | 278 | 40,540 s | 0.293 GB | |
| | stem_cell_diff (d=8) | 278 | 57,788 s | 0.292 GB | |
| | stem_cell_diff (d=16) | 278 | 82,595 s | 0.293 GB | |
| | emt_72 (d=2) | 3,881 | 16,508 s | N/A | |
| | Larry_3000 (d=2) | 34,317 | 92,686 s | N/A | |
| **PROFET** | stem_cell_diff (d=2) | 278 | 487 s | 0.362 GB | Greedy forward (backprop-free) Step 1 + simulation-free Step 2 |
| | stem_cell_diff (d=4) | 278 | 619 s | 0.363 GB | |
| | stem_cell_diff (d=8) | 278 | 1,033 s | 0.362 GB | |
| | stem_cell_diff (d=16) | 278 | 1,085 s | 0.361 GB | |
| | emt_72 (d=2) | 3,881 | 423 s | 0.470 GB | |
| | Larry_3000 (d=2) | 34,317 | 2,743 s | 3.954 GB | |
| **CellOT** | emt_72 (d=2) | 3,881 | 2,024 s | 0.427 GB | |
| | Larry_3000 (d=2) | 34,317 | 2,349 s | 6.907 GB | |

**Table S5. Comparison of predicted and observed gene expression distributions at the held-out time points (time points 1 and 3) in the mESC dataset, related to Figure 3G.** – provided as a standalone file

**Table S6. Comparison of predicted and observed gene expression distributions at the held-out time point (day 2) in the EMT dataset, related to Figure 4E.** – provided as a standalone file

**Table S7. List of 121 genes previously implicated in the response to palbociclib, including genes involved in cell cycle regulation, MYC and estrogen receptor (ER) signaling, growth factor pathways, the Hippo pathway, and inflammatory processes related to the section "Tracing Phenotypic Shifts Associated with Palbociclib Therapy in HR+ Breast Cancer".** – provided as a standalone file

**Table S8. Coefficients of variation (CVs) of the predicted cell-state trajectory displacements for the in vitro dataset and patients PA3, 862, and 887, related to Figure 5B.**

| experiment label | mean_displacement | std_displacement | iqr_displacement | cv_displacement | entropy | experiment memo |
|---|---|---|---|---|---|---|
| Patient_887 | 1.0204561 | 0.26583105 | 0.32117099 | 0.2605022 | 3.27391317 | Palbo_887_nofibroblast_malignant_Rgene_dim2-f_Lip=5e-2-t_size=50-network=64_64_64 |
| Patient_PA3 | 1.9689372 | 0.8479199 | 1.44968212 | 0.43064854 | 3.47874554 | Palbo_BMC_nofibroblast_malignant_Rgene_dim2-f_Lip=5e-2-t_size=50-network=64_64_64 |
| Patient_862 | 1.852924 | 0.5552303 | 0.98318267 | 0.29965088 | 3.37245175 | Palbo_862_nofibroblast_malignant_Rgene_dim2-f_Lip=5e-2-t_size=50-network=64_64_64 |
| MCF7A | 3.4400735 | 0.35988832 | 0.57818079 | 0.10461646 | 3.57442794 | Palbo_NDPR_nofibroblast_malignant_Rgene_dim2-f_Lip=5e-2-t_size=50-network=64_64_64 |

**Table S9. Differential gene expression (DEG) analysis comparing post-treatment and pre-treatment cells in the in vitro dataset and patients PA3, 862, and 887, related to Figure 6E.** – provided as a standalone file

**Table S10. Immune cell marker genes used for inferCNV analysis of patient PA3 from Luo et al, related to the section "Methods: Preprocessing of breast cancer patient data".**

| marker | type | details |
|---|---|---|
| PTPRC | immune | CD45 |
| CD2 | immune | T CELLS & NK CELLS |
| CD3G | immune | T CELLS |
| CD8A | immune | CYTOTOXIC T CELLS |
| CD4 | immune | HELPER T CELLS |
| CD19 | immune | B CELLS |
| MS4A1 | immune | B CELLS; CD20 |
| CD79A | immune | B CELLS |
| CD79B | immune | B CELLS |
| CD14 | immune | MACROPHAGES, NEUTROPHILS, DC |
| CD68 | immune | MACROPHAGES, MONOCYTES |
| CD163 | immune | MACROPHAGES, MONOCYTES |
| CSF1R | immune | MACROPHAGES, MONOCYTES |
| NCAM1 | immune | NK CELLS; CD56 |
| ITGAX | immune | DC; CD11C |
| IL3RA | immune | DC; CD123 |
| LY6G6D | immune | NEUTROPHILS, MONOCYTES, GRANULOCYTES |
| FUT4 | immune | NEUTROPHILS, MYELOID CELLS; CD15 |
| FCGR3A | immune | NK CELLS, NEUTROPHILS, MONOCYTES, MACROPHAGES; CD16 |
| ITGAM | immune | NK CELLS, NEUTROPHILS, MONOCYTES, GRANULOCYTES, MACROPHAGES; CD11B |
| CD34 | immune | PROGENITOR |

### Supplemental Notes

#### Supplemental Note 1: Wasserstein Gradient Flow and Lipschitz-Regularized KL Divergence

*This note is related to the sections "Overview of PROFET" and "Gradient flow simulation with Generative Particles Algorithm (GPA)".*

We briefly review the mathematical foundation behind the Generative Particles Algorithm (GPA), which simulates Wasserstein gradient flows driven by divergences between probability measures.

**Wasserstein gradient flow of probability measures.** Let $\rho_t$ be a time-indexed family of probability distributions evolving over $\mathbb{R}^d$. The Wasserstein gradient flow describes their evolution as a solution to the continuity equation

$$\partial_t \rho_t + \nabla \cdot (\rho_t v_t) = 0, \tag{1}$$

where the velocity field $v_t$ is given by the Wasserstein gradient of an energy functional $\mathcal{F} : \mathcal{P}(\mathbb{R}^d) \to \mathbb{R}$, namely:

$$v_t = -\nabla \frac{\delta \mathcal{F}}{\delta \rho}(\rho_t), \tag{2}$$

with $\frac{\delta \mathcal{F}}{\delta \rho}(\rho_t)$ denoting the first variation of $\mathcal{F}$ with respect to $\rho_t$. This framework provides a natural interpretation of dynamics over probability measures as steepest descent flows in the space of distributions.

**KL divergence and its regularization.** A common choice for the energy functional is the Kullback–Leibler (KL) divergence, defined as $\mathrm{KL}(\rho_t \| \pi) = \mathbb{E}_{\rho_t}\left[\log\left(\frac{d\rho_t}{d\pi}\right)\right]$, where $\pi$ is a fixed target distribution. This formulation requires that $\rho_t$ is absolutely continuous with respect to $\pi$ (i.e., $\rho_t \ll \pi$), so that the Radon–Nikodym derivative $\frac{d\rho_t}{d\pi}$ exists. In practice, both $\rho_t$ and $\pi$ may be represented only by samples, and neither density may be available in closed form.

In such cases, we instead consider the variational formulation of f-divergences, which allows estimation directly from samples:

$$D_f(\rho_t \| \pi) = \sup_{\phi \in C_b(\mathbb{R}^d)} \left\{ \mathbb{E}_{\rho_t}[\phi] - \inf_{\nu \in \mathbb{R}} \mathbb{E}_\pi[f^\star(\phi - \nu) + \nu] \right\}, \tag{3}$$

where $f(x) = x \log x$ for KL divergence and $f^\star$ is its Legendre transform. This formulation does not require access to densities or absolute continuity with respect to Lebesgue measure, making it particularly suitable for sample-based approximation.

However, for empirical distributions, KL divergence may diverge or become non-differentiable due to lack of absolute continuity. To address this, we use a Lipschitz-regularized $f$-divergence with $f(x) = x \log x$, which remains well-defined even for discrete measures. It admits a dual variational representation:

$$D_f^{\mathsf{Lip}_L(\mathbb{R}^d)}(\rho_t \| \pi) = \sup_{\phi \in \mathsf{Lip}_L(\mathbb{R}^d)} \left\{ \mathbb{E}_{\rho_t}[\phi] - \inf_{\nu \in \mathbb{R}} \mathbb{E}_\pi[f^\star(\phi - \nu) + \nu] \right\}, \tag{4}$$

where $f^\star$ is the Legendre transform of $f$, and the test function $\phi$ is constrained to be $L$-Lipschitz. The optimal potential $\phi_t^{L,*}$ obtained from this formulation approximates the first variation $\frac{\delta \mathcal{F}}{\delta \rho}(\rho_t)$, ensuring that the induced velocity field $v_t = -\nabla \phi_t^{L,*}$ is both numerically stable and biologically plausible.

This formulation avoids the need for explicit densities or absolute continuity with respect to Lebesgue measure, and its robustness under minimal moment assumptions is further strengthened when combined with Lipschitz constraints, as discussed in Supplemental Note 2.

#### Supplemental Note 2: Robustness through Lipschitz regularization

*This note is related to the sections "Overview of PROFET" and "The role of Lipschitz regularization in empirical gradient flow modeling ".*

In Wasserstein gradient flows driven by KL divergence, the velocity field is given by the gradient of the first variation of the divergence functional. For the unregularized KL divergence, this first variation is the log-density ratio:

$$\frac{\delta}{\delta\rho}\mathrm{KL}(\rho\|\pi) = \log\left(\frac{d\rho}{d\pi}\right), \tag{5}$$

which defines the velocity field as:

$$v_t(x) = -\nabla \log\left(\frac{d\rho_t}{d\pi}\right). \tag{6}$$

However, this expression is only well-defined when the source distribution $\rho_t$ is absolutely continuous with respect to the target distribution $\pi$. In practical applications—especially when $\rho_t$ is an empirical distribution represented by discrete particles—this condition typically fails, resulting in an infinite KL divergence and an undefined first variation. As a result, the velocity field breaks down and cannot be used for gradient flow simulation.

Lipschitz regularization overcomes this limitation by defining a relaxed divergence:

$$D_f^{\mathsf{Lip}_L}(\rho_t\|\pi) = \sup_{\phi\in\mathsf{Lip}_L}\left\{\mathbb{E}_{\rho_t}[\phi] - \inf_{\nu\in\mathbb{R}}\mathbb{E}_\pi[f^\star(\phi-\nu)+\nu]\right\}, \tag{7}$$

which remains finite and admits a well-defined first variation as long as $\rho_t$ has finite first moment[1]—even when $\rho_t$ is not absolutely continuous w.r.t. $\pi$, and even when $\pi$ is heavy-tailed, supported on low-dimensional manifolds, or discrete, as in our case. The existence of the first variation is essential in Wasserstein gradient flows; without it, the velocity field is not well-defined. This allows our method, which learns dynamics through Lipschitz-regularized gradient flows, to remain agnostic to the parametric form or regularity of the target distribution, enabling a fully data-driven modeling of transitions. The optimal potential $\phi_t^{L,*}$ serves as a proxy for $\log\left(\frac{d\rho_t}{d\pi}\right)$; more precisely, $\phi_t^{L,*}$ approximates $\log\left(\frac{d\rho_t}{d\pi}\right)$ using $\log\left(\frac{d\sigma_t}{d\pi}\right)$, where $\sigma_t$ is an intermediate measure that minimizes the KL divergence to $\pi$ while remaining close to $\rho_t$ in Wasserstein-1 distance. This formulation effectively estimates the gradient of the log-likelihood while maintaining robustness and boundedness.

Furthermore, the Lipschitz constraint implies a uniform bound on the velocity magnitude: since $\phi_t^{L,*}\in\mathrm{Lip}_L(\mathbb{R}^d)$, we have $\|\nabla\phi_t^{L,*}(x)\| \le L$ for all $x$ and $t$. This finite speed bound is critical for the forward Euler discretization used in GPA: in analogy with the Courant–Friedrichs–Lewy (CFL) condition, a bounded velocity ensures that the maximum particle displacement per step is bounded by $L\Delta t$, preventing overshoot and guaranteeing stability without requiring implicit solvers or adaptive step-size controllers.

Consequently, the velocity field

$$v_t(x) = -\nabla\phi_t^{L,*}(x) \tag{8}$$

remains numerically stable throughout training and enables particle evolution under ill-posed conditions that would render the standard KL-based gradient flow inapplicable.

#### Supplemental Note 3: Comparison to flow matching in score-based generative modeling

*This note is related to the section "Overview of PROFET".*

After GPA simulation, we learn a continuous-time Eulerian velocity field $v_\theta(x,s)$ using force-matching. This distills a sequence of local, time-indexed potentials $\{\phi_n^{L,*}\}$ into a single neural vector field, enabling amortized learning of the global dynamics.

Structurally, our force-matching loss closely resembles the denoising score matching (DSM) loss used in score-based generative models. The DSM objective takes the form:

$$\mathcal{L}_{\mathsf{DSM}}(\theta) = \mathbb{E}_{t\sim\mathcal{U}(0,T)}\mathbb{E}_{x_0\sim p\mathsf{data},\, x_t\sim\mathcal{N}(x_0,\sigma_t^2 I)}\left[\left|s_\theta(x_t,t) - \frac{1}{\sigma_t^2}(x_0-x_t)\right|^2\right], \tag{9}$$

where a score network $s_\theta$ is trained to recover the gradient of the log-density at perturbed inputs, assuming an analytically defined forward process (e.g., Gaussian noise).

In contrast, our force-matching objective:

$$\mathbb{E}_{(x,s)}\left[\left|v_\theta(x,s)+\nabla\phi_s^{L,*}(x)\right|^2\right], \tag{10}$$

supervises the neural velocity field using gradients derived from GPA-based simulations, without relying on a parametric form for the forward dynamics.

This key distinction enables our approach to flexibly model complex systems in which analytic transition distributions (e.g., Gaussian noise or OT interpolation) are inadequate or biologically unrealistic. Whereas traditional flow-matching methods[2] and score-based models[3] typically impose Gaussian smoothing or regular OT priors, our method infers transitions directly from empirical distributions. This allows it to capture biologically relevant phenomena such as bifurcations, convergence to terminal states, or heterogeneous treatment responses—dynamics that arise naturally in developmental and disease progression settings.

#### Supplemental Note 4: Simulation setup to model epithelial-mesenchymal transition

*This note is related to the section "Overview of PROFET".*

**Capturing biological variability in EMT dynamics** Various studies profiling EMT at single-cell resolution have reported heterogeneity in gene expression dynamics even in a genetically homogeneous population[4–6]. To capture this behavior, we simulated Eq. (6) (Methods) for an ensemble of parameter sets and for multiple random initial conditions for each parameter set in the ensemble. Parameter sets in the ensemble were sampled in a manner that captures the range of biologically plausible behaviors (see[7] for a detailed description of the sampling scheme). In the absence of noise (*i.e.*, for $D = 0$), the EMT gene regulatory network in our setup exhibits multi-stability, exhibiting stable states that span the spectrum from epithelial to mesenchymal phenotypes[8]. To simplify the analysis, we consider only those parameter sets that exhibit tri-stability for $D = 0$; the three steady states for each parameter set can correspond to epithelial, hybrid epithelial / mesenchymal, and mesenchymal phenotypes which are the key categories of phenotypic states studied in EMT. Parameter sets with fewer or more stable states when no noise is present were discarded from the ensemble. For each such parameter set, we simulated the dynamics starting from $10$ random initial conditions from $t = 0$ to $t = 1000$ (shown in Figure S27AB). Each simulated trajectory in our setup can then be interpreted as corresponding to the dynamical behavior of a single cell. Note that the time here is simulation time and its mapping to real time will depend on the timescale associated with different regulatory interactions in the gene network.

**Simulating perturbation-induced transition from epithelial to mesenchymal state** Simulating dynamics for a given parameter set starting from a random initial condition can result in a trajectory that leads to an epithelial, hybrid, or mesenchymal state. Since we want to focus on transition from epithelial to mesenchymal states, we discarded trajectories that failed to remain in an epithelial state (characterized by high expression of CDH1 and low expression of VIM) between $t = 200$ and $t = 500$. At $t = 500$, we introduced a perturbation that can trigger a transition from an epithelial state to a mesenchymal state: 100-fold increase in the production rate of SNAIL and 100-fold decrease in the production rate of miR-34. This perturbation lasted till $t = 750$ at which point it was withdrawn and SNAIL and miR-34 production rates restored to their original values (see shaded region in Figure S27AB). The perturbation causes cells (represented by trajectories) to start transitioning to mesenchymal states characterized by low expression of CDH1 and high expression of VIM. All cells in our setup do not transition at the same time within the perturbation window, consistent with experimental data[6], and cells can remain in the mesenchymal state even after the EMT-inducing perturbation has been withdrawn (*i.e.*, for $t > 750$). Note that all cells in our simulation setup do not transition to a mesenchymal state in response to the perturbation and among those that do transition, not all remain in the mesenchymal state after the perturbation is withdrawn; trajectories corresponding to such behavior were also discarded. Figure S27AB show numerous simulated trajectories retained in the ensemble and their response to the perturbation.

To generate a synthetic dataset for PROFET training, we aligned the simulated trajectories by the time point at which the log2-transformed VIM expression level first becomes positive (representing the transition to a more mesenchymal state). Figure S27C-D show the trajectories shifted by their transition time points. This step is equivalent to the pseudotime analysis done in various single-cell studies to group together cells that are at different stages of transition from one phenotype to another[5;6], and is needed since all cells in a population do not undergo phenotypic transition at the same rate (a behavior that is recapitulated in our simulation setup). Thereafter, we sampled the simulated cell population at five points in shifted time (indicated by dashed vertical lines in Figure S27C-D). Sampled cell states at these time points (before, during, and after EMT) constituted the synthetic longitudinal dataset that was then used to train the PROFET model and to test the performance of the overall approach.

**Use of synthetic data for hyperparameter selection** The synthetic EMT dataset described above was further used to guide the selection of key hyperparameters in the PROFET framework. Specifically, we leveraged the known temporal structure of the simulated trajectories to perform systematic validation using held-out intermediate time points. From the simulated trajectories, we sampled five discrete time points to mimic temporally sparse scRNA-seq measurements. PROFET was then trained using a subset of these time points (e.g., days 0, 2, and 4), while the remaining intermediate time points (e.g., days 1 and 3) were withheld for evaluation.

Hyperparameters governing both the GPA simulation and the force-matching step—including the stopping criterion, Lipschitz regularization strength, latent dimensionality, and network architecture—were empirically tuned to ensure that the reconstructed trajectories accurately matched the distributions at these withheld time points. In particular, we selected parameter ranges that produced stable, non-divergent dynamics while minimizing distributional discrepancies (measured by $W_2$, Sinkhorn divergence, and MMD) between predicted and simulated cell states.

The resulting hyperparameter configurations were found to be consistent across different realizations of the synthetic dataset and were subsequently fixed for application to real datasets. This procedure provides a principled strategy for parameter selection by leveraging a controlled setting in which ground-truth temporal dynamics are known, ensuring that the learned model generalizes to experimentally observed biological systems.

#### Supplemental Note 5: Full description of novelty

*This note is related to the section "Model Validation and Benchmarking Across Four Biological Datasets".*

**Gradient flow-based modeling from empirical data.** We propose an agnostic and data-driven framework for modeling temporal dynamics via Wasserstein gradient flows. At the core of our approach is GPA, which simulates particle trajectories that follow the gradient flow of an energy functional—specifically, a Lipschitz-regularized KL divergence. Unlike OT-based methods that optimize static couplings or CNFs that rely on a predefined base distribution, our method directly learns smooth dynamics from solely empirical source and target distributions, without assuming specific distributional forms or mechanistic priors. The learned trajectories interpolate between observed single-cell snapshots in a fully nonparametric framework that remains robust to complex, irregular, or discrete data supports, enabling flexible and interpretable modeling of biological processes.

**Robustness and stability via Lipschitz-regularzied divergence.** A key strength of our method lies in the robustness and stability of the Lipschitz-regularized KL divergence used to define the gradient flow. This regularization ensures that the induced velocity field is bounded by a constant $L$, which makes the simulation numerically stable and imposes a biologically sensible constraint on transport speeds—unlike unregularized OT flows with unbounded gradients or diffusion models with predefined stochasticity. As shown in[1], the resulting divergence $D_f^{\Gamma_L}(\rho\|\pi)$ is well-defined and admits stable variational gradients for arbitrary target distributions $\pi$, as long as the current distribution $\rho$—which evolves step-by-step during the GPA simulation—has finite first moment. This makes the method particularly robust in challenging settings where the distributions are heavy-tailed, lie on low-dimensional or fractal manifolds, or are purely discrete—as is often the case in single-cell data.

**Adaptive resolution and robust trajectory learning.** The dissipative structure of Wasserstein gradient flows naturally provides a criterion for adaptive resolution: as the particle distribution approaches the target, the expected kinetic energy decays (Theorem 2, Gu et al. 2024), signaling convergence. PROFET exploits this by discretizing the gradient flow via forward Euler with greedy optimization at each step. While the step size $\Delta t$ is fixed to be small for numerical stability, the time horizon—i.e., the total number of forward Euler steps—adapts to the complexity of the data geometry: more steps for geometrically complex dynamics such as branching or merging, fewer for simpler transitions (see Methods, “Stopping time for GPA”). The resulting variable time horizon is then rescaled to the physical time interval between adjacent snapshots, so the resolution is adaptively determined by the data rather than prescribed a priori. This stands in contrast to DTO methods, where the full computational graph must be retained across integration steps for back-propagation, creating a fundamental trade-off between expressivity and computational cost: increasing the number of steps requires backpropagating through a proportionally deeper graph, forcing these methods to fix the step count a priori, often at a resolution insufficient for complex geometries. PROFET avoids this limitation, faithfully resolving the transport at whatever resolution the geometry demands.

**Learning generalizable velocity fields via force matching.** After GPA simulation, we learn a continuous-time Eulerian velocity field $v_\theta(x, s)$ using force-matching, distilling local potentials $\{\phi_n^{L,*}\}$ into a unified neural vector field. This process enables generalization across time and space, without relying on an analytical form of the dynamics. While the structure of our objective is reminiscent of denoising score matching or flow-matching approaches, a detailed comparison—including its connection and distinction from score-based generative models—is provided in Supplemental Note 3.

#### Supplemental Note 6: Simulation results on TrajectoryNet

*This note is related to the section “Model Validation and Benchmarking Across Four Biological Datasets”.*

TrajectoryNet was excluded from Figure 2C–E because its simulated trajectories diverged substantially from the data support, yielding prediction errors orders of magnitude larger than all other methods. This bias arises from its reliance on continuous normalizing flows (CNFs), which transport particles through an auxiliary Gaussian prior, pulling trajectories away from the true data manifold. Full simulation results for TrajectoryNet are shown in Figure S28 in this Supplemental note.

#### Supplemental Note 7: Ablation studies and robustness analyses

*This note is related to the sections ”Ablation Analysis, Robustness, and Computational Scalability”,“The role of Lipschitz regularization in empirical gradient flow modeling” and “Stopping time for GPA”.*

To further assess the design and robustness of the PROFET framework, we performed a series of ablation studies and sensitivity analyses focusing on the role of the force-matching step, performance under limited data, and dependence on key hyperparameters.

**Ablation of Step 2 (force-matching).** To evaluate the contribution of the force-matching step, we compared trajectories obtained from Step 1 alone (GPA-based gradient flow simulation) with those from the full two-step PROFET framework. Temporal consistency was quantified using the cosine distance between velocity fields evaluated at shared intermediate time points, which measures directional discrepancies in the inferred dynamics.

In the piecewise GPA construction, the endpoint distribution of one segment matches the initial distribution of the subsequent segment at the population level. However, GPA does not provide explicit one-to-one correspondence between individual cells across adjacent segments. To establish cell-level continuity, we constructed correspondences using a $k$-nearest neighbors (KNN) approach at the linking time points. Specifically, for each cell at the end of one segment, we identify its $k$ nearest neighbors in the subsequent segment based on proximity in the latent space. The velocity at the linking point is then estimated by averaging the velocity vectors of these neighboring cells, providing a proxy for the forward-time velocity of each cell across segments.

Using this construction, we compute the cosine distance between the velocity inferred from the first segment and the predicted velocity from the subsequent segment for each cell, and aggregate these discrepancies across all cells. To ensure robustness, we varied $k$ between 3 and 7 and confirmed that the results are consistent across this range.

After neural force-matching, a global continuous-time velocity field is learned, allowing direct evaluation of velocities at the same linking time points without requiring explicit cell matching. This enables a direct comparison of velocity fields before and after force-matching using the same cosine distance metric.

Across datasets, the inclusion of Step 2 leads to a marked reduction in temporal inconsistency. Using a threshold of 0.2 to indicate substantial mismatch, velocity vectors inferred from the full PROFET model consistently fall below this threshold, whereas those obtained from Step 1 alone remain above it. This indicates that GPA captures locally valid transport directions but does not enforce global coherence across time, resulting in misaligned dynamics. The force-matching step resolves this issue by integrating local flows into a unified velocity field that is smooth and temporally consistent (Figure 2G).

**Two-time-point ablation and data efficiency.** We next considered the setting with only two observed time points, where Step 1 alone is sufficient to produce interpolation trajectories. In this regime, we evaluated the effect of Step 2 on robustness and generalization.

To assess data efficiency, we performed uniform subsampling experiments, retaining between 10% and 50% of cells while preserving the underlying distribution. Prediction accuracy was quantified using the $W_2$ distance between predicted and observed distributions at a masked intermediate time point (for the EMT dataset, day 2 was used for evaluation, with days 0 and 4 used for trajectory inference).

In the subsampled setting, GPA (Step 1) learns trajectories only for the selected subset of cells. In contrast, the full PROFET model leverages the second step to learn a global velocity field via force-matching, effectively extending the Lagrangian particle dynamics from Step 1 to an Eulerian representation defined over the entire state space. This enables prediction of trajectories for cells that were not included in the subsampled training set.

As a result, the full PROFET model maintains stable performance under subsampling. For example, the $W_2$ error is 0.96 when using the full dataset and remains comparable (average $\approx 1.02$) across subsampled settings. These results indicate that the force-matching step provides a stabilizing effect by learning a global representation of the dynamics, thereby improving robustness and generalization in data-limited settings (Figure 2H).

**Sensitivity analysis of hyperparameters.** We further examined the dependence of PROFET on key modeling parameters through a grid search.

**GPA stopping criterion and PCA latent dimensionality.** Across both EMT and mESC datasets, distributional errors (measured by $W_2$, Sinkhorn divergence, and MMD) increase gradually with the dimensionality of the latent space (tested at dimensions 2, 4, 8, and 16), consistent with known challenges in high-dimensional distance estimation. Despite this trend, the relative performance of PROFET compared to baseline methods (e.g., WOT and null models) remains stable across all tested dimensions, indicating robustness to the choice of latent representation.

We also evaluated the impact of the GPA stopping criterion, determined based on various divergence thresholds derived from the Lipschitz-regularized $f$-divergence. The resulting trajectories were found to be stable and accurately recover the masked intermediate time points across datasets, with all prediction errors remaining lower than those of baseline methods. These results indicate that the method is not sensitive to the choice of stopping criterion (Figure S7).

**Lipschitz constraint parameter $L$.** The effect of the Lipschitz regularization parameter $L$ was evaluated over the range $L \in [1, 5]$. In the EMT dataset, performance remains stable across this range, consistently yielding the lowest error among all methods. In the mESC dataset, errors increase with larger values of $L$, reflecting increased sensitivity to outliers when the velocity constraint is relaxed (Figure S7).

This behavior is consistent with the role of the Lipschitz constraint in bounding velocity magnitudes and controlling the smoothness of the inferred dynamics. Smaller values enforce smoother and more stable trajectories, while larger values allow sharper transitions that may amplify noise in empirical distributions.

**Summary.** Together, these analyses indicate that the two-step structure of PROFET is essential for achieving temporally consistent dynamics, while also providing robustness to sampling variability and modeling choices. The method remains stable across a wide range of hyperparameters and maintains predictive performance even under substantial data reduction.

#### Supplemental Note 8: Computational Efficiency and Scalability

*This note is related to the section "Ablation Analysis, Robustness, and Computational Scalability".*

To assess the computational cost and scalability of PROFET, we systematically evaluated runtime and memory usage across datasets of varying sizes and dimensionalities. We further compared these metrics against representative baseline methods, including adjoint-based continuous normalizing flow (CNF) models and backpropagation-through-time approaches.

**Computational design of PROFET.** PROFET consists of two stages with distinct computational characteristics:

*Step 1 (GPA: particle-based gradient flow):* This stage performs forward Euler integration, where each step involves solving a dual-potential optimization problem. The computational cost is controlled by two key design choices. First, each Euler step uses only a small number of inner optimization steps (5 iterations), analogous to critic updates in WGAN training, which significantly reduces per-step overhead. Second, the method follows a greedy local optimization, where optimization is performed locally at the particle level without constructing a global computational graph across time. As a result, Step 1 is fully backpropagation-free, avoiding the memory overhead associated with storing intermediate states.

*Step 2 (neural force-matching):* This stage learns a global, time-dependent vector field from precomputed trajectories and is simulation-free. It does not require backpropagation through trajectories or ODE solvers, making it computationally lightweight.

**Runtime benchmarks.** We measured wall-clock runtime across three representative datasets: stem cell differentiation, EMT, and LARRY (3000-cell benchmark).

Across all datasets, PROFET demonstrates competitive runtime. On the largest dataset (LARRY_3000), PROFET completes in approximately 2,743 seconds, compared to TIGON ($\sim$36,388 s; $\sim$13$\times$ slower), TrajectoryNet ($\sim$92,686 s; $\sim$34$\times$ slower), and DeepRUOT ($\sim$5,243 s; $\sim$2$\times$ slower).

Only fully simulation-free methods (e.g., VGFM and MMFM) achieve faster runtimes; however, these methods do not model stochastic dynamics or learn an underlying energy landscape.

Runtime increases moderately with dimensionality (e.g., $\sim$486 s to $\sim$1,085 s for dimensions 2 to 16), indicating that computational cost scales primarily with integration resolution rather than with expensive gradient backpropagation.

**Memory usage.** Peak memory usage remains low and scales favorably across datasets. For the stem cell dataset, memory usage is approximately 0.36 GB and remains nearly constant across dimensions. For the largest dataset, peak memory usage is approximately 3.95 GB.

This is comparable to PRESCIENT and PI-SDE, and significantly lower than MIOFlow ($\sim$6.7 GB) and DeepRUOT ($\sim$7.4 GB). This efficiency arises from the OTD design, which avoids storing intermediate states for backpropagation.

**Scalability analysis.** PROFET scales efficiently with both dataset size and dimensionality. With respect to dataset size, the method exhibits approximately linear scaling with the number of particles. With respect to dimensionality, runtime increases moderately, while memory usage remains largely unaffected.

Unlike adjoint-based methods (e.g., TrajectoryNet and TIGON), which require solving backward ODEs, or discretize-then-optimize (DTO) approaches (e.g., MIOFlow and DeepRUOT), which backpropagate through full trajectories, PROFET performs local updates without maintaining a trajectory-wide computational graph.

**Summary.** These results demonstrate that PROFET achieves a favorable balance between computational efficiency and modeling flexibility. Its particle-based OTD formulation enables scalable learning with low memory overhead, while avoiding the computational bottlenecks associated with backpropagation-heavy or adjoint-based methods.

#### Supplemental Note 9: Velocity growth in KL gradient flow and the role of Lipschitz regularization

*This note is related to the section "The role of Lipschitz regularization in empirical gradient flow modeling".*

While the KL divergence is well-defined for many continuous distributions, its associated gradient flow can yield unbounded velocity fields, making it challenging to simulate numerically. To illustrate, consider two distributions from the exponential family:

$$p(x) = h(x)\exp\left(\langle \theta_p, T(x)\rangle - A(\theta_p)\right), \quad q(x) = h(x)\exp\left(\langle \theta_q, T(x)\rangle - A(\theta_q)\right). \tag{11}$$

with log-density ratio and gradient:

$$\log\frac{p(x)}{q(x)} = \langle \theta_p - \theta_q, T(x)\rangle - \left(A(\theta_p) - A(\theta_q)\right), \quad \nabla_x \log\frac{p(x)}{q(x)} = J_T(x)^\top(\theta_p - \theta_q). \tag{12}$$

When the sufficient statistics $T(x)$ contain polynomial components, the gradient can grow without bound as $|x| \to \infty$. This leads to a velocity field that diverges in magnitude, rendering the corresponding gradient flow ill-posed or numerically unstable.

However, empirical distributions are inherently supported on compact subsets of $\mathbb{R}^d$. Within this bounded support, the gradient field can be well-approximated by a Lipschitz-constrained potential $-\nabla\phi_t^{L,*}$, where the Lipschitz constant $L$ sets an upper bound on velocity magnitude. When $L$ is chosen sufficiently large, the resulting flow closely mimics the unregularized KL-induced dynamics — such as the Ornstein–Uhlenbeck (OU) process.

Figure S29 illustrates this behavior, showing that GPA with a large $L$ (e.g., $L = 10$) can recover the convergence dynamics of the OU process. With smaller $L$ values (e.g. $L = 1$), convergence is slower due to the more restricted transport velocity, and the converged distribution may exhibit slightly larger error relative to the true stationary distribution — reflecting convergence to a proximal solution of the regularized functional rather than to the exact stationary distribution of the unregularized KL flow.

#### Supplemental Note 10: Theoretical guarantee for latent-space modeling

*This note is related to the section "Handling high-dimensionality via latent-space modeling".*

Let $\mathcal{E} : \mathbb{R}^D \to \mathbb{R}^d$ and $\mathcal{D} : \mathbb{R}^d \to \mathbb{R}^D$be Lipschitz encoder and decoder maps, and let $P$ and $Q$ be probability measures in the latent and ambient spaces respectively. The Lipschitz-regularized $f$-divergence satisfies the inequality[10]:

$$D_f^{\mathsf{Lip}_L}(\mathcal{D}_\# P|Q) \le D_f^{a_\mathcal{D}\mathsf{Lip}_L}(P|\mathcal{E}_\# Q), \tag{13}$$

where $a_\mathcal{D}$ is the Lipschitz constant of the decoder.

This inequality implies that the modeling error incurred by transporting distributions in latent space and decoding back to ambient space is controlled. In particular, if the decoder has bounded distortion, the divergence in the ambient space remains upper-bounded by a Lipschitz-scaled divergence in the latent space. This justifies the use of GPA in reduced dimensions, especially in biological applications where the intrinsic data manifold may be significantly lower-dimensional.

#### Supplemental Note 11: PROFET algorithm

*This note is related to the section "Model architecture and training algorithm".*

**Algorithm 1** Particle-based Reconstruction Of generative Force-matched Expression Trajectories (PRO-FET)

**Require:** Time-stamped gene expression snapshots $\{P_t\}_{t=0}^{T}$
**Ensure:** Learned continuous-time velocity field $v_\theta(x, s)$

**Step 1: Trajectory Inference via GPA**

**for** each consecutive pair $(P_{t_i}, P_{t_{i+1}})$ **do**
  Initialize particles $\{Y_0^i\}_{i=1}^{N} \sim P_{t_i}$, $\{X^j\}_{j=1}^{M} \sim P_{t_{i+1}}$
  **for** $n = 0$ to $n_T - 1$ **do**
    Solve dual potential $\phi_n^{L,*} = \arg\max_{\phi \in \mathsf{Lip}_L} \left\{ \frac{1}{N} \sum_i \phi(Y_n^i) - \frac{1}{M} \sum_j f^\star(\phi(X^j)) \right\}$
    Update particles: $Y_{n+1}^i \leftarrow Y_n^i - \Delta t \nabla \phi_n^{L,*}(Y_n^i)$
  **end for**
  Collect supervised samples: $(x, s, v) \leftarrow (Y_n^i,\ n\Delta t,\ -\nabla \phi_n^{L,*}(Y_n^i))$
**end for**

**Step 2: Flow Model Training**

Initialize neural network $v_\theta(x, s)$
**while** not converged **do**
  Sample batch $\{(x_k, s_k, v_k)\}$ from all collected trajectories
  Compute flow matching loss: $\mathcal{L}(\theta) = \frac{1}{K} \sum_k \|v_\theta(x_k, s_k) - v_k\|^2$
  Update $\theta$ via gradient descent
**end while**
**return** $v_\theta(x, s)$
